# Rare missense variants of the leukocyte common antigen related receptor (LAR) display reduced activity in transcellular adhesion and synapse formation

**DOI:** 10.1101/2025.02.16.638491

**Authors:** Mathias Kaas, Nicolas Chofflet, Deniz Bicer, Sune Skeldal, Jinjie Duan, Benjamin Feller, Joachim Vilstrup, Rosa Groth, Suganya Sivagurunathan, Hesam Dashti, Jan Skov Pedersen, Thomas Werge, Anders D. Børglum, Beth A. Cimini, Thouis R. Jones, Melina Claussnitzer, Peder Madsen, Hideto Takahashi, Ditte Demontis, Søren Thirup, Simon Glerup

## Abstract

The leukocyte common antigen related receptor (LAR) is a member of the LAR receptor protein tyrosine phosphatase (RPTP) family of synaptic adhesion molecules that contribute to the proper alignment and specialization of synaptic connections in the mammalian brain. LAR-RPTP members have been genetically associated with neuropsychiatric disorders, but the molecular consequences of genetic perturbations of LAR remain unstudied. Using exome sequencing data from psychiatric patients and controls, we identify rare missense variants of LAR that render the extracellular domain (ECD) unstable and susceptible to proteolytic cleavage. Using recombinant and cellular systems, we describe three variants that cause disruption of the LAR:NGL-3 interaction, which results in loss of transcellular adhesion and synaptogenic effects. Furthermore, we show that overexpression of two of these variants elicit altered morphological phenotypes in an imaging-based morphological profiling assay compared to wild type LAR, suggesting that destabilization of the LAR ECD has broad effects on LAR function. In conclusion, our study identifies three rare, missense variants in LAR that could provide insights into LAR involvement with psychiatric pathobiology.

## Introduction

Synapse formation is a highly complex process partly orchestrated by alignment of pre- and postsynaptic cell-surface adhesion molecules, of which combinatorial interactions contribute to the specification of synaptic subtypes [1-3]. One class of synaptic adhesion molecules is the type IIa receptor protein tyrosine phosphatases constituting the leukocyte common antigen related receptor (LAR), PTPδ and PTPσ, here denoted LAR-RPTPs [4, 5]. These receptors are present at synaptic terminals and axonal surfaces where they engage with postsynaptic adhesion molecules, heparan and chondroitin sulphate proteoglycans (HSPGs and CSPGs), presynaptic receptors and adaptor proteins, to properly guide axons to their targets and ensure transsynaptic tethering and function [6-10]. Like several other synaptic adhesion molecules, LAR-RPTPs have been associated to neurological and psychiatric disorders including autism spectrum disorders, Alzheimer’s Disease, attention deficit hyperactivity disorder (ADHD) and schizophrenia [11-19].

The three LAR-RPTPs share a common protein domain structure with three N-terminal Immunoglobulin-like (Ig-like) domains, up to eight fibronectin III (FN) domains, followed by a transmembrane helix and two intracellular phosphatase domains [4]. Presynaptic LAR-RPTPs interact with several postsynaptic partners across the synapse, including tropomyosin receptor kinase C (TrkC), interleukin 1 receptor accessory protein like 1 (IL1RAPL1) and the interleukin-1 receptor accessory protein (IL-1RAcP), Slit- and Trk-like receptors (Slitrks), synaptic adhesion-like molecules (SALMs) and Netrin-G ligand 3 (NGL-3) [5, 6, 20-28]. While most of these interactions are through the LAR-RPTP Ig-like domains and are regulated by mini-exon A and B (meA/B) insertions [4], interactions with NGL-3 are uniquely mapped to the FN1-2 domains [24]. In this context, LAR-RPTP:NGL-3 interactions induce excitatory synapse connections seemingly without affecting inhibitory synapses, and indeed, the FN1-2 domains of LAR-RPTPs are capable of inducing postsynaptic differentiation by themselves [24, 25].

Recent studies have demonstrated that the intracellular domains (ICDs) of LAR-RPTPs play an important role in their functions. LAR-RPTPs interact with Liprin-α(s) through their C-terminal phosphatase domain (D2), mediating lateral clustering of LAR-RPTP:Liprin-α complexes that are necessary for presynaptic differentiation [29-31]. The interaction of LAR ICDs with additional proteins such as Trio, PDGFR, catenin-β and TrkB have implicated LAR in actin remodeling, a process important for axon guidance during neuronal development [32-34].

Several of the postsynaptic and intracellular proteins interacting with LAR-RPTPs have been associated to psychiatric and neurological disorders, indicating that LAR-RPTPs participate in processes that are important for key pathways in CNS pathologies [14, 35, 36]. Furthermore, transgenic mice with perturbations of the *Ptprf, Ptprd* and *Ptprs* genes, encoding murine LAR, PTPδ and PTPσ, have been studied and show behavioral phenotypes such as hyperactivity, impaired spatial learning and memory, motor deficits in addition to a variety of electrophysiological phenotypes [11, 37-39].

In the present study, we used exome sequencing data from the iPSYCH cohort to map rare, missense variants to our previously reported high-resolution structures of the LAR FN1-4 domains [40]. Using recombinant protein and cellular systems we identified three damaging variants in the LAR FN1-2 domains that render LAR unable to bind its postsynaptic ligand NGL-3 and reduce synaptogenic effects *in vitro*. Using an imaging-based morphological profiling assay, we show that two of these variants elicit different morphological profiles than the WT receptor, mainly causing perturbations of actin morphology. These data indicate that disruption of the LAR ectodomain can perturb several molecular functions of LAR and potentially increase susceptibility of developing psychiatric disorders.

## Results

### LAR missense variant mapping and processing

*PTPRF*, the gene encoding human LAR, has been associated with ADHD and schizophrenia in genome-wide association studies, but the molecular consequences of *PTPRF* perturbations have remained largely unstudied [13-15]. To investigate the effects of LAR/*PTPRF* variation, we evaluated rare missense variants in *PTPRF* (allele count ≤5 in the iPSYCH and non-Finnish Europeans from the nonpsychiatric exome subset of gnomAD [41] cohorts combined) in whole-exome sequencing data from the Danish iPSYCH cohort. These data consist of whole-exome sequences of 9,084 controls and 19,364 individuals diagnosed with at least one of the following psychiatric disorders: ADHD, autism, schizophrenia and affective disorders [42]. After QC, we identified rare missense variants at 164 sites in *PTPRF*, distributed among 183 patients and 66 controls (Fig. 1a). The increased number of missense variants observed in patients was not significant in whole-gene burden analysis (OR=1.3 (0.98 – 1.76), p=0.07, Fisher’s exact test). To evaluate whether there were domain-specific increased burden of missense variants, we supplemented the control data with rare missense variant counts from the gnomAD (non-psychiatric non-Finish European ancestry exomes subset) and found an increased burden of rare missense variants in the LAR FN2 domain in individuals with psychiatric disorders (OR=3 (1.33 – 7.27), p=0.006 Fisher’s exact test, Fig. 1b). We also identified 13 non-rare variants with high individual odds ratios (OR>3, p<0.05 Fisher’s exact test, no multiple testing correction) which were used in the following experimental studies but were not included in the burden tests described above.

**Figure 1:** **a)** Rare missense variants identified in exome-sequencing data of individuals with psychiatric disorders and controls in the iPSYCH cohort. Variants are mapped onto a schematic illustration of the LAR receptor with the Ig1-FN8 domain on top and juxtamembraneous-D2 domain on the bottom. “Control variants” are found in individuals without any psychiatric diagnosis as defined in the iPSYCH cohorts. “Shared variants” refer to missense variants found in both patients and controls. **b)** Domain-wise burden analysis of rare, missense variants of LAR with OR and 95 % confidence interval boxes. FN2 domain shows OR=3, 95 % CI: 1.33 – 7.27, two-sided uncorrected P-value = 0.006 (Fisher exact test). **c)** Western blot analysis of selected missense variants from expression in CHO-K1 cells. Cells were left for 24 hours after transfection before lysis. Western blotting was performed with a LAR ECD specific antibody.

To evaluate potential consequences of missense variation in LAR, we initially screened the effects of missense perturbations in a LAR processing assay. LAR is initially expressed as a single-chain receptor (approximately 200 kDa) and undergoes proteolytic cleavage of the extracellular domain upon exposure to relevant extracellular proteases, rendering E (extracellular, 150 kDa) and P (phosphatase, 85 kDa) subunits [43, 44]. We expressed LAR WT and 164 rare missense variants, as well as the 13 non-rare missense variants, in CHO-K1 cells and evaluated their processing by western blotting (Fig. S1). While a large proportion of the variants did not show any obvious processing changes, some variants showed clear differences in posttranslational processing (Fig. 1c). Here, S51F, P62L and R92W showed a band around 90 kDa not present in the WT. R956L showed a lower molecular weight of the ECD, while Y1356S showed an extra band between the usual one- and two-chain bands. Most noticeably, V389M and P417L showed different molecular weights of the ECD, and several breakdown bands were observed with lower molecular weight, suggesting that the specific amino acid changes in these variants cause destabilization and proteolytic cleavage of the ECD (Fig. 1c).

Missense variants in the FN1-4 domain cause decreased thermal stability and altered NGL-3 binding in solution

We previously reported high-resolution structures of the LAR FN1-4 domains where the V389M and P417L variants are located [40]. Investigating the side chain positions of V389 and P417 on the experimental structures of the FN1-4 and FN1-2 domains showed that the side chains were oriented towards the center of the beta-sandwiches of FN1 and FN2 respectively (Fig. 2a and S2a-b). As side chains that are facing the inside of protein structures are often important for their structural integrity, we investigated the effects of these missense variants on protein stability using recombinant LAR FN1-4. To this end, we cloned a subset of LAR variants into a construct encoding the LAR FN1-4 domain and expressed them in E. coli as previously described [40]. We initially evaluated the stability of these variants using the Tycho NT.6 apparatus to measure the unfolding response. Here, WT LAR FN1-4 showed an inflection temperature (T_i_) of 59.2 °C. Several variants, including V389M and P417L showed a reduced T_i_, indicating that these missense variants indeed cause destabilization of the FN1-4 domain of LAR (Fig. 2b and S2d). We also observed that mutations at P381 (S/L) also caused decreased stability, although the sidechain of P381 is exposed to the hinge region between FN1 and FN2 (Fig. S2c).

**Figure 2:** **a)** Missense variation sites on the experimental structure of LAR FN1-4 (from PDB: 6TPW [40]). Sidechains of V389 and P417 are facing the core of their respective FN domains, while P381 is facing the hinge region between the FN1-2 domains. **b)** Thermal stability of selected variants using Tycho setup. Data are represented as first-derivatives of the fluorescence 330nm/350nm ratio (n=2 pr measurement). **c)** ThermoFluor plots with Boltzmann Sigmoidal fits for selected missense variant. Horizontal dashed line indicates T_i_ (n=3 pr variant, n=3 pr WT batch using three batches for a total of n=9 replicates). **d)** Table showing alteration in melting transition point (ºC change) compared to WT LAR FN1-4 for Tycho and ThermoFluor. **e-k)** MST traces for two replicates of LAR FN1-4 WT and indicated variants. Note that for P381L, V389M and P417L the traces cross during the MST-ON time (after purple vertical bar), indicative of temperature-dependent alterations.

To verify these findings, we customized a fluorescence based thermostability assay to LAR FN1-4 using SYPRO Orange. Here, SYPRO Orange will increase its fluorescent signal when exposed to the hydrophobic cores of proteins whilst they unfold [45]. We screened the unfolding response of FN1-4 missense variants to confirm the data seen from the Tycho assay. This assay revealed a similar pattern of changes in unfolding response of the LAR FN1-4 mutants, although with a lower baseline T_i_ for the WT, but also higher resolution for the changes caused by missense mutations (Fig. 2c-d, S2e). In general, variants in the FN1 domain and the small hinge region between the FN1 and FN2 domains had a stronger effect on thermostability compared to variants in the other three FN domains. Again, the P381L/S, V389M and P417L variants were demonstrated to have marked effects and caused reduction of the thermal stability of the LAR FN1-4 domain. The T_i_ for these variants was significantly different from WT when comparing the V50 values from the Boltzmann sigmoidal fit (p<0.001).

As the FN1-2 domains of LAR are solely responsible for the LAR:NGL-3 interaction [25], we investigated whether missense variants in this region would affect LAR:NGL-3 binding properties. To this end, we set up a microscale thermophoresis (MST) assay to assess LAR FN1-4 binding to the NGL-3 ECD. Measurements of the affinity constant for the WT LAR:NGL-3 interaction were found to have a K_D_ of 3.68 µM (Fig. S2f). When testing the P381L, V389M and P417L variants, we found distinct MST patterns differing from the WT, indicative of a temperature dependent alteration of the LAR:NGL-3 interaction (Fig. 2f,i,k). The P381S variant showed a higher affinity for NGL-3 than LAR WT (Fig. 2g), but as this variant showed NGL-3 dependent differences in initial fluorescence (not shown), the K_D_ should be interpreted with caution. The remaining variants tested did not show notable changes in affinity nor MST trace patterns (Fig. 2h,j, S2g and data not shown).

Taken together, these data suggests that rare missense variants found in the FN1-2 domain of LAR alter the stability of these domains and, for three specific variants, potentially cause disruption of the interaction with its postsynaptic ligand NGL-3.

### LAR variants P381L, V389M and P417L are unable to induce transcellular adhesion

To gain a better understanding of the functional consequences of the variants in the FN1-2 domains of LAR, we evaluated the effects of the identified missense variants on NGL-3 binding in cellular systems. A primary function of LAR-RPTPs is to induce transcellular adhesion by interacting with postsynaptic ligands [4]. This can be assessed in non-neuronal systems, as cells (over)expressing LAR-RPTPs will aggregate with cells expressing their postsynaptic ligands [24, 46]. As the data from recombinant FN1-4 variants in cell-free assays indicated that several variants, including P381L, V389M and P417L, changed the structural stability of the FN1-2 domains known to interact with NGL-3, we explored whether these variants caused perturbations of LAR:NGL-3 transcellular interactions by the cellular aggregation assay [24, 25]. Accordingly, we co-transfected HEK293 cells with LAR variants and mCherry or NGL-3 and GFP and mixed the cells in suspension 24 hours post transfection followed by imaging analysis (Fig. 3a). In agreement with previous studies, WT LAR and NGL-3 readily induced transcellular adhesion, but introduction of variants P381L, V389M or P417L resulted in >50 % mean, significant decrease in cellular aggregation (Fig. 3b-c). We also assessed 19 other variants throughout the LAR protein to determine whether this effect was specific for P381L, V389M and P417L. Although only a subset of missense variants was tested, none showed significant differences from WT LAR (Fig. S3a-b), suggesting that the observed loss of adhesion is specific to P381L, V389M and P417L. Notably, T408M, which was as unstable as V389M in the stability assays, and P381S which had apparent higher affinity for NGL-3 in MST, did not differ from LAR WT in this assay.

**Figure 3:** **a)** Illustration of the transcellular aggregation assay using LAR and NGL-3. Here, separate wells of HEK293T cells are co-transfected with NGL-3 + GFP or LAR + mCherry and subsequently resuspended and mixed in solution and then analyzed with fluorescence microscopy to evaluate amount of aggregated cells **b)** Quantification of cell aggregation index (area) for LAR WT and selected variants as well as IgSF8 (negative control). Bar coloring indicates case-control status (control variants: white, case variants: dark grey). The experiment constitutes at least three biological replicates of n=4 images (12 in total) pr variant. **p=0.0047, ***p=0.0001, ****p<0.0001 (Kruskal-Wallis test). Data is presented as mean ± SD. **c)** Representative images from fluorescent microscopy of cell aggregation solutions as described in a.

### Loss of NGL-3 binding renders LAR variants unable to induce synapse formation

To determine whether the loss of transcellular aggregation was a result of loss of NGL-3 interaction *in vitro*, we tested the binding of full-length LAR variants to NGL-3 at the cell surface. We transfected COS7 cells with WT LAR or P381L, V389M and P417L variants, and subjected the cells to a soluble NGL-3-Fc fusion protein (Fig. 4a). NGL-3-Fc was detected at significantly lower levels at the surface of COS7 cells transfected with variants P381L, V389M and P417L compared to WT control and variants P381S and R388H (Fig. 4b-c). Quantifications were normalized to the surface expression of LAR, where all tested variants showed significantly reduced surface expression (Fig. 4d).

**Figure 4:** **a)** Illustration of *in situ* binding assay using LAR variant transfected HEK293T cells and Fc-NGL-3 fusion protein. Cells were transfected on coverslips and exposed to 2.5 µM soluble Fc-NGL-3 and immunostained (without permeabilization) to visualize the amount of Fc-NGL-3 binding. **b)** Representative fluorescent images of LAR transfected cells with surface-bound NGL-3. **c)** Quantification of surface NGL-3 binding for LAR WT and selected variants, as well as CD4 as negative control binder for NGL-3 (at least three biological replicates and n=49 cells pr replicate. ***p=0.002,****p<0.0001 (Kruskal Wallis test). Data are presented as mean ± SD. **d)** LAR Surface expression quantification from NGL-3 *in situ* binding assay. All variants showed less signal at the cell surface compared to WT. *p=0.0141, ****p<0.0001 (Kruskal Wallis test). **e)** Illustration of artificial synaptogenesis assay using LAR transfected HEK293T cells seeded onto rat hippocampal neurons. The cells were co-cultured for 24 hours before PSD-95 assessment. **f)** Representative images of LAR transfected HEK293T cells in co-culture with hippocampal neurons stained for PSD-95 clustering. **g)** Quantification of PSD-95 clusters as markers of excitatory synapse formation onto LAR transfected HEK293T cells, using CD4 as a negative control. Data consists of three biological replicates with n>10 images pr replicate. **p<0.005, ****p<0.001 (Kruskal Wallis test). Data are presented as mean ± SD. Bar coloring in c), d) and g) indicates case-control status (control variants: white, case variants: dark grey)

As LAR is capable of inducing excitatory postsynaptic differentiation through NGL-3 and other postsynaptic ligands [24, 25, 47], we explored whether the loss of transcellular adhesion and NGL-3 interaction shown by the P381L, V389M and P417L variants translated to loss of postsynaptic induction in an artificial synapse formation assay. Here, we transfected HEK293 cells with LAR WT or missense variants and seeded them onto DIV14 rat hippocampal neurons. This mixed culture was PFA fixed 24 hours after HEK293 cell seeding and immunostained for PSD-95 (Fig. 4e). HEK293 cells expressing LAR WT readily induced postsynaptic differentiation from the co-cultured neurons as demonstrated by increased PSD-95 signal in proximity to these cells, which was not observed for the negative controls transfected with CD4 (Fig. 4f-g). In line with the loss of transcellular adhesion, the P381L, V389M and P417L missense variants displayed a significant decrease in postsynaptic induction, while variants P381S and R388H showed no difference in accumulation of PSD-95 clustering when compared to WT LAR (Fig. 4f-g). In this experiments, we selected HEK293 cells with similar surface expression of each LAR construct in order to assess the effect of the reduction of LAR:NGL3 trans-interaction and not LAR surface expression itself (Fig. S4a).

Since the full-length LAR constructs used so far do not contain meA or meB inserts, it is unlikely that they could induce postsynaptic differentiation through their Ig-like domains, since the majority of known postsynaptic adhesion partners through these domains depend on combinatorial meA/B insertions [22, 48]. Thus, directly interpreting whether the loss of synaptogenic effects, seen with P381L, V389M and P417L, is caused by loss of NGL-3 binding or FN1-2 domain destabilization (and potential breakdown of the N-terminal part of the ECD) is difficult. To evaluate if these variants retained functional integrity of the N-terminal Ig-like domains, we tested their ability to bind postsynaptic partners through these domains when expressing the meA and meB insertions. In a similar cell surface binding assay as performed above, we tested LAR^meA+/meB+^ variants binding to SALM5-Fc [26]. Here, P417L was the only variant that showed decreased binding to SALM5-Fc (Fig. S4b+c). Furthermore, P417L was also the only variant that showed lower surface exposure among the variants (Fig. S4d).

### The LAR FN1-4 domains are disrupted by variants P381L, V389M and P417L

As cellular processing of full-length LAR, as well as thermal stability and MST analysis of the (recombinant) FN1-4 domains indicated destabilization by the P381L, V389M and P417L variants, we speculated whether these variants caused unfolding of the FN1-2 domains which then in turn affected the interaction with NGL-3. To test if the P381L, V389M and P417L variants had generalized effects on the structure of the FN1-2 domains, we used small-angle x-ray scattering (SAXS) analysis for the recombinant FN1-4 variants, as we had previously used this method to determine solution structures of the WT LAR FN1-4 domain [40]. The P381L, V389M and P417 variants all showed large differences in scattering profiles (log(I(q)) versus log(q)) and pair-distance distribution functions (PDDF, a histogram of distances between pair of points weighted by the excess scattering length in the points) compared to WT LAR FN1-4, indicating altered protein structure where the PDDF suggests asymmetric and/or elongated structures of the variants (Fig. 5a, S5a). Moreover, dimensionless Kratky analysis showed that variant P381L was highly flexible and probably denature easily (Fig. 5b). Guinier fit analyses are shown in Fig S5b-e and other SAXS statistics in figure 5c.

**Figure 5:** **a)** Pair distance distribution function (indirect Fourier transformation [IFT] of SAXS data) of LAR variants for analysis of distance distribution histogram to estimate the shape of protein, whoing elongated plots for the three tested variants indicating elongated protein shapes. **b)** Dimensionless Kratky plot based on Radius of gyration (Rg) of LAR variants for compactness and folding state analysis.**c)** Structural parameters of LAR rare variants calculated from SAXS data based on Guinier analysis and IFT. I (0), forward scattering intensity, MW, molecular weight estimate, Rg(Å) Radius of gyration, and Dmax(Å), maximum particle dimension. **d)** Structural modelling of LAR FN 1-4 and NGL-3 interaction using AlphaFold2 and **e)** Molecular docking program HDOCK.

As no experimental structures are available for the LAR:NGL-3 complex, we used AlphaFold2 [49] and molecular docking simulations to model the LAR FN1-4:NGL-3 interaction to estimate the position of P381, V389 and P417 in relation to NGL-3. Here, both AlphaFold2 (available through ColabFold [50]) and HDOCK simulations [51], using the sequences or published (individual) structures of LAR FN1-4 domains and NGL-3 ECD domains respectively, suggested that the FN1-2 domains of LAR interacts with the concave surface of the NGL-3 LRR domain (Fig. 5d-e, S6a-b). The interaction involved beta-sheet surfaces of FN1 as well as the FN1-2 hinge region and N-terminal FN2 loops (Fig. 5d-e and S6a-b). AlphaFold and HDOCK, however, predicted opposite beta-sandwich surfaces of the FN1 domain to be involved. In the AlphaFold model, the P381L variant was seemingly in the interaction surface of LAR:NGL-3, whereas the V389M and P417L variants were placed too far from NGL-3 to be directly involved in the interaction (Fig. 5d). Neither of the variants were found to be directly involved in the LAR:NGL-3 interaction in the HDOCK model (Fig. 5e). Both models were high-confidence predictions as illustrated by PAE plots for AlphaFold (Fig. S6c) and HDOCK docking score of -295.67 and confidence score 0.9485.

Taken together, SAXS and computational modeling suggest that P381L, V389M and P417L potentially cause unfolding of the FN1-2 domains to limit interactions with NGL-3.

### Imaging-based morphological profiling illustrates broad effects of damaging LAR missense variants

LAR-RPTPs are involved in several cellular processes other than synapse formation including neurite outgrowth, dephosphorylation of intracellular targets, actin remodeling, filopodia formation and cell migration [32, 33, 52, 53]. Thus, damaging LAR missense variation might have effects beyond those evaluated in classical adhesion and synaptogenic assays. In an attempt to evaluate whether LAR missense variation could have broader molecular effects, we adapted the recently published LipocyteProfiler assay (itself an adaption of the Cell Painting assay [54, 55]), that utilizes high-content confocal imaging and computational feature extraction to capture thousands of morphological features from single cells and systematically interrogate genetic or chemical perturbations [56, 57]. We transfected U2OS cells with LAR WT or the P381S/L, R388H, V389M, P416S and P417L variants, and after 24 hours we performed an adapted Lipocyte Profiler staining protocol, using MitoTracker, Wheat Germ Agglutinin (WGA), Phalloidin, Hoechst and an α-LAR antibody, the latter replacing the SYTO-14 and BODIPY stains. Using automated confocal imaging, we captured images of U2OS cells and performed a single cell analysis pipeline that identified transfected cells based on the α-LAR signal and extracted ∼3000 features per cell across five imaging channels. Using pr well mean readouts for each feature, we calculated per-feature Z-scores and p-values for each LAR variant compared to WT (Fig. 6a-b).

**Figure 6:** **a)** Graphical illustration of adapted LP assay to evaluate morphological features elicited by LAR WT or variant overexpression in U2OS cells. Cells were multi-stained with Hoechst, MitoTracker, Phalloidin, WGA and anti-LAR in 384 well plates. Images were acquired with automated, high-content confocal microscopy and feature profiles were extracted for single cells and aggregated for each well before undergoing customized analysis. **b)** Representative images of the four channels from WT, P381L, V389M and P417L illustrating staining of specific compartments and identification of LAR-positive cells. **c-e)** Volcano plots illustrating changed feature types between WT LAR and P381L, V389M and P417L respectively. Dot size is scaled dependent on the p-value as illustrated. Data is from n=5-6 wells with a minimum of 20 LAR-positive cells pr well. p-value from t-test. X-axis are clipped at -1.5 and 1.5.

When comparing WT LAR with the variants P381L, V389M or P417L, there was a clear change in morphological feature characteristics for V389M and P417L (Fig. 6c-e). Most notably different from LAR WT was P417L with 329 features significantly changed from the WT, while V389M had 112 features and P381L had only 13 significant changes (p<0.001, t-test, Fig. 6c-e). Common for V389M and P417L comparisons with WT were a high proportion of Actin-Golgi-Plasma membrane (AGP) features altered between the groups, of which most were related to cytoplasmic texture features (Supplementary Table 1). Analysis of LAR WT against P381S, R388H and P416S showed only 4, 5 and 1 features significantly changed, suggesting that these variants are generally well tolerated and do not induce large changes to LAR function (Fig. S6a-c). Overall, the imaging-based profiling suggested that the V389M and P417L LAR variants perturb functions of LAR not specific for synapse formation.

## Discussion

In the present study, we used rare missense variants identified in individuals diagnosed with psychiatric disorders and controls to investigate the biological effects that might underlie the genetic association of *PTPRF*/LAR with psychiatric disorders. We identified three rare missense variants in the LAR FN1-2 domains that caused loss of synaptogenic and adhesion properties of LAR. Among these variants, P381L was only found in controls, while V389M and P417L, that also elicited more general perturbations of LAR function, were found in patients only.

Studies of full-length LAR processing, SAXS and the adapted LP assay suggest that destabilization of the FN1-2 domains by V389M and P417L may result in LAR instability and/or accelerated turnover. This could result in increased loss of (fragments of) the LAR ECD, which in turn could also cause general loss of synaptic adhesion through the LAR extracellular domain. Cell surface shedding of synaptic adhesion molecules has been described as a key regulatory mechanism for their activity and function [58], and soluble LAR-RPTP ectodomains have been shown to regulate neurite growth and response to nerve injury [59-63]. Furthermore, the excess soluble LAR ectodomain fragments at synaptic terminals could act as decoy receptors for postsynaptic LAR-RPTP ligands to prevent transsynaptic complex formation, as proposed for other synaptic adhesion molecules [58]. During development, excess soluble LAR ectodomain could act as guidance cues for neighboring neurons during development while the LAR expressing neurons would lack membrane anchored LAR to participate in axon guidance, providing potential mechanisms by which the identified variants could participate in psychiatric pathogenesis.

The loss of synaptogenic potential of the LAR P381L, V389M and P417L variants shown in this study is plausibly explained by loss of NGL-3 binding, since the same variants expressed in a LAR construct containing meA and B inserts could bind the postsynaptic ligand SALM5. Thus, LAR could potentially still induce synapse formation, through its Ig-like domains, in the absence of NGL-3 binding. MeA and B insertions in the Ig-like domains are important for the majority of interactions of the Ig-like domains of LAR-RPTPs, and LAR-RPTPs devoid of both these inserts generally have low affinity for postsynaptic ligands interacting with their Ig-like domains [5]. A recent study has thoroughly characterized the presence of mini-exon inserts in LAR-RPTPs throughout several brain regions, neuronal subtypes and projections in mice [48]. Specifically, they showed that the majority of excitatory neurons in the cortex and thalamus are devoid of the meA and B inserts, suggesting the importance of LAR:NGL-3 binding in specific circuits. Thus, the NGL-3 specific synaptic perturbations caused by the P381L, V389M and P417L LAR variants could be highly context or spatially dependent *in vivo*.

Lateral clustering of LAR-RPTPs is critical for synaptogenesis and actin remodeling and is mediated by both intracellular adaptor molecules, such as Liprin-αs, and extracellular ligands [30, 33, 64, 65]. The LAR D1 domain has phosphatase activity which is inhibited after Liprin-α induced clustering, suggesting that LAR-RPTP signaling is regulated by receptor dimerization [31]. In addition, the LAR-RPTP ICD undergoes redistribution after shedding of the ectodomain, indicating that release of the ECD could have consequences for downstream intracellular signaling of LAR [66]. While we did not investigate the distribution or processing of the LAR ICD in the present work, the changed morphological features elicited by expression of the V389M and P417L variants could be a manifestation of altered LAR clustering and/or intracellular signaling caused by destabilization of the LAR ectodomain. Additionally, the ECDs of LAR-RPTPs are well described to interact with extracellular matrix proteins such as laminin and syndecans and other HSPG/CSPGs to modulate focal adhesions and actin morphology [53, 67-70], and distortion of such extracellular interactions through ectodomain destabilization could in turn perturb LAR clustering and downstream signaling. As such, the altered AGP morphology observed using imaging-based morphological profiling could be caused by dysfunctional regulation of the LAR ICD due to ectodomain destabilization. Thus, the LAR V389M and P417L variants could potentially contribute to psychiatric pathobiology through several mechanisms *in vivo*, in line with the observed effects in the adapted LP assay. Indeed, unbiased morphological phenotyping represents a more powerful approach to evaluate the effects of a large number of missense variants, and can potentially capture both known and unknown functions of the proteins and/or variants in question [71].

The three highlighted variants are rare, and in the present study we lacked statistical power to argue any genetic association with psychiatric disorders, and indeed, P381L was only found in controls. The understanding of psychiatric genetics has improved over the last decade due to large GWAS and rare variant studies and have demonstrated that psychiatric disorders have a highly heterogenous genetic makeup with contributions from thousands of both rare and common variants [14, 15, 72]. The presence of damaging, rare variants on top of a high polygenic risk score for psychiatric disorders contribute significantly to psychiatric risk, but by themselves, these variants are not necessarily causing psychiatric disorders [14, 73, 74]. In addition, data from gnomAD suggests that *PTPRF* is evolutionarily constrained as indicated by LOEUF and missense Z scores of 0.35 and 4.05 respectively [75], and as such, the damaging phenotypes elicited by P381L, V389M and P417L could be due to general missense intolerance of *PTPRF* rather than specific association with psychiatric disorders. However, the findings in the present study argue for further attention to missense variation in LAR-RPTPs and their potential association with psychiatric disorders.

Taken together, we present the identification of three rare missense variants in the LAR FN1-2 domains that perturb transcellular adhesion, synapse formation and broader LAR functions *in vitro* and propose that such damaging missense variation in LAR could contribute to psychiatric pathobiology.

## Supporting information

Illum correction code

Main figures

Analysis code

## Author contributions

Study design: MK, DD, SSk, JPV, SG, PM. Design of experiments: MK, SSk, JPV, NC, BF, MC, SG, HT, ST. Protein production: PM, JPV, RG. Performance of experiments: MK, SSk, NC, BF, RG, DB. Data analysis and interpretation: MK, DD, JD, SSk, JJ, JPV, SG, PM, NC, BF, HD, JSP, ST, DB, SSi, RJ, BC. Resources: MK, SG, PM, DD, ADB, ST, HT, MC, JSP, BC. Supervision of genetic analyses: DD, ADB, TW. Writing: MK, SG.

## Acknowledgements

This study was funded by the Lundbeck Foundation (grant number R263-2017-3678 (MK)) and the Novo Nordisk Foundation (grant numbers NNF21SA0072102, NNF21OC0072675 (MK)) and supported by the Canadian Institutes of Health Research grants (MOP-133517, PTJ-191947), National Institutes of Health (NIH) (NIMH R01MH077303) and Fonds de la Recherche du Québec Research Scholars (FRQS Junior 2 (29106) and senior (251655)) grants to H.T., and the FRQS doctoral award (353400) and the IRCM doctoral scholarship to N.C. The iPSYCH team was supported by grants from the Lundbeck Foundation (R102-A9118, R155-2014-1724, and R248-2017-2003), NIH/NIMH (1R01MH124851-01 to A.D.B.). High-performance computer capacity for handling and statistical analysis of iPSYCH data on the GenomeDK HPC facility was provided by the Center for Genomics and Personalized Medicine and the Centre for Integrative Sequencing, iSEQ, Aarhus University, Denmark (grant to A.D.B.). Mitra Shamsali is thanked for excellent technical assistance.

## Figure legends

**Supplemental Figure 1:** Western blots of all rare missense variants identified in the iPSYCH cohort in transfected CHO-K1 cells. Blots were developed with a polyclonal antibody against the extracellular domain of LAR. Media were also analyzed for receptor shedding. A GAPDH antibody was used to show that cell viability was comparable between transfected cells. Of note, P416S was analyzed with a new plasmid prep and looked like WT LAR (data not shown). GAPDH indicated with asterisks are from a different blot with the same samples, but the mock and WT lanes were switched. The last blot illustrates the described non-rare variants identified.

**Supplemental Figure 2: a-c)** Side chain positions for V389 (a), P417L (b) and P381 (c) are indicated in red with intermolecular sidechains within 4 Å shown as grey sticks (from experimental structure 6TPV of LAR FN1-2 [40]). **d)** Thermostability assessment using Tycho for remaining tested variants in this assay (n=2 pr variant) **e)** Fluorescent changes from SYPRO Orange ThermoFluor assay showing increases in normalized SYPRO Orange fluorescence during protein unfolding for remaining tested variants (n=3 pr variant, n=3 pr batch WT, 9 replicates in total). **f)** LAR WT vs. NGL-3 with observed changes in F-norm and K_D_ assessment. **g)** Table showing changes in K_D_ observed for the different variants.

**Supplemental Figure 3: a)** Representative images of the remaining variants tested in transcellular adhesion assay. **b)** Quantification of remaining variants from the assay. No variants were significantly different (Kruskal Wallis test). Bar coloring indicates case-control status (control variants: white, shared variants: light grey, case variants: dark grey).

**Supplemental Figure 4: a)** Quantitative assessment of LAR variant surface expression in HEK293 cells in the artificial synapse formation assay. Of note, cells were selected based on their similar expression for uniform PSD-95 analysis. None of the tested variants had significantly different levels at the cell surface in this assay (Kruskal Wallis test). Data are presented as mean ± SD. **b)** Representative images from SALM5-Fc surface binding to LAR meA+meB variants in transfected HEK293 cells. The scale bar represents 30 μm. **c)** Quantification of SALM5-Fc surface binding to LAR meA+meB variants in transfected HEK293 cells. Among the missense variants, only P417L showed significantly decreased SALM5 binding (p=0.0007, Kruskal Wallis test). Data are presented as mean ± SD. **d)** Surface expression of the same LAR variants as in c). Only P417L showed significant changes from WT (p=0.0052, Kruskal Wallis test). Data are presented as mean ± SD. Bar coloring indicates case-control status (control variants: white, case variants: dark grey).

**Supplemental Figure 5: a)** SAXS data plotted at log-log scale of LAR wild-type and rare variants. Y-axis depicts the scattering intensity, I(q), and x-axis shows the momentum transfer vector, q. **b-e)** SAXS data points that are used for Guinier analysis and their normalized residual plots. Points that are on the red lines are used to calculate Rg (Å). Very low angle scattering points are neglected due to parasitic scattering and aggregation artefacts.

**Supplemental Figure 6: a-b)** Additional model structure illustrations from AlphaFold2 and HDOCK modelling of the LAR FN1-4:NGL-3 interface. **c)** Predicted aligned error (PAE) plots of the three best AlphaFold2 models demonstrating high confidence in the interposition of the LRR domain of NGL-3 and the FN1 domain of LAR. The color code indicates the expected distance error in Ångströms (Å).

**Supplemental Figure 7: a-c)** Pairwise comparison of LAR WT and missense variants as indicated. Dot size is scaled dependent on the p-value as illustrated. Data is from n=5-6 wells with a minimum of 20 LAR-positive cells pr well.

**Supplementary Table 1:** List of significantly altered features for the different variants tested in the adapted LP assay.

