## Supplementary material for "Rare missense variants of the leukocyte common antigen related receptor (LAR) display reduced activity in transcellular adhesion and synapse formation": Illum correction code

```
CellProfiler Pipeline: http://www.cellprofiler.org
Version:5
DateRevision:424
GitHash:
ModuleCount:21
HasImagePlaneDetails:False

LoadData:[module_num:1|svn_version:'Unknown'|variable_revision_number:6|show_window:False|notes:
[]|batch_state:array([], dtype=uint8)|enabled:True|wants_pause:False]
    Input data file location:Default Input Folder sub-folder|Desktop\CellReceptormutants
    Name of the file:load_data.csv
    Load images based on this data?:Yes
    Base image location:None|
    Process just a range of rows?:No
    Rows to process:1,100000
    Group images by metadata?:Yes
    Select metadata tags for grouping:Plate
    Rescale intensities?:Yes

Resize:[module_num:2|svn_version:'Unknown'|variable_revision_number:5|show_window:False|notes:
[]|batch_state:array([], dtype=uint8)|enabled:True|wants_pause:False]
    Select the input image:OrigDNA
    Name the output image:DownsampledDNA
    Resizing method:Resize by a fraction or multiple of the original size
    X Resizing factor:0.1
    Y Resizing factor:0.1
    Z Resizing factor:1.0
    Width (x) of the final image:100
    Height (y) of the final image:100
    # of planes (z) in the final image:10
    Interpolation method:Bilinear
    Method to specify the dimensions:Manual
    Select the image with the desired dimensions:None
    Additional image count:0

CorrectIlluminationCalculate:
[module_num:3|svn_version:'Unknown'|variable_revision_number:2|show_window:False|notes:[]|batch_state:array([],
dtype=uint8)|enabled:True|wants_pause:False]
    Select the input image:DownsampledDNA
    Name the output image:IllumDNA
    Select how the illumination function is calculated:Regular
    Dilate objects in the final averaged image?:No
    Dilation radius:1
    Block size:60
    Rescale the illumination function?:Yes
    Calculate function for each image individually, or based on all images?:All: Across cycles
    Smoothing method:Median Filter
    Method to calculate smoothing filter size:Manually
    Approximate object diameter:10
    Smoothing filter size:40
    Retain the averaged image?:No
    Name the averaged image:IllumBlueAvg
    Retain the dilated image?:No
```

Name the dilated image:IllumBlueDilated
Automatically calculate spline parameters?:Yes
Background mode:auto
Number of spline points:5
Background threshold:2
Image resampling factor:2
Maximum number of iterations:40
Residual value for convergence:0.001

Resize:[module_num:4|svn_version:'Unknown'|variable_revision_number:5|show_window:False|notes:[]|batch_state:array([], dtype=uint8)|enabled:True|wants_pause:False]
Select the input image:IllumDNA
Name the output image:UpsampledIllumDNA
Resizing method:Resize by a fraction or multiple of the original size
X Resizing factor:10
Y Resizing factor:10
Z Resizing factor:1.0
Width (x) of the final image:100
Height (y) of the final image:100
### of planes (z) in the final image:10
Interpolation method:Bilinear
Method to specify the dimensions:Manual
Select the image with the desired dimensions:None
Additional image count:0

Resize:[module_num:5|svn_version:'Unknown'|variable_revision_number:5|show_window:False|notes:[]|batch_state:array([], dtype=uint8)|enabled:True|wants_pause:False]
Select the input image:OrigAGP
Name the output image:DownsampledAGP
Resizing method:Resize by a fraction or multiple of the original size
X Resizing factor:0.1
Y Resizing factor:0.1
Z Resizing factor:1.0
Width (x) of the final image:100
Height (y) of the final image:100
### of planes (z) in the final image:10
Interpolation method:Bilinear
Method to specify the dimensions:Manual
Select the image with the desired dimensions:None
Additional image count:0

CorrectIlluminationCalculate:[module_num:6|svn_version:'Unknown'|variable_revision_number:2|show_window:False|notes:[]|batch_state:array([], dtype=uint8)|enabled:True|wants_pause:False]
Select the input image:DownsampledAGP
Name the output image:IllumAGP
Select how the illumination function is calculated:Regular
Dilate objects in the final averaged image?:No
Dilation radius:1
Block size:60
Rescale the illumination function?:Yes
Calculate function for each image individually, or based on all images?:All: Across cycles
Smoothing method:Median Filter
Method to calculate smoothing filter size:Manually

Approximate object diameter:10
Smoothing filter size:40
Retain the averaged image?:No
Name the averaged image:IllumBlueAvg
Retain the dilated image?:No
Name the dilated image:IllumBlueDilated
Automatically calculate spline parameters?:Yes
Background mode:auto
Number of spline points:5
Background threshold:2
Image resampling factor:2
Maximum number of iterations:40
Residual value for convergence:0.001

Resize:[module_num:7|svn_version:'Unknown'|variable_revision_number:5|show_window:False|notes:[]|batch_state:array([], dtype=uint8)|enabled:True|wants_pause:False]
Select the input image:IllumAGP
Name the output image:UpsampledIllumAGP
Resizing method:Resize by a fraction or multiple of the original size
X Resizing factor:10
Y Resizing factor:10
Z Resizing factor:1.0
Width (x) of the final image:100
Height (y) of the final image:100
### of planes (z) in the final image:10
Interpolation method:Bilinear
Method to specify the dimensions:Manual
Select the image with the desired dimensions:None
Additional image count:0

Resize:[module_num:8|svn_version:'Unknown'|variable_revision_number:5|show_window:False|notes:[]|batch_state:array([], dtype=uint8)|enabled:True|wants_pause:False]
Select the input image:OrigMito
Name the output image:DownsampledMito
Resizing method:Resize by a fraction or multiple of the original size
X Resizing factor:0.1
Y Resizing factor:0.1
Z Resizing factor:1.0
Width (x) of the final image:100
Height (y) of the final image:100
### of planes (z) in the final image:10
Interpolation method:Bilinear
Method to specify the dimensions:Manual
Select the image with the desired dimensions:None
Additional image count:0

CorrectIlluminationCalculate:[module_num:9|svn_version:'Unknown'|variable_revision_number:2|show_window:False|notes:[]|batch_state:array([], dtype=uint8)|enabled:True|wants_pause:False]
Select the input image:DownsampledMito
Name the output image:IllumMito
Select how the illumination function is calculated:Regular
Dilate objects in the final averaged image?:No
Dilation radius:1

Block size:60
Rescale the illumination function?:Yes
Calculate function for each image individually, or based on all images?:All: Across cycles
Smoothing method:Median Filter
Method to calculate smoothing filter size:Manually
Approximate object diameter:10
Smoothing filter size:40
Retain the averaged image?:No
Name the averaged image:IllumBlueAvg
Retain the dilated image?:No
Name the dilated image:IllumBlueDilated
Automatically calculate spline parameters?:Yes
Background mode:auto
Number of spline points:5
Background threshold:2
Image resampling factor:2
Maximum number of iterations:40
Residual value for convergence:0.001

Resize:[module_num:10|svn_version:'Unknown'|variable_revision_number:5|show_window:False|notes:[]|batch_state:array([], dtype=uint8)|enabled:True|wants_pause:False]
Select the input image:IllumMito
Name the output image:UpsampledIllumMito
Resizing method:Resize by a fraction or multiple of the original size
X Resizing factor:10
Y Resizing factor:10
Z Resizing factor:1.0
Width (x) of the final image:100
Height (y) of the final image:100
### of planes (z) in the final image:10
Interpolation method:Bilinear
Method to specify the dimensions:Manual
Select the image with the desired dimensions:None
Additional image count:0

Resize:[module_num:11|svn_version:'Unknown'|variable_revision_number:5|show_window:False|notes:[]|batch_state:array([], dtype=uint8)|enabled:True|wants_pause:False]
Select the input image:OrigBrightfield
Name the output image:DownsampledBrightfield
Resizing method:Resize by a fraction or multiple of the original size
X Resizing factor:0.1
Y Resizing factor:0.1
Z Resizing factor:1.0
Width (x) of the final image:100
Height (y) of the final image:100
### of planes (z) in the final image:10
Interpolation method:Bilinear
Method to specify the dimensions:Manual
Select the image with the desired dimensions:None
Additional image count:0

CorrectIlluminationCalculate:
[module_num:12|svn_version:'Unknown'|variable_revision_number:2|show_window:False|notes:[]|batch_state:array([], dtype=uint8)|enabled:True|wants_pause:False]

    Select the input image:DownsampledBrightfield
    Name the output image:IllumBrightfield
    Select how the illumination function is calculated:Regular
    Dilate objects in the final averaged image?:No
    Dilation radius:1
    Block size:60
    Rescale the illumination function?:Yes
    Calculate function for each image individually, or based on all images?:All: Across cycles
    Smoothing method:Median Filter
    Method to calculate smoothing filter size:Manually
    Approximate object diameter:10
    Smoothing filter size:40
    Retain the averaged image?:No
    Name the averaged image:IllumBlueAvg
    Retain the dilated image?:No
    Name the dilated image:IllumBlueDilated
    Automatically calculate spline parameters?:Yes
    Background mode:auto
    Number of spline points:5
    Background threshold:2
    Image resampling factor:2
    Maximum number of iterations:40
    Residual value for convergence:0.001

Resize:[module_num:13|svn_version:'Unknown'|variable_revision_number:5|show_window:False|notes:
[]|batch_state:array([], dtype=uint8)|enabled:True|wants_pause:False]
    Select the input image:IllumBrightfield
    Name the output image:UpsampledIllumBrightfield
    Resizing method:Resize by a fraction or multiple of the original size
    X Resizing factor:10
    Y Resizing factor:10
    Z Resizing factor:1.0
    Width (x) of the final image:100
    Height (y) of the final image:100
    # of planes (z) in the final image:10
    Interpolation method:Bilinear
    Method to specify the dimensions:Manual
    Select the image with the desired dimensions:None
    Additional image count:0

Resize:[module_num:14|svn_version:'Unknown'|variable_revision_number:5|show_window:False|notes:
[]|batch_state:array([], dtype=uint8)|enabled:True|wants_pause:False]
    Select the input image:OrigMarker
    Name the output image:DownsampledMarker
    Resizing method:Resize by a fraction or multiple of the original size
    X Resizing factor:0.1
    Y Resizing factor:0.1
    Z Resizing factor:1.0
    Width (x) of the final image:100
    Height (y) of the final image:100
    # of planes (z) in the final image:10
    Interpolation method:Bilinear
    Method to specify the dimensions:Manual
    Select the image with the desired dimensions:None

Additional image count:0

CorrectIlluminationCalculate:
[module_num:15|svn_version:'Unknown'|variable_revision_number:2|show_window:False|notes:[]|batch_state:array([], dtype=uint8)|enabled:True|wants_pause:False]
Select the input image:DownsampledMarker
Name the output image:IllumMarker
Select how the illumination function is calculated:Regular
Dilate objects in the final averaged image?:No
Dilation radius:1
Block size:60
Rescale the illumination function?:Yes
Calculate function for each image individually, or based on all images?:All: Across cycles
Smoothing method:Median Filter
Method to calculate smoothing filter size:Manually
Approximate object diameter:10
Smoothing filter size:40
Retain the averaged image?:No
Name the averaged image:IllumBlueAvg
Retain the dilated image?:No
Name the dilated image:IllumBlueDilated
Automatically calculate spline parameters?:Yes
Background mode:auto
Number of spline points:5
Background threshold:2
Image resampling factor:2
Maximum number of iterations:40
Residual value for convergence:0.001

Resize:[module_num:16|svn_version:'Unknown'|variable_revision_number:5|show_window:False|notes:[]|batch_state:array([], dtype=uint8)|enabled:True|wants_pause:False]
Select the input image:IllumMarker
Name the output image:UpsampledIllumMarker
Resizing method:Resize by a fraction or multiple of the original size
X Resizing factor:10
Y Resizing factor:10
Z Resizing factor:1.0
Width (x) of the final image:100
Height (y) of the final image:100
### of planes (z) in the final image:10
Interpolation method:Bilinear
Method to specify the dimensions:Manual
Select the image with the desired dimensions:None
Additional image count:0

SaveImages:[module_num:17|svn_version:'Unknown'|variable_revision_number:16|show_window:False|notes:[]|batch_state:array([], dtype=uint8)|enabled:True|wants_pause:False]
Select the type of image to save:Image
Select the image to save:UpsampledIllumDNA
Select method for constructing file names:Single name
Select image name for file prefix:None
Enter single file name:\g<Plate>_IllumDNA
Number of digits:4
Append a suffix to the image file name?:No

Text to append to the image name:
Saved file format:npy
Output file location:Default Output Folder|
Image bit depth:8-bit integer
Overwrite existing files without warning?:No
When to save:Last cycle
Record the file and path information to the saved image?:No
Create subfolders in the output folder?:No
Base image folder:Elsewhere...|
How to save the series:T (Time)
Save with lossless compression?:Yes

SaveImages:[module_num:18|svn_version:'Unknown'|variable_revision_number:16|show_window:False|notes:[]|batch_state:array([], dtype=uint8)|enabled:True|wants_pause:False]
Select the type of image to save:Image
Select the image to save:UpsampledIllumAGP
Select method for constructing file names:Single name
Select image name for file prefix:None
Enter single file name:\g<Plate>_IllumAGP
Number of digits:4
Append a suffix to the image file name?:No
Text to append to the image name:
Saved file format:npy
Output file location:Default Output Folder|
Image bit depth:8-bit integer
Overwrite existing files without warning?:No
When to save:Last cycle
Record the file and path information to the saved image?:No
Create subfolders in the output folder?:No
Base image folder:Elsewhere...|
How to save the series:T (Time)
Save with lossless compression?:Yes

SaveImages:[module_num:19|svn_version:'Unknown'|variable_revision_number:16|show_window:False|notes:[]|batch_state:array([], dtype=uint8)|enabled:True|wants_pause:False]
Select the type of image to save:Image
Select the image to save:UpsampledIllumMito
Select method for constructing file names:Single name
Select image name for file prefix:None
Enter single file name:\g<Plate>_IllumMito
Number of digits:4
Append a suffix to the image file name?:No
Text to append to the image name:
Saved file format:npy
Output file location:Default Output Folder|
Image bit depth:8-bit integer
Overwrite existing files without warning?:No
When to save:Last cycle
Record the file and path information to the saved image?:No
Create subfolders in the output folder?:No
Base image folder:Elsewhere...|
How to save the series:T (Time)
Save with lossless compression?:Yes

```
SaveImages:[module_num:20|svn_version:'Unknown'|variable_revision_number:16|show_window:False|notes:
[]|batch_state:array([], dtype=uint8)|enabled:True|wants_pause:False]
    Select the type of image to save:Image
    Select the image to save:UpsampledIllumBrightfield
    Select method for constructing file names:Single name
    Select image name for file prefix:None
    Enter single file name:\g<Plate>_IllumBrightfield
    Number of digits:4
    Append a suffix to the image file name?:No
    Text to append to the image name:
    Saved file format:npy
    Output file location:Default Output Folder|
    Image bit depth:8-bit integer
    Overwrite existing files without warning?:No
    When to save:Last cycle
    Record the file and path information to the saved image?:No
    Create subfolders in the output folder?:No
    Base image folder:Elsewhere...|
    How to save the series:T (Time)
    Save with lossless compression?:Yes

SaveImages:[module_num:21|svn_version:'Unknown'|variable_revision_number:16|show_window:False|notes:
[]|batch_state:array([], dtype=uint8)|enabled:True|wants_pause:False]
    Select the type of image to save:Image
    Select the image to save:UpsampledIllumMarker
    Select method for constructing file names:Single name
    Select image name for file prefix:None
    Enter single file name:\g<Plate>_IllumMarker
    Number of digits:4
    Append a suffix to the image file name?:No
    Text to append to the image name:
    Saved file format:npy
    Output file location:Default Output Folder|
    Image bit depth:8-bit integer
    Overwrite existing files without warning?:No
    When to save:Last cycle
    Record the file and path information to the saved image?:No
    Create subfolders in the output folder?:No
    Base image folder:Elsewhere...|
    How to save the series:T (Time)
    Save with lossless compression?:Yes
```
