## Supplementary material for "Rare missense variants of the leukocyte common antigen related receptor (LAR) display reduced activity in transcellular adhesion and synapse formation": Materials and methods

### Whole Exome Sequencing analysis

In our study, we utilized the Case-Cohort dataset, iPSYCH2012 [1], established by the Integrative Psychiatric Research (iPSYCH) consortium, rooted in the Danish demographic. The dataset comprises all samples linked to a psychiatric diagnosis (as specified below) and 30.000 randomly chosen population-based controls. A subset of the iPSYCH2012 dataset, spanning data from 34,544 individuals, presents whole exome sequencing (WES) data of both control participants and those diagnosed with any of the five salient psychiatric maladies as logged in the Danish Psychiatric Central Research Register by the end of 2015. These disorders encompass attention-deficit/hyperactivity disorder (ICD10: F90.0), autism spectrum disorder (ICD10: F84.0, F84.1, F84.5, F84.8 or F84.9), bipolar disorder (ICD10: F30-F31), schizophrenia spectrum (ICD10: F20-F29), and affective disorder (ICD10: F30-F39).

WES analyses were performed on DNA extracted from dried blood spots; a method that has previously been utilized for generating high-quality WES data [2]. After DNA extraction the samples were whole-genome-amplified in triplicate as previously described [3]. Using the Illumina Nextera capture kit the coding regions were isolated and subsequently sequenced on an Illumina HiSeq platform at the Genomics Platform at the Broad Institute of MIT and Harvard, Boston, MA. Raw sequencing data was aligned to the reference genome (Hg19). Genotype-calling was performed as suggested by Genome Analysis Toolkit v.3.4. and QC steps were performed with Hail 0.1 (Hail Team, https://github.com/hail-is/hail). Variants were only included in the analysis if they were located in high-confidence regions in both the iPSYCH and GnomAD databases [4] defined as >80% of the samples having more than 10 x coverage in the genomic region in question. After these QC steps, the dataset was streamlined to represent 28,448 participants, which includes 19,364 cases and 9,084 controls, and a set of 1,352,490 high-quality genetic variants. Annotation of these variants was achieved through SnpEff [5] version 4.3t. Additional allele count details from the gnomAD [4] exomes r2.1.1 database were annotated via SnpSift [6] version 4.3t. Within the scope of our research, we classified variants as 'rare' if their allele count was no greater than 5 across our iPSYCH cohort (n=28,448) and the subset of non-psych non-Finnish European of gnomAD (n=44,779).

### Mapping of rare missense variants and bioinformatical analysis

From the WES data subset of iPSYCH2012 we extracted all variants of *PTPRF* within the gene positions chromosome 1 43991708-44089343. The genomic positions and annotated nucleotide changes were subjected to the Variant Effect Predictor (VEP) [7] tool, using the GRCh37.p13 assembly. As *PTPRF* has many transcripts and thus for most (coding) variants VEP presents >5 consequences pr mutation, the longest transcript of *PTPRF* was selected as the reference, and this transcript represents the canonical Uniprot sequence (ID: P10586). To gain more power for the subsequent domain-wise analysis, we incorporated variant counts from the non-neuro non-Finnish gnomAD cohort into the analysis as well. Whole-gene and per domain burden tests were done with Fisher's Exact test.

### Plasmid preparation

The human *PTPRF* construct was purchased from Genscript inserted into a pcDNA3.1/Zeo(+) vector. This parent plasmid was exposed to high-throughput site-directed mutagenesis as offered by Genscript to create 164 PTPRF rare and 13 non-rare missense variants for cellular experiments.

Human *PTPRF* construct encoding the FN1-4 domains was purchased from Genscript inserted into a pET-9a vector as previously described [8]. The plasmids contained N-terminal 6xhis tag bridged by a TEV cleavage site. This construct was also subjected to mutagenesis to obtain missense variants of the FN1-4 domain.

Plasmid for Fc-NGL-3 was produced as previously described [9]. In brief, the annotated sequence for the extracellular domain has been inserted in frame using 5' NotI and 3' XbaI sites in a modified pCMV6-XL4 vector. This vector inserts the coding region between a C-terminally fused Fc protein bridged by a 3CPro cleavage site (LEVLFQ/GP) and N-terminally prolactin leader peptide (MDSKGSSQKGSRLLLLLVVSNLLLCQGVVSTPVV) and Flag-tag (DYKDDDDK).

For use in cell-surface binding, cellular aggregation and artificial synapse formation assays, the following constructs have previously been described: extracellularly HA-tagged human CD4 (HA-CD4) [10], intracellularly CFP-tagged rat IgSF8 (IgSF8-CFP) [11] and NGL-3 (NGL-3-CFP) [12]. The plasmid pCAGGS-mCherry was a gift from Phil Sharp (Addgene plasmid # 41583) [13].The plasmid pCAG-GFP was a gift from Connie Cepko (Addgene plasmid # 11150) [14].

### Cell culturing and rare variant processing analysis

CHO-K1 cells were maintained at 37 °C and 5 % $CO_2$ in either SFM Hybridoma (cat) or F-12 (cat) media supplemented with 10% FBS. For full length LAR variant analysis, CHO-K1 cells were seeded in 24 well plates at ~100K cells/well. 24 hours after seeding, medium was replaced with SFM Hybridoma media with no additives. Cells were transfected with LAR variants with FuGENE (Promega), using 256 ng DNA and 1 µL FuGENE pr well, and left for 24 hours before the medium was harvested and cells were lysed in 100 µL TNE lysis buffer with cOmplete and PhosStop for 20 minutes on ice. Cells were scraped off and transferred to 1.5 mL Eppendorf tubes and spun at 4500 rpm for 10 minutes at 4 °C before being used for western blotting.

### SDS-PAGE and Western blotting

Cell lysates were spun down at 4500 rpm at 4°C for 10 minutes and cell pellets were discarded. 10-20 µL of lysate was mixed with ¼ of total volume LDS Sample Buffer (Invitrogen, #NP0007) and 1/10 of total volume DTT. The protein mixture was heated to 95 °C for 5 minutes before being loaded on a 4-12 % Bis-Tris gel (Invitrogen) in NuPAGE MOPS buffer (ThermoFisher, #NP0001)). The proteins were transferred to nitrocellulose membrane using iBlot2 Transfer Stack kit (Invitrogen, NB301001) and iBlot2 Dry Blot System (Invitrogen, #IB1001). The membrane was blocked in 50mM Tris-base, 500mM NaCl, 2% Milk powder and 2% Tween-20) for at least 30 minutes and incubated with primary LAR antibody (R&D systems, AF3004) recognizing the LAR ECD overnight (ON). The antibodies were diluted in blocking buffer. The next day, the membrane was washed three times for 10 minutes in washing buffer (CaCl2 2mM, MgCl2 1mM, HEPES 10mM, NaCl 140mM, 0.2% Milk powder and 0.5% Tween-20) and incubated with HRP-conjugated secondary antibody (diluted 1:2000 in blocking buffer, rabbit α-goat (DAKO #0160) for 1 hour at room temperature. Next, the membrane was washed 3x5 min in washing buffer and bands developed using Amersham ECL Western Blotting Detection Reagent (GE Healthcare, # RPN2106) or SuperSignal West Femto Maximum Sensitivity Substrate (ThermoFisher Scientific, #34096) - following supplier's instructions - and detected and analyzed using LAS-4000 and appurtenant software (GE Healthcare) or iBright™ CL1500 Imaging System (A44114, Invitrogen). The intensity of bands was quantified by densitometric analysis using Multi Gauge V3.2 software.

### Protein production and purification

The production of LAR FN1-4 constructs was performed as previously described [8]. The plasmids for FN1-4 WT or mutants were transformed into E. coli BL21(DE3) cells. Transformed cells were cultured at 37°C in LB broth medium containing 100 μg/mL kanamycin. When optical density reached 0.85 at 600 nm the culture was supplied with 1 mM isopropyl B-D-1-thiogalactopyranoside (IPTG) to induce protein expression and the culturing temperature was lowered to 30°C and left ON for 14-16 hours at 225 rpm. The cells were harvested by centrifugation at 6000g for 20 minutes at 4°C and pellets were stored at -20°C. The cells were lysed in by sonication pulses of 30 % for 3 x 5 minutes on ice. Bacterial debris was removed by ultracentrifugation at 10,000g for x min at 4°C. His-tagged protein was extracted by either application to a HisTrap HP column (GE Healthcare) at 1 ml/min or by application onto Talon beads. In both cases, the column/beads were washed in 50 mM Tris pH 7.4, 300 mM NaCl, 20 mM imidazole, and the protein eluted in elution buffer containing 50 mM Tris pH 7.4, 300 mM NaCl, 400 mM imidazole. Eluted protein was dialyzed overnight at 4 °C against PBS pH 7.5 containing 10 % glycerol The dialyzed protein solution was exposed to size-exclusion chromatography (SEC) using a Superdex 200 Increase 16/600 column (GE Healthcare) after equilibration to a SEC buffer containing 50 mM HEPES pH 7.5, 50 mM NaCl. Peak SEC fractions were analysed with SDS-PAGE and upon confirmation of protein size were pooled and concentrated to ~3 mg/mL before storage at -80°C.

Production of Fc-fusion proteins was performed using the ExpiCHO system (Gibco A29133). ExpiCHO-S™ cells were maintained in suspension in 125-mL polycarbonate Erlenmeyer shaker flasks at 37 °C, >80% humidity and 8 % $CO_2$ at ~125 rpm using ExpiCHO™ Expression Medium. One day before transfection, cells were seeded at 4*10^6 cells/mL in 120 mL Expression Medium. On the day of transfection, cells were diluted to a final concentration of 6*10^6 cells/mL. 1 μg of DNA pr mL of culture was diluted in ice-cold OptiPRO medium and mixed with ice-cold ExpiFectamine-OptiPRO mixture and left at RT for 5 minutes before adding to the shaker flask containing the ExpiCHO-S cells. 18-22 hours post transfection, the cultures were supplemented with freshly prepared mixture of ExpiFectamine CHO Enhancer and ExpiCHO Feed. Protein was harvested at 8-10 days after transfection by centrifugation at 6000g for 10 min at 4°C. For purification, the solution was subjected to a Protein A column at 1 mL $min^{-1}$ at 4 °C ON. The Fc-region was cleaved off utilizing the 3C cleavage site with 3C protease. The protein solution was exposed to size-exclusion chromatography (SEC) using a Superdex 200 Increase 16/600 column (GE Healthcare) after equilibration to a SEC buffer containing 50 mM HEPES pH 7.5, 50 mM NaCl. Peak SEC fractions were analysed with SDS-

PAGE and pooled and concentrated to ~3 mg/mL and stored in 50 mM HEPES pH 7.4, 50 mM NaCl at -80 °C.

### Thermostability using Tycho NT.6

Alterations to LAR FN1-4 stability were assessed using the Tycho NT.6 apparatus from NanoTemper. LAR FN1-4 variant samples were prepared at ~25 µg/mL in 50 mM HEPES pH 7.5, 50 mM NaCl. The samples were subjected to a temperature increment from 30–95 °C and the intrinsic fluorescence of tryptophane and tyrosine aromatic rings at 330 nm and 350 nm were measured throughout. The ratio of 330/350 nm fluorescence was used to analyze the thermal transition temperatures. Inflection temperatures were calculated automatically with the inherent Tycho NT.6 software.

### ThermoFlour assay:

Screening of melting transition point of LAR FN1-4 mutants was performed using protein mixtures with SYPRO Orange. A mixture of 4 µM LAR FN1-4 protein and 5 x SYPRO orange was diluted in 50 mM HEPES pH 7.5 50 mM NaCl and loaded into 96-well qPCR plates in triplicates. Several LAR FN1-4 WT batches were spread across the plate to eliminate positioning effects. Melting transition was assessed using an AriaMx Real-time PCR System (Agilent) with FAM filter. The AriaMx was programmed to perform a melting curve with a temperature curve from 40-95 °C in 0.2 °C increments, with each increment lasting 5 seconds. Intensity signals were measured with the SYBR/FAM filter. Raw intensities were normalized to the highest value and values above 60 °C were omitted from the analysis. Melting transition points were analyzed by Boltzmann sigmoidal fits in Graphpad Prism.

### Microscale Thermophoresis

LAR FN1-4 variants were analyzed for binding to NGL-3 using Microscale thermophoresis (MST). His-tagged LAR FN1-4 WT and mutants were labelled with RED-tris-NTA 2$^{nd}$ generation dye (MO-L018, NanoTemper). Optimal labeling conditions were determined as 200 nM LAR FN1-4 and 25 nM fluorophore in 25 mM HEPES pH 7.5, 150 mM NaCl, 0.01 % Tween20 with incubation for 20 min at RT followed by centrifugation for 10 min at 15000g at 4 °C. 1:1 dilution series of NGL-3 were prepared in 25 mM HEPES pH 7.5, 150 mM NaCl in PCR tubes and labeled LAR FN1-4 was added to a final concentration of 100 nM and 0.005 % Tween20. Samples were briefly spun down and incubated at RT for 1 h protected from light. Samples were then loaded into MonolithTM NT.115 Monolith Premium Capillary Chips and run on a Monolith NT.115 (NanoTemper Technologies GmbH). MST was run with 100 % LED power, medium MST power with 3 seconds initial fluorescence, 20 seconds MST on time and 1 second back diffusion. The data was acquired with MO.Control software and analyzed using MO.Affinity Analysis (NanoTemper Technologies GmbH). MST on time for analysis was set to 2.5 s and the data was fitted using a 1:1 stoichiometry $K_D$ model inherent to the software.

### Cellular aggregation assays

Two groups of HEK293T cells grown in 12-well plates were transfected with the appropriate plasmids using TransIT-LT1 (Mirus Bio. LLC; #MIR2305) and cultured in Dulbecco's modified Eagle's medium (DMEM) (Gibco; #11965118) containing 10% (v/v) fetal bovine serum (FBS) (Wisent; #080-150). After 48 hours, HEK293T cells were trypsinized and resuspended in 250 µl of DMEM containing 10% (v/v) FBS before mixing of appropriate GFP- and mCherry-expressing cells to a 1:1 ratio. Mixed HEK293T groups were rotated at room temperature for 2 hours to allow cells to aggregate. Cell mixtures (500 µl) were added to 24-well plates and then imaged on an IncuCyte S3 Live-Cell Analysis System (Sartorius) with a 4x air objective. Images were acquired as 16-bit grayscale.

### *In situ* surface binding assays using soluble proteins

Protein binding assays were performed as described previously [15, 16]. Briefly, to assess protein binding on the surface of COS7 cells, the cells were transfected using TransIT-LT1 (Mirus Bio. LLC; #MIR2305) with appropriate plasmids and then cultured in DMEM (Gibco; #11965118) containing 10% (v/v) FBS (Wisent; #080-150). Twenty-four hours after transfection, the transfected COS7 cells were washed with extracellular solution (ECS) containing 168 mM NaCl, 2.4 mM KCl, 20 mM HEPES, pH 7.4, 10 mM D-glucose, 2 mM $CaCl_2$, 1.3 mM $MgCl_2$, and 100 µg/ml bovine serum albumin (BSA; Sigma, #A9647); the cells were then incubated for 1 hour at 4 °C with NGL-3-Fc or SALM5-Fc (R&D Systems; #9385-SA) proteins diluted in ECS to 2.5 and 0.2 µM respectively. Cells were washed with ECS and subsequently fixed in prewarmed 4% (v/v) paraformaldehyde (PFA)/4% (w/v) sucrose in PBS for 12 minutes and blocked in 5% (v/v) normal donkey serum (NDS) and 3% (w/v) BSA in PBS for 1 hour at room temperature. COS7 cells were incubated with anti-LAR (0.1 µg/ml; goat; R&D Systems; #AF3004) or anti-HA (0.5 µg/ml; rabbit; Abcam; #ab9110) without permeabilization overnight at 4 °C. Cells were then incubated with highly cross-adsorbed Alexa dye–conjugated secondary antibodies generated in donkey toward the appropriate species (1.5 µg/ml; Jackson ImmunoResearch or ThermoFisher). Images were acquired on a Leica DM6 fluorescence microscope with a 40x 1.25 NA oil objective and a Hamamatsu C11440 ORCA-Flash 4.0 camera using LasX software (Leica). Images were acquired as 16-bit grayscale. For quantification, sets of cells were stained simultaneously and imaged with identical settings.

### Artificial synapse formation assay

All animal experiments were carried out in accordance with Canadian Council on Animal Care guidelines and approved by the IRCM Animal Care Committee. Cocultures of rat hippocampal neurons with HEK293T cells were set up as previously [15, 16]. Briefly, hippocampal neurons from E18 rat embryos were cultured on poly-L-lysine-coated glass coverslips in neurobasal medium (Gibco; #21103-049) supplemented with NeuroCult SM1 (StemCell; #05711) and GlutaMaX (Gibco; #35050061). For neuron-HEK293T coculture assays, appropriate plasmids were transfected in HEK293T cells using TransIT-LT1 (Mirus Bio. LLC; catalog number: MIR2305) and cultured in DMEM containing 10% (v/v) FBS. Twenty-four hours after transfection, cells were harvested by trypsinization and seeded on the neuron cultures at 14 days *in vitro* (DIV). After 24 hours of coculture, cells were fixed in prewarmed 4% (v/v) PFA/4% (w/v) sucrose in PBS for 12 min and blocked with 5% (v/v) NDS and 3% (w/v) BSA in PBS for

1 hr at room temperature. Cells were incubated with anti-LAR (0.1 µg/ml; goat; R&D Systems; #AF3004) or anti-HA (0.5 µg/ml; rabbit; Abcam; #ab9110) overnight at 4 °C to label transfected surface-expressed proteins. Cells were then permeabilized in 0.2% (v/v) Triton X-100 in PBS and incubated with anti-PSD95 (3.3 µg/ml; mouse; ThermoFisher, #MA1-045) overnight at 4 °C. Cells were then incubated with highly cross-adsorbed Alexa dye–conjugated secondary antibodies generated in donkey toward the appropriate species (1.5 µg/ml; Jackson ImmunoResearch or ThermoFisher). Images were acquired on a Leica DM6 fluorescence microscope with a 63x 1.40 numerical aperture (NA) oil objective and a Hamamatsu C11440 ORCA-Flash 4.0 camera using LasX software (Leica). Images were acquired as 16-bit grayscale. For quantification, sets of cells were stained simultaneously and imaged with identical settings.

### Image analysis

To quantify binding levels and cell surface expression levels, we measured the average intensity of each channel within the delineated COS7 or HEK293T cell area subtracted by the average intensity of the off-cell background. For *in situ* binding assays, the average intensity of bound soluble NGL-3-Fc protein was normalized using the average surface intensity of the LAR or CD4 protein signal. COS7 cells expressing similar levels of LAR or CD4 proteins were selected to quantify bound soluble NGL-3-Fc protein. Analyses were performed using Volocity 6.0, Excel for Microsoft 365 (Microsoft), and Prism 9 (GraphPad Software, Inc). For the artificial synapse formation assays, HEK293T cells displaying similar surface levels of LAR or CD4 proteins were imaged without considering the channel corresponding to PSD95 signal. To assess glutamatergic postsynaptic differentiation, the fluorescence channel corresponding to PSD95 was thresholded, and the total tresholded intensity within regions positive for surface LAR or HA was measured and normalized to HEK293T cells area. Analysis was performed using Metamorph 7.8 (Molecular Devices), Excel for Microsoft 365, and Prism 9. For the cellular aggregation assays, the aggregation index was calculated by dividing the number of particles with an area higher that of a single cell by the total number of particles. Analysis was performed using the Analyze Particles function of the FIJI processing package (ImageJ). Agostino's K-squared test was used to assess normal distributions, and Bartlett's test was used to check whether standard deviations (SDs) were significantly different across groups. To compare three or more groups with normal distribution, Welch's ANOVA with Dunnett's T3 post hoc analysis was used. For comparisons between three or more groups without normal distribution, Kruskal Wallis tests with Dunn's post hoc analysis were used. Statistical significance was examined with appropriate tests as indicated in the figure legends. All data are reported as the mean ± SEM from at least three independent experiments, and statistical significance was defined as $p < 0.05$.

### Small Angle X-ray Scattering

Purified LAR variants are concentrated to different series (1 mg/ml, 1.5 mg/ml, 2 mg/ml and 2.5 mg/ml) in gel filtration buffer (150 mM NaCl, 100 mM Tris pH 7.5). SAXS data were collected at in-house instrument at iNANO at Aarhus University [17] at 20°C for 1800 second measurement for each sample. Data collection methodology is detailly described in [17]. Raw data processing, water scattering, background and beam-stop shadow corrections, data reduction and buffer subtraction are done by in-house software package Supersaxs (Oliveira

& Pedersen, unpublished). The resulting data are subjected to Guinier fit analysis, Kratky analysis and IFT(GNOM) by using BioXTAS RAW software package [18]. Scattering curves are normalized to concentration and plotted in log-log scale with the associated error bars. Ab-initio models are determined by dummy residue modelling that are constructed from scattering profiles by using DAMMIF which is part of the ATSAS package and scattering data are fitted to models by using CRYSOL in ATSAS software package [19]. All the structural models are prepared by using The PyMOL Molecular Graphics System, Version 3.0 Schrödinger, LLC.

### Adapted Lipocyte Profiling

This assay is an adaptation of LipocyteProfiler and Cell Painting [20, 21]. U2OS cells were maintained in McCoy's 5A medium supplemented with 10 % FBS and were plated in 384 well PhenoPlates (PerkinElmer # 6057302) at a density of 2K cells / well. After 24 hours, the medium was changed to McCoy's 5A without FBS and the cells were transfected with GFP, LAR WT or missense variants using Lipofectamine™ 3000 Transfection Reagent (Invitrogen L3000015). To minimize plate-layout effects, each replicate of the different transfections (n=6) was placed as suggested from: https://carpenter-singh-lab.broadinstitute.org/blog/how-normalize-cell-painting-data. 24 hours after the transfection, the medium was supplemented with MitoTracker Deep Red (Invitrogen #M22426) to a final concentration of 500 nM and incubated for 30 min at 37 °C. Then all wells were subjected to a 16 % PFA solution to a final concentration of 4 % and kept for 20 min protected from light. This solution was then removed, and the wells were washed twice with 1x HBSS (Gibco #14025076) and incubated with a multi-stain solution containing 1 unit/mL Alexa Fluor™ 568 Phalloidin (Invitrogen A12380), 10 µg/mL Hoechst 33342 (Invitrogen H3570), 1.5 µg/mL Wheat Germ Agglutinin Alexa Fluor™ 555 Conjugate (Invitrogen W32464), 0.1 % Triton X-100 (Sigma Aldrich #X100) in 1x HBSS containing 1 % BSA and incubated for 30 min at RT protected from light. The solution was removed, wells washed twice in 1x HBSS and anti-LAR (R&D, AF3004) in 1x HBSS containing 1 % BSA was added to the wells and left ON at 4 °C. On the next day, the wells were washed twice with 1x HBSS and secondary stained with an Alexa Fluor™ 488 conjugated antibody (Sigma Aldrich # A-11055 for 2 hours at RT protected from light. Finally, the wells were washed three times with 1x HBSS and imaged using the Opera Phenix High Content screening system in confocal mode with the 20x objective and numerical aperture of 1.

Feature extraction and quantification was performed using CellProfiler. Prior to image processing, the images were flat field illumination correction was performed using mean averaging across all image sets, after which median smoothing was applied and the resulting image from the illumination correction function is then used to divide from the original image. Nuclei were identified with the Hoechst stain and then used to identify single cells through the WGA/Phalloidin stain. Cytoplasmic regions were found by subtracting the nuclei segments from the whole cell segments. For each of these compartments, measures related to size, shape, intensity, granularity, texture, colocalization and distance to neighboring objects were extracted. The data was normalized across the plate using subtraction of the median of each variable and dividing it with the interquartile range. α-LAR signal was used to identify transfected cells and only aggregate these for downstream analysis. Subsequent manual filtering steps included total LAR positive cell count >25 and removal of artefacts. The features were normalized across the plate and pr well means were calculated for each feature and used to generate per-feature Z-scores and p-values for each LAR variant. This normalized, filtered data was analyzed using R4.3.1.

Of note, the U2OS cells were used for their compatibility with the high-throughput format and easy transfection and growth in imaging plates. Analysis was performed directly between variants and WT, since several studies have shown that finding relevant negative controls in these highly sensitive morphological studies [22, 23].

All code used in the manuscript will be made available.

### Methods references
