## Supplementary figures for "Rare missense variants of the leukocyte common antigen related receptor (LAR) display reduced activity in transcellular adhesion and synapse formation"

### Supplementary Figure 1

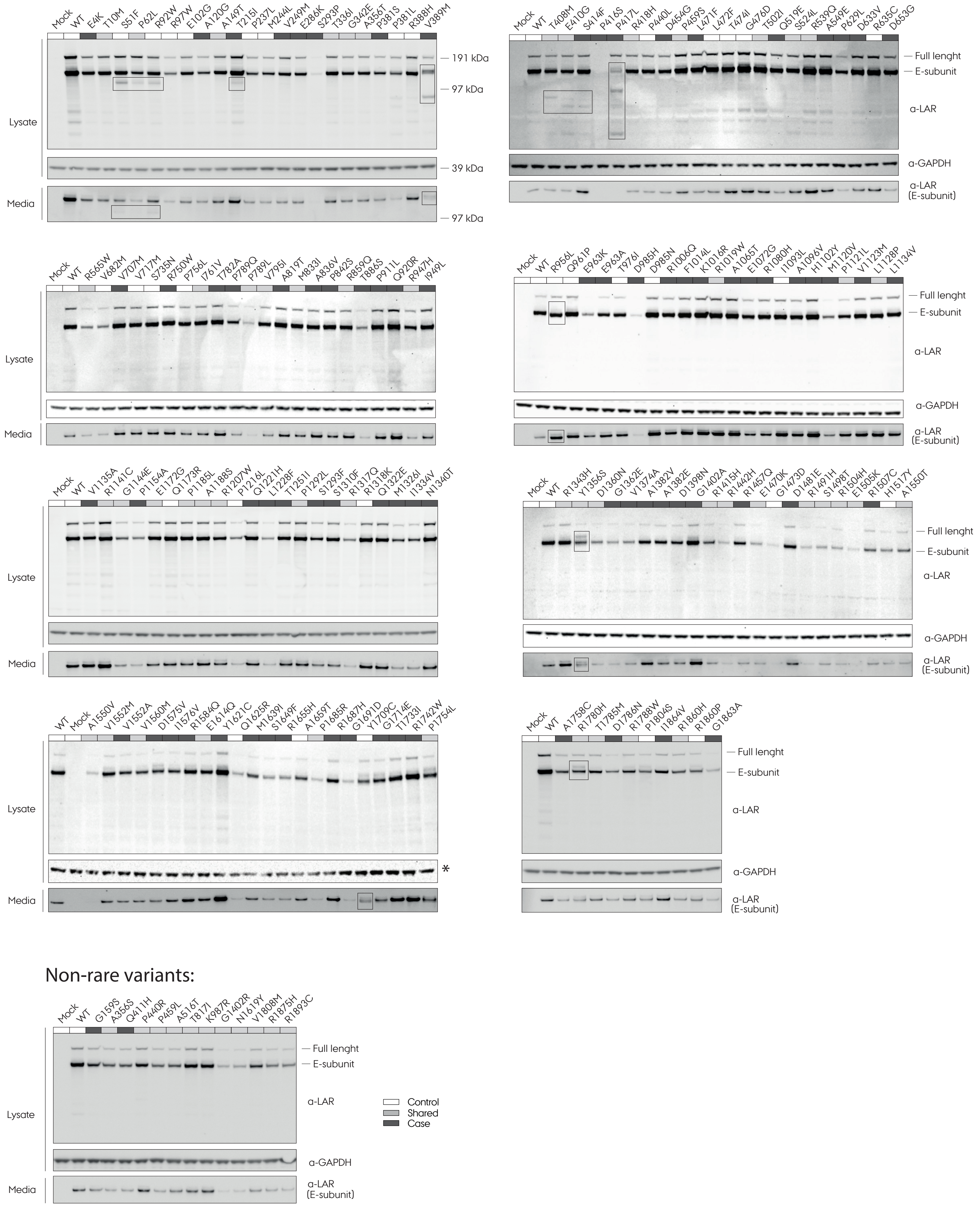

### Supplementary Figure 2

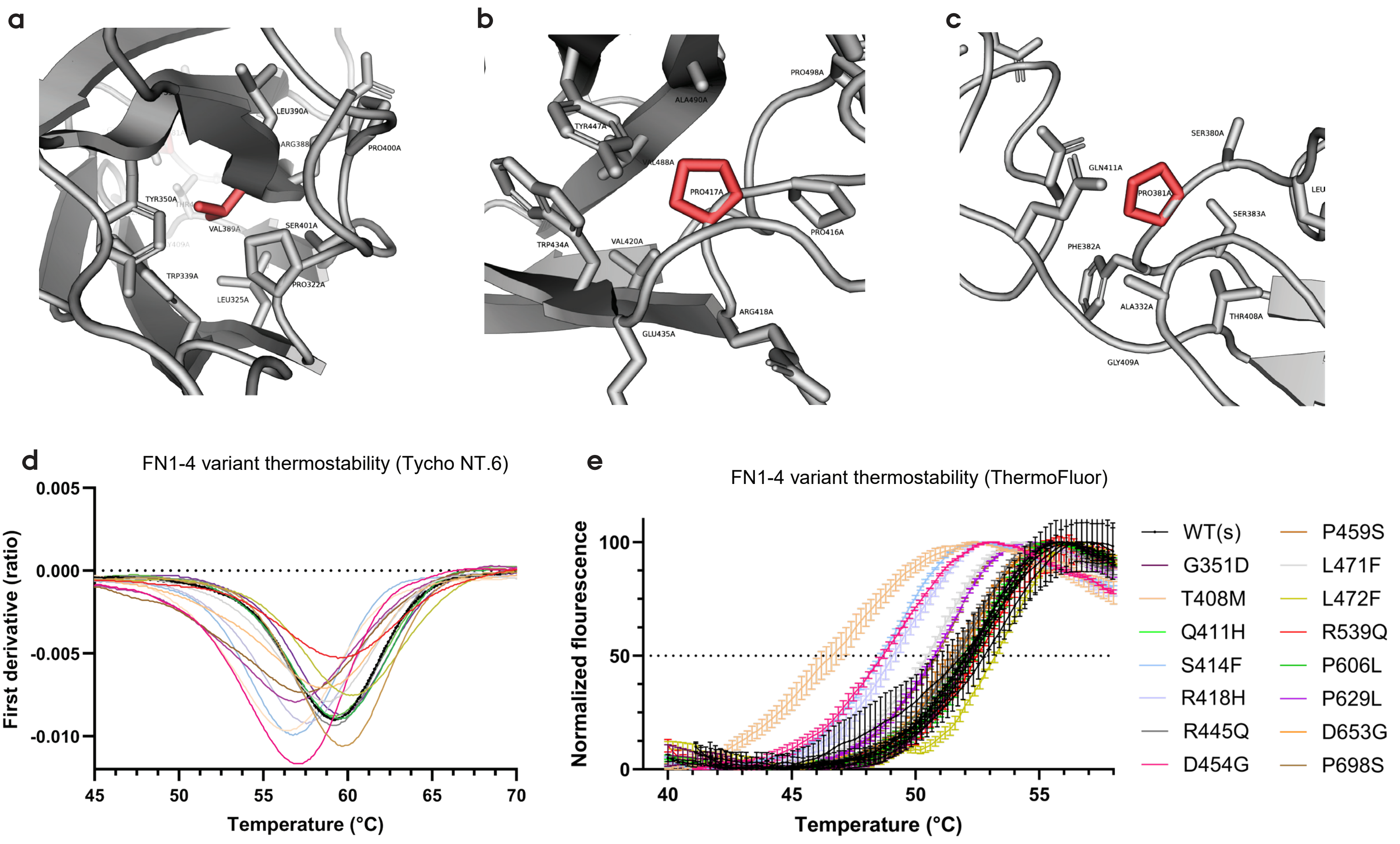


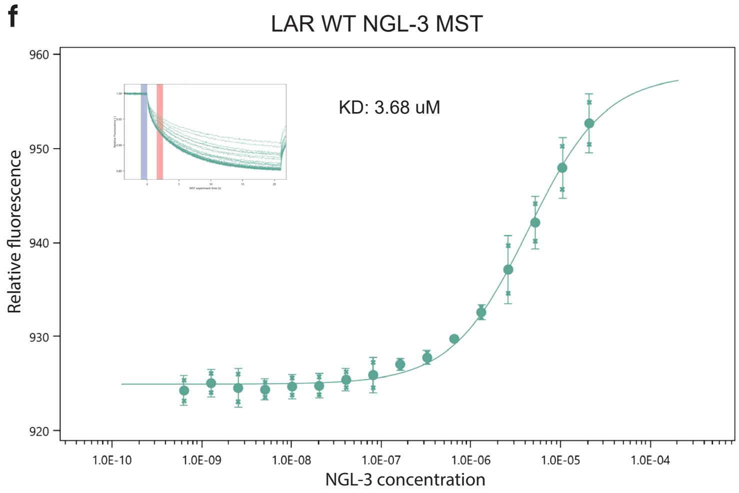


g

| | KD μM | KD change (μM) | Affinity FC |
|---|---|---|---|
| WT | 3,68 | | |
| P381S* | 0,10 | -3,58 | 38,33 |
| P381L | N/A | N/A | N/A |
| R388H | 6,63 | 2,95 | 0,56 |
| V389M | N/A | N/A | N/A |
| P416S | 5,14 | 1,46 | 0,72 |
| P417L | N/A | N/A | N/A |
| R454G | 2,97 | -0,71 | 1,24 |
| L471F | 3,90 | 0,22 | 0,94 |
| L472F | 2,45 | -1,23 | 1,50 |
| R539Q | 2,53 | -1,15 | 1,46 |
| D653G | 6,06 | 2,38 | 0,61 |

#### Supplementary Figure 3

**a**

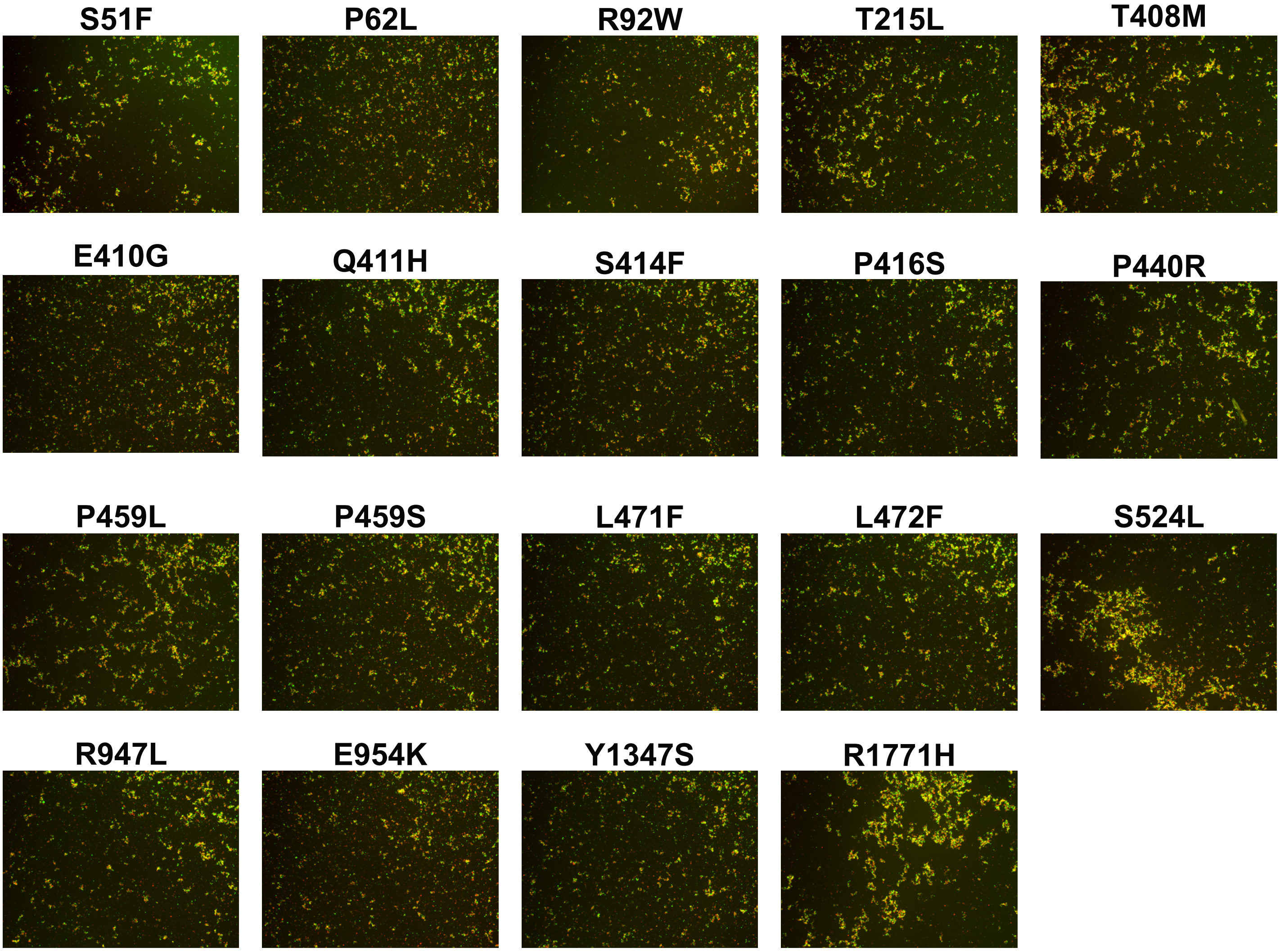


**b**

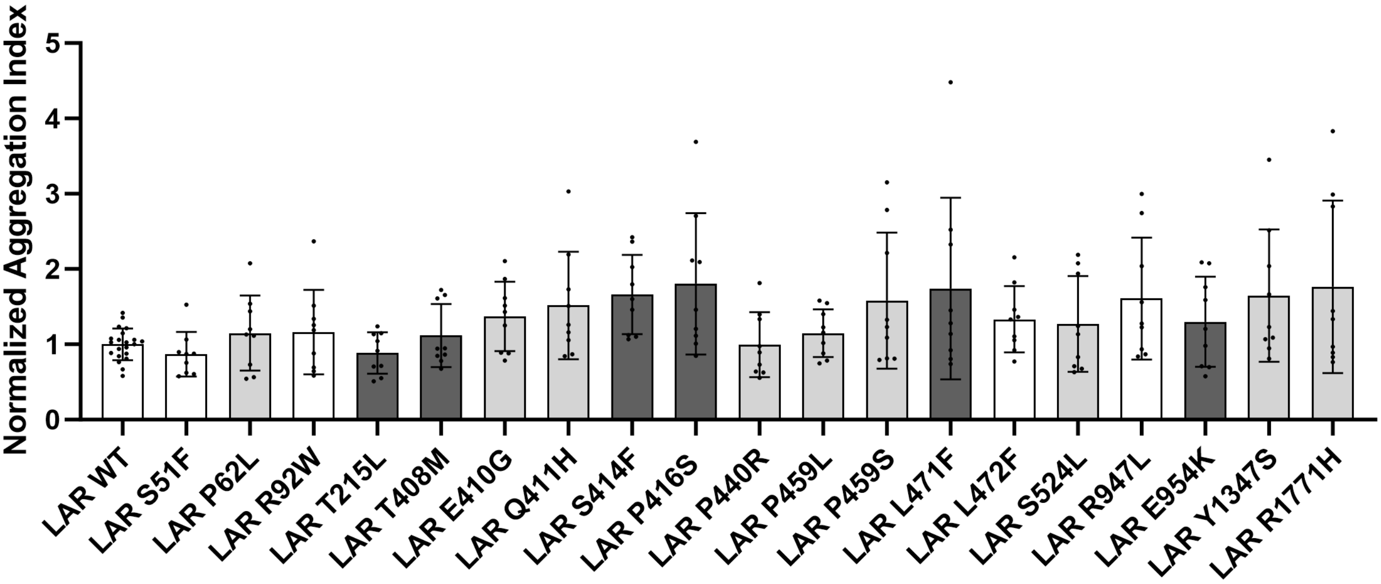

### Supplementary Figure 4

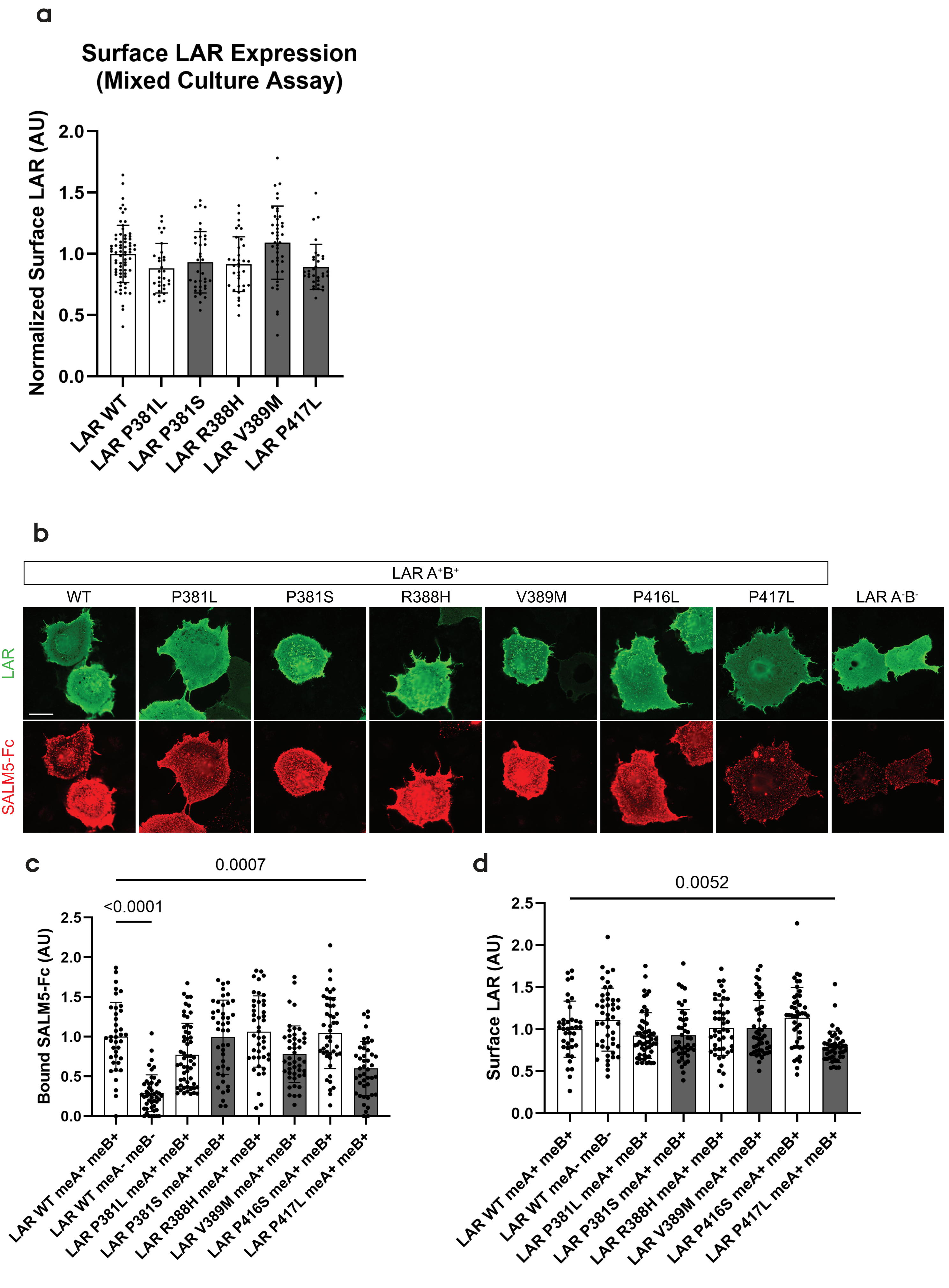

### Supplementary Figure 5

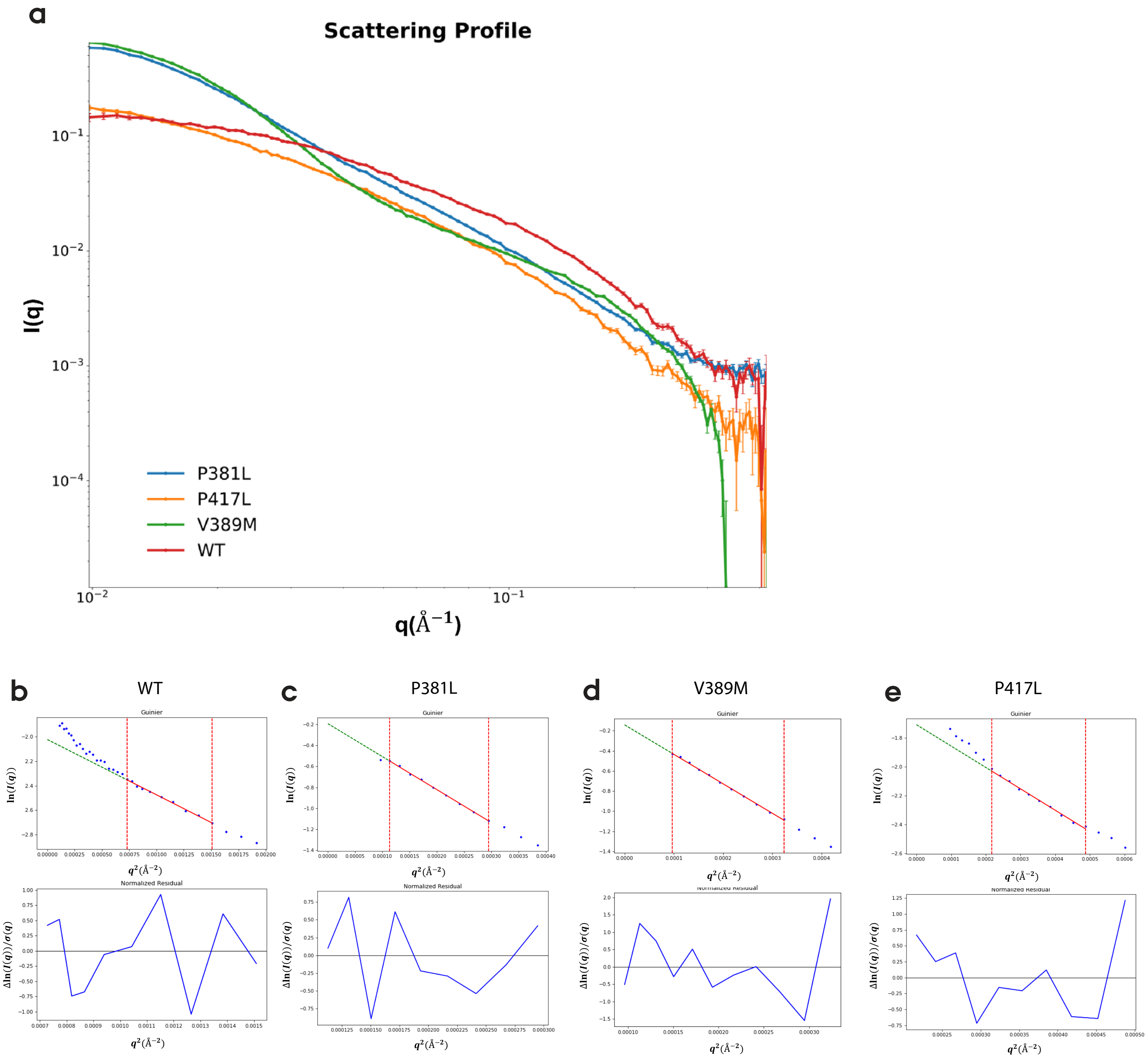

#### Supplementary Figure 6

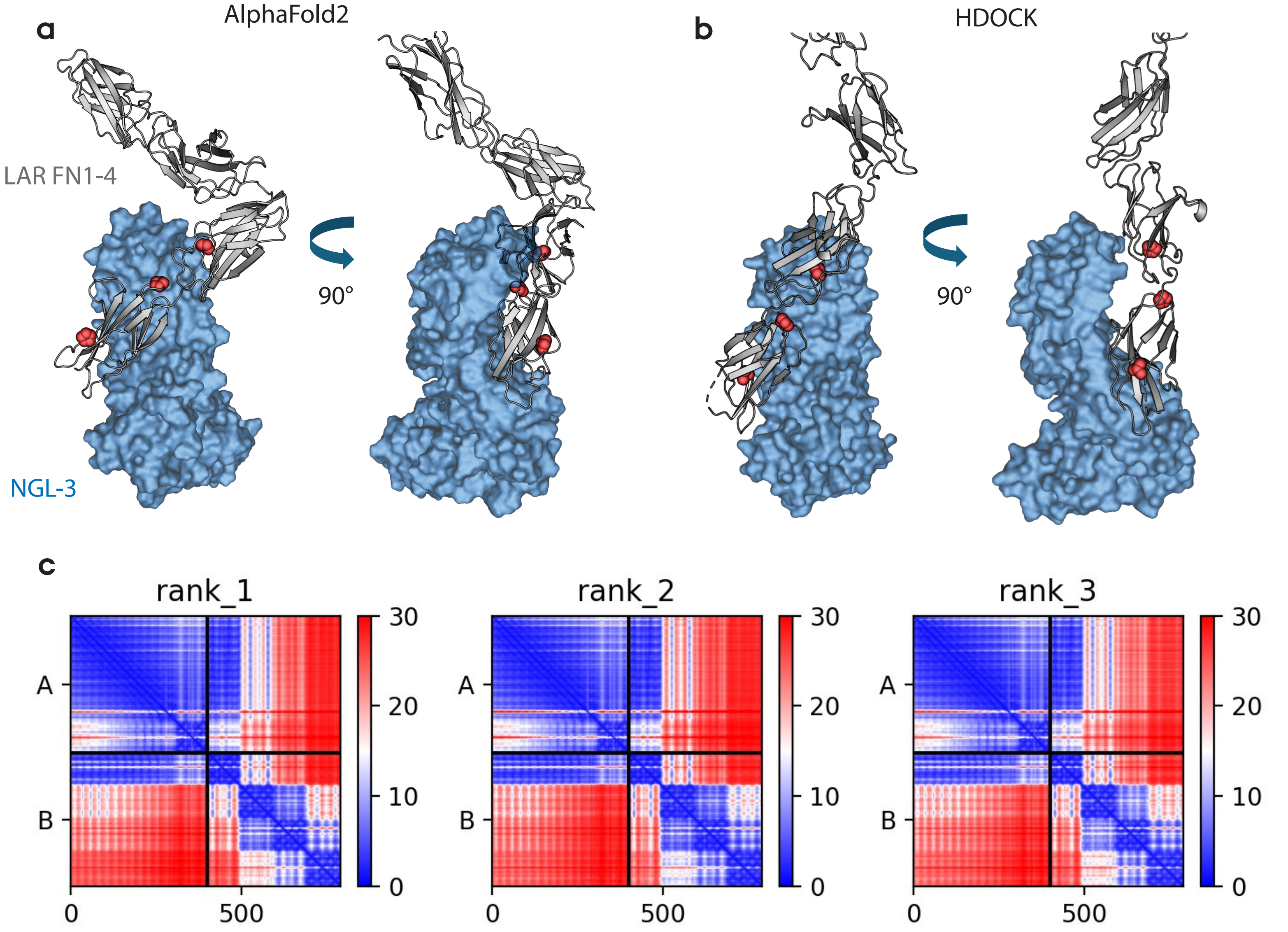

### Supplementary Figure 7

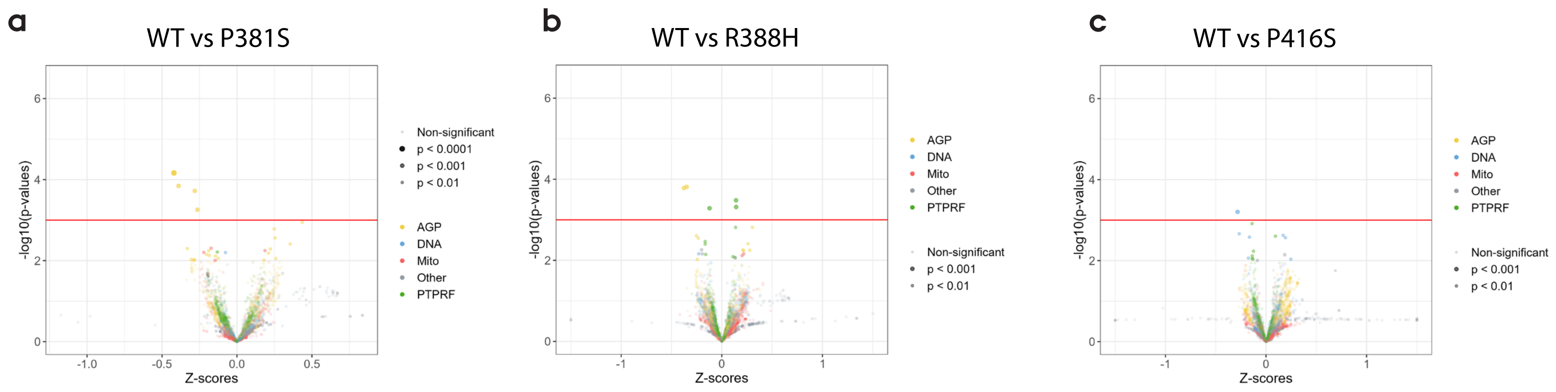
