## Supplementary material for "Rare missense variants of the leukocyte common antigen related receptor (LAR) display reduced activity in transcellular adhesion and synapse formation": Main figures

### Figure 1

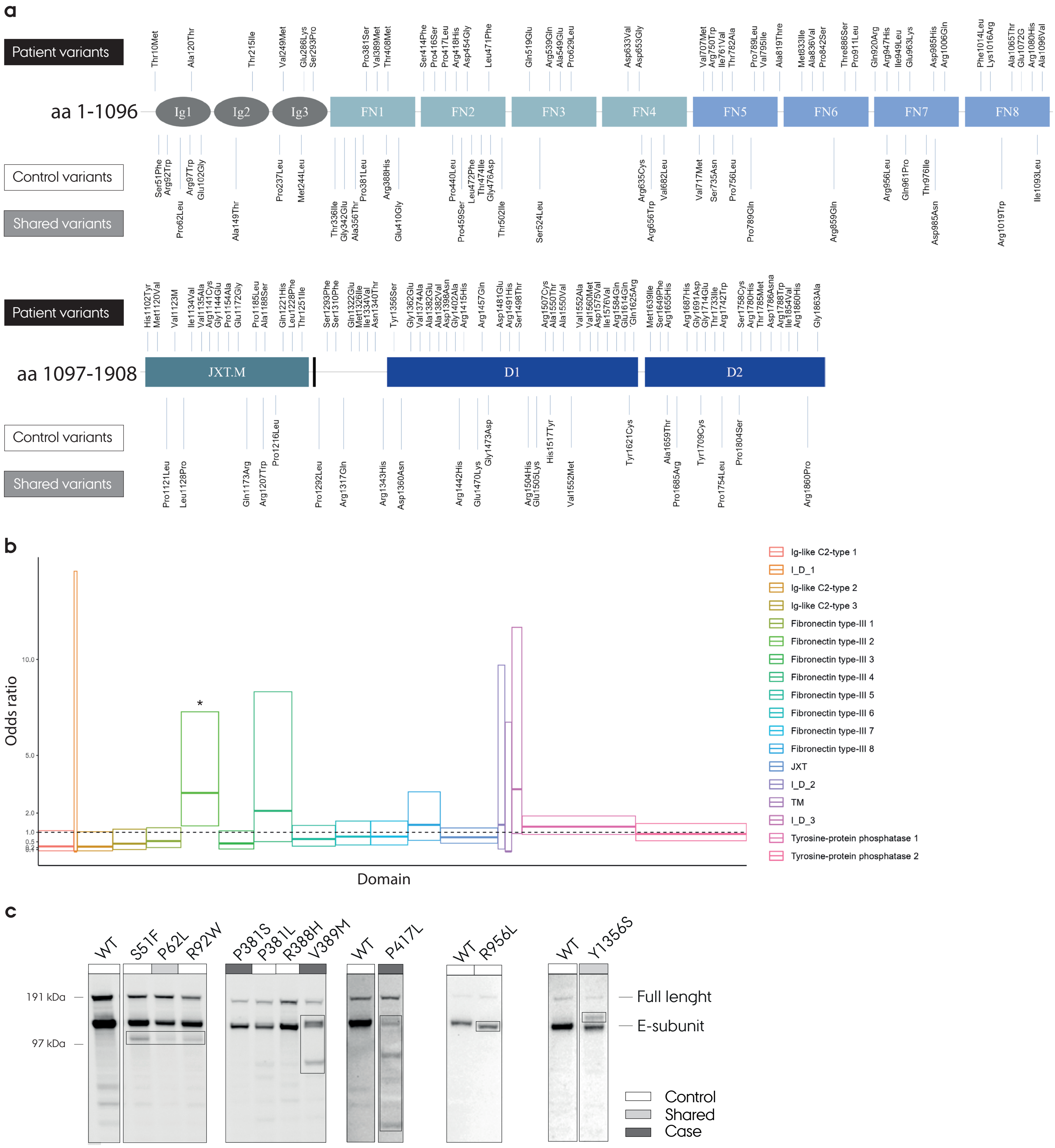
a
Patient variants
Control variants
Shared variants
aa 1-1096
Ig1
Ig2
Ig3
FN1
FN2
FN3
FN4
FN5
FN6
FN7
FN8
aa 1097-1908
JXT.M
D1
D2
b
Odds ratio
Domain
Ig-like C2-type 1
I_D_1
Ig-like C2-type 2
Ig-like C2-type 3
Fibronectin type-III 1
Fibronectin type-III 2
Fibronectin type-III 3
Fibronectin type-III 4
Fibronectin type-III 5
Fibronectin type-III 6
Fibronectin type-III 7
Fibronectin type-III 8
JXT
I_D_2
TM
I_D_3
Tyrosine-protein phosphatase 1
Tyrosine-protein phosphatase 2
c
WT
S51F
P62L
R92W
P381S
P381L
R388H
V389M
P417L
R956L
Y1356S
191 kDa
97 kDa
Full lenght
E-subunit
Control
Shared
Case

### Figure 2

a

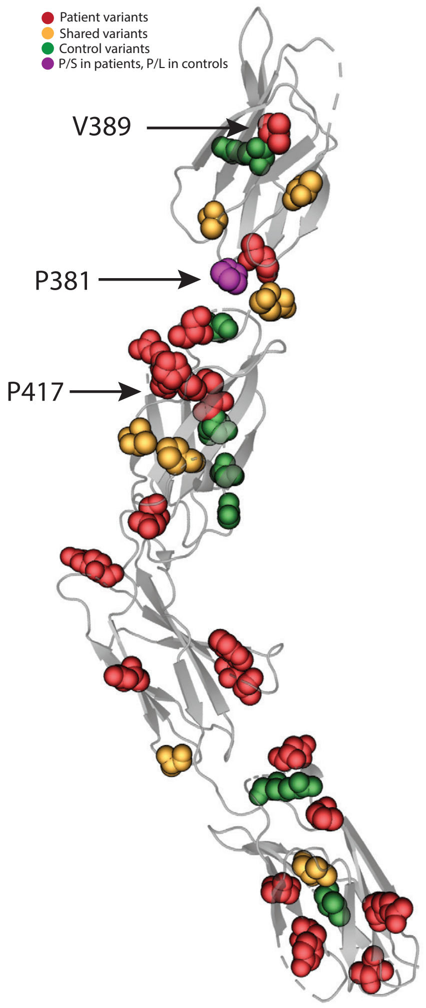


b

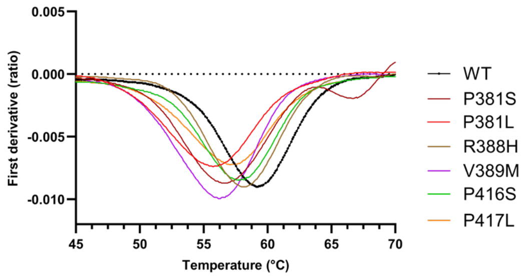


c

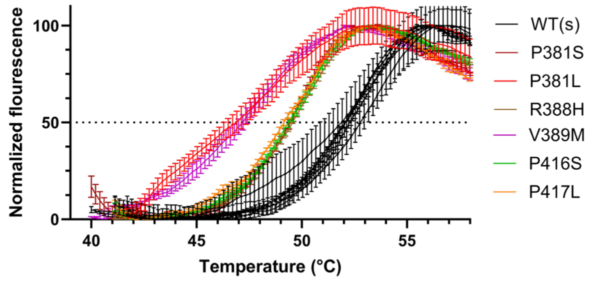


d

| | ΔT (ºC) | |
|---|---|---|
| Variant | ThermoFluor | Tycho |
| G351D | 0.1 | 0.4 |
| P381S | -5.2 | -2.6 |
| P381L | -5.4 | -4.0 |
| R388H | -5.0 | -1.6 |
| V389M | -9.4 | -3.5 |
| T408M | -10.6 | -3.4 |
| Q411H | 0.1 | -0.5 |
| S414F | -6.7 | -2.9 |
| P416S | -5.3 | -1.9 |
| P417L | -5.6 | -2.5 |
| R418H | -5.6 | -2.1 |
| R445Q | -0.8 | -0.5 |
| D454G | -6.7 | -2.6 |
| P459S | -0.7 | 0.1 |
| L471F | -3.1 | -1.0 |
| L472F | 2.3 | 0.6 |
| R539Q | 0.7 | -0.2 |
| P606L | 0.4 | -0.2 |
| P629L | -2.6 | -2.8 |
| D653G | -0.8 | -1.0 |
| P698S | -2.5 | -2.3 |

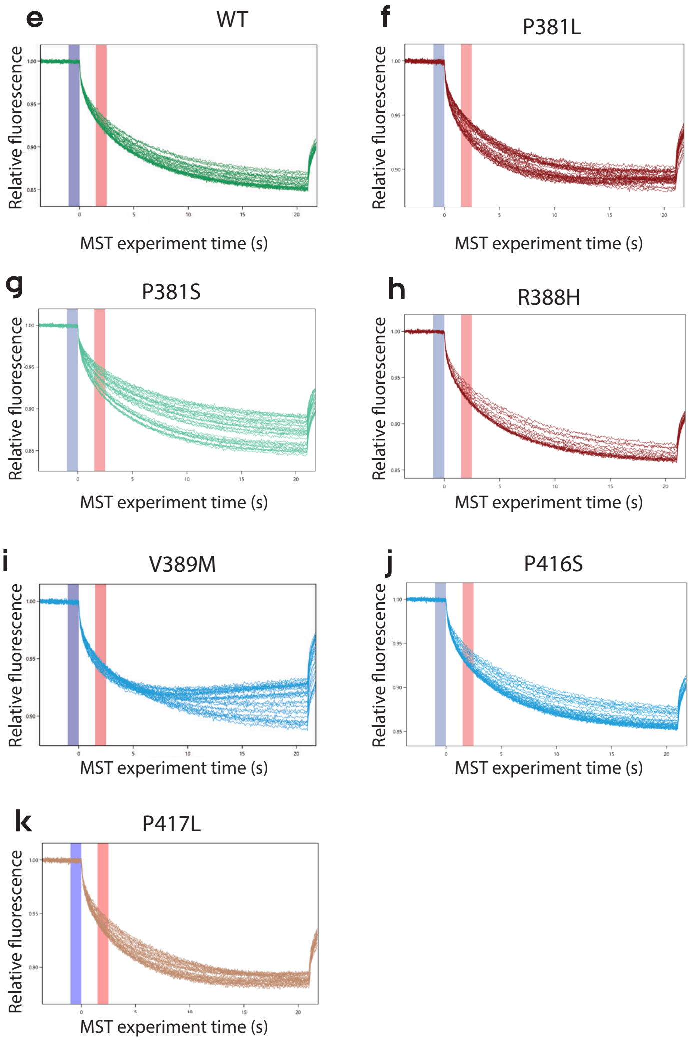

### Figure 3

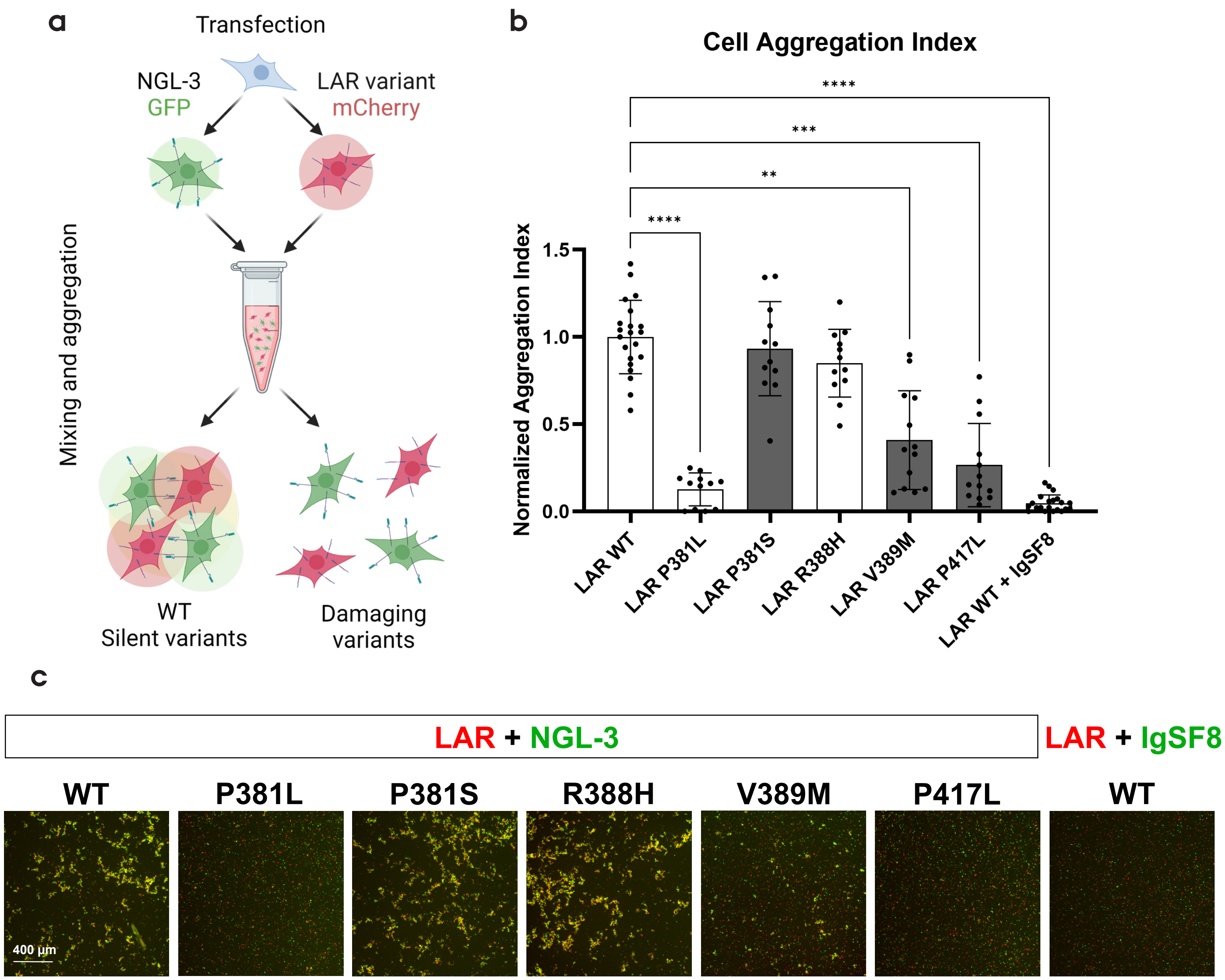
a
Transfection
NGL-3
GFP
LAR variant
mCherry
Mixing and aggregation
WT
Silent variants
Damaging
variants
b
Cell Aggregation Index
Normalized Aggregation Index
1.5
1.0
0.5
0.0
****
***
**
****
LAR WT
LAR P381L
LAR P381S
LAR R388H
LAR V389M
LAR P417L
LAR WT + IgSF8
c
LAR + NGL-3
LAR + IgSF8
WT
P381L
P381S
R388H
V389M
P417L
WT
400 μm

### Figure 4

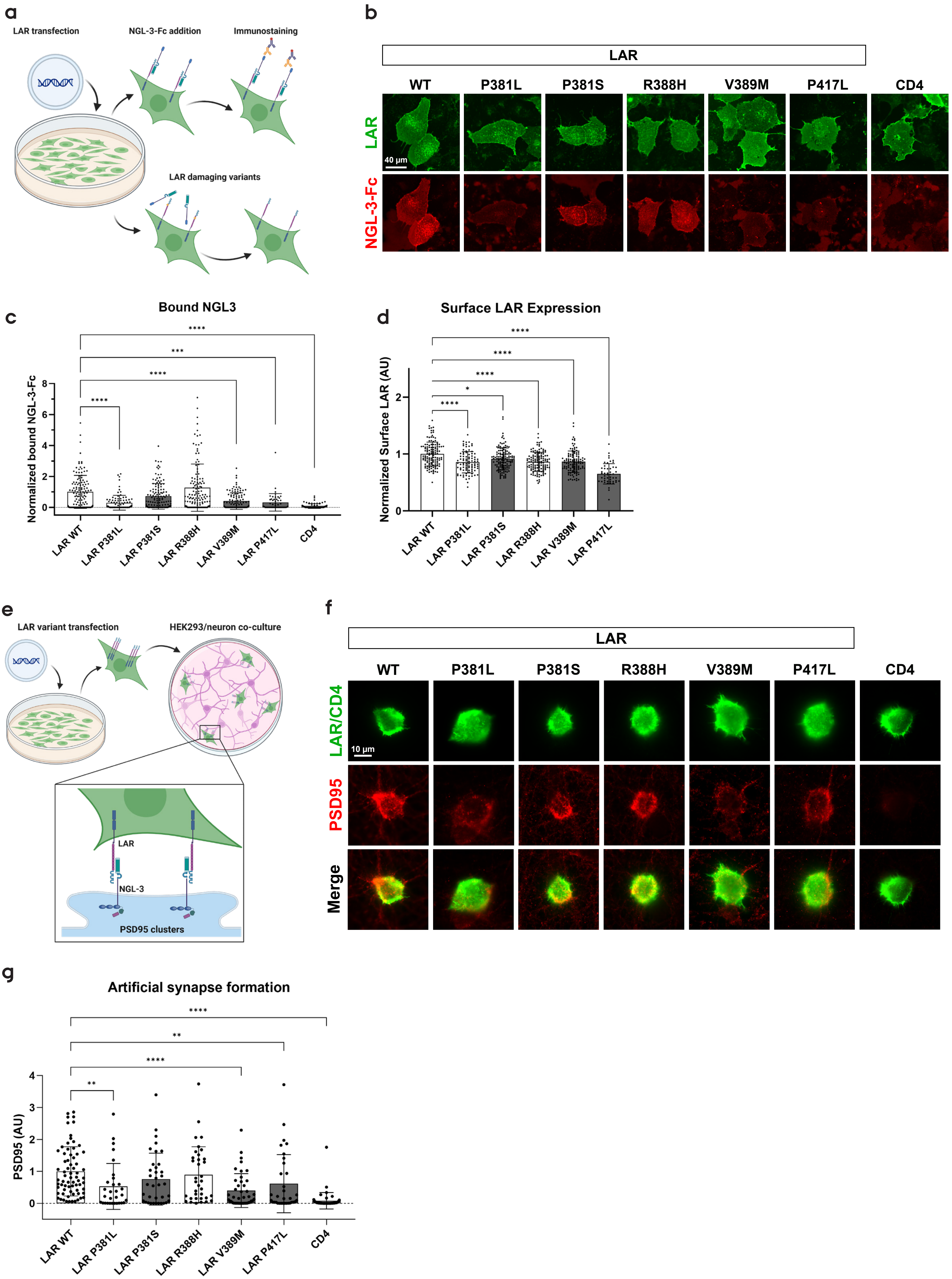

a
LAR transfection
NGL-3-Fc addition
Immunostaining
LAR damaging variants
b
LAR
WT
P381L
P381S
R388H
V389M
P417L
CD4
LAR
NGL-3-Fc
40 μm
c
Bound NGL3
Normalized bound NGL-3-Fc
****
***
****
****
LAR WT
LAR P381L
LAR P381S
LAR R388H
LAR V389M
LAR P417L
CD4
d
Surface LAR Expression
Normalized Surface LAR (AU)
****
****
****
*
****
LAR WT
LAR P381L
LAR P381S
LAR R388H
LAR V389M
LAR P417L
e
LAR variant transfection
HEK293/neuron co-culture
LAR
NGL-3
PSD95 clusters
f
LAR
WT
P381L
P381S
R388H
V389M
P417L
CD4
LAR/CD4
PSD95
Merge
10 μm
g
Artificial synapse formation
PSD95 (AU)
****
**
****
**
LAR WT
LAR P381L
LAR P381S
LAR R388H
LAR V389M
LAR P417L
CD4

### Figure 5

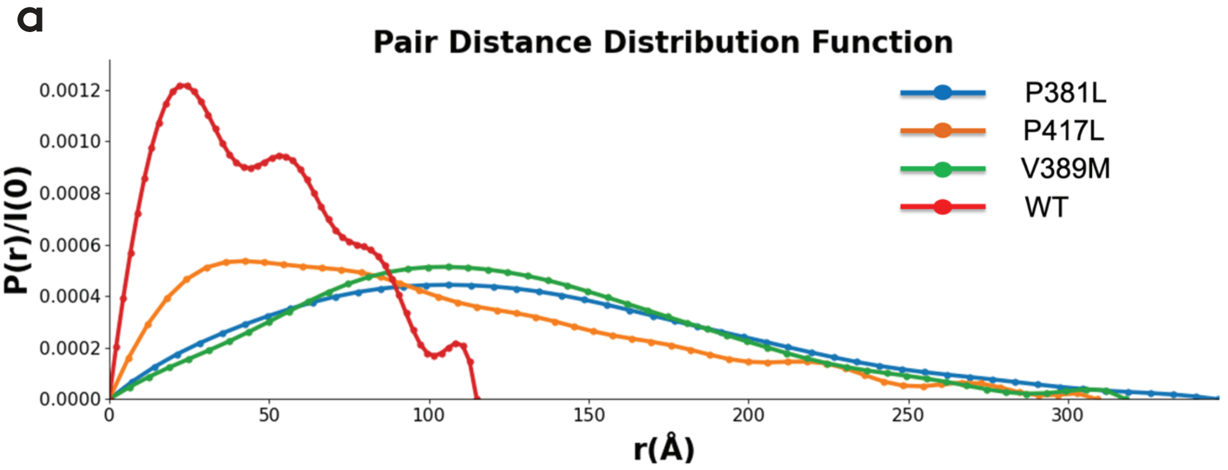


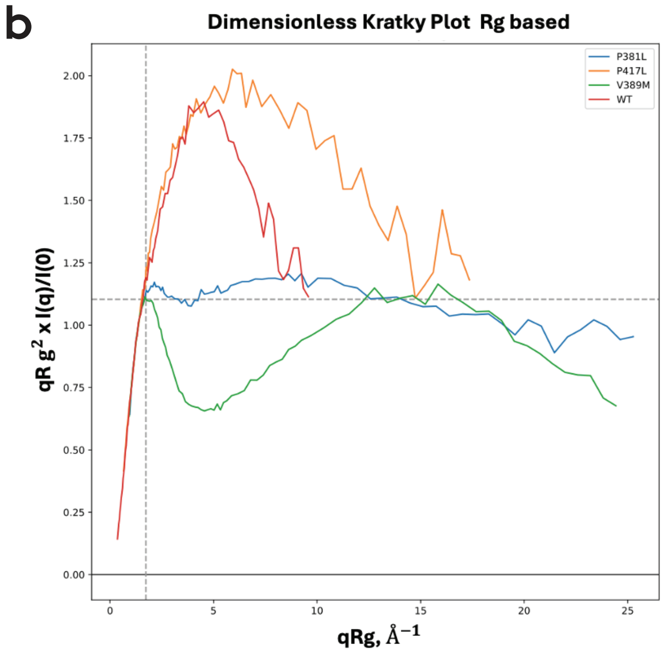


c

| Variant | Concentration (mg/ml) | I(0) | MW (kDa) | Rg (Å) | Dmax (Å) |
|---|---|---|---|---|---|
| Wild Type | 1.25 | 0.13 | 41 | 36 (±0.68) | 115 |
| P381L | 1.60 | 0.82 | 587 | 97 (±0.75) | 347 |
| V389M | 1.25 | 0.87 | 386 | 93 (±0.56) | 318 |
| P417L | 1.25 | 0.18 | 211 | 66 (±1.39) | 309 |

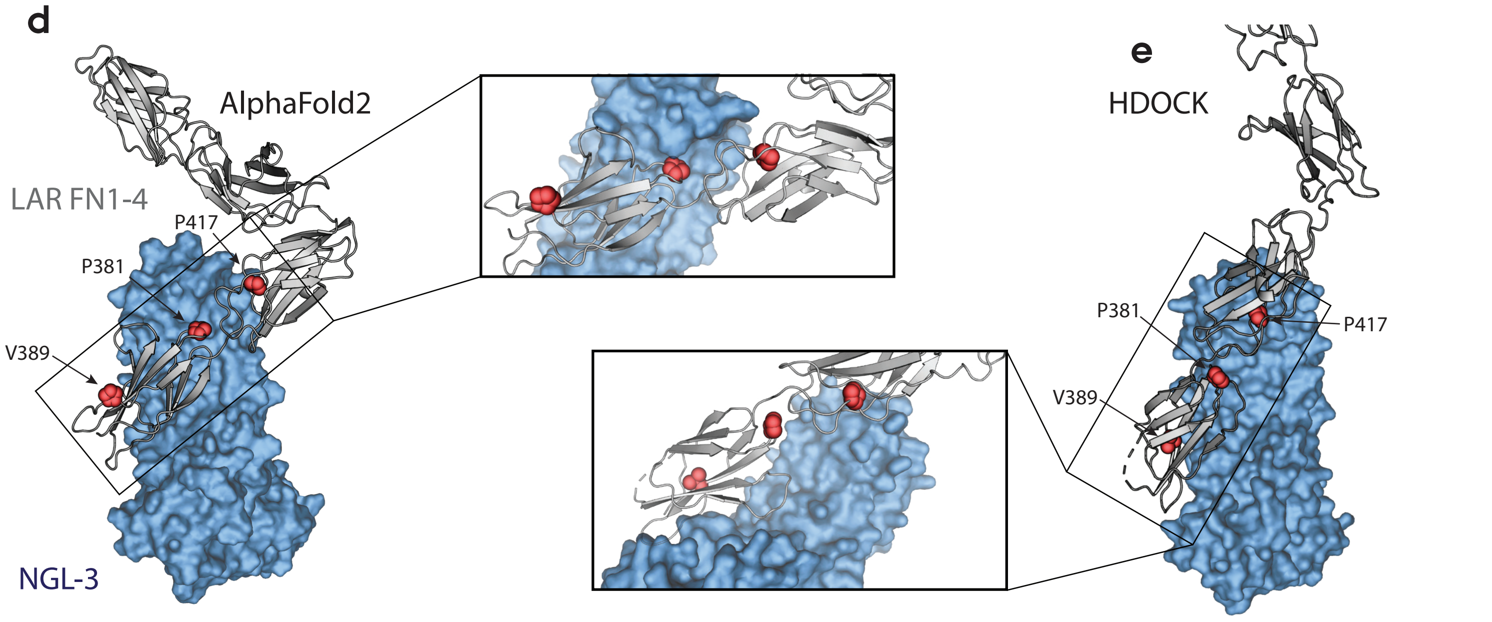

### Figure 6

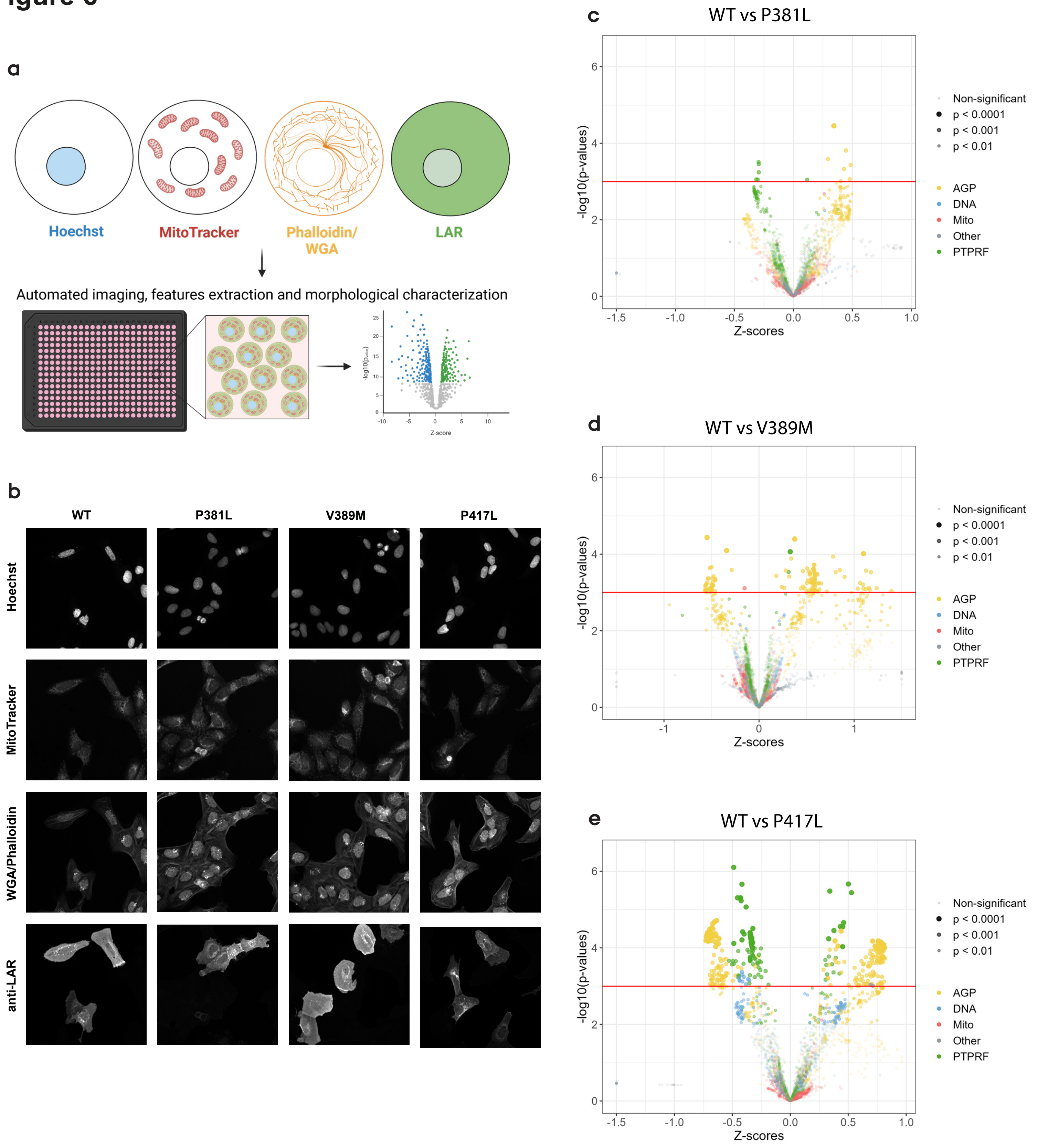
a
Hoechst
MitoTracker
Phalloidin/
WGA
LAR
Automated imaging, features extraction and morphological characterization
-log10(pvalue)
Z-score
b
WT
P381L
V389M
P417L
Hoechst
MitoTracker
WGA/Phalloidin
anti-LAR
c
WT vs P381L
-log10(p-values)
Z-scores
Non-significant
p < 0.0001
p < 0.001
p < 0.01
AGP
DNA
Mito
Other
PTPRF
d
WT vs V389M
-log10(p-values)
Z-scores
Non-significant
p < 0.0001
p < 0.001
p < 0.01
AGP
DNA
Mito
Other
PTPRF
e
WT vs P417L
-log10(p-values)
Z-scores
Non-significant
p < 0.0001
p < 0.001
p < 0.01
AGP
DNA
Mito
Other
PTPRF
