## Supplementary material for "Rare missense variants of the leukocyte common antigen related receptor (LAR) display reduced activity in transcellular adhesion and synapse formation": Analysis code

```
CellProfiler Pipeline: http://www.cellprofiler.org
Version:5
DateRevision:424
GitHash:
ModuleCount:40
HasImagePlaneDetails:False

LoadData:[module_num:1|svn_version:'Unknown'|variable_revision_number:6|show_window:False|notes:[]|batch_state:array([], dtype=uint8)|enabled:True|wants_pause:False]
    Input data file location:Elsewhere...|D:\\load_data_csv\\2023_07_18_Batch2\\230625_U2OS_PTPRF_bin1_21field_plate1
    Name of the file:load_data_with_illum.csv
    Load images based on this data?:Yes
    Base image location:None|
    Process just a range of rows?:No
    Rows to process:1,100000
    Group images by metadata?:Yes
    Select metadata tags for grouping:Plate,Well,Site
    Rescale intensities?:Yes

MeasureImageQuality:[module_num:2|svn_version:'Unknown'|variable_revision_number:6|show_window:False|notes:[]|batch_state:array([], dtype=uint8)|enabled:True|wants_pause:True]
    Calculate metrics for which images?:Select...
    Image count:1
    Scale count:4
    Threshold count:1
    Select the images to measure:OrigAGP, OrigBrightfield, OrigDNA, OrigMarker, OrigMito
    Include the image rescaling value?:Yes
    Calculate blur metrics?:Yes
    Spatial scale for blur measurements:5
    Spatial scale for blur measurements:10
    Spatial scale for blur measurements:20
    Spatial scale for blur measurements:50
    Calculate saturation metrics?:Yes
    Calculate intensity metrics?:Yes
    Calculate thresholds?:No
    Use all thresholding methods?:Yes
    Select a thresholding method:Otsu
    Typical fraction of the image covered by objects:0.1
    Two-class or three-class thresholding?:Two classes
    Minimize the weighted variance or the entropy?:Weighted variance
    Assign pixels in the middle intensity class to the foreground or the background?:Foreground

CorrectIlluminationApply:[module_num:3|svn_version:'Unknown'|variable_revision_number:5|show_window:False|notes:[]|batch_state:array([], dtype=uint8)|enabled:True|wants_pause:False]
    Select the input image:OrigDNA
    Name the output image:DNA
    Select the illumination function:IllumDNA
    Select how the illumination function is applied:Divide
    Select the input image:OrigMarker
    Name the output image:Marker
    Select the illumination function:IllumMarker
```

Select how the illumination function is applied:Divide
Select the input image:OrigAGP
Name the output image:AGP
Select the illumination function:IllumAGP
Select how the illumination function is applied:Divide
Select the input image:OrigBrightfield
Name the output image:Brightfield
Select the illumination function:IllumBrightfield
Select how the illumination function is applied:Divide
Select the input image:OrigMito
Name the output image:Mito
Select the illumination function:IllumMito
Select how the illumination function is applied:Divide
Set output image values less than 0 equal to 0?:Yes
Set output image values greater than 1 equal to 1?:Yes

IdentifyPrimaryObjects:
[module_num:4|svn_version:'Unknown'|variable_revision_number:15|show_window:False|notes:
['']|batch_state:array(b'', dtype='|S1')|enabled:True|wants_pause:False]
Select the input image:DNA
Name the primary objects to be identified:Nuclei
Typical diameter of objects, in pixel units (Min,Max):20,200
Discard objects outside the diameter range?:Yes
Discard objects touching the border of the image?:Yes
Method to distinguish clumped objects:Shape
Method to draw dividing lines between clumped objects:Shape
Size of smoothing filter:10
Suppress local maxima that are closer than this minimum allowed distance:8
Speed up by using lower-resolution image to find local maxima?:No
Fill holes in identified objects?:After declumping only
Automatically calculate size of smoothing filter for declumping?:Yes
Automatically calculate minimum allowed distance between local maxima?:Yes
Handling of objects if excessive number of objects identified:Continue
Maximum number of objects:500
Use advanced settings?:Yes
Threshold setting version:12
Threshold strategy:Global
Thresholding method:Minimum Cross-Entropy
Threshold smoothing scale:1
Threshold correction factor:1
Lower and upper bounds on threshold:0.01,1
Manual threshold:0.0
Select the measurement to threshold with:None
Two-class or three-class thresholding?:Three classes
Log transform before thresholding?:No
Assign pixels in the middle intensity class to the foreground or the background?:Background
Size of adaptive window:100
Lower outlier fraction:0.05
Upper outlier fraction:0.05
Averaging method:Mean
Variance method:Standard deviation
### of deviations:2
Thresholding method:Otsu

IdentifySecondaryObjects:
[module_num:5|svn_version:'Unknown'|variable_revision_number:10|show_window:False|notes:
['']|batch_state:array(b'', dtype='|S1')|enabled:True|wants_pause:False]
    Select the input objects:Nuclei
    Name the objects to be identified:Cells_small
    Select the method to identify the secondary objects:Watershed - Image
    Select the input image:Mito
    Number of pixels by which to expand the primary objects:10
    Regularization factor:0.01
    Discard secondary objects touching the border of the image?:No
    Discard the associated primary objects?:No
    Name the new primary objects:FilteredNuclei
    Fill holes in identified objects?:Yes
    Threshold setting version:12
    Threshold strategy:Global
    Thresholding method:Otsu
    Threshold smoothing scale:0
    Threshold correction factor:0.8
    Lower and upper bounds on threshold:0.015,.06
    Manual threshold:0.0
    Select the measurement to threshold with:None
    Two-class or three-class thresholding?:Three classes
    Log transform before thresholding?:Yes
    Assign pixels in the middle intensity class to the foreground or the background?:Foreground
    Size of adaptive window:100
    Lower outlier fraction:0.05
    Upper outlier fraction:0.05
    Averaging method:Mean
    Variance method:Standard deviation
    # of deviations:2
    Thresholding method:Otsu

IdentifySecondaryObjects:
[module_num:6|svn_version:'Unknown'|variable_revision_number:10|show_window:False|notes:[]|batch_state:array([],
dtype=uint8)|enabled:True|wants_pause:False]
    Select the input objects:Cells_small
    Name the objects to be identified:Cells
    Select the method to identify the secondary objects:Propagation
    Select the input image:AGP
    Number of pixels by which to expand the primary objects:10
    Regularization factor:0.0001
    Discard secondary objects touching the border of the image?:No
    Discard the associated primary objects?:No
    Name the new primary objects:FilteredNuclei
    Fill holes in identified objects?:Yes
    Threshold setting version:12
    Threshold strategy:Global
    Thresholding method:Otsu
    Threshold smoothing scale:1.3488
    Threshold correction factor:0.6
    Lower and upper bounds on threshold:0.002,1.0
    Manual threshold:0.0
    Select the measurement to threshold with:None
    Two-class or three-class thresholding?:Three classes

Log transform before thresholding?:Yes
Assign pixels in the middle intensity class to the foreground or the background?:Foreground
Size of adaptive window:50
Lower outlier fraction:0.05
Upper outlier fraction:0.05
Averaging method:Mean
Variance method:Standard deviation
### of deviations:2.0
Thresholding method:Minimum Cross-Entropy

IdentifyTertiaryObjects:[module_num:7|svn_version:'Unknown'|variable_revision_number:3|show_window:False|notes:[]|batch_state:array(b'', dtype='|S1')|enabled:True|wants_pause:False]
Select the larger identified objects:Cells
Select the smaller identified objects:Nuclei
Name the tertiary objects to be identified:Cytoplasm
Shrink smaller object prior to subtraction?:Yes

Threshold:[module_num:8|svn_version:'Unknown'|variable_revision_number:12|show_window:False|notes:[]|batch_state:array([], dtype=uint8)|enabled:True|wants_pause:False]
Select the input image:Marker
Name the output image:ThresholdMarker
Threshold strategy:Global
Thresholding method:Minimum Cross-Entropy
Threshold smoothing scale:0.0
Threshold correction factor:1.0
Lower and upper bounds on threshold:0.007,1.0
Manual threshold:0.0
Select the measurement to threshold with:None
Two-class or three-class thresholding?:Two classes
Log transform before thresholding?:Yes
Assign pixels in the middle intensity class to the foreground or the background?:Foreground
Size of adaptive window:50
Lower outlier fraction:0.05
Upper outlier fraction:0.05
Averaging method:Mean
Variance method:Standard deviation
### of deviations:2.0
Thresholding method:Minimum Cross-Entropy

MaskObjects:[module_num:9|svn_version:'Unknown'|variable_revision_number:3|show_window:False|notes:[]|batch_state:array([], dtype=uint8)|enabled:True|wants_pause:False]
Select objects to be masked:Cells
Name the masked objects:TransfectedCells
Mask using a region defined by other objects or by binary image?:Image
Select the masking object:None
Select the masking image:ThresholdMarker
Handling of objects that are partially masked:Remove depending on overlap
Fraction of object that must overlap:0.5
Numbering of resulting objects:Renumber
Invert the mask?:No

MaskObjects:[module_num:10|svn_version:'Unknown'|variable_revision_number:3|show_window:False|notes:[]|batch_state:array([], dtype=uint8)|enabled:True|wants_pause:False]
Select objects to be masked:Cells

Name the masked objects:UntransfectedCells
Mask using a region defined by other objects or by binary image?:Objects
Select the masking object:TransfectedCells
Select the masking image:ThresholdMarker
Handling of objects that are partially masked:Keep
Fraction of object that must overlap:0.5
Numbering of resulting objects:Renumber
Invert the mask?:Yes

MaskObjects:[module_num:11|svn_version:'Unknown'|variable_revision_number:3|show_window:False|notes:[]|batch_state:array([], dtype=uint8)|enabled:True|wants_pause:False]
Select objects to be masked:Nuclei
Name the masked objects:UntransfectedNuclei
Mask using a region defined by other objects or by binary image?:Objects
Select the masking object:TransfectedCells
Select the masking image:ThresholdMarker
Handling of objects that are partially masked:Keep
Fraction of object that must overlap:0.5
Numbering of resulting objects:Renumber
Invert the mask?:Yes

MaskObjects:[module_num:12|svn_version:'Unknown'|variable_revision_number:3|show_window:False|notes:[]|batch_state:array([], dtype=uint8)|enabled:True|wants_pause:False]
Select objects to be masked:Cytoplasm
Name the masked objects:UntransfectedCytoplasm
Mask using a region defined by other objects or by binary image?:Objects
Select the masking object:TransfectedCells
Select the masking image:ThresholdMarker
Handling of objects that are partially masked:Keep
Fraction of object that must overlap:0.5
Numbering of resulting objects:Renumber
Invert the mask?:Yes

MaskObjects:[module_num:13|svn_version:'Unknown'|variable_revision_number:3|show_window:False|notes:[]|batch_state:array([], dtype=uint8)|enabled:True|wants_pause:False]
Select objects to be masked:Nuclei
Name the masked objects:TransfectedNuclei
Mask using a region defined by other objects or by binary image?:Objects
Select the masking object:TransfectedCells
Select the masking image:ThresholdMarker
Handling of objects that are partially masked:Keep
Fraction of object that must overlap:0.5
Numbering of resulting objects:Renumber
Invert the mask?:No

MaskObjects:[module_num:14|svn_version:'Unknown'|variable_revision_number:3|show_window:False|notes:[]|batch_state:array([], dtype=uint8)|enabled:True|wants_pause:False]
Select objects to be masked:Cytoplasm
Name the masked objects:TransfectedCytoplasm
Mask using a region defined by other objects or by binary image?:Objects
Select the masking object:TransfectedCells
Select the masking image:ThresholdMarker
Handling of objects that are partially masked:Keep
Fraction of object that must overlap:0.5

Numbering of resulting objects:Renumber
Invert the mask?:No

MeasureColocalization:
[module_num:15|svn_version:'Unknown'|variable_revision_number:5|show_window:False|notes:
['']|batch_state:array(b'', dtype='|S1')|enabled:True|wants_pause:True]
Select images to measure:AGP, Brightfield, DNA, Marker, Mito
Set threshold as percentage of maximum intensity for the images:15.0
Select where to measure correlation:Within objects
Select objects to measure:TransfectedCells, TransfectedCytoplasm, TransfectedNuclei, UntransfectedCells, UntransfectedCytoplasm, UntransfectedNuclei
Run all metrics?:Yes
Calculate correlation and slope metrics?:Yes
Calculate the Manders coefficients?:Yes
Calculate the Rank Weighted Colocalization coefficients?:Yes
Calculate the Overlap coefficients?:Yes
Calculate the Manders coefficients using Costes auto threshold?:No
Method for Costes thresholding:Faster

MeasureGranularity:[module_num:16|svn_version:'Unknown'|variable_revision_number:4|show_window:False|notes:
[]|batch_state:array(b'', dtype='|S1')|enabled:True|wants_pause:False]
Select images to measure:AGP, Brightfield, DNA, Marker, Mito
Measure within objects?:Yes
Select objects to measure:TransfectedCells, TransfectedCytoplasm, TransfectedNuclei, UntransfectedCells, UntransfectedCytoplasm, UntransfectedNuclei
Subsampling factor for granularity measurements:0.25
Subsampling factor for background reduction:0.25
Radius of structuring element:10
Range of the granular spectrum:16

MeasureObjectIntensity:
[module_num:17|svn_version:'Unknown'|variable_revision_number:4|show_window:False|notes:[]|batch_state:array(b'', dtype='|S1')|enabled:True|wants_pause:False]
Select images to measure:AGP, Brightfield, DNA, Marker, Mito
Select objects to measure:TransfectedCells, TransfectedCytoplasm, TransfectedNuclei, UntransfectedCells, UntransfectedCytoplasm, UntransfectedNuclei

MeasureObjectNeighbors:
[module_num:18|svn_version:'Unknown'|variable_revision_number:3|show_window:False|notes:[]|batch_state:array(b'', dtype='|S1')|enabled:False|wants_pause:False]
Select objects to measure:TransfectedCells
Select neighboring objects to measure:TransfectedCells
Method to determine neighbors:Within a specified distance
Neighbor distance:10
Consider objects discarded for touching image border?:Yes
Retain the image of objects colored by numbers of neighbors?:No
Name the output image:ObjectNeighborCount
Select colormap:Default
Retain the image of objects colored by percent of touching pixels?:No
Name the output image:PercentTouching
Select colormap:Default

MeasureObjectNeighbors:
[module_num:19|svn_version:'Unknown'|variable_revision_number:3|show_window:False|notes:[]|batch_state:array(b'',

dtype='|S1')|enabled:False|wants_pause:False]
    Select objects to measure:TransfectedNuclei
    Select neighboring objects to measure:TransfectedNuclei
    Method to determine neighbors:Within a specified distance
    Neighbor distance:2
    Consider objects discarded for touching image border?:Yes
    Retain the image of objects colored by numbers of neighbors?:No
    Name the output image:ObjectNeighborCount
    Select colormap:Default
    Retain the image of objects colored by percent of touching pixels?:No
    Name the output image:PercentTouching
    Select colormap:Default

MeasureObjectNeighbors:
[module_num:20|svn_version:'Unknown'|variable_revision_number:3|show_window:False|notes:[]|batch_state:array(b'', dtype='|S1')|enabled:False|wants_pause:False]
    Select objects to measure:TransfectedCells
    Select neighboring objects to measure:TransfectedCells
    Method to determine neighbors:Adjacent
    Neighbor distance:5
    Consider objects discarded for touching image border?:Yes
    Retain the image of objects colored by numbers of neighbors?:No
    Name the output image:ObjectNeighborCount
    Select colormap:Default
    Retain the image of objects colored by percent of touching pixels?:No
    Name the output image:PercentTouching
    Select colormap:Default

MeasureObjectNeighbors:
[module_num:21|svn_version:'Unknown'|variable_revision_number:3|show_window:False|notes:[]|batch_state:array([], dtype=uint8)|enabled:False|wants_pause:False]
    Select objects to measure:UntransfectedCells
    Select neighboring objects to measure:UntransfectedCells
    Method to determine neighbors:Within a specified distance
    Neighbor distance:10
    Consider objects discarded for touching image border?:Yes
    Retain the image of objects colored by numbers of neighbors?:No
    Name the output image:ObjectNeighborCount
    Select colormap:Default
    Retain the image of objects colored by percent of touching pixels?:No
    Name the output image:PercentTouching
    Select colormap:Default

MeasureObjectNeighbors:
[module_num:22|svn_version:'Unknown'|variable_revision_number:3|show_window:False|notes:[]|batch_state:array([], dtype=uint8)|enabled:False|wants_pause:False]
    Select objects to measure:UntransfectedNuclei
    Select neighboring objects to measure:UntransfectedNuclei
    Method to determine neighbors:Within a specified distance
    Neighbor distance:2
    Consider objects discarded for touching image border?:Yes
    Retain the image of objects colored by numbers of neighbors?:No
    Name the output image:ObjectNeighborCount
    Select colormap:Default

Retain the image of objects colored by percent of touching pixels?:No
Name the output image:PercentTouching
Select colormap:Default

MeasureObjectNeighbors:
[module_num:23|svn_version:'Unknown'|variable_revision_number:3|show_window:False|notes:[]|batch_state:array([], dtype=uint8)|enabled:False|wants_pause:False]
Select objects to measure:UntransfectedCells
Select neighboring objects to measure:UntransfectedCells
Method to determine neighbors:Adjacent
Neighbor distance:5
Consider objects discarded for touching image border?:Yes
Retain the image of objects colored by numbers of neighbors?:No
Name the output image:ObjectNeighborCount
Select colormap:Default
Retain the image of objects colored by percent of touching pixels?:No
Name the output image:PercentTouching
Select colormap:Default

MeasureObjectIntensityDistribution:
[module_num:24|svn_version:'Unknown'|variable_revision_number:6|show_window:False|notes:[]|batch_state:array([], dtype=uint8)|enabled:True|wants_pause:False]
Select images to measure:AGP, Brightfield, DNA, Marker, Mito
Hidden:6
Hidden:1
Hidden:0
Calculate intensity Zernikes?:None
Maximum zernike moment:9
Select objects to measure:TransfectedNuclei
Object to use as center?:These objects
Select objects to use as centers:None
Select objects to measure:UntransfectedNuclei
Object to use as center?:These objects
Select objects to use as centers:None
Select objects to measure:TransfectedCytoplasm
Object to use as center?:These objects
Select objects to use as centers:None
Select objects to measure:UntransfectedCytoplasm
Object to use as center?:These objects
Select objects to use as centers:None
Select objects to measure:TransfectedCells
Object to use as center?:These objects
Select objects to use as centers:None
Select objects to measure:UntransfectedCells
Object to use as center?:These objects
Select objects to use as centers:None
Scale the bins?:Yes
Number of bins:4
Maximum radius:100

MeasureObjectSizeShape:
[module_num:25|svn_version:'Unknown'|variable_revision_number:3|show_window:False|notes:[]|batch_state:array(b'', dtype='|S1')|enabled:True|wants_pause:False]
Select object sets to measure:TransfectedCells, TransfectedCytoplasm, TransfectedNuclei, UntransfectedCells,

UntransfectedCytoplasm, UntransfectedNuclei
Calculate the Zernike features?:Yes
Calculate the advanced features?:No

MeasureTexture:[module_num:26|svn_version:'Unknown'|variable_revision_number:7|show_window:False|notes:[]|batch_state:array(b'', dtype='|S1')|enabled:True|wants_pause:False]
Select images to measure:AGP, Brightfield, DNA, Marker, Mito
Select objects to measure:TransfectedCells, TransfectedCytoplasm, TransfectedNuclei, UntransfectedCells, UntransfectedCytoplasm, UntransfectedNuclei
Enter how many gray levels to measure the texture at:256
Hidden:4
Measure whole images or objects?:Both
Texture scale to measure:3
Texture scale to measure:5
Texture scale to measure:10
Texture scale to measure:20

MeasureImageIntensity:[module_num:27|svn_version:'Unknown'|variable_revision_number:4|show_window:False|notes:['Measures the ntensity of the whole image (AGP, BFHigh, BFLow, Brightfield, DNA, ER, Mito, RNA) as well as the intensity of the image only in regions without cells (* _BackgroundOnly).']|batch_state:array([], dtype=uint8)|enabled:True|wants_pause:False]
Select images to measure:AGP, Brightfield, DNA, Marker, Mito
Measure the intensity only from areas enclosed by objects?:No
Select input object sets:
Calculate custom percentiles:No
Specify percentiles to measure:10,90

OverlayOutlines:[module_num:28|svn_version:'Unknown'|variable_revision_number:4|show_window:False|notes:[]|batch_state:array([], dtype=uint8)|enabled:True|wants_pause:False]
Display outlines on a blank image?:Yes
Select image on which to display outlines:None
Name the output image:TransfectedNucleiOutlines
Outline display mode:Grayscale
Select method to determine brightness of outlines:Max of image
How to outline:Inner
Select outline color:Red
Select objects to display:TransfectedNuclei

OverlayOutlines:[module_num:29|svn_version:'Unknown'|variable_revision_number:4|show_window:False|notes:[]|batch_state:array([], dtype=uint8)|enabled:True|wants_pause:False]
Display outlines on a blank image?:Yes
Select image on which to display outlines:None
Name the output image:TransfectedCellOutlines
Outline display mode:Grayscale
Select method to determine brightness of outlines:Max of image
How to outline:Inner
Select outline color:Red
Select objects to display:TransfectedCells

OverlayOutlines:[module_num:30|svn_version:'Unknown'|variable_revision_number:4|show_window:False|notes:[]|batch_state:array([], dtype=uint8)|enabled:True|wants_pause:False]
Display outlines on a blank image?:Yes
Select image on which to display outlines:ThresholdMarker

Name the output image:TransfectedCytoplasmOutlines
Outline display mode:Color
Select method to determine brightness of outlines:Max of image
How to outline:Inner
Select outline color:Red
Select objects to display:TransfectedCytoplasm

OverlayOutlines:[module_num:31|svn_version:'Unknown'|variable_revision_number:4|show_window:False|notes:[]|batch_state:array([], dtype=uint8)|enabled:True|wants_pause:False]
Display outlines on a blank image?:Yes
Select image on which to display outlines:None
Name the output image:UnTransfectedNucleiOutlines
Outline display mode:Grayscale
Select method to determine brightness of outlines:Max of image
How to outline:Inner
Select outline color:Red
Select objects to display:UntransfectedNuclei

OverlayOutlines:[module_num:32|svn_version:'Unknown'|variable_revision_number:4|show_window:False|notes:[]|batch_state:array([], dtype=uint8)|enabled:True|wants_pause:False]
Display outlines on a blank image?:Yes
Select image on which to display outlines:None
Name the output image:UnTransfectedCellOutlines
Outline display mode:Grayscale
Select method to determine brightness of outlines:Max of image
How to outline:Inner
Select outline color:Red
Select objects to display:UntransfectedCells

OverlayOutlines:[module_num:33|svn_version:'Unknown'|variable_revision_number:4|show_window:False|notes:[]|batch_state:array([], dtype=uint8)|enabled:True|wants_pause:False]
Display outlines on a blank image?:Yes
Select image on which to display outlines:ThresholdMarker
Name the output image:UnTransfectedCytoplasmOutlines
Outline display mode:Color
Select method to determine brightness of outlines:Max of image
How to outline:Inner
Select outline color:Red
Select objects to display:UntransfectedCytoplasm

SaveImages:[module_num:34|svn_version:'Unknown'|variable_revision_number:16|show_window:False|notes:[]|batch_state:array([], dtype=uint8)|enabled:True|wants_pause:False]
Select the type of image to save:Image
Select the image to save:TransfectedNucleiOutlines
Select method for constructing file names:Single name
Select image name for file prefix:OrigDNA
Enter single file name:\g<Well>_\g<Site>--transfected_nuclei_outlines
Number of digits:4
Append a suffix to the image file name?:Yes
Text to append to the image name:_nuclei
Saved file format:png
Output file location:Default Output Folder|\g<Plate>-\g<Well>-\g<Site>/outlines
Image bit depth:8-bit integer
Overwrite existing files without warning?:Yes

    When to save:Every cycle
    Record the file and path information to the saved image?:Yes
    Create subfolders in the output folder?:No
    Base image folder:Default Input Folder
    How to save the series:T (Time)
    Save with lossless compression?:No

SaveImages:[module_num:35|svn_version:'Unknown'|variable_revision_number:16|show_window:False|notes:[]|batch_state:array([], dtype=uint8)|enabled:True|wants_pause:False]
    Select the type of image to save:Image
    Select the image to save:TransfectedCellOutlines
    Select method for constructing file names:Single name
    Select image name for file prefix:OrigDNA
    Enter single file name:\g<Well>_\g<Site>--transfected_cell_outlines
    Number of digits:4
    Append a suffix to the image file name?:Yes
    Text to append to the image name:_nuclei
    Saved file format:png
    Output file location:Default Output Folder|\g<Plate>-\g<Well>-\g<Site>/outlines
    Image bit depth:8-bit integer
    Overwrite existing files without warning?:Yes
    When to save:Every cycle
    Record the file and path information to the saved image?:Yes
    Create subfolders in the output folder?:No
    Base image folder:Default Input Folder
    How to save the series:T (Time)
    Save with lossless compression?:No

SaveImages:[module_num:36|svn_version:'Unknown'|variable_revision_number:16|show_window:False|notes:[]|batch_state:array([], dtype=uint8)|enabled:True|wants_pause:False]
    Select the type of image to save:Image
    Select the image to save:TransfectedCytoplasmOutlines
    Select method for constructing file names:Single name
    Select image name for file prefix:OrigDNA
    Enter single file name:\g<Well>_\g<Site>--transfected_cytoplasm_outlines
    Number of digits:4
    Append a suffix to the image file name?:Yes
    Text to append to the image name:_nuclei
    Saved file format:png
    Output file location:Default Output Folder|\g<Plate>-\g<Well>-\g<Site>/outlines
    Image bit depth:8-bit integer
    Overwrite existing files without warning?:Yes
    When to save:Every cycle
    Record the file and path information to the saved image?:Yes
    Create subfolders in the output folder?:No
    Base image folder:Default Input Folder
    How to save the series:T (Time)
    Save with lossless compression?:No

SaveImages:[module_num:37|svn_version:'Unknown'|variable_revision_number:16|show_window:False|notes:[]|batch_state:array([], dtype=uint8)|enabled:True|wants_pause:False]
    Select the type of image to save:Image
    Select the image to save:UnTransfectedNucleiOutlines
    Select method for constructing file names:Single name

Select image name for file prefix:OrigDNA
Enter single file name:\g<Well>_\g<Site>--untransfected_nuclei_outlines
Number of digits:4
Append a suffix to the image file name?:Yes
Text to append to the image name:_nuclei
Saved file format:png
Output file location:Default Output Folder|\g<Plate>-\g<Well>-\g<Site>/outlines
Image bit depth:8-bit integer
Overwrite existing files without warning?:No
When to save:Every cycle
Record the file and path information to the saved image?:Yes
Create subfolders in the output folder?:No
Base image folder:Default Input Folder
How to save the series:T (Time)
Save with lossless compression?:No

SaveImages:[module_num:38|svn_version:'Unknown'|variable_revision_number:16|show_window:False|notes:[]|batch_state:array([], dtype=uint8)|enabled:True|wants_pause:False]
Select the type of image to save:Image
Select the image to save:UnTransfectedCellOutlines
Select method for constructing file names:Single name
Select image name for file prefix:OrigDNA
Enter single file name:\g<Well>_\g<Site>--untransfected_cell_outlines
Number of digits:4
Append a suffix to the image file name?:Yes
Text to append to the image name:_nuclei
Saved file format:png
Output file location:Default Output Folder|\g<Plate>-\g<Well>-\g<Site>/outlines
Image bit depth:8-bit integer
Overwrite existing files without warning?:Yes
When to save:Every cycle
Record the file and path information to the saved image?:Yes
Create subfolders in the output folder?:No
Base image folder:Default Input Folder
How to save the series:T (Time)
Save with lossless compression?:No

SaveImages:[module_num:39|svn_version:'Unknown'|variable_revision_number:16|show_window:False|notes:[]|batch_state:array([], dtype=uint8)|enabled:True|wants_pause:False]
Select the type of image to save:Image
Select the image to save:UnTransfectedCytoplasmOutlines
Select method for constructing file names:Single name
Select image name for file prefix:OrigDNA
Enter single file name:\g<Well>_\g<Site>--untransfected_cytoplasm_outlines
Number of digits:4
Append a suffix to the image file name?:Yes
Text to append to the image name:_nuclei
Saved file format:png
Output file location:Default Output Folder|\g<Plate>-\g<Well>-\g<Site>/outlines
Image bit depth:8-bit integer
Overwrite existing files without warning?:Yes
When to save:Every cycle
Record the file and path information to the saved image?:Yes
Create subfolders in the output folder?:No

Base image folder:Default Input Folder
How to save the series:T (Time)
Save with lossless compression?:No

ExportToSpreadsheet:
[module_num:40|svn_version:'Unknown'|variable_revision_number:13|show_window:False|notes:[''|batch_state:array([], dtype=uint8)|enabled:True|wants_pause:False]
Select the column delimiter:Comma (",")
Add image metadata columns to your object data file?:No
Add image file and folder names to your object data file?:No
Select the measurements to export:No
Calculate the per-image mean values for object measurements?:No
Calculate the per-image median values for object measurements?:No
Calculate the per-image standard deviation values for object measurements?:No
Output file location:Default Output Folder|\g<Plate>-\g<Well>-\g<Site>
Create a GenePattern GCT file?:No
Select source of sample row name:Metadata
Select the image to use as the identifier:None
Select the metadata to use as the identifier:None
Export all measurement types?:No
Press button to select measurements:Cells|AreaShape_Zernike_9_1,Cells|AreaShape_Zernike_9_3,Cells|AreaShape_Zernike_9_9,Cells|AreaShape_Zernike_9_5,Cells|AreaShape_Zernike_9_7,Cells|AreaShape_Zernike_8_6,Cells|AreaShape_Zernike_8_8,Cells|AreaShape_Zernike_8_4,Cells|AreaShape_Zernike_8_0,Cells|AreaShape_Zernike_8_2,Cells|AreaShape_Zernike_3_3,Cells|AreaShape_Zernike_3_1,Cells|AreaShape_Zernike_4_2,Cells|AreaShape_Zernike_4_0,Cells|AreaShape_Zernike_4_4,Cells|AreaShape_Zernike_2_2,Cells|AreaShape_Zernike_2_0,Cells|AreaShape_Zernike_5_5,Cells|AreaShape_Zernike_5_3,Cells|AreaShape_Zernike_5_1,Cells|AreaShape_Zernike_7_3,Cells|AreaShape_Zernike_7_1,Cells|AreaShape_Zernike_7_7,Cells|AreaShape_Zernike_7_5,Cells|AreaShape_Zernike_0_0,Cells|AreaShape_Zernike_1_1,Cells|AreaShape_Zernike_6_0,Cells|AreaShape_Zernike_6_4,Cells|AreaShape_Zernike_6_6,Cells|AreaShape_Zernike_6_2,Cells|AreaShape_MinorAxisLength,Cells|AreaShape_Solidity,Cells|AreaShape_Area,Cells|AreaShape_BoundingBoxMaximum_X,Cells|AreaShape_BoundingBoxMaximum_Y,Cells|AreaShape_MinFeretDiameter,Cells|AreaShape_MaxFeretDiameter,Cells|AreaShape_MedianRadius,Cells|AreaShape_BoundingBoxMinimum_Y,Cells|AreaShape_BoundingBoxMinimum_X,Cells|AreaShape_EquivalentDiameter,Cells|AreaShape_EulerNumber,Cells|AreaShape_FormFactor,Cells|AreaShape_Compactness,Cells|AreaShape_ConvexArea,Cells|AreaShape_MajorAxisLength,Cells|AreaShape_Perimeter,Cells|AreaShape_Center_Y,Cells|AreaShape_Center_X,Cells|AreaShape_Orientation,Cells|AreaShape_Extent,Cells|AreaShape_MaximumRadius,Cells|AreaShape_Eccentricity,Cells|AreaShape_BoundingBoxArea,Cells|AreaShape_MeanRadius,Cells|Texture_Contrast_Phalloidin_20_03_256,Cells|Texture_Contrast_Phalloidin_20_02_256,Cells|Texture_Contrast_Phalloidin_20_00_256,Cells|Texture_Contrast_Phalloidin_20_01_256,Cells|Texture_Contrast_Phalloidin_5_02_256,Cells|Texture_Contrast_Phalloidin_5_01_256,Cells|Texture_Contrast_Phalloidin_5_03_256,Cells|Texture_Contrast_Phalloidin_5_00_256,Cells|Texture_Contrast_Phalloidin_3_02_256,Cells|Texture_Contrast_Phalloidin_3_01_256,Cells|Texture_Contrast_Phalloidin_3_00_256,Cells|Texture_Contrast_Phalloidin_3_03_256,Cells|Texture_Contrast_Phalloidin_10_00_256,Cells|Texture_Contrast_Phalloidin_10_02_256,Cells|Texture_Contrast_Phalloidin_10_03_256,Cells|Texture_Contrast_Phalloidin_10_01_256,Cells|Texture_Contrast_DNA_3_01_256,Cells|Texture_Contrast_DNA_3_00_256,Cells|Texture_Contrast_DNA_3_03_256,Cells|Texture_Contrast_DNA_3_02_256,Cells|Texture_Contrast_DNA_20_03_256,Cells|Texture_Contrast_DNA_20_01_256,Cells|Texture_Contrast_DNA_20_02_256,Cells|Texture_Contrast_DNA_20_00_256,Cells|Texture_Contrast_DNA_10_02_256,Cells|Texture_Contrast_DNA_10_00_256,Cells|Texture_Contrast_DNA_10_01_256,Cells|Texture_Contrast_DNA_10_03_256,Cells|Texture_Contrast_DNA_5_00_256,Cells|Texture_Contrast_DNA_5_03_256,Cells|Texture_Contrast_DNA_5_01_256,Cells|Texture_Contrast_DNA_5_02_256,Cells|Texture_Contrast_WGA_10_03_256,Cells|Texture_Contrast_WGA_10_02_256,Cells|Texture_Contrast_WGA_10_01_256,Cells|Texture_Contrast_WGA_10_00_256,Cells|Texture_Contrast_WGA_3_03_256,Cells|Texture_Contrast_WGA_3_00_256,Cells|Texture_Contrast_WGA_3_02_256,Cells|Texture_Contrast_WGA_3_01_256,Cells|Texture_Contrast_WGA_20_01_256,Cells|Texture_Contrast_WGA_20_00_256,Cells|Texture_Contrast_WGA_20_02_256,Cells|Texture_Contrast_WGA_20_03_256,Cells|Texture_Contrast_WGA_5_01_256,Cells|Texture_Contrast_WGA_5_02_256,Cells|Texture_Contrast_WGA_5_03_256,Cells|Texture_Contrast_WGA_5_00_256,Cells|Texture_Contrast_Vinculin_3_03_256

,Cells|Texture_Contrast_Vinculin_3_02_256,Cells|Texture_Contrast_Vinculin_3_00_256,Cells|Texture_Contrast_Vinc
ulin_3_01_256,Cells|Texture_Contrast_Vinculin_10_03_256,Cells|Texture_Contrast_Vinculin_10_01_256,Cells|Textur
e_Contrast_Vinculin_10_00_256,Cells|Texture_Contrast_Vinculin_10_02_256,Cells|Texture_Contrast_Vinculin_20_0
0_256,Cells|Texture_Contrast_Vinculin_20_01_256,Cells|Texture_Contrast_Vinculin_20_02_256,Cells|Texture_Contr
ast_Vinculin_20_03_256,Cells|Texture_Contrast_Vinculin_5_01_256,Cells|Texture_Contrast_Vinculin_5_00_256,Cell
s|Texture_Contrast_Vinculin_5_02_256,Cells|Texture_Contrast_Vinculin_5_03_256,Cells|Texture_Correlation_Phalloi
din_10_02_256,Cells|Texture_Correlation_Phalloidin_10_00_256,Cells|Texture_Correlation_Phalloidin_10_01_256,Ce
lls|Texture_Correlation_Phalloidin_10_03_256,Cells|Texture_Correlation_Phalloidin_5_03_256,Cells|Texture_Correlat
ion_Phalloidin_5_02_256,Cells|Texture_Correlation_Phalloidin_5_01_256,Cells|Texture_Correlation_Phalloidin_5_00
_256,Cells|Texture_Correlation_Phalloidin_3_03_256,Cells|Texture_Correlation_Phalloidin_3_01_256,Cells|Texture_
Correlation_Phalloidin_3_02_256,Cells|Texture_Correlation_Phalloidin_3_00_256,Cells|Texture_Correlation_Phalloidi
n_20_02_256,Cells|Texture_Correlation_Phalloidin_20_03_256,Cells|Texture_Correlation_Phalloidin_20_01_256,Cell
s|Texture_Correlation_Phalloidin_20_00_256,Cells|Texture_Correlation_DNA_5_03_256,Cells|Texture_Correlation_D
NA_5_02_256,Cells|Texture_Correlation_DNA_5_00_256,Cells|Texture_Correlation_DNA_5_01_256,Cells|Texture_
Correlation_DNA_20_02_256,Cells|Texture_Correlation_DNA_20_01_256,Cells|Texture_Correlation_DNA_20_00_2
56,Cells|Texture_Correlation_DNA_20_03_256,Cells|Texture_Correlation_DNA_3_01_256,Cells|Texture_Correlation
_DNA_3_03_256,Cells|Texture_Correlation_DNA_3_00_256,Cells|Texture_Correlation_DNA_3_02_256,Cells|Textur
e_Correlation_DNA_10_01_256,Cells|Texture_Correlation_DNA_10_02_256,Cells|Texture_Correlation_DNA_10_00
_256,Cells|Texture_Correlation_DNA_10_03_256,Cells|Texture_Correlation_WGA_5_01_256,Cells|Texture_Correlati
on_WGA_5_00_256,Cells|Texture_Correlation_WGA_5_02_256,Cells|Texture_Correlation_WGA_5_03_256,Cells|Te
xture_Correlation_WGA_10_01_256,Cells|Texture_Correlation_WGA_10_02_256,Cells|Texture_Correlation_WGA_1
0_00_256,Cells|Texture_Correlation_WGA_10_03_256,Cells|Texture_Correlation_WGA_20_01_256,Cells|Texture_C
orrelation_WGA_20_03_256,Cells|Texture_Correlation_WGA_20_02_256,Cells|Texture_Correlation_WGA_20_00_2
56,Cells|Texture_Correlation_WGA_3_01_256,Cells|Texture_Correlation_WGA_3_03_256,Cells|Texture_Correlation_
WGA_3_00_256,Cells|Texture_Correlation_WGA_3_02_256,Cells|Texture_Correlation_Vinculin_10_03_256,Cells|Te
xture_Correlation_Vinculin_10_01_256,Cells|Texture_Correlation_Vinculin_10_00_256,Cells|Texture_Correlation_Vi
nculin_10_02_256,Cells|Texture_Correlation_Vinculin_20_03_256,Cells|Texture_Correlation_Vinculin_20_02_256,Ce
lls|Texture_Correlation_Vinculin_20_00_256,Cells|Texture_Correlation_Vinculin_20_01_256,Cells|Texture_Correlatio
n_Vinculin_3_00_256,Cells|Texture_Correlation_Vinculin_3_03_256,Cells|Texture_Correlation_Vinculin_3_01_256,C
ells|Texture_Correlation_Vinculin_3_02_256,Cells|Texture_Correlation_Vinculin_5_00_256,Cells|Texture_Correlation
_Vinculin_5_02_256,Cells|Texture_Correlation_Vinculin_5_01_256,Cells|Texture_Correlation_Vinculin_5_03_256,Ce
lls|Texture_InfoMeas2_Phalloidin_5_00_256,Cells|Texture_InfoMeas2_Phalloidin_5_02_256,Cells|Texture_InfoMeas2
_Phalloidin_5_01_256,Cells|Texture_InfoMeas2_Phalloidin_5_03_256,Cells|Texture_InfoMeas2_Phalloidin_10_01_25
6,Cells|Texture_InfoMeas2_Phalloidin_10_03_256,Cells|Texture_InfoMeas2_Phalloidin_10_00_256,Cells|Texture_Inf
oMeas2_Phalloidin_10_02_256,Cells|Texture_InfoMeas2_Phalloidin_20_01_256,Cells|Texture_InfoMeas2_Phalloidin
_20_02_256,Cells|Texture_InfoMeas2_Phalloidin_20_03_256,Cells|Texture_InfoMeas2_Phalloidin_20_00_256,Cells|T
exture_InfoMeas2_Phalloidin_3_02_256,Cells|Texture_InfoMeas2_Phalloidin_3_01_256,Cells|Texture_InfoMeas2_Ph
alloidin_3_03_256,Cells|Texture_InfoMeas2_Phalloidin_3_00_256,Cells|Texture_InfoMeas2_Vinculin_5_01_256,Cell
s|Texture_InfoMeas2_Vinculin_5_03_256,Cells|Texture_InfoMeas2_Vinculin_5_00_256,Cells|Texture_InfoMeas2_Vi
nculin_5_02_256,Cells|Texture_InfoMeas2_Vinculin_10_01_256,Cells|Texture_InfoMeas2_Vinculin_10_03_256,Cells
|Texture_InfoMeas2_Vinculin_10_02_256,Cells|Texture_InfoMeas2_Vinculin_10_00_256,Cells|Texture_InfoMeas2_V
inculin_3_01_256,Cells|Texture_InfoMeas2_Vinculin_3_00_256,Cells|Texture_InfoMeas2_Vinculin_3_02_256,Cells|T
exture_InfoMeas2_Vinculin_3_03_256,Cells|Texture_InfoMeas2_Vinculin_20_03_256,Cells|Texture_InfoMeas2_Vinc
ulin_20_02_256,Cells|Texture_InfoMeas2_Vinculin_20_01_256,Cells|Texture_InfoMeas2_Vinculin_20_00_256,Cells|
Texture_InfoMeas2_WGA_3_00_256,Cells|Texture_InfoMeas2_WGA_3_02_256,Cells|Texture_InfoMeas2_WGA_3_
01_256,Cells|Texture_InfoMeas2_WGA_3_03_256,Cells|Texture_InfoMeas2_WGA_10_02_256,Cells|Texture_InfoMe
as2_WGA_10_01_256,Cells|Texture_InfoMeas2_WGA_10_00_256,Cells|Texture_InfoMeas2_WGA_10_03_256,Cells
|Texture_InfoMeas2_WGA_5_03_256,Cells|Texture_InfoMeas2_WGA_5_01_256,Cells|Texture_InfoMeas2_WGA_5_
02_256,Cells|Texture_InfoMeas2_WGA_5_00_256,Cells|Texture_InfoMeas2_WGA_20_02_256,Cells|Texture_InfoMe
as2_WGA_20_01_256,Cells|Texture_InfoMeas2_WGA_20_03_256,Cells|Texture_InfoMeas2_WGA_20_00_256,Cells
|Texture_InfoMeas2_DNA_3_02_256,Cells|Texture_InfoMeas2_DNA_3_00_256,Cells|Texture_InfoMeas2_DNA_3_0
3_256,Cells|Texture_InfoMeas2_DNA_3_01_256,Cells|Texture_InfoMeas2_DNA_5_03_256,Cells|Texture_InfoMeas2
_DNA_5_01_256,Cells|Texture_InfoMeas2_DNA_5_00_256,Cells|Texture_InfoMeas2_DNA_5_02_256,Cells|Texture
_InfoMeas2_DNA_10_00_256,Cells|Texture_InfoMeas2_DNA_10_03_256,Cells|Texture_InfoMeas2_DNA_10_01_25

6,Cells|Texture_InfoMeas2_DNA_10_02_256,Cells|Texture_InfoMeas2_DNA_20_02_256,Cells|Texture_InfoMeas2_
DNA_20_00_256,Cells|Texture_InfoMeas2_DNA_20_01_256,Cells|Texture_InfoMeas2_DNA_20_03_256,Cells|Textu
re_Variance_Phalloidin_10_03_256,Cells|Texture_Variance_Phalloidin_10_02_256,Cells|Texture_Variance_Phalloidin
_10_00_256,Cells|Texture_Variance_Phalloidin_10_01_256,Cells|Texture_Variance_Phalloidin_20_03_256,Cells|Text
ure_Variance_Phalloidin_20_00_256,Cells|Texture_Variance_Phalloidin_20_02_256,Cells|Texture_Variance_Phalloidi
n_20_01_256,Cells|Texture_Variance_Phalloidin_3_03_256,Cells|Texture_Variance_Phalloidin_3_00_256,Cells|Textu
re_Variance_Phalloidin_3_02_256,Cells|Texture_Variance_Phalloidin_3_01_256,Cells|Texture_Variance_Phalloidin_5
_03_256,Cells|Texture_Variance_Phalloidin_5_01_256,Cells|Texture_Variance_Phalloidin_5_02_256,Cells|Texture_V
ariance_Phalloidin_5_00_256,Cells|Texture_Variance_WGA_20_01_256,Cells|Texture_Variance_WGA_20_03_256,C
ells|Texture_Variance_WGA_20_02_256,Cells|Texture_Variance_WGA_20_00_256,Cells|Texture_Variance_WGA_3
_02_256,Cells|Texture_Variance_WGA_3_01_256,Cells|Texture_Variance_WGA_3_03_256,Cells|Texture_Variance_
WGA_3_00_256,Cells|Texture_Variance_WGA_5_03_256,Cells|Texture_Variance_WGA_5_00_256,Cells|Texture_V
ariance_WGA_5_01_256,Cells|Texture_Variance_WGA_5_02_256,Cells|Texture_Variance_WGA_10_00_256,Cells|T
exture_Variance_WGA_10_02_256,Cells|Texture_Variance_WGA_10_01_256,Cells|Texture_Variance_WGA_10_03_
256,Cells|Texture_Variance_DNA_10_00_256,Cells|Texture_Variance_DNA_10_03_256,Cells|Texture_Variance_DN
A_10_02_256,Cells|Texture_Variance_DNA_10_01_256,Cells|Texture_Variance_DNA_20_02_256,Cells|Texture_Var
iance_DNA_20_01_256,Cells|Texture_Variance_DNA_20_00_256,Cells|Texture_Variance_DNA_20_03_256,Cells|Te
xture_Variance_DNA_3_03_256,Cells|Texture_Variance_DNA_3_01_256,Cells|Texture_Variance_DNA_3_00_256,C
ells|Texture_Variance_DNA_3_02_256,Cells|Texture_Variance_DNA_5_00_256,Cells|Texture_Variance_DNA_5_02_
256,Cells|Texture_Variance_DNA_5_03_256,Cells|Texture_Variance_DNA_5_01_256,Cells|Texture_Variance_Vincul
in_3_00_256,Cells|Texture_Variance_Vinculin_3_02_256,Cells|Texture_Variance_Vinculin_3_03_256,Cells|Texture_
Variance_Vinculin_3_01_256,Cells|Texture_Variance_Vinculin_5_01_256,Cells|Texture_Variance_Vinculin_5_03_25
6,Cells|Texture_Variance_Vinculin_5_02_256,Cells|Texture_Variance_Vinculin_5_00_256,Cells|Texture_Variance_Vi
nculin_10_03_256,Cells|Texture_Variance_Vinculin_10_00_256,Cells|Texture_Variance_Vinculin_10_02_256,Cells|T
exture_Variance_Vinculin_10_01_256,Cells|Texture_Variance_Vinculin_20_01_256,Cells|Texture_Variance_Vinculin
_20_02_256,Cells|Texture_Variance_Vinculin_20_00_256,Cells|Texture_Variance_Vinculin_20_03_256,Cells|Texture
_DifferenceVariance_Phalloidin_20_00_256,Cells|Texture_DifferenceVariance_Phalloidin_20_03_256,Cells|Texture_
DifferenceVariance_Phalloidin_20_01_256,Cells|Texture_DifferenceVariance_Phalloidin_20_02_256,Cells|Texture_Di
fferenceVariance_Phalloidin_10_02_256,Cells|Texture_DifferenceVariance_Phalloidin_10_00_256,Cells|Texture_Diff
erenceVariance_Phalloidin_10_01_256,Cells|Texture_DifferenceVariance_Phalloidin_10_03_256,Cells|Texture_Differ
enceVariance_Phalloidin_5_02_256,Cells|Texture_DifferenceVariance_Phalloidin_5_00_256,Cells|Texture_Difference
Variance_Phalloidin_5_01_256,Cells|Texture_DifferenceVariance_Phalloidin_5_03_256,Cells|Texture_DifferenceVari
ance_Phalloidin_3_01_256,Cells|Texture_DifferenceVariance_Phalloidin_3_02_256,Cells|Texture_DifferenceVariance
_Phalloidin_3_00_256,Cells|Texture_DifferenceVariance_Phalloidin_3_03_256,Cells|Texture_DifferenceVariance_Vin
culin_5_01_256,Cells|Texture_DifferenceVariance_Vinculin_5_00_256,Cells|Texture_DifferenceVariance_Vinculin_5
_03_256,Cells|Texture_DifferenceVariance_Vinculin_5_02_256,Cells|Texture_DifferenceVariance_Vinculin_20_01_2
56,Cells|Texture_DifferenceVariance_Vinculin_20_00_256,Cells|Texture_DifferenceVariance_Vinculin_20_03_256,C
ells|Texture_DifferenceVariance_Vinculin_20_02_256,Cells|Texture_DifferenceVariance_Vinculin_3_00_256,Cells|Te
xture_DifferenceVariance_Vinculin_3_03_256,Cells|Texture_DifferenceVariance_Vinculin_3_01_256,Cells|Texture_D
ifferenceVariance_Vinculin_3_02_256,Cells|Texture_DifferenceVariance_Vinculin_10_02_256,Cells|Texture_Differen
ceVariance_Vinculin_10_03_256,Cells|Texture_DifferenceVariance_Vinculin_10_00_256,Cells|Texture_DifferenceVa
riance_Vinculin_10_01_256,Cells|Texture_DifferenceVariance_WGA_5_02_256,Cells|Texture_DifferenceVariance_W
GA_5_01_256,Cells|Texture_DifferenceVariance_WGA_5_03_256,Cells|Texture_DifferenceVariance_WGA_5_00_25
6,Cells|Texture_DifferenceVariance_WGA_3_00_256,Cells|Texture_DifferenceVariance_WGA_3_03_256,Cells|Textu
re_DifferenceVariance_WGA_3_01_256,Cells|Texture_DifferenceVariance_WGA_3_02_256,Cells|Texture_Differenc
eVariance_WGA_20_01_256,Cells|Texture_DifferenceVariance_WGA_20_02_256,Cells|Texture_DifferenceVariance
_WGA_20_00_256,Cells|Texture_DifferenceVariance_WGA_20_03_256,Cells|Texture_DifferenceVariance_WGA_10
_03_256,Cells|Texture_DifferenceVariance_WGA_10_02_256,Cells|Texture_DifferenceVariance_WGA_10_00_256,C
ells|Texture_DifferenceVariance_WGA_10_01_256,Cells|Texture_DifferenceVariance_DNA_10_03_256,Cells|Texture
_DifferenceVariance_DNA_10_00_256,Cells|Texture_DifferenceVariance_DNA_10_01_256,Cells|Texture_Difference
Variance_DNA_10_02_256,Cells|Texture_DifferenceVariance_DNA_20_02_256,Cells|Texture_DifferenceVariance_D
NA_20_01_256,Cells|Texture_DifferenceVariance_DNA_20_00_256,Cells|Texture_DifferenceVariance_DNA_20_03_
256,Cells|Texture_DifferenceVariance_DNA_3_02_256,Cells|Texture_DifferenceVariance_DNA_3_03_256,Cells|Text
ure_DifferenceVariance_DNA_3_00_256,Cells|Texture_DifferenceVariance_DNA_3_01_256,Cells|Texture_Differenc

eVariance_DNA_5_02_256,Cells|Texture_DifferenceVariance_DNA_5_00_256,Cells|Texture_DifferenceVariance_DNA_5_03_256,Cells|Texture_DifferenceVariance_DNA_5_01_256,Cells|Texture_InfoMeas1_Vinculin_5_02_256,Cells|Texture_InfoMeas1_Vinculin_5_03_256,Cells|Texture_InfoMeas1_Vinculin_5_00_256,Cells|Texture_InfoMeas1_Vinculin_5_01_256,Cells|Texture_InfoMeas1_Vinculin_20_01_256,Cells|Texture_InfoMeas1_Vinculin_20_03_256,Cells|Texture_InfoMeas1_Vinculin_20_00_256,Cells|Texture_InfoMeas1_Vinculin_20_02_256,Cells|Texture_InfoMeas1_Vinculin_10_01_256,Cells|Texture_InfoMeas1_Vinculin_10_00_256,Cells|Texture_InfoMeas1_Vinculin_10_03_256,Cells|Texture_InfoMeas1_Vinculin_10_02_256,Cells|Texture_InfoMeas1_Vinculin_3_02_256,Cells|Texture_InfoMeas1_Vinculin_3_01_256,Cells|Texture_InfoMeas1_Vinculin_3_00_256,Cells|Texture_InfoMeas1_Vinculin_3_03_256,Cells|Texture_InfoMeas1_DNA_5_02_256,Cells|Texture_InfoMeas1_DNA_5_03_256,Cells|Texture_InfoMeas1_DNA_5_00_256,Cells|Texture_InfoMeas1_DNA_5_01_256,Cells|Texture_InfoMeas1_DNA_10_03_256,Cells|Texture_InfoMeas1_DNA_10_02_256,Cells|Texture_InfoMeas1_DNA_10_01_256,Cells|Texture_InfoMeas1_DNA_10_00_256,Cells|Texture_InfoMeas1_DNA_20_03_256,Cells|Texture_InfoMeas1_DNA_20_01_256,Cells|Texture_InfoMeas1_DNA_20_02_256,Cells|Texture_InfoMeas1_DNA_20_00_256,Cells|Texture_InfoMeas1_DNA_3_02_256,Cells|Texture_InfoMeas1_DNA_3_01_256,Cells|Texture_InfoMeas1_DNA_3_03_256,Cells|Texture_InfoMeas1_DNA_3_00_256,Cells|Texture_InfoMeas1_WGA_10_02_256,Cells|Texture_InfoMeas1_WGA_10_01_256,Cells|Texture_InfoMeas1_WGA_10_00_256,Cells|Texture_InfoMeas1_WGA_10_03_256,Cells|Texture_InfoMeas1_WGA_20_01_256,Cells|Texture_InfoMeas1_WGA_20_03_256,Cells|Texture_InfoMeas1_WGA_20_00_256,Cells|Texture_InfoMeas1_WGA_20_02_256,Cells|Texture_InfoMeas1_WGA_3_02_256,Cells|Texture_InfoMeas1_WGA_3_00_256,Cells|Texture_InfoMeas1_WGA_3_01_256,Cells|Texture_InfoMeas1_WGA_3_03_256,Cells|Texture_InfoMeas1_WGA_5_02_256,Cells|Texture_InfoMeas1_WGA_5_03_256,Cells|Texture_InfoMeas1_WGA_5_00_256,Cells|Texture_InfoMeas1_WGA_5_01_256,Cells|Texture_InfoMeas1_Phalloidin_3_00_256,Cells|Texture_InfoMeas1_Phalloidin_3_01_256,Cells|Texture_InfoMeas1_Phalloidin_3_02_256,Cells|Texture_InfoMeas1_Phalloidin_3_03_256,Cells|Texture_InfoMeas1_Phalloidin_20_00_256,Cells|Texture_InfoMeas1_Phalloidin_20_01_256,Cells|Texture_InfoMeas1_Phalloidin_20_02_256,Cells|Texture_InfoMeas1_Phalloidin_20_03_256,Cells|Texture_InfoMeas1_Phalloidin_5_02_256,Cells|Texture_InfoMeas1_Phalloidin_5_03_256,Cells|Texture_InfoMeas1_Phalloidin_5_01_256,Cells|Texture_InfoMeas1_Phalloidin_5_00_256,Cells|Texture_InfoMeas1_Phalloidin_10_00_256,Cells|Texture_InfoMeas1_Phalloidin_10_03_256,Cells|Texture_InfoMeas1_Phalloidin_10_02_256,Cells|Texture_InfoMeas1_Phalloidin_10_01_256,Cells|Texture_AngularSecondMoment_DNA_10_00_256,Cells|Texture_AngularSecondMoment_DNA_10_01_256,Cells|Texture_AngularSecondMoment_DNA_10_03_256,Cells|Texture_AngularSecondMoment_DNA_10_02_256,Cells|Texture_AngularSecondMoment_DNA_5_00_256,Cells|Texture_AngularSecondMoment_DNA_5_02_256,Cells|Texture_AngularSecondMoment_DNA_5_01_256,Cells|Texture_AngularSecondMoment_DNA_5_03_256,Cells|Texture_AngularSecondMoment_DNA_20_03_256,Cells|Texture_AngularSecondMoment_DNA_20_01_256,Cells|Texture_AngularSecondMoment_DNA_20_00_256,Cells|Texture_AngularSecondMoment_DNA_20_02_256,Cells|Texture_AngularSecondMoment_DNA_3_03_256,Cells|Texture_AngularSecondMoment_DNA_3_00_256,Cells|Texture_AngularSecondMoment_DNA_3_01_256,Cells|Texture_AngularSecondMoment_DNA_3_02_256,Cells|Texture_AngularSecondMoment_Vinculin_5_03_256,Cells|Texture_AngularSecondMoment_Vinculin_5_02_256,Cells|Texture_AngularSecondMoment_Vinculin_5_01_256,Cells|Texture_AngularSecondMoment_Vinculin_5_00_256,Cells|Texture_AngularSecondMoment_Vinculin_10_01_256,Cells|Texture_AngularSecondMoment_Vinculin_10_02_256,Cells|Texture_AngularSecondMoment_Vinculin_10_00_256,Cells|Texture_AngularSecondMoment_Vinculin_10_03_256,Cells|Texture_AngularSecondMoment_Vinculin_3_03_256,Cells|Texture_AngularSecondMoment_Vinculin_3_01_256,Cells|Texture_AngularSecondMoment_Vinculin_3_00_256,Cells|Texture_AngularSecondMoment_Vinculin_3_02_256,Cells|Texture_AngularSecondMoment_Vinculin_20_03_256,Cells|Texture_AngularSecondMoment_Vinculin_20_00_256,Cells|Texture_AngularSecondMoment_Vinculin_20_02_256,Cells|Texture_AngularSecondMoment_Vinculin_20_01_256,Cells|Texture_AngularSecondMoment_Phalloidin_10_03_256,Cells|Texture_AngularSecondMoment_Phalloidin_10_02_256,Cells|Texture_AngularSecondMoment_Phalloidin_10_00_256,Cells|Texture_AngularSecondMoment_Phalloidin_10_01_256,Cells|Texture_AngularSecondMoment_Phalloidin_20_02_256,Cells|Texture_AngularSecondMoment_Phalloidin_20_00_256,Cells|Texture_AngularSecondMoment_Phalloidin_20_03_256,Cells|Texture_AngularSecondMoment_Phalloidin_20_01_256,Cells|Texture_AngularSecondMoment_Phalloidin_5_03_256,Cells|Texture_AngularSecondMoment_Phalloidin_5_00_256,Cells|Texture_AngularSecondMoment_Phalloidin_5_02_256,Cells|Texture_AngularSecondMoment_Phalloidin_5_01_256,Cells|Texture_AngularSecondMoment_Phalloidin_3_03_256,Cells|Texture_AngularSecondMoment_Phalloidin_3_01_256,Cells|Texture_AngularSecondMoment_Phalloidin_3_02_256,Cells|Texture_AngularSecondMoment_Phalloidin_3_00_256,Cells|Texture_AngularSecondMoment_WGA_5_01_256,Cells|Texture_AngularSecondMoment_WGA_5_02_256,Cells|Texture_AngularSecondMoment_WGA_5_00_256,Cells|Texture_AngularSecondMoment_WGA_5_03_256,Cells|Texture_AngularSecondMoment_WGA_10_03_256,Cells|Texture_AngularSecondMoment_WGA_10_01_256,Cells|Texture_AngularSecondMoment_WG

A_10_02_256,Cells|Texture_AngularSecondMoment_WGA_10_00_256,Cells|Texture_AngularSecondMoment_WGA
_20_02_256,Cells|Texture_AngularSecondMoment_WGA_20_00_256,Cells|Texture_AngularSecondMoment_WGA_2
0_03_256,Cells|Texture_AngularSecondMoment_WGA_20_01_256,Cells|Texture_AngularSecondMoment_WGA_3_0
0_256,Cells|Texture_AngularSecondMoment_WGA_3_01_256,Cells|Texture_AngularSecondMoment_WGA_3_02_25
6,Cells|Texture_AngularSecondMoment_WGA_3_03_256,Cells|Texture_DifferenceEntropy_DNA_20_01_256,Cells|T
exture_DifferenceEntropy_DNA_20_02_256,Cells|Texture_DifferenceEntropy_DNA_20_00_256,Cells|Texture_Differ
enceEntropy_DNA_20_03_256,Cells|Texture_DifferenceEntropy_DNA_3_01_256,Cells|Texture_DifferenceEntropy_
DNA_3_02_256,Cells|Texture_DifferenceEntropy_DNA_3_00_256,Cells|Texture_DifferenceEntropy_DNA_3_03_256
,Cells|Texture_DifferenceEntropy_DNA_10_02_256,Cells|Texture_DifferenceEntropy_DNA_10_01_256,Cells|Texture
_DifferenceEntropy_DNA_10_00_256,Cells|Texture_DifferenceEntropy_DNA_10_03_256,Cells|Texture_DifferenceE
ntropy_DNA_5_03_256,Cells|Texture_DifferenceEntropy_DNA_5_00_256,Cells|Texture_DifferenceEntropy_DNA_5
_01_256,Cells|Texture_DifferenceEntropy_DNA_5_02_256,Cells|Texture_DifferenceEntropy_Phalloidin_20_00_256,
Cells|Texture_DifferenceEntropy_Phalloidin_20_01_256,Cells|Texture_DifferenceEntropy_Phalloidin_20_02_256,Cell
s|Texture_DifferenceEntropy_Phalloidin_20_03_256,Cells|Texture_DifferenceEntropy_Phalloidin_10_02_256,Cells|Te
xture_DifferenceEntropy_Phalloidin_10_01_256,Cells|Texture_DifferenceEntropy_Phalloidin_10_03_256,Cells|Textur
e_DifferenceEntropy_Phalloidin_10_00_256,Cells|Texture_DifferenceEntropy_Phalloidin_3_00_256,Cells|Texture_Dif
ferenceEntropy_Phalloidin_3_02_256,Cells|Texture_DifferenceEntropy_Phalloidin_3_03_256,Cells|Texture_Differenc
eEntropy_Phalloidin_3_01_256,Cells|Texture_DifferenceEntropy_Phalloidin_5_03_256,Cells|Texture_DifferenceEntro
py_Phalloidin_5_01_256,Cells|Texture_DifferenceEntropy_Phalloidin_5_02_256,Cells|Texture_DifferenceEntropy_Ph
alloidin_5_00_256,Cells|Texture_DifferenceEntropy_Vinculin_3_01_256,Cells|Texture_DifferenceEntropy_Vinculin_3
_02_256,Cells|Texture_DifferenceEntropy_Vinculin_3_03_256,Cells|Texture_DifferenceEntropy_Vinculin_3_00_256,
Cells|Texture_DifferenceEntropy_Vinculin_10_03_256,Cells|Texture_DifferenceEntropy_Vinculin_10_01_256,Cells|T
exture_DifferenceEntropy_Vinculin_10_02_256,Cells|Texture_DifferenceEntropy_Vinculin_10_00_256,Cells|Texture_
DifferenceEntropy_Vinculin_5_00_256,Cells|Texture_DifferenceEntropy_Vinculin_5_03_256,Cells|Texture_Differenc
eEntropy_Vinculin_5_02_256,Cells|Texture_DifferenceEntropy_Vinculin_5_01_256,Cells|Texture_DifferenceEntropy
_Vinculin_20_00_256,Cells|Texture_DifferenceEntropy_Vinculin_20_01_256,Cells|Texture_DifferenceEntropy_Vincu
lin_20_03_256,Cells|Texture_DifferenceEntropy_Vinculin_20_02_256,Cells|Texture_DifferenceEntropy_WGA_20_01
_256,Cells|Texture_DifferenceEntropy_WGA_20_03_256,Cells|Texture_DifferenceEntropy_WGA_20_00_256,Cells|T
exture_DifferenceEntropy_WGA_20_02_256,Cells|Texture_DifferenceEntropy_WGA_5_03_256,Cells|Texture_Differ
enceEntropy_WGA_5_00_256,Cells|Texture_DifferenceEntropy_WGA_5_02_256,Cells|Texture_DifferenceEntropy_
WGA_5_01_256,Cells|Texture_DifferenceEntropy_WGA_3_00_256,Cells|Texture_DifferenceEntropy_WGA_3_03_2
56,Cells|Texture_DifferenceEntropy_WGA_3_01_256,Cells|Texture_DifferenceEntropy_WGA_3_02_256,Cells|Textur
e_DifferenceEntropy_WGA_10_02_256,Cells|Texture_DifferenceEntropy_WGA_10_03_256,Cells|Texture_Difference
Entropy_WGA_10_00_256,Cells|Texture_DifferenceEntropy_WGA_10_01_256,Cells|Texture_SumAverage_Phalloidi
n_3_03_256,Cells|Texture_SumAverage_Phalloidin_3_01_256,Cells|Texture_SumAverage_Phalloidin_3_00_256,Cells
|Texture_SumAverage_Phalloidin_3_02_256,Cells|Texture_SumAverage_Phalloidin_20_01_256,Cells|Texture_SumAv
erage_Phalloidin_20_03_256,Cells|Texture_SumAverage_Phalloidin_20_00_256,Cells|Texture_SumAverage_Phalloidi
n_20_02_256,Cells|Texture_SumAverage_Phalloidin_10_02_256,Cells|Texture_SumAverage_Phalloidin_10_01_256,C
ells|Texture_SumAverage_Phalloidin_10_00_256,Cells|Texture_SumAverage_Phalloidin_10_03_256,Cells|Texture_Su
mAverage_Phalloidin_5_03_256,Cells|Texture_SumAverage_Phalloidin_5_00_256,Cells|Texture_SumAverage_Phallo
idin_5_01_256,Cells|Texture_SumAverage_Phalloidin_5_02_256,Cells|Texture_SumAverage_Vinculin_3_03_256,Cell
s|Texture_SumAverage_Vinculin_3_00_256,Cells|Texture_SumAverage_Vinculin_3_02_256,Cells|Texture_SumAvera
ge_Vinculin_3_01_256,Cells|Texture_SumAverage_Vinculin_5_00_256,Cells|Texture_SumAverage_Vinculin_5_03_2
56,Cells|Texture_SumAverage_Vinculin_5_01_256,Cells|Texture_SumAverage_Vinculin_5_02_256,Cells|Texture_Su
mAverage_Vinculin_10_02_256,Cells|Texture_SumAverage_Vinculin_10_00_256,Cells|Texture_SumAverage_Vincul
in_10_03_256,Cells|Texture_SumAverage_Vinculin_10_01_256,Cells|Texture_SumAverage_Vinculin_20_01_256,Cel
ls|Texture_SumAverage_Vinculin_20_03_256,Cells|Texture_SumAverage_Vinculin_20_02_256,Cells|Texture_SumAv
erage_Vinculin_20_00_256,Cells|Texture_SumAverage_WGA_5_03_256,Cells|Texture_SumAverage_WGA_5_00_25
6,Cells|Texture_SumAverage_WGA_5_01_256,Cells|Texture_SumAverage_WGA_5_02_256,Cells|Texture_SumAver
age_WGA_20_02_256,Cells|Texture_SumAverage_WGA_20_01_256,Cells|Texture_SumAverage_WGA_20_00_256,
Cells|Texture_SumAverage_WGA_20_03_256,Cells|Texture_SumAverage_WGA_10_03_256,Cells|Texture_SumAver
age_WGA_10_00_256,Cells|Texture_SumAverage_WGA_10_01_256,Cells|Texture_SumAverage_WGA_10_02_256,
Cells|Texture_SumAverage_WGA_3_03_256,Cells|Texture_SumAverage_WGA_3_02_256,Cells|Texture_SumAverag
e_WGA_3_01_256,Cells|Texture_SumAverage_WGA_3_00_256,Cells|Texture_SumAverage_DNA_5_01_256,Cells|T

exture_SumAverage_DNA_5_03_256,Cells|Texture_SumAverage_DNA_5_02_256,Cells|Texture_SumAverage_DNA
_5_00_256,Cells|Texture_SumAverage_DNA_10_01_256,Cells|Texture_SumAverage_DNA_10_00_256,Cells|Texture
_SumAverage_DNA_10_02_256,Cells|Texture_SumAverage_DNA_10_03_256,Cells|Texture_SumAverage_DNA_3_
01_256,Cells|Texture_SumAverage_DNA_3_03_256,Cells|Texture_SumAverage_DNA_3_00_256,Cells|Texture_Sum
Average_DNA_3_02_256,Cells|Texture_SumAverage_DNA_20_01_256,Cells|Texture_SumAverage_DNA_20_02_25
6,Cells|Texture_SumAverage_DNA_20_00_256,Cells|Texture_SumAverage_DNA_20_03_256,Cells|Texture_InverseD
ifferenceMoment_Vinculin_3_01_256,Cells|Texture_InverseDifferenceMoment_Vinculin_3_00_256,Cells|Texture_Inv
erseDifferenceMoment_Vinculin_3_03_256,Cells|Texture_InverseDifferenceMoment_Vinculin_3_02_256,Cells|Textur
e_InverseDifferenceMoment_Vinculin_10_03_256,Cells|Texture_InverseDifferenceMoment_Vinculin_10_01_256,Cell
s|Texture_InverseDifferenceMoment_Vinculin_10_02_256,Cells|Texture_InverseDifferenceMoment_Vinculin_10_00_
256,Cells|Texture_InverseDifferenceMoment_Vinculin_20_00_256,Cells|Texture_InverseDifferenceMoment_Vinculin
_20_03_256,Cells|Texture_InverseDifferenceMoment_Vinculin_20_01_256,Cells|Texture_InverseDifferenceMoment_
Vinculin_20_02_256,Cells|Texture_InverseDifferenceMoment_Vinculin_5_00_256,Cells|Texture_InverseDifferenceM
oment_Vinculin_5_02_256,Cells|Texture_InverseDifferenceMoment_Vinculin_5_03_256,Cells|Texture_InverseDiffere
nceMoment_Vinculin_5_01_256,Cells|Texture_InverseDifferenceMoment_WGA_5_03_256,Cells|Texture_InverseDiff
erenceMoment_WGA_5_00_256,Cells|Texture_InverseDifferenceMoment_WGA_5_01_256,Cells|Texture_InverseDiff
erenceMoment_WGA_5_02_256,Cells|Texture_InverseDifferenceMoment_WGA_3_02_256,Cells|Texture_InverseDiff
erenceMoment_WGA_3_03_256,Cells|Texture_InverseDifferenceMoment_WGA_3_00_256,Cells|Texture_InverseDiff
erenceMoment_WGA_3_01_256,Cells|Texture_InverseDifferenceMoment_WGA_10_01_256,Cells|Texture_InverseDi
fferenceMoment_WGA_10_00_256,Cells|Texture_InverseDifferenceMoment_WGA_10_02_256,Cells|Texture_Inverse
DifferenceMoment_WGA_10_03_256,Cells|Texture_InverseDifferenceMoment_WGA_20_02_256,Cells|Texture_Inve
rseDifferenceMoment_WGA_20_03_256,Cells|Texture_InverseDifferenceMoment_WGA_20_01_256,Cells|Texture_In
verseDifferenceMoment_WGA_20_00_256,Cells|Texture_InverseDifferenceMoment_Phalloidin_5_03_256,Cells|Text
ure_InverseDifferenceMoment_Phalloidin_5_01_256,Cells|Texture_InverseDifferenceMoment_Phalloidin_5_02_256,C
ells|Texture_InverseDifferenceMoment_Phalloidin_5_00_256,Cells|Texture_InverseDifferenceMoment_Phalloidin_20_
03_256,Cells|Texture_InverseDifferenceMoment_Phalloidin_20_02_256,Cells|Texture_InverseDifferenceMoment_Pha
lloidin_20_00_256,Cells|Texture_InverseDifferenceMoment_Phalloidin_20_01_256,Cells|Texture_InverseDifferenceM
oment_Phalloidin_3_02_256,Cells|Texture_InverseDifferenceMoment_Phalloidin_3_00_256,Cells|Texture_InverseDiff
erenceMoment_Phalloidin_3_01_256,Cells|Texture_InverseDifferenceMoment_Phalloidin_3_03_256,Cells|Texture_In
verseDifferenceMoment_Phalloidin_10_00_256,Cells|Texture_InverseDifferenceMoment_Phalloidin_10_01_256,Cells|
Texture_InverseDifferenceMoment_Phalloidin_10_03_256,Cells|Texture_InverseDifferenceMoment_Phalloidin_10_02
_256,Cells|Texture_InverseDifferenceMoment_DNA_20_00_256,Cells|Texture_InverseDifferenceMoment_DNA_20_0
3_256,Cells|Texture_InverseDifferenceMoment_DNA_20_02_256,Cells|Texture_InverseDifferenceMoment_DNA_20_
01_256,Cells|Texture_InverseDifferenceMoment_DNA_10_02_256,Cells|Texture_InverseDifferenceMoment_DNA_10
_00_256,Cells|Texture_InverseDifferenceMoment_DNA_10_01_256,Cells|Texture_InverseDifferenceMoment_DNA_1
0_03_256,Cells|Texture_InverseDifferenceMoment_DNA_5_00_256,Cells|Texture_InverseDifferenceMoment_DNA_5
_03_256,Cells|Texture_InverseDifferenceMoment_DNA_5_01_256,Cells|Texture_InverseDifferenceMoment_DNA_5_
02_256,Cells|Texture_InverseDifferenceMoment_DNA_3_01_256,Cells|Texture_InverseDifferenceMoment_DNA_3_0
2_256,Cells|Texture_InverseDifferenceMoment_DNA_3_00_256,Cells|Texture_InverseDifferenceMoment_DNA_3_03
_256,Cells|Texture_SumVariance_WGA_20_01_256,Cells|Texture_SumVariance_WGA_20_03_256,Cells|Texture_Su
mVariance_WGA_20_02_256,Cells|Texture_SumVariance_WGA_20_00_256,Cells|Texture_SumVariance_WGA_5_0
1_256,Cells|Texture_SumVariance_WGA_5_02_256,Cells|Texture_SumVariance_WGA_5_03_256,Cells|Texture_Su
mVariance_WGA_5_00_256,Cells|Texture_SumVariance_WGA_10_00_256,Cells|Texture_SumVariance_WGA_10_0
1_256,Cells|Texture_SumVariance_WGA_10_03_256,Cells|Texture_SumVariance_WGA_10_02_256,Cells|Texture_S
umVariance_WGA_3_02_256,Cells|Texture_SumVariance_WGA_3_01_256,Cells|Texture_SumVariance_WGA_3_00
_256,Cells|Texture_SumVariance_WGA_3_03_256,Cells|Texture_SumVariance_DNA_20_01_256,Cells|Texture_Sum
Variance_DNA_20_02_256,Cells|Texture_SumVariance_DNA_20_00_256,Cells|Texture_SumVariance_DNA_20_03_
256,Cells|Texture_SumVariance_DNA_3_01_256,Cells|Texture_SumVariance_DNA_3_02_256,Cells|Texture_SumVa
riance_DNA_3_00_256,Cells|Texture_SumVariance_DNA_3_03_256,Cells|Texture_SumVariance_DNA_5_01_256,C
ells|Texture_SumVariance_DNA_5_02_256,Cells|Texture_SumVariance_DNA_5_03_256,Cells|Texture_SumVariance
_DNA_5_00_256,Cells|Texture_SumVariance_DNA_10_01_256,Cells|Texture_SumVariance_DNA_10_00_256,Cells|
Texture_SumVariance_DNA_10_02_256,Cells|Texture_SumVariance_DNA_10_03_256,Cells|Texture_SumVariance_
Vinculin_20_03_256,Cells|Texture_SumVariance_Vinculin_20_00_256,Cells|Texture_SumVariance_Vinculin_20_02_
256,Cells|Texture_SumVariance_Vinculin_20_01_256,Cells|Texture_SumVariance_Vinculin_5_03_256,Cells|Texture_

SumVariance_Vinculin_5_00_256,Cells|Texture_SumVariance_Vinculin_5_02_256,Cells|Texture_SumVariance_Vinc
ulin_5_01_256,Cells|Texture_SumVariance_Vinculin_3_00_256,Cells|Texture_SumVariance_Vinculin_3_01_256,Cell
s|Texture_SumVariance_Vinculin_3_03_256,Cells|Texture_SumVariance_Vinculin_3_02_256,Cells|Texture_SumVari
ance_Vinculin_10_02_256,Cells|Texture_SumVariance_Vinculin_10_01_256,Cells|Texture_SumVariance_Vinculin_1
0_03_256,Cells|Texture_SumVariance_Vinculin_10_00_256,Cells|Texture_SumVariance_Phalloidin_3_01_256,Cells|
Texture_SumVariance_Phalloidin_3_00_256,Cells|Texture_SumVariance_Phalloidin_3_02_256,Cells|Texture_SumVar
iance_Phalloidin_3_03_256,Cells|Texture_SumVariance_Phalloidin_5_00_256,Cells|Texture_SumVariance_Phalloidin
_5_02_256,Cells|Texture_SumVariance_Phalloidin_5_01_256,Cells|Texture_SumVariance_Phalloidin_5_03_256,Cells
|Texture_SumVariance_Phalloidin_10_00_256,Cells|Texture_SumVariance_Phalloidin_10_01_256,Cells|Texture_Sum
Variance_Phalloidin_10_02_256,Cells|Texture_SumVariance_Phalloidin_10_03_256,Cells|Texture_SumVariance_Phal
loidin_20_00_256,Cells|Texture_SumVariance_Phalloidin_20_03_256,Cells|Texture_SumVariance_Phalloidin_20_01_
256,Cells|Texture_SumVariance_Phalloidin_20_02_256,Cells|Texture_Entropy_Phalloidin_10_00_256,Cells|Texture_
Entropy_Phalloidin_10_02_256,Cells|Texture_Entropy_Phalloidin_10_03_256,Cells|Texture_Entropy_Phalloidin_10_0
1_256,Cells|Texture_Entropy_Phalloidin_20_03_256,Cells|Texture_Entropy_Phalloidin_20_00_256,Cells|Texture_Entr
opy_Phalloidin_20_01_256,Cells|Texture_Entropy_Phalloidin_20_02_256,Cells|Texture_Entropy_Phalloidin_3_03_25
6,Cells|Texture_Entropy_Phalloidin_3_02_256,Cells|Texture_Entropy_Phalloidin_3_00_256,Cells|Texture_Entropy_Ph
alloidin_3_01_256,Cells|Texture_Entropy_Phalloidin_5_00_256,Cells|Texture_Entropy_Phalloidin_5_01_256,Cells|Te
xture_Entropy_Phalloidin_5_03_256,Cells|Texture_Entropy_Phalloidin_5_02_256,Cells|Texture_Entropy_DNA_5_02
_256,Cells|Texture_Entropy_DNA_5_00_256,Cells|Texture_Entropy_DNA_5_03_256,Cells|Texture_Entropy_DNA_5
_01_256,Cells|Texture_Entropy_DNA_3_02_256,Cells|Texture_Entropy_DNA_3_03_256,Cells|Texture_Entropy_DN
A_3_00_256,Cells|Texture_Entropy_DNA_3_01_256,Cells|Texture_Entropy_DNA_10_01_256,Cells|Texture_Entropy
_DNA_10_02_256,Cells|Texture_Entropy_DNA_10_00_256,Cells|Texture_Entropy_DNA_10_03_256,Cells|Texture_
Entropy_DNA_20_01_256,Cells|Texture_Entropy_DNA_20_00_256,Cells|Texture_Entropy_DNA_20_03_256,Cells|T
exture_Entropy_DNA_20_02_256,Cells|Texture_Entropy_WGA_3_02_256,Cells|Texture_Entropy_WGA_3_01_256,C
ells|Texture_Entropy_WGA_3_00_256,Cells|Texture_Entropy_WGA_3_03_256,Cells|Texture_Entropy_WGA_20_00_
256,Cells|Texture_Entropy_WGA_20_01_256,Cells|Texture_Entropy_WGA_20_03_256,Cells|Texture_Entropy_WGA
_20_02_256,Cells|Texture_Entropy_WGA_10_01_256,Cells|Texture_Entropy_WGA_10_00_256,Cells|Texture_Entrop
y_WGA_10_03_256,Cells|Texture_Entropy_WGA_10_02_256,Cells|Texture_Entropy_WGA_5_00_256,Cells|Texture
_Entropy_WGA_5_03_256,Cells|Texture_Entropy_WGA_5_01_256,Cells|Texture_Entropy_WGA_5_02_256,Cells|Te
xture_Entropy_Vinculin_10_01_256,Cells|Texture_Entropy_Vinculin_10_00_256,Cells|Texture_Entropy_Vinculin_10
_02_256,Cells|Texture_Entropy_Vinculin_10_03_256,Cells|Texture_Entropy_Vinculin_3_02_256,Cells|Texture_Entro
py_Vinculin_3_03_256,Cells|Texture_Entropy_Vinculin_3_01_256,Cells|Texture_Entropy_Vinculin_3_00_256,Cells|T
exture_Entropy_Vinculin_5_03_256,Cells|Texture_Entropy_Vinculin_5_02_256,Cells|Texture_Entropy_Vinculin_5_0
1_256,Cells|Texture_Entropy_Vinculin_5_00_256,Cells|Texture_Entropy_Vinculin_20_01_256,Cells|Texture_Entropy
_Vinculin_20_03_256,Cells|Texture_Entropy_Vinculin_20_00_256,Cells|Texture_Entropy_Vinculin_20_02_256,Cells|
Texture_SumEntropy_Phalloidin_20_03_256,Cells|Texture_SumEntropy_Phalloidin_20_00_256,Cells|Texture_SumEn
tropy_Phalloidin_20_02_256,Cells|Texture_SumEntropy_Phalloidin_20_01_256,Cells|Texture_SumEntropy_Phalloidin
_3_03_256,Cells|Texture_SumEntropy_Phalloidin_3_00_256,Cells|Texture_SumEntropy_Phalloidin_3_02_256,Cells|T
exture_SumEntropy_Phalloidin_3_01_256,Cells|Texture_SumEntropy_Phalloidin_5_01_256,Cells|Texture_SumEntrop
y_Phalloidin_5_00_256,Cells|Texture_SumEntropy_Phalloidin_5_03_256,Cells|Texture_SumEntropy_Phalloidin_5_02
_256,Cells|Texture_SumEntropy_Phalloidin_10_03_256,Cells|Texture_SumEntropy_Phalloidin_10_01_256,Cells|Text
ure_SumEntropy_Phalloidin_10_00_256,Cells|Texture_SumEntropy_Phalloidin_10_02_256,Cells|Texture_SumEntrop
y_Vinculin_3_03_256,Cells|Texture_SumEntropy_Vinculin_3_00_256,Cells|Texture_SumEntropy_Vinculin_3_01_25
6,Cells|Texture_SumEntropy_Vinculin_3_02_256,Cells|Texture_SumEntropy_Vinculin_5_03_256,Cells|Texture_Sum
Entropy_Vinculin_5_01_256,Cells|Texture_SumEntropy_Vinculin_5_00_256,Cells|Texture_SumEntropy_Vinculin_5_
02_256,Cells|Texture_SumEntropy_Vinculin_10_00_256,Cells|Texture_SumEntropy_Vinculin_10_02_256,Cells|Textu
re_SumEntropy_Vinculin_10_03_256,Cells|Texture_SumEntropy_Vinculin_10_01_256,Cells|Texture_SumEntropy_Vi
nculin_20_02_256,Cells|Texture_SumEntropy_Vinculin_20_01_256,Cells|Texture_SumEntropy_Vinculin_20_00_256,
Cells|Texture_SumEntropy_Vinculin_20_03_256,Cells|Texture_SumEntropy_WGA_5_00_256,Cells|Texture_SumEntr
opy_WGA_5_03_256,Cells|Texture_SumEntropy_WGA_5_01_256,Cells|Texture_SumEntropy_WGA_5_02_256,Cells
|Texture_SumEntropy_WGA_10_02_256,Cells|Texture_SumEntropy_WGA_10_01_256,Cells|Texture_SumEntropy_
WGA_10_03_256,Cells|Texture_SumEntropy_WGA_10_00_256,Cells|Texture_SumEntropy_WGA_3_01_256,Cells|T
exture_SumEntropy_WGA_3_00_256,Cells|Texture_SumEntropy_WGA_3_02_256,Cells|Texture_SumEntropy_WGA
_3_03_256,Cells|Texture_SumEntropy_WGA_20_02_256,Cells|Texture_SumEntropy_WGA_20_03_256,Cells|Texture

_SumEntropy_WGA_20_01_256,Cells|Texture_SumEntropy_WGA_20_00_256,Cells|Texture_SumEntropy_DNA_5_00_256,Cells|Texture_SumEntropy_DNA_5_01_256,Cells|Texture_SumEntropy_DNA_5_02_256,Cells|Texture_SumEntropy_DNA_5_03_256,Cells|Texture_SumEntropy_DNA_3_02_256,Cells|Texture_SumEntropy_DNA_3_01_256,Cells|Texture_SumEntropy_DNA_3_03_256,Cells|Texture_SumEntropy_DNA_3_00_256,Cells|Texture_SumEntropy_DNA_10_03_256,Cells|Texture_SumEntropy_DNA_10_02_256,Cells|Texture_SumEntropy_DNA_10_00_256,Cells|Texture_SumEntropy_DNA_10_01_256,Cells|Texture_SumEntropy_DNA_20_02_256,Cells|Texture_SumEntropy_DNA_20_00_256,Cells|Texture_SumEntropy_DNA_20_03_256,Cells|Texture_SumEntropy_DNA_20_01_256,Cells|Granularity_2_Phalloidin,Cells|Granularity_2_Vinculin,Cells|Granularity_2_WGA,Cells|Granularity_2_DNA,Cells|Granularity_1_Vinculin,Cells|Granularity_1_DNA,Cells|Granularity_1_WGA,Cells|Granularity_1_Phalloidin,Cells|Granularity_14_Phalloidin,Cells|Granularity_14_Vinculin,Cells|Granularity_14_WGA,Cells|Granularity_14_DNA,Cells|Granularity_9_Phalloidin,Cells|Granularity_9_Vinculin,Cells|Granularity_9_DNA,Cells|Granularity_9_WGA,Cells|Granularity_7_WGA,Cells|Granularity_7_Phalloidin,Cells|Granularity_7_Vinculin,Cells|Granularity_7_DNA,Cells|Granularity_12_DNA,Cells|Granularity_12_Vinculin,Cells|Granularity_12_WGA,Cells|Granularity_12_Phalloidin,Cells|Granularity_10_Phalloidin,Cells|Granularity_10_Vinculin,Cells|Granularity_10_DNA,Cells|Granularity_10_WGA,Cells|Granularity_11_WGA,Cells|Granularity_11_Phalloidin,Cells|Granularity_11_Vinculin,Cells|Granularity_11_DNA,Cells|Granularity_15_Phalloidin,Cells|Granularity_15_WGA,Cells|Granularity_15_DNA,Cells|Granularity_15_Vinculin,Cells|Granularity_8_Phalloidin,Cells|Granularity_8_Vinculin,Cells|Granularity_8_WGA,Cells|Granularity_8_DNA,Cells|Granularity_5_Phalloidin,Cells|Granularity_5_DNA,Cells|Granularity_5_Vinculin,Cells|Granularity_5_WGA,Cells|Granularity_16_Vinculin,Cells|Granularity_16_WGA,Cells|Granularity_16_DNA,Cells|Granularity_16_Phalloidin,Cells|Granularity_6_DNA,Cells|Granularity_6_Vinculin,Cells|Granularity_6_WGA,Cells|Granularity_6_Phalloidin,Cells|Granularity_3_WGA,Cells|Granularity_3_Vinculin,Cells|Granularity_3_DNA,Cells|Granularity_3_Phalloidin,Cells|Granularity_13_Phalloidin,Cells|Granularity_13_Vinculin,Cells|Granularity_13_DNA,Cells|Granularity_13_WGA,Cells|Granularity_4_Phalloidin,Cells|Granularity_4_WGA,Cells|Granularity_4_Vinculin,Cells|Granularity_4_DNA,Cells|RadialDistribution_RadialCV_DNA_4of4,Cells|RadialDistribution_RadialCV_DNA_1of4,Cells|RadialDistribution_RadialCV_DNA_2of4,Cells|RadialDistribution_RadialCV_DNA_3of4,Cells|RadialDistribution_RadialCV_Phalloidin_1of4,Cells|RadialDistribution_RadialCV_Phalloidin_3of4,Cells|RadialDistribution_RadialCV_Phalloidin_2of4,Cells|RadialDistribution_RadialCV_Phalloidin_4of4,Cells|RadialDistribution_RadialCV_Vinculin_2of4,Cells|RadialDistribution_RadialCV_Vinculin_4of4,Cells|RadialDistribution_RadialCV_Vinculin_1of4,Cells|RadialDistribution_RadialCV_Vinculin_3of4,Cells|RadialDistribution_RadialCV_WGA_4of4,Cells|RadialDistribution_RadialCV_WGA_1of4,Cells|RadialDistribution_RadialCV_WGA_2of4,Cells|RadialDistribution_RadialCV_WGA_3of4,Cells|RadialDistribution_MeanFrac_WGA_3of4,Cells|RadialDistribution_MeanFrac_WGA_1of4,Cells|RadialDistribution_MeanFrac_WGA_2of4,Cells|RadialDistribution_MeanFrac_WGA_4of4,Cells|RadialDistribution_MeanFrac_Phalloidin_4of4,Cells|RadialDistribution_MeanFrac_Phalloidin_2of4,Cells|RadialDistribution_MeanFrac_Phalloidin_1of4,Cells|RadialDistribution_MeanFrac_Phalloidin_3of4,Cells|RadialDistribution_MeanFrac_Vinculin_1of4,Cells|RadialDistribution_MeanFrac_Vinculin_2of4,Cells|RadialDistribution_MeanFrac_Vinculin_3of4,Cells|RadialDistribution_MeanFrac_Vinculin_4of4,Cells|RadialDistribution_MeanFrac_DNA_2of4,Cells|RadialDistribution_MeanFrac_DNA_1of4,Cells|RadialDistribution_MeanFrac_DNA_4of4,Cells|RadialDistribution_MeanFrac_DNA_3of4,Cells|RadialDistribution_FracAtD_WGA_2of4,Cells|RadialDistribution_FracAtD_WGA_1of4,Cells|RadialDistribution_FracAtD_WGA_3of4,Cells|RadialDistribution_FracAtD_WGA_4of4,Cells|RadialDistribution_FracAtD_Vinculin_3of4,Cells|RadialDistribution_FracAtD_Vinculin_1of4,Cells|RadialDistribution_FracAtD_Vinculin_2of4,Cells|RadialDistribution_FracAtD_Vinculin_4of4,Cells|RadialDistribution_FracAtD_DNA_2of4,Cells|RadialDistribution_FracAtD_DNA_4of4,Cells|RadialDistribution_FracAtD_DNA_3of4,Cells|RadialDistribution_FracAtD_DNA_1of4,Cells|RadialDistribution_FracAtD_Phalloidin_1of4,Cells|RadialDistribution_FracAtD_Phalloidin_2of4,Cells|RadialDistribution_FracAtD_Phalloidin_4of4,Cells|RadialDistribution_FracAtD_Phalloidin_3of4,Cells|Neighbors_SecondClosestDistance_10,Cells|Neighbors_SecondClosestDistance_Adjacent,Cells|Neighbors_AngleBetweenNeighbors_10,Cells|Neighbors_AngleBetweenNeighbors_Adjacent,Cells|Neighbors_SecondClosestObjectNumber_10,Cells|Neighbors_SecondClosestObjectNumber_Adjacent,Cells|Neighbors_PercentTouching_10,Cells|Neighbors_PercentTouching_Adjacent,Cells|Neighbors_NumberOfNeighbors_10,Cells|Neighbors_NumberOfNeighbors_Adjacent,Cells|Neighbors_FirstClosestObjectNumber_10,Cells|Neighbors_FirstClosestObjectNumber_Adjacent,Cells|Neighbors_FirstClosestDistance_Adjacent,Cells|Neighbors_FirstClosestDistance_10,Cells|Correlation_RWC_DNA_WGA,Cells|Correlation_RWC_DNA_Phalloidin,Cells|Correlation_RWC_DNA_Vinculin,Cells|Correlation_RWC_Vinculin_WGA,Cells|Correlation_RWC_Vinculin_DNA,Cells|Correlation_RWC_Vinculin_Phalloidin,Cells|Correlation_RWC_Phalloidin_WGA,Cells|Correlation_RWC_Phalloidin_DNA,Cells|Correlation_RWC_Phalloidin_Vinculin,Cells|Correlation_RWC_WGA_Phalloidin,Cells|Correlation_RWC_WGA_DNA,Cells|Correlation_RWC_WGA_Vinculin,Cells|Correlation_K_WGA_Phalloidin,Cells|Correlation_K_WGA_DNA,Cells|Correlation_K_WGA_Vinculin,Cells|Correlation_K_Phalloidin_WGA,Cells|Co rrelation_K_Phalloidin_Vinculin,Cells|Correlation_K_Phalloidin_DNA,Cells|Correlation_K_DNA_WGA,Cells|Correlation_K_DNA_Vinculin,Cells|Correlation_K_DNA_Phalloidin,Cells|Correlation_K_Vinculin_WGA,Cells|Correlation_K_Vinculin_Phalloidin,Cells|Correlation_K_Vinculin_DNA,Cells|Correlation_Overlap_DNA_WGA,Cells|Correlation_Overlap_DNA_Phalloidin,Cells|Correlation_Overlap_DNA_Vinculin,Cells|Correlation_Overlap_Phalloidin_Vinculin,Cells|Correlation_Overlap_Phalloidin_WGA,Cells|Correlation_Overlap_Vinculin_WGA,Cells|Correlation_Correlation_DNA_WGA,Cells|Correlation_Correlation_DNA_Phalloidin,Cells|Correlation_Correlation_DNA_Vinculin,Cells|Correlation_Correlation_Phalloidin_WGA,Cells|Correlation_Correlation_Phalloidin_Vinculin,Cells|Correlation_Correlation_Vinculin_WGA,Cells|Correlation_Manders_Vinculin_DNA,Cells|Correlation_Manders_Vinculin_WGA,Cells|Correlation_Manders_Vinculin_Phalloidin,Cells|Correlation_Manders_DNA_Phalloidin,Cells|Correlation_Manders_DNA_Vinculin,Cells|Correlation_Manders_DNA_WGA,Cells|Correlation_Manders_Phalloidin_WGA,Cells|Correlation_Manders_Phalloidin_Vinculin,Cells|Correlation_Manders_Phalloidin_DNA,Cells|Correlation_Manders_WGA_Vinculin,Cells|Correlation_Manders_WGA_DNA,Cells|Correlation_Manders_WGA_Phalloidin,Cells|Intensity_MedianIntensity_WGA,Cells|Intensity_MedianIntensity_DNA,Cells|Intensity_MedianIntensity_Phalloidin,Cells|Intensity_MedianIntensity_Vinculin,Cells|Intensity_MinIntensityEdge_Vinculin,Cells|Intensity_MinIntensityEdge_Phalloidin,Cells|Intensity_MinIntensityEdge_WGA,Cells|Intensity_MinIntensityEdge_DNA,Cells|Intensity_MeanIntensity_DNA,Cells|Intensity_MeanIntensity_WGA,Cells|Intensity_MeanIntensity_Phalloidin,Cells|Intensity_MeanIntensity_Vinculin,Cells|Intensity_IntegratedIntensityEdge_Phalloidin,Cells|Intensity_IntegratedIntensityEdge_WGA,Cells|Intensity_IntegratedIntensityEdge_DNA,Cells|Intensity_IntegratedIntensityEdge_Vinculin,Cells|Intensity_MeanIntensityEdge_Vinculin,Cells|Intensity_MeanIntensityEdge_Phalloidin,Cells|Intensity_MeanIntensityEdge_WGA,Cells|Intensity_MeanIntensityEdge_DNA,Cells|Intensity_IntegratedIntensity_DNA,Cells|Intensity_IntegratedIntensity_Vinculin,Cells|Intensity_IntegratedIntensity_WGA,Cells|Intensity_IntegratedIntensity_Phalloidin,Cells|Intensity_UpperQuartileIntensity_DNA,Cells|Intensity_UpperQuartileIntensity_WGA,Cells|Intensity_UpperQuartileIntensity_Phalloidin,Cells|Intensity_UpperQuartileIntensity_Vinculin,Cells|Intensity_MaxIntensity_Vinculin,Cells|Intensity_MaxIntensity_WGA,Cells|Intensity_MaxIntensity_Phalloidin,Cells|Intensity_MaxIntensity_DNA,Cells|Intensity_MADIntensity_Phalloidin,Cells|Intensity_MADIntensity_DNA,Cells|Intensity_MADIntensity_Vinculin,Cells|Intensity_MADIntensity_WGA,Cells|Intensity_MaxIntensityEdge_Phalloidin,Cells|Intensity_MaxIntensityEdge_WGA,Cells|Intensity_MaxIntensityEdge_DNA,Cells|Intensity_MaxIntensityEdge_Vinculin,Cells|Intensity_StdIntensityEdge_WGA,Cells|Intensity_StdIntensityEdge_Phalloidin,Cells|Intensity_StdIntensityEdge_Vinculin,Cells|Intensity_StdIntensityEdge_DNA,Cells|Intensity_MinIntensity_WGA,Cells|Intensity_MinIntensity_Vinculin,Cells|Intensity_MinIntensity_DNA,Cells|Intensity_MinIntensity_Phalloidin,Cells|Intensity_StdIntensity_DNA,Cells|Intensity_StdIntensity_Vinculin,Cells|Intensity_StdIntensity_WGA,Cells|Intensity_StdIntensity_Phalloidin,Cells|Intensity_LowerQuartileIntensity_Vinculin,Cells|Intensity_LowerQuartileIntensity_WGA,Cells|Intensity_LowerQuartileIntensity_Phalloidin,Cells|Intensity_LowerQuartileIntensity_DNA,Cells|Intensity_MassDisplacement_Vinculin,Cells|Intensity_MassDisplacement_DNA,Cells|Intensity_MassDisplacement_WGA,Cells|Intensity_MassDisplacement_Phalloidin,Cells|Location_MaxIntensity_Z_Vinculin,Cells|Location_MaxIntensity_Z_WGA,Cells|Location_MaxIntensity_Z_Phalloidin,Cells|Location_MaxIntensity_Z_DNA,Cells|Location_MaxIntensity_X_Phalloidin,Cells|Location_MaxIntensity_X_DNA,Cells|Location_MaxIntensity_X_WGA,Cells|Location_MaxIntensity_X_Vinculin,Cells|Location_MaxIntensity_Y_WGA,Cells|Location_MaxIntensity_Y_DNA,Cells|Location_MaxIntensity_Y_Phalloidin,Cells|Location_MaxIntensity_Y_Vinculin,Cells|Location_CenterMassIntensity_Z_DNA,Cells|Location_CenterMassIntensity_Z_WGA,Cells|Location_CenterMassIntensity_Z_Phalloidin,Cells|Location_CenterMassIntensity_Z_Vinculin,Cells|Location_CenterMassIntensity_X_Vinculin,Cells|Location_CenterMassIntensity_X_DNA,Cells|Location_CenterMassIntensity_X_Phalloidin,Cells|Location_CenterMassIntensity_X_WGA,Cells|Location_CenterMassIntensity_Y_DNA,Cells|Location_CenterMassIntensity_Y_Vinculin,Cells|Location_CenterMassIntensity_Y_WGA,Cells|Location_CenterMassIntensity_Y_Phalloidin,Cells|Children_Cytoplasm_Count,Image|Texture_InfoMeas2_Vinculin_3_02_256,Image|Texture_InfoMeas2_Vinculin_3_00_256,Image|Texture_InfoMeas2_Vinculin_3_01_256,Image|Texture_InfoMeas2_Vinculin_3_03_256,Image|Texture_InfoMeas2_Vinculin_5_02_256,Image|Texture_InfoMeas2_Vinculin_5_03_256,Image|Texture_InfoMeas2_Vinculin_5_00_256,Image|Texture_InfoMeas2_Vinculin_5_01_256,Image|Texture_InfoMeas2_Vinculin_10_01_256,Image|Texture_InfoMeas2_Vinculin_10_02_256,Image|Texture_InfoMeas2_Vinculin_10_00_256,Image|Texture_InfoMeas2_Vinculin_10_03_256,Image|Texture_InfoMeas2_Vinculin_20_02_256,Image|Texture_InfoMeas2_Vinculin_20_03_256,Image|Texture_InfoMeas2_Vinculin_20_01_256,Image|Texture_InfoMeas2_Vinculin_20_00_256,Image|Texture_InfoMeas2_DNA_3_03_256,Image|Texture_InfoMeas2_DNA_3_01_256,Image|Texture_InfoMeas2_DNA_3_00_256,Image|Texture_InfoMeas2_DNA_3_02_256,Image|Texture_InfoMeas2_DNA_5_00_256,Image|Texture_InfoMeas2_DNA_5_01_256,Image|Texture_InfoMeas2_DNA_5_03_256,Image|Texture_InfoMeas2_DNA_5_02_256,Image|Texture_InfoMeas2_DNA_10_00_256,Image|Texture_InfoMeas2_DNA_10_03_256,Image|Texture_InfoMeas2_DNA_10_01_256,Image|Texture_InfoMeas2_DNA_10_02_256,Image|Texture_InfoMeas2_DNA_20_02_256,Image|Textu

re_InfoMeas2_DNA_20_03_256,Image|Texture_InfoMeas2_DNA_20_00_256,Image|Texture_InfoMeas2_DNA_20_0
1_256,Image|Texture_InfoMeas2_Phalloidin_20_00_256,Image|Texture_InfoMeas2_Phalloidin_20_03_256,Image|Text
ure_InfoMeas2_Phalloidin_20_02_256,Image|Texture_InfoMeas2_Phalloidin_20_01_256,Image|Texture_InfoMeas2_P
halloidin_10_03_256,Image|Texture_InfoMeas2_Phalloidin_10_00_256,Image|Texture_InfoMeas2_Phalloidin_10_02_
256,Image|Texture_InfoMeas2_Phalloidin_10_01_256,Image|Texture_InfoMeas2_Phalloidin_3_01_256,Image|Texture
_InfoMeas2_Phalloidin_3_00_256,Image|Texture_InfoMeas2_Phalloidin_3_02_256,Image|Texture_InfoMeas2_Phalloi
din_3_03_256,Image|Texture_InfoMeas2_Phalloidin_5_00_256,Image|Texture_InfoMeas2_Phalloidin_5_01_256,Imag
e|Texture_InfoMeas2_Phalloidin_5_02_256,Image|Texture_InfoMeas2_Phalloidin_5_03_256,Image|Texture_InfoMeas
2_WGA_5_03_256,Image|Texture_InfoMeas2_WGA_5_00_256,Image|Texture_InfoMeas2_WGA_5_01_256,Image|T
exture_InfoMeas2_WGA_5_02_256,Image|Texture_InfoMeas2_WGA_3_00_256,Image|Texture_InfoMeas2_WGA_3_
03_256,Image|Texture_InfoMeas2_WGA_3_01_256,Image|Texture_InfoMeas2_WGA_3_02_256,Image|Texture_Info
Meas2_WGA_20_02_256,Image|Texture_InfoMeas2_WGA_20_03_256,Image|Texture_InfoMeas2_WGA_20_01_256
,Image|Texture_InfoMeas2_WGA_20_00_256,Image|Texture_InfoMeas2_WGA_10_01_256,Image|Texture_InfoMeas
2_WGA_10_00_256,Image|Texture_InfoMeas2_WGA_10_03_256,Image|Texture_InfoMeas2_WGA_10_02_256,Imag
e|Texture_SumEntropy_WGA_5_03_256,Image|Texture_SumEntropy_WGA_5_00_256,Image|Texture_SumEntropy_
WGA_5_01_256,Image|Texture_SumEntropy_WGA_5_02_256,Image|Texture_SumEntropy_WGA_10_00_256,Image
|Texture_SumEntropy_WGA_10_03_256,Image|Texture_SumEntropy_WGA_10_01_256,Image|Texture_SumEntropy
_WGA_10_02_256,Image|Texture_SumEntropy_WGA_20_02_256,Image|Texture_SumEntropy_WGA_20_00_256,Im
age|Texture_SumEntropy_WGA_20_03_256,Image|Texture_SumEntropy_WGA_20_01_256,Image|Texture_SumEntro
py_WGA_3_01_256,Image|Texture_SumEntropy_WGA_3_00_256,Image|Texture_SumEntropy_WGA_3_02_256,Ima
ge|Texture_SumEntropy_WGA_3_03_256,Image|Texture_SumEntropy_Phalloidin_20_01_256,Image|Texture_SumEnt
ropy_Phalloidin_20_03_256,Image|Texture_SumEntropy_Phalloidin_20_02_256,Image|Texture_SumEntropy_Phalloid
in_20_00_256,Image|Texture_SumEntropy_Phalloidin_3_00_256,Image|Texture_SumEntropy_Phalloidin_3_03_256,I
mage|Texture_SumEntropy_Phalloidin_3_01_256,Image|Texture_SumEntropy_Phalloidin_3_02_256,Image|Texture_S
umEntropy_Phalloidin_10_02_256,Image|Texture_SumEntropy_Phalloidin_10_01_256,Image|Texture_SumEntropy_P
halloidin_10_00_256,Image|Texture_SumEntropy_Phalloidin_10_03_256,Image|Texture_SumEntropy_Phalloidin_5_0
1_256,Image|Texture_SumEntropy_Phalloidin_5_02_256,Image|Texture_SumEntropy_Phalloidin_5_00_256,Image|Te
xture_SumEntropy_Phalloidin_5_03_256,Image|Texture_SumEntropy_Vinculin_20_00_256,Image|Texture_SumEntro
py_Vinculin_20_02_256,Image|Texture_SumEntropy_Vinculin_20_03_256,Image|Texture_SumEntropy_Vinculin_20_
01_256,Image|Texture_SumEntropy_Vinculin_3_03_256,Image|Texture_SumEntropy_Vinculin_3_00_256,Image|Text
ure_SumEntropy_Vinculin_3_02_256,Image|Texture_SumEntropy_Vinculin_3_01_256,Image|Texture_SumEntropy_V
inculin_10_02_256,Image|Texture_SumEntropy_Vinculin_10_03_256,Image|Texture_SumEntropy_Vinculin_10_00_2
56,Image|Texture_SumEntropy_Vinculin_10_01_256,Image|Texture_SumEntropy_Vinculin_5_01_256,Image|Texture_
SumEntropy_Vinculin_5_02_256,Image|Texture_SumEntropy_Vinculin_5_03_256,Image|Texture_SumEntropy_Vincu
lin_5_00_256,Image|Texture_SumEntropy_DNA_10_01_256,Image|Texture_SumEntropy_DNA_10_03_256,Image|Te
xture_SumEntropy_DNA_10_02_256,Image|Texture_SumEntropy_DNA_10_00_256,Image|Texture_SumEntropy_DN
A_20_00_256,Image|Texture_SumEntropy_DNA_20_01_256,Image|Texture_SumEntropy_DNA_20_03_256,Image|Te
xture_SumEntropy_DNA_20_02_256,Image|Texture_SumEntropy_DNA_3_02_256,Image|Texture_SumEntropy_DN
A_3_00_256,Image|Texture_SumEntropy_DNA_3_03_256,Image|Texture_SumEntropy_DNA_3_01_256,Image|Textu
re_SumEntropy_DNA_5_03_256,Image|Texture_SumEntropy_DNA_5_02_256,Image|Texture_SumEntropy_DNA_5_
01_256,Image|Texture_SumEntropy_DNA_5_00_256,Image|Texture_InfoMeas1_DNA_5_00_256,Image|Texture_Info
Meas1_DNA_5_03_256,Image|Texture_InfoMeas1_DNA_5_01_256,Image|Texture_InfoMeas1_DNA_5_02_256,Imag
e|Texture_InfoMeas1_DNA_20_02_256,Image|Texture_InfoMeas1_DNA_20_00_256,Image|Texture_InfoMeas1_DN
A_20_03_256,Image|Texture_InfoMeas1_DNA_20_01_256,Image|Texture_InfoMeas1_DNA_10_02_256,Image|Textu
re_InfoMeas1_DNA_10_03_256,Image|Texture_InfoMeas1_DNA_10_00_256,Image|Texture_InfoMeas1_DNA_10_0
1_256,Image|Texture_InfoMeas1_DNA_3_01_256,Image|Texture_InfoMeas1_DNA_3_02_256,Image|Texture_InfoMe
as1_DNA_3_00_256,Image|Texture_InfoMeas1_DNA_3_03_256,Image|Texture_InfoMeas1_Vinculin_20_00_256,Ima
ge|Texture_InfoMeas1_Vinculin_20_03_256,Image|Texture_InfoMeas1_Vinculin_20_02_256,Image|Texture_InfoMea
s1_Vinculin_20_01_256,Image|Texture_InfoMeas1_Vinculin_10_03_256,Image|Texture_InfoMeas1_Vinculin_10_01_
256,Image|Texture_InfoMeas1_Vinculin_10_02_256,Image|Texture_InfoMeas1_Vinculin_10_00_256,Image|Texture_I
nfoMeas1_Vinculin_5_02_256,Image|Texture_InfoMeas1_Vinculin_5_00_256,Image|Texture_InfoMeas1_Vinculin_5_
03_256,Image|Texture_InfoMeas1_Vinculin_5_01_256,Image|Texture_InfoMeas1_Vinculin_3_02_256,Image|Texture
_InfoMeas1_Vinculin_3_00_256,Image|Texture_InfoMeas1_Vinculin_3_01_256,Image|Texture_InfoMeas1_Vinculin_
3_03_256,Image|Texture_InfoMeas1_Phalloidin_20_03_256,Image|Texture_InfoMeas1_Phalloidin_20_01_256,Image|

Texture_InfoMeas1_Phalloidin_20_02_256,Image|Texture_InfoMeas1_Phalloidin_20_00_256,Image|Texture_InfoMea s1_Phalloidin_3_03_256,Image|Texture_InfoMeas1_Phalloidin_3_00_256,Image|Texture_InfoMeas1_Phalloidin_3_01 _256,Image|Texture_InfoMeas1_Phalloidin_3_02_256,Image|Texture_InfoMeas1_Phalloidin_10_01_256,Image|Textur e_InfoMeas1_Phalloidin_10_02_256,Image|Texture_InfoMeas1_Phalloidin_10_03_256,Image|Texture_InfoMeas1_Pha lloidin_10_00_256,Image|Texture_InfoMeas1_Phalloidin_5_02_256,Image|Texture_InfoMeas1_Phalloidin_5_00_256,I mage|Texture_InfoMeas1_Phalloidin_5_03_256,Image|Texture_InfoMeas1_Phalloidin_5_01_256,Image|Texture_Info Meas1_WGA_5_03_256,Image|Texture_InfoMeas1_WGA_5_01_256,Image|Texture_InfoMeas1_WGA_5_02_256,Im age|Texture_InfoMeas1_WGA_5_00_256,Image|Texture_InfoMeas1_WGA_10_00_256,Image|Texture_InfoMeas1_W GA_10_02_256,Image|Texture_InfoMeas1_WGA_10_01_256,Image|Texture_InfoMeas1_WGA_10_03_256,Image|Te xture_InfoMeas1_WGA_20_00_256,Image|Texture_InfoMeas1_WGA_20_03_256,Image|Texture_InfoMeas1_WGA_ 20_01_256,Image|Texture_InfoMeas1_WGA_20_02_256,Image|Texture_InfoMeas1_WGA_3_02_256,Image|Texture_ InfoMeas1_WGA_3_01_256,Image|Texture_InfoMeas1_WGA_3_03_256,Image|Texture_InfoMeas1_WGA_3_00_256 ,Image|Texture_DifferenceVariance_DNA_3_00_256,Image|Texture_DifferenceVariance_DNA_3_03_256,Image|Text ure_DifferenceVariance_DNA_3_02_256,Image|Texture_DifferenceVariance_DNA_3_01_256,Image|Texture_Differe nceVariance_DNA_20_02_256,Image|Texture_DifferenceVariance_DNA_20_00_256,Image|Texture_DifferenceVaria nce_DNA_20_03_256,Image|Texture_DifferenceVariance_DNA_20_01_256,Image|Texture_DifferenceVariance_DNA _10_03_256,Image|Texture_DifferenceVariance_DNA_10_02_256,Image|Texture_DifferenceVariance_DNA_10_00_2 56,Image|Texture_DifferenceVariance_DNA_10_01_256,Image|Texture_DifferenceVariance_DNA_5_03_256,Image|T exture_DifferenceVariance_DNA_5_02_256,Image|Texture_DifferenceVariance_DNA_5_01_256,Image|Texture_Diff erenceVariance_DNA_5_00_256,Image|Texture_DifferenceVariance_Vinculin_20_03_256,Image|Texture_Difference Variance_Vinculin_20_00_256,Image|Texture_DifferenceVariance_Vinculin_20_01_256,Image|Texture_DifferenceVa riance_Vinculin_20_02_256,Image|Texture_DifferenceVariance_Vinculin_3_01_256,Image|Texture_DifferenceVarian ce_Vinculin_3_03_256,Image|Texture_DifferenceVariance_Vinculin_3_02_256,Image|Texture_DifferenceVariance_Vi nculin_3_00_256,Image|Texture_DifferenceVariance_Vinculin_10_03_256,Image|Texture_DifferenceVariance_Vincul in_10_00_256,Image|Texture_DifferenceVariance_Vinculin_10_01_256,Image|Texture_DifferenceVariance_Vinculin_ 10_02_256,Image|Texture_DifferenceVariance_Vinculin_5_00_256,Image|Texture_DifferenceVariance_Vinculin_5_0 2_256,Image|Texture_DifferenceVariance_Vinculin_5_01_256,Image|Texture_DifferenceVariance_Vinculin_5_03_25 6,Image|Texture_DifferenceVariance_WGA_20_03_256,Image|Texture_DifferenceVariance_WGA_20_00_256,Image| Texture_DifferenceVariance_WGA_20_01_256,Image|Texture_DifferenceVariance_WGA_20_02_256,Image|Texture_ DifferenceVariance_WGA_5_00_256,Image|Texture_DifferenceVariance_WGA_5_02_256,Image|Texture_Difference Variance_WGA_5_01_256,Image|Texture_DifferenceVariance_WGA_5_03_256,Image|Texture_DifferenceVariance_ WGA_10_02_256,Image|Texture_DifferenceVariance_WGA_10_01_256,Image|Texture_DifferenceVariance_WGA_1 0_00_256,Image|Texture_DifferenceVariance_WGA_10_03_256,Image|Texture_DifferenceVariance_WGA_3_00_256 ,Image|Texture_DifferenceVariance_WGA_3_02_256,Image|Texture_DifferenceVariance_WGA_3_03_256,Image|Tex ture_DifferenceVariance_WGA_3_01_256,Image|Texture_DifferenceVariance_Phalloidin_3_00_256,Image|Texture_D ifferenceVariance_Phalloidin_3_03_256,Image|Texture_DifferenceVariance_Phalloidin_3_01_256,Image|Texture_Diff erenceVariance_Phalloidin_3_02_256,Image|Texture_DifferenceVariance_Phalloidin_5_02_256,Image|Texture_Differ enceVariance_Phalloidin_5_00_256,Image|Texture_DifferenceVariance_Phalloidin_5_03_256,Image|Texture_Differen ceVariance_Phalloidin_5_01_256,Image|Texture_DifferenceVariance_Phalloidin_20_01_256,Image|Texture_Differenc eVariance_Phalloidin_20_03_256,Image|Texture_DifferenceVariance_Phalloidin_20_00_256,Image|Texture_Differenc eVariance_Phalloidin_20_02_256,Image|Texture_DifferenceVariance_Phalloidin_10_02_256,Image|Texture_Differenc eVariance_Phalloidin_10_00_256,Image|Texture_DifferenceVariance_Phalloidin_10_03_256,Image|Texture_Differenc eVariance_Phalloidin_10_01_256,Image|Texture_SumAverage_WGA_10_03_256,Image|Texture_SumAverage_WGA _10_00_256,Image|Texture_SumAverage_WGA_10_01_256,Image|Texture_SumAverage_WGA_10_02_256,Image|T exture_SumAverage_WGA_5_01_256,Image|Texture_SumAverage_WGA_5_00_256,Image|Texture_SumAverage_W GA_5_02_256,Image|Texture_SumAverage_WGA_5_03_256,Image|Texture_SumAverage_WGA_3_02_256,Image|Te xture_SumAverage_WGA_3_00_256,Image|Texture_SumAverage_WGA_3_01_256,Image|Texture_SumAverage_WG A_3_03_256,Image|Texture_SumAverage_WGA_20_00_256,Image|Texture_SumAverage_WGA_20_01_256,Image|T exture_SumAverage_WGA_20_03_256,Image|Texture_SumAverage_WGA_20_02_256,Image|Texture_SumAverage_ DNA_10_01_256,Image|Texture_SumAverage_DNA_10_02_256,Image|Texture_SumAverage_DNA_10_00_256,Ima ge|Texture_SumAverage_DNA_10_03_256,Image|Texture_SumAverage_DNA_5_00_256,Image|Texture_SumAverag e_DNA_5_02_256,Image|Texture_SumAverage_DNA_5_01_256,Image|Texture_SumAverage_DNA_5_03_256,Image |Texture_SumAverage_DNA_20_00_256,Image|Texture_SumAverage_DNA_20_02_256,Image|Texture_SumAverage _DNA_20_03_256,Image|Texture_SumAverage_DNA_20_01_256,Image|Texture_SumAverage_DNA_3_00_256,Ima ge|Texture_SumAverage_DNA_3_02_256,Image|Texture_SumAverage_DNA_3_01_256,Image|Texture_SumAverage_DNA_3_03_256,Image|Texture_SumAverage_Vinculin_5_00_256,Image|Texture_SumAverage_Vinculin_5_02_256,Image|Texture_SumAverage_Vinculin_5_03_256,Image|Texture_SumAverage_Vinculin_5_01_256,Image|Texture_SumAverage_Vinculin_10_02_256,Image|Texture_SumAverage_Vinculin_10_03_256,Image|Texture_SumAverage_Vinculin_10_01_256,Image|Texture_SumAverage_Vinculin_10_00_256,Image|Texture_SumAverage_Vinculin_3_01_256,Image|Texture_SumAverage_Vinculin_3_03_256,Image|Texture_SumAverage_Vinculin_3_02_256,Image|Texture_SumAverage_Vinculin_3_00_256,Image|Texture_SumAverage_Vinculin_20_03_256,Image|Texture_SumAverage_Vinculin_20_01_256,Image|Texture_SumAverage_Vinculin_20_02_256,Image|Texture_SumAverage_Vinculin_20_00_256,Image|Texture_SumAverage_Phalloidin_5_02_256,Image|Texture_SumAverage_Phalloidin_5_03_256,Image|Texture_SumAverage_Phalloidin_5_00_256,Image|Texture_SumAverage_Phalloidin_5_01_256,Image|Texture_SumAverage_Phalloidin_3_00_256,Image|Texture_SumAverage_Phalloidin_3_01_256,Image|Texture_SumAverage_Phalloidin_3_03_256,Image|Texture_SumAverage_Phalloidin_3_02_256,Image|Texture_SumAverage_Phalloidin_20_01_256,Image|Texture_SumAverage_Phalloidin_20_03_256,Image|Texture_SumAverage_Phalloidin_20_02_256,Image|Texture_SumAverage_Phalloidin_20_00_256,Image|Texture_SumAverage_Phalloidin_10_03_256,Image|Texture_SumAverage_Phalloidin_10_01_256,Image|Texture_SumAverage_Phalloidin_10_02_256,Image|Texture_SumAverage_Phalloidin_10_00_256,Image|Texture_Contrast_DNA_20_01_256,Image|Texture_Contrast_DNA_20_03_256,Image|Texture_Contrast_DNA_20_02_256,Image|Texture_Contrast_DNA_20_00_256,Image|Texture_Contrast_DNA_5_02_256,Image|Texture_Contrast_DNA_5_01_256,Image|Texture_Contrast_DNA_5_00_256,Image|Texture_Contrast_DNA_5_03_256,Image|Texture_Contrast_DNA_10_02_256,Image|Texture_Contrast_DNA_10_03_256,Image|Texture_Contrast_DNA_10_01_256,Image|Texture_Contrast_DNA_10_00_256,Image|Texture_Contrast_DNA_3_02_256,Image|Texture_Contrast_DNA_3_01_256,Image|Texture_Contrast_DNA_3_03_256,Image|Texture_Contrast_DNA_3_00_256,Image|Texture_Contrast_Phalloidin_20_03_256,Image|Texture_Contrast_Phalloidin_20_00_256,Image|Texture_Contrast_Phalloidin_20_01_256,Image|Texture_Contrast_Phalloidin_20_02_256,Image|Texture_Contrast_Phalloidin_5_00_256,Image|Texture_Contrast_Phalloidin_5_01_256,Image|Texture_Contrast_Phalloidin_5_03_256,Image|Texture_Contrast_Phalloidin_5_02_256,Image|Texture_Contrast_Phalloidin_3_01_256,Image|Texture_Contrast_Phalloidin_3_02_256,Image|Texture_Contrast_Phalloidin_3_00_256,Image|Texture_Contrast_Phalloidin_3_03_256,Image|Texture_Contrast_Phalloidin_10_03_256,Image|Texture_Contrast_Phalloidin_10_01_256,Image|Texture_Contrast_Phalloidin_10_00_256,Image|Texture_Contrast_Phalloidin_10_02_256,Image|Texture_Contrast_WGA_20_01_256,Image|Texture_Contrast_WGA_20_03_256,Image|Texture_Contrast_WGA_20_02_256,Image|Texture_Contrast_WGA_20_00_256,Image|Texture_Contrast_WGA_5_00_256,Image|Texture_Contrast_WGA_5_01_256,Image|Texture_Contrast_WGA_5_03_256,Image|Texture_Contrast_WGA_5_02_256,Image|Texture_Contrast_WGA_3_01_256,Image|Texture_Contrast_WGA_3_03_256,Image|Texture_Contrast_WGA_3_00_256,Image|Texture_Contrast_WGA_3_02_256,Image|Texture_Contrast_WGA_10_00_256,Image|Texture_Contrast_WGA_10_02_256,Image|Texture_Contrast_WGA_10_01_256,Image|Texture_Contrast_WGA_10_03_256,Image|Texture_Contrast_Vinculin_3_02_256,Image|Texture_Contrast_Vinculin_3_01_256,Image|Texture_Contrast_Vinculin_3_00_256,Image|Texture_Contrast_Vinculin_3_03_256,Image|Texture_Contrast_Vinculin_20_03_256,Image|Texture_Contrast_Vinculin_20_01_256,Image|Texture_Contrast_Vinculin_20_02_256,Image|Texture_Contrast_Vinculin_20_00_256,Image|Texture_Contrast_Vinculin_10_02_256,Image|Texture_Contrast_Vinculin_10_00_256,Image|Texture_Contrast_Vinculin_10_03_256,Image|Texture_Contrast_Vinculin_10_01_256,Image|Texture_Contrast_Vinculin_5_02_256,Image|Texture_Contrast_Vinculin_5_03_256,Image|Texture_Contrast_Vinculin_5_00_256,Image|Texture_Contrast_Vinculin_5_01_256,Image|Texture_AngularSecondMoment_Vinculin_5_01_256,Image|Texture_AngularSecondMoment_Vinculin_5_02_256,Image|Texture_AngularSecondMoment_Vinculin_5_00_256,Image|Texture_AngularSecondMoment_Vinculin_5_03_256,Image|Texture_AngularSecondMoment_Vinculin_3_02_256,Image|Texture_AngularSecondMoment_Vinculin_3_00_256,Image|Texture_AngularSecondMoment_Vinculin_3_03_256,Image|Texture_AngularSecondMoment_Vinculin_3_01_256,Image|Texture_AngularSecondMoment_Vinculin_20_01_256,Image|Texture_AngularSecondMoment_Vinculin_20_02_256,Image|Texture_AngularSecondMoment_Vinculin_20_00_256,Image|Texture_AngularSecondMoment_Vinculin_20_03_256,Image|Texture_AngularSecondMoment_Vinculin_10_00_256,Image|Texture_AngularSecondMoment_Vinculin_10_01_256,Image|Texture_AngularSecondMoment_Vinculin_10_02_256,Image|Texture_AngularSecondMoment_Vinculin_10_03_256,Image|Texture_AngularSecondMoment_WGA_5_02_256,Image|Texture_AngularSecondMoment_WGA_5_01_256,Image|Texture_AngularSecondMoment_WGA_5_03_256,Image|Texture_AngularSecondMoment_WGA_5_00_256,Image|Texture_AngularSecondMoment_WGA_3_03_256,Image|Texture_AngularSecondMoment_WGA_3_01_256,Image|Texture_AngularSecondMoment_WGA_3_02_256,Image|Texture_AngularSecondMoment_WGA_3_00_256,Image|Texture_AngularSecondMoment_WGA_10_03_256,Image|Texture_AngularSecondMoment_WGA_10_00_256,Image|Texture_AngularSecondMoment_WGA_10_02_256,Image|Texture_AngularSecondMoment_WGA_10_01_256,Image|Texture_AngularSecondMoment_WGA_20_00

_256,Image|Texture_AngularSecondMoment_WGA_20_01_256,Image|Texture_AngularSecondMoment_WGA_20_03
_256,Image|Texture_AngularSecondMoment_WGA_20_02_256,Image|Texture_AngularSecondMoment_DNA_10_01
_256,Image|Texture_AngularSecondMoment_DNA_10_02_256,Image|Texture_AngularSecondMoment_DNA_10_03_
256,Image|Texture_AngularSecondMoment_DNA_10_00_256,Image|Texture_AngularSecondMoment_DNA_5_00_25
6,Image|Texture_AngularSecondMoment_DNA_5_01_256,Image|Texture_AngularSecondMoment_DNA_5_03_256,I
mage|Texture_AngularSecondMoment_DNA_5_02_256,Image|Texture_AngularSecondMoment_DNA_20_00_256,Im
age|Texture_AngularSecondMoment_DNA_20_03_256,Image|Texture_AngularSecondMoment_DNA_20_02_256,Ima
ge|Texture_AngularSecondMoment_DNA_20_01_256,Image|Texture_AngularSecondMoment_DNA_3_00_256,Image|
Texture_AngularSecondMoment_DNA_3_03_256,Image|Texture_AngularSecondMoment_DNA_3_02_256,Image|Tex
ture_AngularSecondMoment_DNA_3_01_256,Image|Texture_AngularSecondMoment_Phalloidin_3_02_256,Image|Te
xture_AngularSecondMoment_Phalloidin_3_03_256,Image|Texture_AngularSecondMoment_Phalloidin_3_01_256,Im
age|Texture_AngularSecondMoment_Phalloidin_3_00_256,Image|Texture_AngularSecondMoment_Phalloidin_10_00_
256,Image|Texture_AngularSecondMoment_Phalloidin_10_03_256,Image|Texture_AngularSecondMoment_Phalloidin
_10_02_256,Image|Texture_AngularSecondMoment_Phalloidin_10_01_256,Image|Texture_AngularSecondMoment_P
halloidin_5_00_256,Image|Texture_AngularSecondMoment_Phalloidin_5_01_256,Image|Texture_AngularSecondMom
ent_Phalloidin_5_02_256,Image|Texture_AngularSecondMoment_Phalloidin_5_03_256,Image|Texture_AngularSecon
dMoment_Phalloidin_20_03_256,Image|Texture_AngularSecondMoment_Phalloidin_20_02_256,Image|Texture_Angu
larSecondMoment_Phalloidin_20_01_256,Image|Texture_AngularSecondMoment_Phalloidin_20_00_256,Image|Textu
re_Correlation_WGA_5_03_256,Image|Texture_Correlation_WGA_5_01_256,Image|Texture_Correlation_WGA_5_00
_256,Image|Texture_Correlation_WGA_5_02_256,Image|Texture_Correlation_WGA_10_01_256,Image|Texture_Corr
elation_WGA_10_03_256,Image|Texture_Correlation_WGA_10_02_256,Image|Texture_Correlation_WGA_10_00_25
6,Image|Texture_Correlation_WGA_20_00_256,Image|Texture_Correlation_WGA_20_03_256,Image|Texture_Correla
tion_WGA_20_01_256,Image|Texture_Correlation_WGA_20_02_256,Image|Texture_Correlation_WGA_3_03_256,Im
age|Texture_Correlation_WGA_3_01_256,Image|Texture_Correlation_WGA_3_00_256,Image|Texture_Correlation_W
GA_3_02_256,Image|Texture_Correlation_Vinculin_5_00_256,Image|Texture_Correlation_Vinculin_5_01_256,Image|
Texture_Correlation_Vinculin_5_02_256,Image|Texture_Correlation_Vinculin_5_03_256,Image|Texture_Correlation_
Vinculin_20_02_256,Image|Texture_Correlation_Vinculin_20_00_256,Image|Texture_Correlation_Vinculin_20_03_25
6,Image|Texture_Correlation_Vinculin_20_01_256,Image|Texture_Correlation_Vinculin_10_00_256,Image|Texture_C
orrelation_Vinculin_10_02_256,Image|Texture_Correlation_Vinculin_10_03_256,Image|Texture_Correlation_Vinculin
_10_01_256,Image|Texture_Correlation_Vinculin_3_00_256,Image|Texture_Correlation_Vinculin_3_02_256,Image|Te
xture_Correlation_Vinculin_3_01_256,Image|Texture_Correlation_Vinculin_3_03_256,Image|Texture_Correlation_Ph
alloidin_5_02_256,Image|Texture_Correlation_Phalloidin_5_00_256,Image|Texture_Correlation_Phalloidin_5_01_256,
Image|Texture_Correlation_Phalloidin_5_03_256,Image|Texture_Correlation_Phalloidin_10_00_256,Image|Texture_C
orrelation_Phalloidin_10_02_256,Image|Texture_Correlation_Phalloidin_10_03_256,Image|Texture_Correlation_Phall
oidin_10_01_256,Image|Texture_Correlation_Phalloidin_3_02_256,Image|Texture_Correlation_Phalloidin_3_01_256,I
mage|Texture_Correlation_Phalloidin_3_00_256,Image|Texture_Correlation_Phalloidin_3_03_256,Image|Texture_Corr
elation_Phalloidin_20_02_256,Image|Texture_Correlation_Phalloidin_20_01_256,Image|Texture_Correlation_Phalloidi
n_20_03_256,Image|Texture_Correlation_Phalloidin_20_00_256,Image|Texture_Correlation_DNA_3_00_256,Image|T
exture_Correlation_DNA_3_02_256,Image|Texture_Correlation_DNA_3_01_256,Image|Texture_Correlation_DNA_3_
03_256,Image|Texture_Correlation_DNA_10_03_256,Image|Texture_Correlation_DNA_10_01_256,Image|Texture_Co
rrelation_DNA_10_00_256,Image|Texture_Correlation_DNA_10_02_256,Image|Texture_Correlation_DNA_5_00_256
,Image|Texture_Correlation_DNA_5_03_256,Image|Texture_Correlation_DNA_5_02_256,Image|Texture_Correlation_
DNA_5_01_256,Image|Texture_Correlation_DNA_20_03_256,Image|Texture_Correlation_DNA_20_02_256,Image|Te
xture_Correlation_DNA_20_01_256,Image|Texture_Correlation_DNA_20_00_256,Image|Texture_Variance_WGA_10
_00_256,Image|Texture_Variance_WGA_10_01_256,Image|Texture_Variance_WGA_10_02_256,Image|Texture_Vari
ance_WGA_10_03_256,Image|Texture_Variance_WGA_3_00_256,Image|Texture_Variance_WGA_3_03_256,Image|
Texture_Variance_WGA_3_02_256,Image|Texture_Variance_WGA_3_01_256,Image|Texture_Variance_WGA_5_01_
256,Image|Texture_Variance_WGA_5_03_256,Image|Texture_Variance_WGA_5_02_256,Image|Texture_Variance_W
GA_5_00_256,Image|Texture_Variance_WGA_20_03_256,Image|Texture_Variance_WGA_20_02_256,Image|Texture
_Variance_WGA_20_01_256,Image|Texture_Variance_WGA_20_00_256,Image|Texture_Variance_Vinculin_3_03_25
6,Image|Texture_Variance_Vinculin_3_02_256,Image|Texture_Variance_Vinculin_3_01_256,Image|Texture_Variance
_Vinculin_3_00_256,Image|Texture_Variance_Vinculin_5_02_256,Image|Texture_Variance_Vinculin_5_00_256,Imag
e|Texture_Variance_Vinculin_5_01_256,Image|Texture_Variance_Vinculin_5_03_256,Image|Texture_Variance_Vincu
lin_20_03_256,Image|Texture_Variance_Vinculin_20_01_256,Image|Texture_Variance_Vinculin_20_00_256,Image|T

exture_Variance_Vinculin_20_02_256,Image|Texture_Variance_Vinculin_10_01_256,Image|Texture_Variance_Vinculin_10_00_256,Image|Texture_Variance_Vinculin_10_03_256,Image|Texture_Variance_Vinculin_10_02_256,Image|Texture_Variance_Phalloidin_20_02_256,Image|Texture_Variance_Phalloidin_20_00_256,Image|Texture_Variance_Phalloidin_20_01_256,Image|Texture_Variance_Phalloidin_20_03_256,Image|Texture_Variance_Phalloidin_3_03_256,Image|Texture_Variance_Phalloidin_3_02_256,Image|Texture_Variance_Phalloidin_3_00_256,Image|Texture_Variance_Phalloidin_3_01_256,Image|Texture_Variance_Phalloidin_5_03_256,Image|Texture_Variance_Phalloidin_5_00_256,Image|Texture_Variance_Phalloidin_5_01_256,Image|Texture_Variance_Phalloidin_5_02_256,Image|Texture_Variance_Phalloidin_10_03_256,Image|Texture_Variance_Phalloidin_10_01_256,Image|Texture_Variance_Phalloidin_10_00_256,Image|Texture_Variance_Phalloidin_10_02_256,Image|Texture_Variance_DNA_10_00_256,Image|Texture_Variance_DNA_10_03_256,Image|Texture_Variance_DNA_10_02_256,Image|Texture_Variance_DNA_10_01_256,Image|Texture_Variance_DNA_5_00_256,Image|Texture_Variance_DNA_5_03_256,Image|Texture_Variance_DNA_5_02_256,Image|Texture_Variance_DNA_5_01_256,Image|Texture_Variance_DNA_20_02_256,Image|Texture_Variance_DNA_20_00_256,Image|Texture_Variance_DNA_20_03_256,Image|Texture_Variance_DNA_20_01_256,Image|Texture_Variance_DNA_3_03_256,Image|Texture_Variance_DNA_3_01_256,Image|Texture_Variance_DNA_3_02_256,Image|Texture_Variance_DNA_3_00_256,Image|Texture_Entropy_WGA_5_01_256,Image|Texture_Entropy_WGA_5_00_256,Image|Texture_Entropy_WGA_5_03_256,Image|Texture_Entropy_WGA_5_02_256,Image|Texture_Entropy_WGA_20_00_256,Image|Texture_Entropy_WGA_20_01_256,Image|Texture_Entropy_WGA_20_02_256,Image|Texture_Entropy_WGA_20_03_256,Image|Texture_Entropy_WGA_3_03_256,Image|Texture_Entropy_WGA_3_00_256,Image|Texture_Entropy_WGA_3_02_256,Image|Texture_Entropy_WGA_3_01_256,Image|Texture_Entropy_WGA_10_01_256,Image|Texture_Entropy_WGA_10_00_256,Image|Texture_Entropy_WGA_10_02_256,Image|Texture_Entropy_WGA_10_03_256,Image|Texture_Entropy_DNA_5_03_256,Image|Texture_Entropy_DNA_5_02_256,Image|Texture_Entropy_DNA_5_00_256,Image|Texture_Entropy_DNA_5_01_256,Image|Texture_Entropy_DNA_10_00_256,Image|Texture_Entropy_DNA_10_03_256,Image|Texture_Entropy_DNA_10_02_256,Image|Texture_Entropy_DNA_10_01_256,Image|Texture_Entropy_DNA_3_00_256,Image|Texture_Entropy_DNA_3_02_256,Image|Texture_Entropy_DNA_3_01_256,Image|Texture_Entropy_DNA_3_03_256,Image|Texture_Entropy_DNA_20_03_256,Image|Texture_Entropy_DNA_20_00_256,Image|Texture_Entropy_DNA_20_01_256,Image|Texture_Entropy_DNA_20_02_256,Image|Texture_Entropy_Phalloidin_20_03_256,Image|Texture_Entropy_Phalloidin_20_00_256,Image|Texture_Entropy_Phalloidin_20_01_256,Image|Texture_Entropy_Phalloidin_20_02_256,Image|Texture_Entropy_Phalloidin_5_00_256,Image|Texture_Entropy_Phalloidin_5_01_256,Image|Texture_Entropy_Phalloidin_5_03_256,Image|Texture_Entropy_Phalloidin_5_02_256,Image|Texture_Entropy_Phalloidin_3_02_256,Image|Texture_Entropy_Phalloidin_3_03_256,Image|Texture_Entropy_Phalloidin_3_01_256,Image|Texture_Entropy_Phalloidin_3_00_256,Image|Texture_Entropy_Phalloidin_10_01_256,Image|Texture_Entropy_Phalloidin_10_03_256,Image|Texture_Entropy_Phalloidin_10_00_256,Image|Texture_Entropy_Phalloidin_10_02_256,Image|Texture_Entropy_Vinculin_3_02_256,Image|Texture_Entropy_Vinculin_3_00_256,Image|Texture_Entropy_Vinculin_3_03_256,Image|Texture_Entropy_Vinculin_3_01_256,Image|Texture_Entropy_Vinculin_5_03_256,Image|Texture_Entropy_Vinculin_5_00_256,Image|Texture_Entropy_Vinculin_5_01_256,Image|Texture_Entropy_Vinculin_5_02_256,Image|Texture_Entropy_Vinculin_20_02_256,Image|Texture_Entropy_Vinculin_20_03_256,Image|Texture_Entropy_Vinculin_20_00_256,Image|Texture_Entropy_Vinculin_20_01_256,Image|Texture_Entropy_Vinculin_10_02_256,Image|Texture_Entropy_Vinculin_10_03_256,Image|Texture_Entropy_Vinculin_10_01_256,Image|Texture_Entropy_Vinculin_10_00_256,Image|Texture_SumVariance_Phalloidin_10_00_256,Image|Texture_SumVariance_Phalloidin_10_01_256,Image|Texture_SumVariance_Phalloidin_10_02_256,Image|Texture_SumVariance_Phalloidin_10_03_256,Image|Texture_SumVariance_Phalloidin_5_01_256,Image|Texture_SumVariance_Phalloidin_5_02_256,Image|Texture_SumVariance_Phalloidin_5_03_256,Image|Texture_SumVariance_Phalloidin_5_00_256,Image|Texture_SumVariance_Phalloidin_20_00_256,Image|Texture_SumVariance_Phalloidin_20_03_256,Image|Texture_SumVariance_Phalloidin_20_01_256,Image|Texture_SumVariance_Phalloidin_20_02_256,Image|Texture_SumVariance_Phalloidin_3_02_256,Image|Texture_SumVariance_Phalloidin_3_01_256,Image|Texture_SumVariance_Phalloidin_3_00_256,Image|Texture_SumVariance_Phalloidin_3_03_256,Image|Texture_SumVariance_DNA_10_03_256,Image|Texture_SumVariance_DNA_10_02_256,Image|Texture_SumVariance_DNA_10_01_256,Image|Texture_SumVariance_DNA_10_00_256,Image|Texture_SumVariance_DNA_5_00_256,Image|Texture_SumVariance_DNA_5_01_256,Image|Texture_SumVariance_DNA_5_03_256,Image|Texture_SumVariance_DNA_5_02_256,Image|Texture_SumVariance_DNA_20_01_256,Image|Texture_SumVariance_DNA_20_00_256,Image|Texture_SumVariance_DNA_20_02_256,Image|Texture_SumVariance_DNA_20_03_256,Image|Texture_SumVariance_DNA_3_03_256,Image|Texture_SumVariance_DNA_3_00_256,Image|Texture_SumVariance_DNA_3_02_256,Image|Texture_SumVariance_DNA_3_01_256,Image|Texture_SumVariance_WGA_20_01_256,Image|Texture_SumVariance_WGA_20_02_256,Image|Texture_SumVariance_WGA_20_03_256,Image|Texture_SumVariance_WGA_20_00_256,Image|Texture_SumVariance_WGA_3_03_256,Im

age|Texture_SumVariance_WGA_3_02_256,Image|Texture_SumVariance_WGA_3_01_256,Image|Texture_SumVaria
nce_WGA_3_00_256,Image|Texture_SumVariance_WGA_5_01_256,Image|Texture_SumVariance_WGA_5_02_256,I
mage|Texture_SumVariance_WGA_5_03_256,Image|Texture_SumVariance_WGA_5_00_256,Image|Texture_SumVar
iance_WGA_10_02_256,Image|Texture_SumVariance_WGA_10_03_256,Image|Texture_SumVariance_WGA_10_01_
256,Image|Texture_SumVariance_WGA_10_00_256,Image|Texture_SumVariance_Vinculin_20_00_256,Image|Textur
e_SumVariance_Vinculin_20_01_256,Image|Texture_SumVariance_Vinculin_20_02_256,Image|Texture_SumVarianc
e_Vinculin_20_03_256,Image|Texture_SumVariance_Vinculin_3_01_256,Image|Texture_SumVariance_Vinculin_3_0
2_256,Image|Texture_SumVariance_Vinculin_3_00_256,Image|Texture_SumVariance_Vinculin_3_03_256,Image|Text
ure_SumVariance_Vinculin_5_01_256,Image|Texture_SumVariance_Vinculin_5_02_256,Image|Texture_SumVariance
_Vinculin_5_00_256,Image|Texture_SumVariance_Vinculin_5_03_256,Image|Texture_SumVariance_Vinculin_10_01
_256,Image|Texture_SumVariance_Vinculin_10_03_256,Image|Texture_SumVariance_Vinculin_10_02_256,Image|Te
xture_SumVariance_Vinculin_10_00_256,Image|Texture_InverseDifferenceMoment_DNA_3_01_256,Image|Texture_I
nverseDifferenceMoment_DNA_3_03_256,Image|Texture_InverseDifferenceMoment_DNA_3_02_256,Image|Texture
_InverseDifferenceMoment_DNA_3_00_256,Image|Texture_InverseDifferenceMoment_DNA_10_03_256,Image|Text
ure_InverseDifferenceMoment_DNA_10_01_256,Image|Texture_InverseDifferenceMoment_DNA_10_00_256,Image|
Texture_InverseDifferenceMoment_DNA_10_02_256,Image|Texture_InverseDifferenceMoment_DNA_20_02_256,Im
age|Texture_InverseDifferenceMoment_DNA_20_00_256,Image|Texture_InverseDifferenceMoment_DNA_20_03_256
,Image|Texture_InverseDifferenceMoment_DNA_20_01_256,Image|Texture_InverseDifferenceMoment_DNA_5_01_2
56,Image|Texture_InverseDifferenceMoment_DNA_5_02_256,Image|Texture_InverseDifferenceMoment_DNA_5_03_
256,Image|Texture_InverseDifferenceMoment_DNA_5_00_256,Image|Texture_InverseDifferenceMoment_Vinculin_1
0_00_256,Image|Texture_InverseDifferenceMoment_Vinculin_10_02_256,Image|Texture_InverseDifferenceMoment_
Vinculin_10_01_256,Image|Texture_InverseDifferenceMoment_Vinculin_10_03_256,Image|Texture_InverseDifferenc
eMoment_Vinculin_3_03_256,Image|Texture_InverseDifferenceMoment_Vinculin_3_01_256,Image|Texture_InverseD
ifferenceMoment_Vinculin_3_02_256,Image|Texture_InverseDifferenceMoment_Vinculin_3_00_256,Image|Texture_I
nverseDifferenceMoment_Vinculin_5_00_256,Image|Texture_InverseDifferenceMoment_Vinculin_5_02_256,Image|T
exture_InverseDifferenceMoment_Vinculin_5_01_256,Image|Texture_InverseDifferenceMoment_Vinculin_5_03_256,
Image|Texture_InverseDifferenceMoment_Vinculin_20_02_256,Image|Texture_InverseDifferenceMoment_Vinculin_2
0_01_256,Image|Texture_InverseDifferenceMoment_Vinculin_20_00_256,Image|Texture_InverseDifferenceMoment_
Vinculin_20_03_256,Image|Texture_InverseDifferenceMoment_Phalloidin_3_00_256,Image|Texture_InverseDifferenc
eMoment_Phalloidin_3_02_256,Image|Texture_InverseDifferenceMoment_Phalloidin_3_03_256,Image|Texture_Invers
eDifferenceMoment_Phalloidin_3_01_256,Image|Texture_InverseDifferenceMoment_Phalloidin_20_00_256,Image|Te
xture_InverseDifferenceMoment_Phalloidin_20_02_256,Image|Texture_InverseDifferenceMoment_Phalloidin_20_01_
256,Image|Texture_InverseDifferenceMoment_Phalloidin_20_03_256,Image|Texture_InverseDifferenceMoment_Phall
oidin_5_03_256,Image|Texture_InverseDifferenceMoment_Phalloidin_5_00_256,Image|Texture_InverseDifferenceMo
ment_Phalloidin_5_02_256,Image|Texture_InverseDifferenceMoment_Phalloidin_5_01_256,Image|Texture_InverseDif
ferenceMoment_Phalloidin_10_02_256,Image|Texture_InverseDifferenceMoment_Phalloidin_10_03_256,Image|Textu
re_InverseDifferenceMoment_Phalloidin_10_01_256,Image|Texture_InverseDifferenceMoment_Phalloidin_10_00_256
,Image|Texture_InverseDifferenceMoment_WGA_10_03_256,Image|Texture_InverseDifferenceMoment_WGA_10_01
_256,Image|Texture_InverseDifferenceMoment_WGA_10_02_256,Image|Texture_InverseDifferenceMoment_WGA_1
0_00_256,Image|Texture_InverseDifferenceMoment_WGA_20_02_256,Image|Texture_InverseDifferenceMoment_W
GA_20_00_256,Image|Texture_InverseDifferenceMoment_WGA_20_03_256,Image|Texture_InverseDifferenceMome
nt_WGA_20_01_256,Image|Texture_InverseDifferenceMoment_WGA_5_02_256,Image|Texture_InverseDifferenceM
oment_WGA_5_01_256,Image|Texture_InverseDifferenceMoment_WGA_5_03_256,Image|Texture_InverseDifference
Moment_WGA_5_00_256,Image|Texture_InverseDifferenceMoment_WGA_3_00_256,Image|Texture_InverseDifferen
ceMoment_WGA_3_03_256,Image|Texture_InverseDifferenceMoment_WGA_3_01_256,Image|Texture_InverseDiffer
enceMoment_WGA_3_02_256,Image|Texture_DifferenceEntropy_WGA_10_02_256,Image|Texture_DifferenceEntrop
y_WGA_10_03_256,Image|Texture_DifferenceEntropy_WGA_10_00_256,Image|Texture_DifferenceEntropy_WGA_1
0_01_256,Image|Texture_DifferenceEntropy_WGA_20_02_256,Image|Texture_DifferenceEntropy_WGA_20_01_256,
Image|Texture_DifferenceEntropy_WGA_20_03_256,Image|Texture_DifferenceEntropy_WGA_20_00_256,Image|Tex
ture_DifferenceEntropy_WGA_3_03_256,Image|Texture_DifferenceEntropy_WGA_3_02_256,Image|Texture_Differe
nceEntropy_WGA_3_00_256,Image|Texture_DifferenceEntropy_WGA_3_01_256,Image|Texture_DifferenceEntropy_
WGA_5_02_256,Image|Texture_DifferenceEntropy_WGA_5_01_256,Image|Texture_DifferenceEntropy_WGA_5_00_
256,Image|Texture_DifferenceEntropy_WGA_5_03_256,Image|Texture_DifferenceEntropy_Vinculin_20_00_256,Imag
e|Texture_DifferenceEntropy_Vinculin_20_02_256,Image|Texture_DifferenceEntropy_Vinculin_20_03_256,Image|Tex

ture_DifferenceEntropy_Vinculin_20_01_256,Image|Texture_DifferenceEntropy_Vinculin_5_02_256,Image|Texture_DifferenceEntropy_Vinculin_5_03_256,Image|Texture_DifferenceEntropy_Vinculin_5_00_256,Image|Texture_DifferenceEntropy_Vinculin_5_01_256,Image|Texture_DifferenceEntropy_Vinculin_10_00_256,Image|Texture_DifferenceEntropy_Vinculin_10_01_256,Image|Texture_DifferenceEntropy_Vinculin_10_03_256,Image|Texture_DifferenceEntropy_Vinculin_10_02_256,Image|Texture_DifferenceEntropy_Vinculin_3_01_256,Image|Texture_DifferenceEntropy_Vinculin_3_03_256,Image|Texture_DifferenceEntropy_Vinculin_3_02_256,Image|Texture_DifferenceEntropy_Vinculin_3_00_256,Image|Texture_DifferenceEntropy_DNA_20_00_256,Image|Texture_DifferenceEntropy_DNA_20_02_256,Image|Texture_DifferenceEntropy_DNA_20_01_256,Image|Texture_DifferenceEntropy_DNA_20_03_256,Image|Texture_DifferenceEntropy_DNA_3_03_256,Image|Texture_DifferenceEntropy_DNA_3_01_256,Image|Texture_DifferenceEntropy_DNA_3_02_256,Image|Texture_DifferenceEntropy_DNA_3_00_256,Image|Texture_DifferenceEntropy_DNA_10_02_256,Image|Texture_DifferenceEntropy_DNA_10_01_256,Image|Texture_DifferenceEntropy_DNA_10_03_256,Image|Texture_DifferenceEntropy_DNA_10_00_256,Image|Texture_DifferenceEntropy_DNA_5_03_256,Image|Texture_DifferenceEntropy_DNA_5_01_256,Image|Texture_DifferenceEntropy_DNA_5_00_256,Image|Texture_DifferenceEntropy_DNA_5_02_256,Image|Texture_DifferenceEntropy_Phalloidin_10_02_256,Image|Texture_DifferenceEntropy_Phalloidin_10_03_256,Image|Texture_DifferenceEntropy_Phalloidin_10_01_256,Image|Texture_DifferenceEntropy_Phalloidin_10_00_256,Image|Texture_DifferenceEntropy_Phalloidin_3_02_256,Image|Texture_DifferenceEntropy_Phalloidin_3_00_256,Image|Texture_DifferenceEntropy_Phalloidin_3_03_256,Image|Texture_DifferenceEntropy_Phalloidin_3_01_256,Image|Texture_DifferenceEntropy_Phalloidin_20_02_256,Image|Texture_DifferenceEntropy_Phalloidin_20_00_256,Image|Texture_DifferenceEntropy_Phalloidin_20_01_256,Image|Texture_DifferenceEntropy_Phalloidin_20_03_256,Image|Texture_DifferenceEntropy_Phalloidin_5_01_256,Image|Texture_DifferenceEntropy_Phalloidin_5_00_256,Image|Texture_DifferenceEntropy_Phalloidin_5_02_256,Image|Texture_DifferenceEntropy_Phalloidin_5_03_256,Image|ImageQuality_LocalFocusScore_OrigVinculin_50,Image|ImageQuality_LocalFocusScore_OrigVinculin_10,Image|ImageQuality_LocalFocusScore_OrigVinculin_20,Image|ImageQuality_LocalFocusScore_OrigVinculin_5,Image|ImageQuality_LocalFocusScore_OrigPhalloidin_5,Image|ImageQuality_LocalFocusScore_OrigPhalloidin_50,Image|ImageQuality_LocalFocusScore_OrigPhalloidin_20,Image|ImageQuality_LocalFocusScore_OrigPhalloidin_10,Image|ImageQuality_LocalFocusScore_OrigWGA_20,Image|ImageQuality_LocalFocusScore_OrigWGA_5,Image|ImageQuality_LocalFocusScore_OrigWGA_50,Image|ImageQuality_LocalFocusScore_OrigWGA_10,Image|ImageQuality_LocalFocusScore_OrigDNA_20,Image|ImageQuality_LocalFocusScore_OrigDNA_50,Image|ImageQuality_LocalFocusScore_OrigDNA_10,Image|ImageQuality_LocalFocusScore_OrigDNA_5,Image|ImageQuality_FocusScore_OrigPhalloidin,Image|ImageQuality_FocusScore_OrigDNA,Image|ImageQuality_FocusScore_OrigWGA,Image|ImageQuality_FocusScore_OrigVinculin,Image|ImageQuality_TotalArea_OrigPhalloidin,Image|ImageQuality_TotalArea_OrigWGA,Image|ImageQuality_TotalArea_OrigDNA,Image|ImageQuality_TotalArea_OrigVinculin,Image|ImageQuality_Scaling_OrigDNA,Image|ImageQuality_Scaling_OrigVinculin,Image|ImageQuality_Scaling_OrigPhalloidin,Image|ImageQuality_Scaling_OrigWGA,Image|ImageQuality_PercentMinimal_OrigVinculin,Image|ImageQuality_PercentMinimal_OrigWGA,Image|ImageQuality_PercentMinimal_OrigPhalloidin,Image|ImageQuality_PercentMinimal_OrigDNA,Image|ImageQuality_Correlation_OrigDNA_50,Image|ImageQuality_Correlation_OrigDNA_10,Image|ImageQuality_Correlation_OrigDNA_20,Image|ImageQuality_Correlation_OrigDNA_5,Image|ImageQuality_Correlation_OrigVinculin_50,Image|ImageQuality_Correlation_OrigVinculin_10,Image|ImageQuality_Correlation_OrigVinculin_20,Image|ImageQuality_Correlation_OrigVinculin_5,Image|ImageQuality_Correlation_OrigPhalloidin_50,Image|ImageQuality_Correlation_OrigPhalloidin_10,Image|ImageQuality_Correlation_OrigPhalloidin_5,Image|ImageQuality_Correlation_OrigPhalloidin_20,Image|ImageQuality_Correlation_OrigWGA_10,Image|ImageQuality_Correlation_OrigWGA_5,Image|ImageQuality_Correlation_OrigWGA_50,Image|ImageQuality_Correlation_OrigWGA_20,Image|ImageQuality_PercentMaximal_OrigDNA,Image|ImageQuality_PercentMaximal_OrigWGA,Image|ImageQuality_PercentMaximal_OrigVinculin,Image|ImageQuality_PercentMaximal_OrigPhalloidin,Image|ImageQuality_StdIntensity_OrigPhalloidin,Image|ImageQuality_StdIntensity_OrigDNA,Image|ImageQuality_StdIntensity_OrigVinculin,Image|ImageQuality_StdIntensity_OrigWGA,Image|ImageQuality_MADIntensity_OrigDNA,Image|ImageQuality_MADIntensity_OrigPhalloidin,Image|ImageQuality_MADIntensity_OrigVinculin,Image|ImageQuality_MADIntensity_OrigWGA,Image|ImageQuality_TotalIntensity_OrigVinculin,Image|ImageQuality_TotalIntensity_OrigWGA,Image|ImageQuality_TotalIntensity_OrigPhalloidin,Image|ImageQuality_TotalIntensity_OrigDNA,Image|ImageQuality_PowerLogLogSlope_OrigDNA,Image|ImageQuality_PowerLogLogSlope_OrigPhalloidin,Image|ImageQuality_PowerLogLogSlope_OrigWGA,Image|ImageQuality_PowerLogLogSlope_OrigVinculin,Image|ImageQuality_MaxIntensity_OrigWGA,Image|ImageQuality_MaxIntensity_OrigPhalloidin,Image|ImageQuality_MaxIntensity_OrigVinculin,Image|ImageQuality_MaxIntensity_OrigDNA,Image|ImageQuality_MeanIntensity_OrigWGA,Image|ImageQuality_MeanIntensity_OrigDNA,Image|ImageQuality_MeanIntensity_OrigVinculin,Image|ImageQuality_MeanIntensity_OrigPhalloidin,Image|ImageQuality_MinIntensity_OrigDNA,Image|ImageQuality_M

inIntensity_OrigWGA,Image|ImageQuality_MinIntensity_OrigVinculin,Image|ImageQuality_MinIntensity_OrigPhalloi
din,Image|ImageQuality_MedianIntensity_OrigWGA,Image|ImageQuality_MedianIntensity_OrigPhalloidin,Image|Ima
geQuality_MedianIntensity_OrigDNA,Image|ImageQuality_MedianIntensity_OrigVinculin,Image|ImageQuality_Thres
holdOtsu_OrigVinculin_3FW,Image|ImageQuality_ThresholdOtsu_OrigDNA_2W,Image|Metadata_Row,Image|Metad
ata_Col,Image|Metadata_PositionZ,Image|Metadata_ImageResolutionY,Image|Metadata_BinningY,Image|Metadata_C
hannelID,Image|Metadata_ObjectiveMagnification,Image|Metadata_MainExcitationWavelength,Image|Metadata_Binni
ngX,Image|Metadata_Site,Image|Metadata_MainEmissionWavelength,Image|Metadata_PositionX,Image|Metadata_Abs
PositionZ,Image|Metadata_ExposureTime,Image|Metadata_ChannelName,Image|Metadata_FieldID,Image|Metadata_O
bjectiveNA,Image|Metadata_PositionY,Image|Metadata_ImageSizeY,Image|Metadata_ImageResolutionX,Image|Metad
ata_ImageSizeX,Image|Metadata_MaxIntensity,Image|Metadata_PlaneID,Image|Metadata_Plate,Image|Metadata_Well,
Image|Metadata_AbsTime,Image|Granularity_4_Phalloidin,Image|Granularity_4_Vinculin,Image|Granularity_4_DNA,I
mage|Granularity_4_WGA,Image|Granularity_12_Vinculin,Image|Granularity_12_DNA,Image|Granularity_12_WGA,I
mage|Granularity_12_Phalloidin,Image|Granularity_3_Phalloidin,Image|Granularity_3_Vinculin,Image|Granularity_3_
DNA,Image|Granularity_3_WGA,Image|Granularity_2_Phalloidin,Image|Granularity_2_DNA,Image|Granularity_2_W
GA,Image|Granularity_2_Vinculin,Image|Granularity_1_WGA,Image|Granularity_1_DNA,Image|Granularity_1_Phallo
idin,Image|Granularity_1_Vinculin,Image|Granularity_11_Phalloidin,Image|Granularity_11_DNA,Image|Granularity_1
1_Vinculin,Image|Granularity_11_WGA,Image|Granularity_5_WGA,Image|Granularity_5_Phalloidin,Image|Granularit
y_5_Vinculin,Image|Granularity_5_DNA,Image|Granularity_13_WGA,Image|Granularity_13_Vinculin,Image|Granular
ity_13_DNA,Image|Granularity_13_Phalloidin,Image|Granularity_15_DNA,Image|Granularity_15_WGA,Image|Granul
arity_15_Phalloidin,Image|Granularity_15_Vinculin,Image|Granularity_6_Vinculin,Image|Granularity_6_Phalloidin,Im
age|Granularity_6_DNA,Image|Granularity_6_WGA,Image|Granularity_10_WGA,Image|Granularity_10_Vinculin,Ima
ge|Granularity_10_DNA,Image|Granularity_10_Phalloidin,Image|Granularity_14_Vinculin,Image|Granularity_14_DN
A,Image|Granularity_14_Phalloidin,Image|Granularity_14_WGA,Image|Granularity_7_Phalloidin,Image|Granularity_7
_Vinculin,Image|Granularity_7_WGA,Image|Granularity_7_DNA,Image|Granularity_16_DNA,Image|Granularity_16_
WGA,Image|Granularity_16_Phalloidin,Image|Granularity_16_Vinculin,Image|Granularity_9_WGA,Image|Granularity
_9_Phalloidin,Image|Granularity_9_DNA,Image|Granularity_9_Vinculin,Image|Granularity_8_Phalloidin,Image|Granul
arity_8_Vinculin,Image|Granularity_8_DNA,Image|Granularity_8_WGA,Image|FileName_IllumVinculin,Image|FileNa
me_CellOutlines,Image|FileName_IllumPhalloidin,Image|FileName_OrigWGA,Image|FileName_OrigPhalloidin,Image
|FileName_OrigVinculin,Image|FileName_NucleiOutlines,Image|FileName_IllumDNA,Image|FileName_OrigDNA,Im
age|FileName_IllumWGA,Image|ModuleError_07CorrectIlluminationApply,Image|ModuleError_27OverlayOutlines,I
mage|ModuleError_22MaskImage,Image|ModuleError_29SaveImages,Image|ModuleError_25MeasureImageIntensity,I
mage|ModuleError_03MeasureImageQuality,Image|ModuleError_15MeasureObjectNeighbors,Image|ModuleError_08I
dentifyPrimaryObjects,Image|ModuleError_26OverlayOutlines,Image|ModuleError_02CorrectIlluminationApply,Image
|ModuleError_09IdentifySecondaryObjects,Image|ModuleError_16MeasureObjectNeighbors,Image|ModuleError_06Co
rrectIlluminationCalculate,Image|ModuleError_23MaskImage,Image|ModuleError_13MeasureGranularity,Image|Modul
eError_19MeasureObjectSizeShape,Image|ModuleError_10FilterObjects,Image|ModuleError_12MeasureColocalization
,Image|ModuleError_20MeasureTexture,Image|ModuleError_14MeasureObjectIntensity,Image|ModuleError_04Measur
eImageQuality,Image|ModuleError_18MeasureObjectIntensityDistribution,Image|ModuleError_24MaskImage,Image|M
oduleError_28SaveImages,Image|ModuleError_01LoadData,Image|ModuleError_11IdentifyTertiaryObjects,Image|Mod
uleError_17MeasureObjectNeighbors,Image|ModuleError_21MaskImage,Image|ExecutionTime_10FilterObjects,Image|
ExecutionTime_07CorrectIlluminationApply,Image|ExecutionTime_24MaskImage,Image|ExecutionTime_04MeasureI
mageQuality,Image|ExecutionTime_27OverlayOutlines,Image|ExecutionTime_18MeasureObjectIntensityDistribution,I
mage|ExecutionTime_26OverlayOutlines,Image|ExecutionTime_22MaskImage,Image|ExecutionTime_17MeasureObje
ctNeighbors,Image|ExecutionTime_29SaveImages,Image|ExecutionTime_25MeasureImageIntensity,Image|ExecutionTi
me_12MeasureColocalization,Image|ExecutionTime_23MaskImage,Image|ExecutionTime_11IdentifyTertiaryObjects,I
mage|ExecutionTime_08IdentifyPrimaryObjects,Image|ExecutionTime_19MeasureObjectSizeShape,Image|ExecutionTi
me_02CorrectIlluminationApply,Image|ExecutionTime_09IdentifySecondaryObjects,Image|ExecutionTime_13Measure
Granularity,Image|ExecutionTime_14MeasureObjectIntensity,Image|ExecutionTime_03MeasureImageQuality,Image|E
xecutionTime_16MeasureObjectNeighbors,Image|ExecutionTime_15MeasureObjectNeighbors,Image|ExecutionTime_
20MeasureTexture,Image|ExecutionTime_21MaskImage,Image|ExecutionTime_28SaveImages,Image|ExecutionTime_
06CorrectIlluminationCalculate,Image|ExecutionTime_01LoadData,Image|Intensity_TotalIntensity_WGA,Image|Intens
ity_TotalIntensity_WGA__BackgroundOnly,Image|Intensity_TotalIntensity_Vinculin,Image|Intensity_TotalIntensity_V
inculin__BackgroundOnly,Image|Intensity_TotalIntensity_DNA,Image|Intensity_TotalIntensity_DNA_BackgroundOnl
y,Image|Intensity_TotalIntensity_Phalloidin,Image|Intensity_TotalIntensity_Phalloidin__BackgroundOnly,Image|Intens

ity_MADIntensity_WGA,Image|Intensity_MADIntensity_WGA__BackgroundOnly,Image|Intensity_MADIntensity_D
NA,Image|Intensity_MADIntensity_DNA_BackgroundOnly,Image|Intensity_MADIntensity_Vinculin,Image|Intensity_
MADIntensity_Vinculin__BackgroundOnly,Image|Intensity_MADIntensity_Phalloidin,Image|Intensity_MADIntensity
_Phalloidin__BackgroundOnly,Image|Intensity_MeanIntensity_Vinculin,Image|Intensity_MeanIntensity_Vinculin__Ba
ckgroundOnly,Image|Intensity_MeanIntensity_DNA,Image|Intensity_MeanIntensity_DNA_BackgroundOnly,Image|Int
ensity_MeanIntensity_WGA,Image|Intensity_MeanIntensity_WGA__BackgroundOnly,Image|Intensity_MeanIntensity_
Phalloidin,Image|Intensity_MeanIntensity_Phalloidin__BackgroundOnly,Image|Intensity_MedianIntensity_Phalloidin,I
mage|Intensity_MedianIntensity_Phalloidin__BackgroundOnly,Image|Intensity_MedianIntensity_DNA,Image|Intensity
_MedianIntensity_DNA_BackgroundOnly,Image|Intensity_MedianIntensity_Vinculin,Image|Intensity_MedianIntensity
_Vinculin__BackgroundOnly,Image|Intensity_MedianIntensity_WGA,Image|Intensity_MedianIntensity_WGA__Backg
roundOnly,Image|Intensity_MinIntensity_WGA,Image|Intensity_MinIntensity_WGA__BackgroundOnly,Image|Intensit
y_MinIntensity_Vinculin,Image|Intensity_MinIntensity_Vinculin__BackgroundOnly,Image|Intensity_MinIntensity_Pha
lloidin,Image|Intensity_MinIntensity_Phalloidin__BackgroundOnly,Image|Intensity_MinIntensity_DNA,Image|Intensit
y_MinIntensity_DNA_BackgroundOnly,Image|Intensity_TotalArea_DNA,Image|Intensity_TotalArea_DNA_Backgrou
ndOnly,Image|Intensity_TotalArea_Vinculin,Image|Intensity_TotalArea_Vinculin__BackgroundOnly,Image|Intensity_
TotalArea_Phalloidin,Image|Intensity_TotalArea_Phalloidin__BackgroundOnly,Image|Intensity_TotalArea_WGA,Ima
ge|Intensity_TotalArea_WGA__BackgroundOnly,Image|Intensity_MaxIntensity_Vinculin,Image|Intensity_MaxIntensit
y_Vinculin__BackgroundOnly,Image|Intensity_MaxIntensity_DNA,Image|Intensity_MaxIntensity_DNA_Background
Only,Image|Intensity_MaxIntensity_Phalloidin,Image|Intensity_MaxIntensity_Phalloidin__BackgroundOnly,Image|Inte
nsity_MaxIntensity_WGA,Image|Intensity_MaxIntensity_WGA__BackgroundOnly,Image|Intensity_StdIntensity_Vinc
ulin,Image|Intensity_StdIntensity_Vinculin__BackgroundOnly,Image|Intensity_StdIntensity_WGA,Image|Intensity_Std
Intensity_WGA__BackgroundOnly,Image|Intensity_StdIntensity_DNA,Image|Intensity_StdIntensity_DNA_Backgroun
dOnly,Image|Intensity_StdIntensity_Phalloidin,Image|Intensity_StdIntensity_Phalloidin__BackgroundOnly,Image|Inten
sity_PercentMaximal_Phalloidin,Image|Intensity_PercentMaximal_Phalloidin__BackgroundOnly,Image|Intensity_Perc
entMaximal_DNA,Image|Intensity_PercentMaximal_DNA_BackgroundOnly,Image|Intensity_PercentMaximal_WGA,I
mage|Intensity_PercentMaximal_WGA__BackgroundOnly,Image|Intensity_PercentMaximal_Vinculin,Image|Intensity_
PercentMaximal_Vinculin__BackgroundOnly,Image|Intensity_LowerQuartileIntensity_Phalloidin,Image|Intensity_Low
erQuartileIntensity_Phalloidin__BackgroundOnly,Image|Intensity_LowerQuartileIntensity_DNA,Image|Intensity_Lowe
rQuartileIntensity_DNA_BackgroundOnly,Image|Intensity_LowerQuartileIntensity_Vinculin,Image|Intensity_LowerQu
artileIntensity_Vinculin__BackgroundOnly,Image|Intensity_LowerQuartileIntensity_WGA,Image|Intensity_LowerQuar
tileIntensity_WGA__BackgroundOnly,Image|Intensity_UpperQuartileIntensity_WGA,Image|Intensity_UpperQuartileIn
tensity_WGA__BackgroundOnly,Image|Intensity_UpperQuartileIntensity_Vinculin,Image|Intensity_UpperQuartileInte
nsity_Vinculin__BackgroundOnly,Image|Intensity_UpperQuartileIntensity_Phalloidin,Image|Intensity_UpperQuartileIn
tensity_Phalloidin__BackgroundOnly,Image|Intensity_UpperQuartileIntensity_DNA,Image|Intensity_UpperQuartileInt
ensity_DNA_BackgroundOnly,Image|PathName_OrigVinculin,Image|PathName_OrigDNA,Image|PathName_NucleiO
utlines,Image|PathName_IllumVinculin,Image|PathName_IllumWGA,Image|PathName_OrigWGA,Image|PathName_O
rigPhalloidin,Image|PathName_IllumDNA,Image|PathName_CellOutlines,Image|PathName_IllumPhalloidin,Image|Wi
dth_OrigDNA,Image|Width_IllumPhalloidin,Image|Width_IllumVinculin,Image|Width_OrigPhalloidin,Image|Width_O
rigWGA,Image|Width_OrigVinculin,Image|Width_IllumDNA,Image|Width_IllumWGA,Image|Threshold_WeightedVa
riance_NucleiIncludingEdges,Image|Threshold_WeightedVariance_CellsIncludingEdges,Image|Threshold_FinalThresh
old_CellsIncludingEdges,Image|Threshold_FinalThreshold_NucleiIncludingEdges,Image|Threshold_SumOfEntropies_
NucleiIncludingEdges,Image|Threshold_SumOfEntropies_CellsIncludingEdges,Image|Threshold_OrigThreshold_Nucle
iIncludingEdges,Image|Threshold_OrigThreshold_CellsIncludingEdges,Image|URL_IllumDNA,Image|URL_OrigDNA,
Image|URL_OrigWGA,Image|URL_IllumPhalloidin,Image|URL_IllumVinculin,Image|URL_IllumWGA,Image|URL_
OrigPhalloidin,Image|URL_OrigVinculin,Image|MD5Digest_OrigWGA,Image|MD5Digest_OrigDNA,Image|MD5Dige
st_IllumDNA,Image|MD5Digest_OrigVinculin,Image|MD5Digest_OrigPhalloidin,Image|MD5Digest_IllumVinculin,I
mage|MD5Digest_IllumPhalloidin,Image|MD5Digest_IllumWGA,Image|Height_IllumDNA,Image|Height_OrigDNA,I
mage|Height_OrigPhalloidin,Image|Height_IllumVinculin,Image|Height_OrigWGA,Image|Height_IllumPhalloidin,Ima
ge|Height_IllumWGA,Image|Height_OrigVinculin,Image|Scaling_IllumDNA,Image|Scaling_IllumVinculin,Image|Scali
ng_OrigDNA,Image|Scaling_OrigWGA,Image|Scaling_IllumWGA,Image|Scaling_OrigPhalloidin,Image|Scaling_Illum
Phalloidin,Image|Scaling_OrigVinculin,Image|Count_CellsIncludingEdges,Image|Count_NucleiIncludingEdges,Image|
Count_Cytoplasm,Image|Count_Nuclei,Image|Group_Index,Image|Group_Number,Nuclei|Intensity_MinIntensityEdge_
Vinculin,Nuclei|Intensity_MinIntensityEdge_WGA,Nuclei|Intensity_MinIntensityEdge_DNA,Nuclei|Intensity_MinInte
nsityEdge_Phalloidin,Nuclei|Intensity_StdIntensityEdge_Phalloidin,Nuclei|Intensity_StdIntensityEdge_WGA,Nuclei|In

tensity_StdIntensityEdge_Vinculin,Nuclei|Intensity_StdIntensityEdge_DNA,Nuclei|Intensity_MaxIntensity_Phalloidin,Nuclei|Intensity_MaxIntensity_Vinculin,Nuclei|Intensity_MaxIntensity_DNA,Nuclei|Intensity_MaxIntensity_WGA,Nuclei|Intensity_LowerQuartileIntensity_WGA,Nuclei|Intensity_LowerQuartileIntensity_Vinculin,Nuclei|Intensity_LowerQuartileIntensity_DNA,Nuclei|Intensity_LowerQuartileIntensity_Phalloidin,Nuclei|Intensity_MassDisplacement_DNA,Nuclei|Intensity_MassDisplacement_Phalloidin,Nuclei|Intensity_MassDisplacement_Vinculin,Nuclei|Intensity_MassDisplacement_WGA,Nuclei|Intensity_MeanIntensity_WGA,Nuclei|Intensity_MeanIntensity_DNA,Nuclei|Intensity_MeanIntensity_Phalloidin,Nuclei|Intensity_MeanIntensity_Vinculin,Nuclei|Intensity_UpperQuartileIntensity_DNA,Nuclei|Intensity_UpperQuartileIntensity_Vinculin,Nuclei|Intensity_UpperQuartileIntensity_WGA,Nuclei|Intensity_UpperQuartileIntensity_Phalloidin,Nuclei|Intensity_StdIntensity_Vinculin,Nuclei|Intensity_StdIntensity_WGA,Nuclei|Intensity_StdIntensity_DNA,Nuclei|Intensity_StdIntensity_Phalloidin,Nuclei|Intensity_IntegratedIntensity_WGA,Nuclei|Intensity_IntegratedIntensity_DNA,Nuclei|Intensity_IntegratedIntensity_Vinculin,Nuclei|Intensity_IntegratedIntensity_Phalloidin,Nuclei|Intensity_MeanIntensityEdge_Vinculin,Nuclei|Intensity_MeanIntensityEdge_Phalloidin,Nuclei|Intensity_MeanIntensityEdge_DNA,Nuclei|Intensity_MeanIntensityEdge_WGA,Nuclei|Intensity_MedianIntensity_DNA,Nuclei|Intensity_MedianIntensity_Phalloidin,Nuclei|Intensity_MedianIntensity_Vinculin,Nuclei|Intensity_MedianIntensity_WGA,Nuclei|Intensity_MADIntensity_Vinculin,Nuclei|Intensity_MADIntensity_WGA,Nuclei|Intensity_MADIntensity_DNA,Nuclei|Intensity_MADIntensity_Phalloidin,Nuclei|Intensity_IntegratedIntensityEdge_Phalloidin,Nuclei|Intensity_IntegratedIntensityEdge_Vinculin,Nuclei|Intensity_IntegratedIntensityEdge_WGA,Nuclei|Intensity_IntegratedIntensityEdge_DNA,Nuclei|Intensity_MaxIntensityEdge_WGA,Nuclei|Intensity_MaxIntensityEdge_Vinculin,Nuclei|Intensity_MaxIntensityEdge_DNA,Nuclei|Intensity_MaxIntensityEdge_Phalloidin,Nuclei|Intensity_MinIntensity_Phalloidin,Nuclei|Intensity_MinIntensity_Vinculin,Nuclei|Intensity_MinIntensity_DNA,Nuclei|Intensity_MinIntensity_WGA,Nuclei|Texture_Entropy_DNA_20_03_256,Nuclei|Texture_Entropy_DNA_20_00_256,Nuclei|Texture_Entropy_DNA_20_02_256,Nuclei|Texture_Entropy_DNA_20_01_256,Nuclei|Texture_Entropy_DNA_5_01_256,Nuclei|Texture_Entropy_DNA_5_03_256,Nuclei|Texture_Entropy_DNA_5_00_256,Nuclei|Texture_Entropy_DNA_5_02_256,Nuclei|Texture_Entropy_DNA_3_02_256,Nuclei|Texture_Entropy_DNA_3_01_256,Nuclei|Texture_Entropy_DNA_3_03_256,Nuclei|Texture_Entropy_DNA_3_00_256,Nuclei|Texture_Entropy_DNA_10_03_256,Nuclei|Texture_Entropy_DNA_10_00_256,Nuclei|Texture_Entropy_DNA_10_02_256,Nuclei|Texture_Entropy_DNA_10_01_256,Nuclei|Texture_Entropy_Phalloidin_3_02_256,Nuclei|Texture_Entropy_Phalloidin_3_00_256,Nuclei|Texture_Entropy_Phalloidin_3_01_256,Nuclei|Texture_Entropy_Phalloidin_3_03_256,Nuclei|Texture_Entropy_Phalloidin_10_03_256,Nuclei|Texture_Entropy_Phalloidin_10_02_256,Nuclei|Texture_Entropy_Phalloidin_10_00_256,Nuclei|Texture_Entropy_Phalloidin_10_01_256,Nuclei|Texture_Entropy_Phalloidin_20_03_256,Nuclei|Texture_Entropy_Phalloidin_20_01_256,Nuclei|Texture_Entropy_Phalloidin_20_02_256,Nuclei|Texture_Entropy_Phalloidin_20_00_256,Nuclei|Texture_Entropy_Phalloidin_5_01_256,Nuclei|Texture_Entropy_Phalloidin_5_03_256,Nuclei|Texture_Entropy_Phalloidin_5_00_256,Nuclei|Texture_Entropy_Phalloidin_5_02_256,Nuclei|Texture_Entropy_WGA_3_01_256,Nuclei|Texture_Entropy_WGA_3_00_256,Nuclei|Texture_Entropy_WGA_3_03_256,Nuclei|Texture_Entropy_WGA_3_02_256,Nuclei|Texture_Entropy_WGA_10_01_256,Nuclei|Texture_Entropy_WGA_10_00_256,Nuclei|Texture_Entropy_WGA_10_02_256,Nuclei|Texture_Entropy_WGA_10_03_256,Nuclei|Texture_Entropy_WGA_5_01_256,Nuclei|Texture_Entropy_WGA_5_02_256,Nuclei|Texture_Entropy_WGA_5_00_256,Nuclei|Texture_Entropy_WGA_5_03_256,Nuclei|Texture_Entropy_WGA_20_03_256,Nuclei|Texture_Entropy_WGA_20_00_256,Nuclei|Texture_Entropy_WGA_20_01_256,Nuclei|Texture_Entropy_WGA_20_02_256,Nuclei|Texture_Entropy_Vinculin_3_03_256,Nuclei|Texture_Entropy_Vinculin_3_00_256,Nuclei|Texture_Entropy_Vinculin_3_01_256,Nuclei|Texture_Entropy_Vinculin_3_02_256,Nuclei|Texture_Entropy_Vinculin_20_03_256,Nuclei|Texture_Entropy_Vinculin_20_00_256,Nuclei|Texture_Entropy_Vinculin_20_02_256,Nuclei|Texture_Entropy_Vinculin_20_01_256,Nuclei|Texture_Entropy_Vinculin_10_00_256,Nuclei|Texture_Entropy_Vinculin_10_02_256,Nuclei|Texture_Entropy_Vinculin_10_03_256,Nuclei|Texture_Entropy_Vinculin_10_01_256,Nuclei|Texture_Entropy_Vinculin_5_01_256,Nuclei|Texture_Entropy_Vinculin_5_02_256,Nuclei|Texture_Entropy_Vinculin_5_00_256,Nuclei|Texture_Entropy_Vinculin_5_03_256,Nuclei|Texture_DifferenceEntropy_Vinculin_20_00_256,Nuclei|Texture_DifferenceEntropy_Vinculin_20_02_256,Nuclei|Texture_DifferenceEntropy_Vinculin_20_01_256,Nuclei|Texture_DifferenceEntropy_Vinculin_20_03_256,Nuclei|Texture_DifferenceEntropy_Vinculin_3_01_256,Nuclei|Texture_DifferenceEntropy_Vinculin_3_03_256,Nuclei|Texture_DifferenceEntropy_Vinculin_3_00_256,Nuclei|Texture_DifferenceEntropy_Vinculin_3_02_256,Nuclei|Texture_DifferenceEntropy_Vinculin_5_01_256,Nuclei|Texture_DifferenceEntropy_Vinculin_5_02_256,Nuclei|Texture_DifferenceEntropy_Vinculin_5_00_256,Nuclei|Texture_DifferenceEntropy_Vinculin_5_03_256,Nuclei|Texture_DifferenceEntropy_Vinculin_10_01_256,Nuclei|Texture_DifferenceEntropy_Vinculin_10_03_256,Nuclei|Texture_DifferenceEntropy_Vinculin_10_00_256,Nuclei|Texture_DifferenceEntropy_Vinculin_10_02_256,Nuclei|Texture_DifferenceEntropy_Phalloidin_5_03_256,Nuclei|Texture_DifferenceEntropy_Phalloidin_5_00_256,Nuclei|Texture_DifferenceEntropy_Phalloidin_5_01_256,Nuclei|Texture_DifferenceEntropy_Phalloidin_5_02_256,Nuclei|Texture_DifferenceEntropy_

Phalloidin_20_02_256,Nuclei|Texture_DifferenceEntropy_Phalloidin_20_00_256,Nuclei|Texture_DifferenceEntropy_P
halloidin_20_03_256,Nuclei|Texture_DifferenceEntropy_Phalloidin_20_01_256,Nuclei|Texture_DifferenceEntropy_Ph
alloidin_3_01_256,Nuclei|Texture_DifferenceEntropy_Phalloidin_3_03_256,Nuclei|Texture_DifferenceEntropy_Phallo
idin_3_00_256,Nuclei|Texture_DifferenceEntropy_Phalloidin_3_02_256,Nuclei|Texture_DifferenceEntropy_Phalloidin
_10_03_256,Nuclei|Texture_DifferenceEntropy_Phalloidin_10_02_256,Nuclei|Texture_DifferenceEntropy_Phalloidin_
10_00_256,Nuclei|Texture_DifferenceEntropy_Phalloidin_10_01_256,Nuclei|Texture_DifferenceEntropy_WGA_3_03
_256,Nuclei|Texture_DifferenceEntropy_WGA_3_02_256,Nuclei|Texture_DifferenceEntropy_WGA_3_00_256,Nuclei
|Texture_DifferenceEntropy_WGA_3_01_256,Nuclei|Texture_DifferenceEntropy_WGA_10_00_256,Nuclei|Texture_D
ifferenceEntropy_WGA_10_03_256,Nuclei|Texture_DifferenceEntropy_WGA_10_01_256,Nuclei|Texture_Difference
Entropy_WGA_10_02_256,Nuclei|Texture_DifferenceEntropy_WGA_20_01_256,Nuclei|Texture_DifferenceEntropy_
WGA_20_00_256,Nuclei|Texture_DifferenceEntropy_WGA_20_02_256,Nuclei|Texture_DifferenceEntropy_WGA_20
_03_256,Nuclei|Texture_DifferenceEntropy_WGA_5_00_256,Nuclei|Texture_DifferenceEntropy_WGA_5_03_256,Nu
clei|Texture_DifferenceEntropy_WGA_5_02_256,Nuclei|Texture_DifferenceEntropy_WGA_5_01_256,Nuclei|Texture
_DifferenceEntropy_DNA_3_01_256,Nuclei|Texture_DifferenceEntropy_DNA_3_03_256,Nuclei|Texture_DifferenceE
ntropy_DNA_3_00_256,Nuclei|Texture_DifferenceEntropy_DNA_3_02_256,Nuclei|Texture_DifferenceEntropy_DNA
_20_01_256,Nuclei|Texture_DifferenceEntropy_DNA_20_02_256,Nuclei|Texture_DifferenceEntropy_DNA_20_03_25
6,Nuclei|Texture_DifferenceEntropy_DNA_20_00_256,Nuclei|Texture_DifferenceEntropy_DNA_10_01_256,Nuclei|T
exture_DifferenceEntropy_DNA_10_00_256,Nuclei|Texture_DifferenceEntropy_DNA_10_02_256,Nuclei|Texture_Dif
ferenceEntropy_DNA_10_03_256,Nuclei|Texture_DifferenceEntropy_DNA_5_01_256,Nuclei|Texture_DifferenceEntr
opy_DNA_5_00_256,Nuclei|Texture_DifferenceEntropy_DNA_5_02_256,Nuclei|Texture_DifferenceEntropy_DNA_5
_03_256,Nuclei|Texture_Correlation_WGA_20_00_256,Nuclei|Texture_Correlation_WGA_20_01_256,Nuclei|Texture
_Correlation_WGA_20_03_256,Nuclei|Texture_Correlation_WGA_20_02_256,Nuclei|Texture_Correlation_WGA_10_
01_256,Nuclei|Texture_Correlation_WGA_10_00_256,Nuclei|Texture_Correlation_WGA_10_02_256,Nuclei|Texture_
Correlation_WGA_10_03_256,Nuclei|Texture_Correlation_WGA_3_01_256,Nuclei|Texture_Correlation_WGA_3_00_
256,Nuclei|Texture_Correlation_WGA_3_02_256,Nuclei|Texture_Correlation_WGA_3_03_256,Nuclei|Texture_Correl
ation_WGA_5_00_256,Nuclei|Texture_Correlation_WGA_5_03_256,Nuclei|Texture_Correlation_WGA_5_01_256,Nu
clei|Texture_Correlation_WGA_5_02_256,Nuclei|Texture_Correlation_DNA_10_01_256,Nuclei|Texture_Correlation_
DNA_10_03_256,Nuclei|Texture_Correlation_DNA_10_00_256,Nuclei|Texture_Correlation_DNA_10_02_256,Nuclei|
Texture_Correlation_DNA_3_03_256,Nuclei|Texture_Correlation_DNA_3_00_256,Nuclei|Texture_Correlation_DNA_
3_01_256,Nuclei|Texture_Correlation_DNA_3_02_256,Nuclei|Texture_Correlation_DNA_5_00_256,Nuclei|Texture_C
orrelation_DNA_5_02_256,Nuclei|Texture_Correlation_DNA_5_01_256,Nuclei|Texture_Correlation_DNA_5_03_256,
Nuclei|Texture_Correlation_DNA_20_03_256,Nuclei|Texture_Correlation_DNA_20_02_256,Nuclei|Texture_Correlati
on_DNA_20_00_256,Nuclei|Texture_Correlation_DNA_20_01_256,Nuclei|Texture_Correlation_Phalloidin_10_00_25
6,Nuclei|Texture_Correlation_Phalloidin_10_03_256,Nuclei|Texture_Correlation_Phalloidin_10_02_256,Nuclei|Textur
e_Correlation_Phalloidin_10_01_256,Nuclei|Texture_Correlation_Phalloidin_20_03_256,Nuclei|Texture_Correlation_
Phalloidin_20_01_256,Nuclei|Texture_Correlation_Phalloidin_20_00_256,Nuclei|Texture_Correlation_Phalloidin_20_
02_256,Nuclei|Texture_Correlation_Phalloidin_5_00_256,Nuclei|Texture_Correlation_Phalloidin_5_03_256,Nuclei|Te
xture_Correlation_Phalloidin_5_01_256,Nuclei|Texture_Correlation_Phalloidin_5_02_256,Nuclei|Texture_Correlation
_Phalloidin_3_01_256,Nuclei|Texture_Correlation_Phalloidin_3_02_256,Nuclei|Texture_Correlation_Phalloidin_3_03
_256,Nuclei|Texture_Correlation_Phalloidin_3_00_256,Nuclei|Texture_Correlation_Vinculin_20_03_256,Nuclei|Textu
re_Correlation_Vinculin_20_02_256,Nuclei|Texture_Correlation_Vinculin_20_01_256,Nuclei|Texture_Correlation_Vi
nculin_20_00_256,Nuclei|Texture_Correlation_Vinculin_10_03_256,Nuclei|Texture_Correlation_Vinculin_10_02_256,
Nuclei|Texture_Correlation_Vinculin_10_00_256,Nuclei|Texture_Correlation_Vinculin_10_01_256,Nuclei|Texture_Co
rrelation_Vinculin_3_00_256,Nuclei|Texture_Correlation_Vinculin_3_03_256,Nuclei|Texture_Correlation_Vinculin_3
_01_256,Nuclei|Texture_Correlation_Vinculin_3_02_256,Nuclei|Texture_Correlation_Vinculin_5_01_256,Nuclei|Text
ure_Correlation_Vinculin_5_02_256,Nuclei|Texture_Correlation_Vinculin_5_00_256,Nuclei|Texture_Correlation_Vin
culin_5_03_256,Nuclei|Texture_Variance_DNA_10_02_256,Nuclei|Texture_Variance_DNA_10_00_256,Nuclei|Textur
e_Variance_DNA_10_03_256,Nuclei|Texture_Variance_DNA_10_01_256,Nuclei|Texture_Variance_DNA_3_02_256,
Nuclei|Texture_Variance_DNA_3_00_256,Nuclei|Texture_Variance_DNA_3_01_256,Nuclei|Texture_Variance_DNA_
3_03_256,Nuclei|Texture_Variance_DNA_20_01_256,Nuclei|Texture_Variance_DNA_20_00_256,Nuclei|Texture_Var
iance_DNA_20_03_256,Nuclei|Texture_Variance_DNA_20_02_256,Nuclei|Texture_Variance_DNA_5_00_256,Nuclei
|Texture_Variance_DNA_5_01_256,Nuclei|Texture_Variance_DNA_5_03_256,Nuclei|Texture_Variance_DNA_5_02_
256,Nuclei|Texture_Variance_WGA_20_03_256,Nuclei|Texture_Variance_WGA_20_02_256,Nuclei|Texture_Variance
_WGA_20_00_256,Nuclei|Texture_Variance_WGA_20_01_256,Nuclei|Texture_Variance_WGA_5_00_256,Nuclei|Te

xture_Variance_WGA_5_01_256,Nuclei|Texture_Variance_WGA_5_02_256,Nuclei|Texture_Variance_WGA_5_03_2
56,Nuclei|Texture_Variance_WGA_3_02_256,Nuclei|Texture_Variance_WGA_3_03_256,Nuclei|Texture_Variance_W
GA_3_01_256,Nuclei|Texture_Variance_WGA_3_00_256,Nuclei|Texture_Variance_WGA_10_00_256,Nuclei|Texture
_Variance_WGA_10_02_256,Nuclei|Texture_Variance_WGA_10_03_256,Nuclei|Texture_Variance_WGA_10_01_25
6,Nuclei|Texture_Variance_Phalloidin_5_01_256,Nuclei|Texture_Variance_Phalloidin_5_03_256,Nuclei|Texture_Vari
ance_Phalloidin_5_00_256,Nuclei|Texture_Variance_Phalloidin_5_02_256,Nuclei|Texture_Variance_Phalloidin_20_0
2_256,Nuclei|Texture_Variance_Phalloidin_20_03_256,Nuclei|Texture_Variance_Phalloidin_20_00_256,Nuclei|Textur
e_Variance_Phalloidin_20_01_256,Nuclei|Texture_Variance_Phalloidin_10_00_256,Nuclei|Texture_Variance_Phalloid
in_10_01_256,Nuclei|Texture_Variance_Phalloidin_10_02_256,Nuclei|Texture_Variance_Phalloidin_10_03_256,Nucle
i|Texture_Variance_Phalloidin_3_00_256,Nuclei|Texture_Variance_Phalloidin_3_01_256,Nuclei|Texture_Variance_Ph
alloidin_3_03_256,Nuclei|Texture_Variance_Phalloidin_3_02_256,Nuclei|Texture_Variance_Vinculin_3_02_256,Nucl
ei|Texture_Variance_Vinculin_3_01_256,Nuclei|Texture_Variance_Vinculin_3_03_256,Nuclei|Texture_Variance_Vinc
ulin_3_00_256,Nuclei|Texture_Variance_Vinculin_10_00_256,Nuclei|Texture_Variance_Vinculin_10_03_256,Nuclei|
Texture_Variance_Vinculin_10_01_256,Nuclei|Texture_Variance_Vinculin_10_02_256,Nuclei|Texture_Variance_Vin
culin_5_01_256,Nuclei|Texture_Variance_Vinculin_5_00_256,Nuclei|Texture_Variance_Vinculin_5_03_256,Nuclei|T
exture_Variance_Vinculin_5_02_256,Nuclei|Texture_Variance_Vinculin_20_00_256,Nuclei|Texture_Variance_Vinculi
n_20_01_256,Nuclei|Texture_Variance_Vinculin_20_03_256,Nuclei|Texture_Variance_Vinculin_20_02_256,Nuclei|T
exture_InfoMeas2_Vinculin_5_02_256,Nuclei|Texture_InfoMeas2_Vinculin_5_00_256,Nuclei|Texture_InfoMeas2_Vi
nculin_5_03_256,Nuclei|Texture_InfoMeas2_Vinculin_5_01_256,Nuclei|Texture_InfoMeas2_Vinculin_20_01_256,Nu
clei|Texture_InfoMeas2_Vinculin_20_00_256,Nuclei|Texture_InfoMeas2_Vinculin_20_03_256,Nuclei|Texture_InfoM
eas2_Vinculin_20_02_256,Nuclei|Texture_InfoMeas2_Vinculin_10_03_256,Nuclei|Texture_InfoMeas2_Vinculin_10_
01_256,Nuclei|Texture_InfoMeas2_Vinculin_10_00_256,Nuclei|Texture_InfoMeas2_Vinculin_10_02_256,Nuclei|Text
ure_InfoMeas2_Vinculin_3_02_256,Nuclei|Texture_InfoMeas2_Vinculin_3_00_256,Nuclei|Texture_InfoMeas2_Vincu
lin_3_03_256,Nuclei|Texture_InfoMeas2_Vinculin_3_01_256,Nuclei|Texture_InfoMeas2_WGA_5_00_256,Nuclei|Tex
ture_InfoMeas2_WGA_5_03_256,Nuclei|Texture_InfoMeas2_WGA_5_01_256,Nuclei|Texture_InfoMeas2_WGA_5_0
2_256,Nuclei|Texture_InfoMeas2_WGA_10_02_256,Nuclei|Texture_InfoMeas2_WGA_10_00_256,Nuclei|Texture_Inf
oMeas2_WGA_10_03_256,Nuclei|Texture_InfoMeas2_WGA_10_01_256,Nuclei|Texture_InfoMeas2_WGA_20_02_2
56,Nuclei|Texture_InfoMeas2_WGA_20_01_256,Nuclei|Texture_InfoMeas2_WGA_20_00_256,Nuclei|Texture_InfoM
eas2_WGA_20_03_256,Nuclei|Texture_InfoMeas2_WGA_3_00_256,Nuclei|Texture_InfoMeas2_WGA_3_01_256,Nu
clei|Texture_InfoMeas2_WGA_3_03_256,Nuclei|Texture_InfoMeas2_WGA_3_02_256,Nuclei|Texture_InfoMeas2_D
NA_3_00_256,Nuclei|Texture_InfoMeas2_DNA_3_03_256,Nuclei|Texture_InfoMeas2_DNA_3_01_256,Nuclei|Textur
e_InfoMeas2_DNA_3_02_256,Nuclei|Texture_InfoMeas2_DNA_10_01_256,Nuclei|Texture_InfoMeas2_DNA_10_03
_256,Nuclei|Texture_InfoMeas2_DNA_10_00_256,Nuclei|Texture_InfoMeas2_DNA_10_02_256,Nuclei|Texture_Info
Meas2_DNA_5_01_256,Nuclei|Texture_InfoMeas2_DNA_5_00_256,Nuclei|Texture_InfoMeas2_DNA_5_02_256,Nuc
lei|Texture_InfoMeas2_DNA_5_03_256,Nuclei|Texture_InfoMeas2_DNA_20_03_256,Nuclei|Texture_InfoMeas2_DN
A_20_02_256,Nuclei|Texture_InfoMeas2_DNA_20_01_256,Nuclei|Texture_InfoMeas2_DNA_20_00_256,Nuclei|Text
ure_InfoMeas2_Phalloidin_20_01_256,Nuclei|Texture_InfoMeas2_Phalloidin_20_02_256,Nuclei|Texture_InfoMeas2_
Phalloidin_20_03_256,Nuclei|Texture_InfoMeas2_Phalloidin_20_00_256,Nuclei|Texture_InfoMeas2_Phalloidin_5_01
_256,Nuclei|Texture_InfoMeas2_Phalloidin_5_02_256,Nuclei|Texture_InfoMeas2_Phalloidin_5_03_256,Nuclei|Textur
e_InfoMeas2_Phalloidin_5_00_256,Nuclei|Texture_InfoMeas2_Phalloidin_10_00_256,Nuclei|Texture_InfoMeas2_Pha
lloidin_10_02_256,Nuclei|Texture_InfoMeas2_Phalloidin_10_03_256,Nuclei|Texture_InfoMeas2_Phalloidin_10_01_2
56,Nuclei|Texture_InfoMeas2_Phalloidin_3_02_256,Nuclei|Texture_InfoMeas2_Phalloidin_3_03_256,Nuclei|Texture_
InfoMeas2_Phalloidin_3_00_256,Nuclei|Texture_InfoMeas2_Phalloidin_3_01_256,Nuclei|Texture_SumVariance_WG
A_20_00_256,Nuclei|Texture_SumVariance_WGA_20_02_256,Nuclei|Texture_SumVariance_WGA_20_03_256,Nucl
ei|Texture_SumVariance_WGA_20_01_256,Nuclei|Texture_SumVariance_WGA_10_00_256,Nuclei|Texture_SumVari
ance_WGA_10_01_256,Nuclei|Texture_SumVariance_WGA_10_02_256,Nuclei|Texture_SumVariance_WGA_10_03
_256,Nuclei|Texture_SumVariance_WGA_3_00_256,Nuclei|Texture_SumVariance_WGA_3_02_256,Nuclei|Texture_
SumVariance_WGA_3_03_256,Nuclei|Texture_SumVariance_WGA_3_01_256,Nuclei|Texture_SumVariance_WGA_
5_03_256,Nuclei|Texture_SumVariance_WGA_5_01_256,Nuclei|Texture_SumVariance_WGA_5_02_256,Nuclei|Text
ure_SumVariance_WGA_5_00_256,Nuclei|Texture_SumVariance_Vinculin_5_00_256,Nuclei|Texture_SumVariance_
Vinculin_5_02_256,Nuclei|Texture_SumVariance_Vinculin_5_03_256,Nuclei|Texture_SumVariance_Vinculin_5_01_
256,Nuclei|Texture_SumVariance_Vinculin_10_03_256,Nuclei|Texture_SumVariance_Vinculin_10_01_256,Nuclei|Te
xture_SumVariance_Vinculin_10_00_256,Nuclei|Texture_SumVariance_Vinculin_10_02_256,Nuclei|Texture_SumVar
iance_Vinculin_3_02_256,Nuclei|Texture_SumVariance_Vinculin_3_00_256,Nuclei|Texture_SumVariance_Vinculin_

3_03_256,Nuclei|Texture_SumVariance_Vinculin_3_01_256,Nuclei|Texture_SumVariance_Vinculin_20_02_256,Nucl
ei|Texture_SumVariance_Vinculin_20_03_256,Nuclei|Texture_SumVariance_Vinculin_20_01_256,Nuclei|Texture_Su
mVariance_Vinculin_20_00_256,Nuclei|Texture_SumVariance_Phalloidin_3_01_256,Nuclei|Texture_SumVariance_P
halloidin_3_02_256,Nuclei|Texture_SumVariance_Phalloidin_3_03_256,Nuclei|Texture_SumVariance_Phalloidin_3_0
0_256,Nuclei|Texture_SumVariance_Phalloidin_5_00_256,Nuclei|Texture_SumVariance_Phalloidin_5_02_256,Nuclei|
Texture_SumVariance_Phalloidin_5_03_256,Nuclei|Texture_SumVariance_Phalloidin_5_01_256,Nuclei|Texture_Sum
Variance_Phalloidin_20_02_256,Nuclei|Texture_SumVariance_Phalloidin_20_03_256,Nuclei|Texture_SumVariance_P
halloidin_20_01_256,Nuclei|Texture_SumVariance_Phalloidin_20_00_256,Nuclei|Texture_SumVariance_Phalloidin_1
0_03_256,Nuclei|Texture_SumVariance_Phalloidin_10_02_256,Nuclei|Texture_SumVariance_Phalloidin_10_01_256,
Nuclei|Texture_SumVariance_Phalloidin_10_00_256,Nuclei|Texture_SumVariance_DNA_10_03_256,Nuclei|Texture_
SumVariance_DNA_10_00_256,Nuclei|Texture_SumVariance_DNA_10_02_256,Nuclei|Texture_SumVariance_DNA_
10_01_256,Nuclei|Texture_SumVariance_DNA_20_03_256,Nuclei|Texture_SumVariance_DNA_20_00_256,Nuclei|T
exture_SumVariance_DNA_20_02_256,Nuclei|Texture_SumVariance_DNA_20_01_256,Nuclei|Texture_SumVariance
_DNA_5_02_256,Nuclei|Texture_SumVariance_DNA_5_01_256,Nuclei|Texture_SumVariance_DNA_5_00_256,Nucl
ei|Texture_SumVariance_DNA_5_03_256,Nuclei|Texture_SumVariance_DNA_3_02_256,Nuclei|Texture_SumVarianc
e_DNA_3_01_256,Nuclei|Texture_SumVariance_DNA_3_03_256,Nuclei|Texture_SumVariance_DNA_3_00_256,Nuc
lei|Texture_SumEntropy_Phalloidin_3_03_256,Nuclei|Texture_SumEntropy_Phalloidin_3_02_256,Nuclei|Texture_Su
mEntropy_Phalloidin_3_00_256,Nuclei|Texture_SumEntropy_Phalloidin_3_01_256,Nuclei|Texture_SumEntropy_Phal
loidin_5_02_256,Nuclei|Texture_SumEntropy_Phalloidin_5_00_256,Nuclei|Texture_SumEntropy_Phalloidin_5_03_25
6,Nuclei|Texture_SumEntropy_Phalloidin_5_01_256,Nuclei|Texture_SumEntropy_Phalloidin_10_03_256,Nuclei|Textu
re_SumEntropy_Phalloidin_10_00_256,Nuclei|Texture_SumEntropy_Phalloidin_10_01_256,Nuclei|Texture_SumEntro
py_Phalloidin_10_02_256,Nuclei|Texture_SumEntropy_Phalloidin_20_01_256,Nuclei|Texture_SumEntropy_Phalloidi
n_20_00_256,Nuclei|Texture_SumEntropy_Phalloidin_20_03_256,Nuclei|Texture_SumEntropy_Phalloidin_20_02_256
,Nuclei|Texture_SumEntropy_WGA_5_02_256,Nuclei|Texture_SumEntropy_WGA_5_03_256,Nuclei|Texture_SumEnt
ropy_WGA_5_01_256,Nuclei|Texture_SumEntropy_WGA_5_00_256,Nuclei|Texture_SumEntropy_WGA_3_00_256,
Nuclei|Texture_SumEntropy_WGA_3_02_256,Nuclei|Texture_SumEntropy_WGA_3_03_256,Nuclei|Texture_SumEnt
ropy_WGA_3_01_256,Nuclei|Texture_SumEntropy_WGA_10_00_256,Nuclei|Texture_SumEntropy_WGA_10_01_25
6,Nuclei|Texture_SumEntropy_WGA_10_02_256,Nuclei|Texture_SumEntropy_WGA_10_03_256,Nuclei|Texture_Su
mEntropy_WGA_20_01_256,Nuclei|Texture_SumEntropy_WGA_20_02_256,Nuclei|Texture_SumEntropy_WGA_20_
03_256,Nuclei|Texture_SumEntropy_WGA_20_00_256,Nuclei|Texture_SumEntropy_DNA_10_00_256,Nuclei|Textur
e_SumEntropy_DNA_10_03_256,Nuclei|Texture_SumEntropy_DNA_10_01_256,Nuclei|Texture_SumEntropy_DNA_
10_02_256,Nuclei|Texture_SumEntropy_DNA_20_01_256,Nuclei|Texture_SumEntropy_DNA_20_00_256,Nuclei|Tex
ture_SumEntropy_DNA_20_03_256,Nuclei|Texture_SumEntropy_DNA_20_02_256,Nuclei|Texture_SumEntropy_DN
A_3_01_256,Nuclei|Texture_SumEntropy_DNA_3_02_256,Nuclei|Texture_SumEntropy_DNA_3_00_256,Nuclei|Text
ure_SumEntropy_DNA_3_03_256,Nuclei|Texture_SumEntropy_DNA_5_03_256,Nuclei|Texture_SumEntropy_DNA_
5_00_256,Nuclei|Texture_SumEntropy_DNA_5_01_256,Nuclei|Texture_SumEntropy_DNA_5_02_256,Nuclei|Texture
_SumEntropy_Vinculin_3_03_256,Nuclei|Texture_SumEntropy_Vinculin_3_01_256,Nuclei|Texture_SumEntropy_Vin
culin_3_02_256,Nuclei|Texture_SumEntropy_Vinculin_3_00_256,Nuclei|Texture_SumEntropy_Vinculin_20_01_256,
Nuclei|Texture_SumEntropy_Vinculin_20_02_256,Nuclei|Texture_SumEntropy_Vinculin_20_03_256,Nuclei|Texture_
SumEntropy_Vinculin_20_00_256,Nuclei|Texture_SumEntropy_Vinculin_10_03_256,Nuclei|Texture_SumEntropy_Vi
nculin_10_01_256,Nuclei|Texture_SumEntropy_Vinculin_10_02_256,Nuclei|Texture_SumEntropy_Vinculin_10_00_2
56,Nuclei|Texture_SumEntropy_Vinculin_5_03_256,Nuclei|Texture_SumEntropy_Vinculin_5_00_256,Nuclei|Texture_
SumEntropy_Vinculin_5_01_256,Nuclei|Texture_SumEntropy_Vinculin_5_02_256,Nuclei|Texture_DifferenceVarianc
e_Phalloidin_5_02_256,Nuclei|Texture_DifferenceVariance_Phalloidin_5_01_256,Nuclei|Texture_DifferenceVariance
_Phalloidin_5_03_256,Nuclei|Texture_DifferenceVariance_Phalloidin_5_00_256,Nuclei|Texture_DifferenceVariance_
Phalloidin_20_00_256,Nuclei|Texture_DifferenceVariance_Phalloidin_20_03_256,Nuclei|Texture_DifferenceVariance
_Phalloidin_20_01_256,Nuclei|Texture_DifferenceVariance_Phalloidin_20_02_256,Nuclei|Texture_DifferenceVarianc
e_Phalloidin_3_00_256,Nuclei|Texture_DifferenceVariance_Phalloidin_3_01_256,Nuclei|Texture_DifferenceVariance
_Phalloidin_3_03_256,Nuclei|Texture_DifferenceVariance_Phalloidin_3_02_256,Nuclei|Texture_DifferenceVariance_
Phalloidin_10_00_256,Nuclei|Texture_DifferenceVariance_Phalloidin_10_02_256,Nuclei|Texture_DifferenceVariance
_Phalloidin_10_03_256,Nuclei|Texture_DifferenceVariance_Phalloidin_10_01_256,Nuclei|Texture_DifferenceVarianc
e_WGA_20_01_256,Nuclei|Texture_DifferenceVariance_WGA_20_03_256,Nuclei|Texture_DifferenceVariance_WG
A_20_00_256,Nuclei|Texture_DifferenceVariance_WGA_20_02_256,Nuclei|Texture_DifferenceVariance_WGA_10_0
0_256,Nuclei|Texture_DifferenceVariance_WGA_10_03_256,Nuclei|Texture_DifferenceVariance_WGA_10_02_256,

Nuclei|Texture_DifferenceVariance_WGA_10_01_256,Nuclei|Texture_DifferenceVariance_WGA_3_01_256,Nuclei|Texture_DifferenceVariance_WGA_3_00_256,Nuclei|Texture_DifferenceVariance_WGA_3_03_256,Nuclei|Texture_DifferenceVariance_WGA_3_02_256,Nuclei|Texture_DifferenceVariance_WGA_5_03_256,Nuclei|Texture_DifferenceVariance_WGA_5_02_256,Nuclei|Texture_DifferenceVariance_WGA_5_00_256,Nuclei|Texture_DifferenceVariance_WGA_5_01_256,Nuclei|Texture_DifferenceVariance_DNA_3_01_256,Nuclei|Texture_DifferenceVariance_DNA_3_03_256,Nuclei|Texture_DifferenceVariance_DNA_3_02_256,Nuclei|Texture_DifferenceVariance_DNA_3_00_256,Nuclei|Texture_DifferenceVariance_DNA_20_03_256,Nuclei|Texture_DifferenceVariance_DNA_20_00_256,Nuclei|Texture_DifferenceVariance_DNA_20_01_256,Nuclei|Texture_DifferenceVariance_DNA_20_02_256,Nuclei|Texture_DifferenceVariance_DNA_5_00_256,Nuclei|Texture_DifferenceVariance_DNA_5_02_256,Nuclei|Texture_DifferenceVariance_DNA_5_03_256,Nuclei|Texture_DifferenceVariance_DNA_5_01_256,Nuclei|Texture_DifferenceVariance_DNA_10_02_256,Nuclei|Texture_DifferenceVariance_DNA_10_01_256,Nuclei|Texture_DifferenceVariance_DNA_10_03_256,Nuclei|Texture_DifferenceVariance_DNA_10_00_256,Nuclei|Texture_DifferenceVariance_Vinculin_5_01_256,Nuclei|Texture_DifferenceVariance_Vinculin_5_03_256,Nuclei|Texture_DifferenceVariance_Vinculin_5_02_256,Nuclei|Texture_DifferenceVariance_Vinculin_5_00_256,Nuclei|Texture_DifferenceVariance_Vinculin_3_00_256,Nuclei|Texture_DifferenceVariance_Vinculin_3_01_256,Nuclei|Texture_DifferenceVariance_Vinculin_3_03_256,Nuclei|Texture_DifferenceVariance_Vinculin_3_02_256,Nuclei|Texture_DifferenceVariance_Vinculin_10_02_256,Nuclei|Texture_DifferenceVariance_Vinculin_10_03_256,Nuclei|Texture_DifferenceVariance_Vinculin_10_01_256,Nuclei|Texture_DifferenceVariance_Vinculin_10_00_256,Nuclei|Texture_DifferenceVariance_Vinculin_20_03_256,Nuclei|Texture_DifferenceVariance_Vinculin_20_01_256,Nuclei|Texture_DifferenceVariance_Vinculin_20_00_256,Nuclei|Texture_DifferenceVariance_Vinculin_20_02_256,Nuclei|Texture_InfoMeas1_Vinculin_5_00_256,Nuclei|Texture_InfoMeas1_Vinculin_5_01_256,Nuclei|Texture_InfoMeas1_Vinculin_5_03_256,Nuclei|Texture_InfoMeas1_Vinculin_5_02_256,Nuclei|Texture_InfoMeas1_Vinculin_3_02_256,Nuclei|Texture_InfoMeas1_Vinculin_3_00_256,Nuclei|Texture_InfoMeas1_Vinculin_3_01_256,Nuclei|Texture_InfoMeas1_Vinculin_3_03_256,Nuclei|Texture_InfoMeas1_Vinculin_20_00_256,Nuclei|Texture_InfoMeas1_Vinculin_20_03_256,Nuclei|Texture_InfoMeas1_Vinculin_20_01_256,Nuclei|Texture_InfoMeas1_Vinculin_20_02_256,Nuclei|Texture_InfoMeas1_Vinculin_10_02_256,Nuclei|Texture_InfoMeas1_Vinculin_10_03_256,Nuclei|Texture_InfoMeas1_Vinculin_10_00_256,Nuclei|Texture_InfoMeas1_Vinculin_10_01_256,Nuclei|Texture_InfoMeas1_DNA_5_02_256,Nuclei|Texture_InfoMeas1_DNA_5_01_256,Nuclei|Texture_InfoMeas1_DNA_5_03_256,Nuclei|Texture_InfoMeas1_DNA_5_00_256,Nuclei|Texture_InfoMeas1_DNA_20_01_256,Nuclei|Texture_InfoMeas1_DNA_20_03_256,Nuclei|Texture_InfoMeas1_DNA_20_02_256,Nuclei|Texture_InfoMeas1_DNA_20_00_256,Nuclei|Texture_InfoMeas1_DNA_3_03_256,Nuclei|Texture_InfoMeas1_DNA_3_00_256,Nuclei|Texture_InfoMeas1_DNA_3_02_256,Nuclei|Texture_InfoMeas1_DNA_3_01_256,Nuclei|Texture_InfoMeas1_DNA_10_01_256,Nuclei|Texture_InfoMeas1_DNA_10_02_256,Nuclei|Texture_InfoMeas1_DNA_10_03_256,Nuclei|Texture_InfoMeas1_DNA_10_00_256,Nuclei|Texture_InfoMeas1_WGA_20_02_256,Nuclei|Texture_InfoMeas1_WGA_20_01_256,Nuclei|Texture_InfoMeas1_WGA_20_03_256,Nuclei|Texture_InfoMeas1_WGA_20_00_256,Nuclei|Texture_InfoMeas1_WGA_3_01_256,Nuclei|Texture_InfoMeas1_WGA_3_00_256,Nuclei|Texture_InfoMeas1_WGA_3_03_256,Nuclei|Texture_InfoMeas1_WGA_3_02_256,Nuclei|Texture_InfoMeas1_WGA_5_03_256,Nuclei|Texture_InfoMeas1_WGA_5_01_256,Nuclei|Texture_InfoMeas1_WGA_5_00_256,Nuclei|Texture_InfoMeas1_WGA_5_02_256,Nuclei|Texture_InfoMeas1_WGA_10_00_256,Nuclei|Texture_InfoMeas1_WGA_10_03_256,Nuclei|Texture_InfoMeas1_WGA_10_02_256,Nuclei|Texture_InfoMeas1_WGA_10_01_256,Nuclei|Texture_InfoMeas1_Phalloidin_20_01_256,Nuclei|Texture_InfoMeas1_Phalloidin_20_02_256,Nuclei|Texture_InfoMeas1_Phalloidin_20_03_256,Nuclei|Texture_InfoMeas1_Phalloidin_20_00_256,Nuclei|Texture_InfoMeas1_Phalloidin_3_03_256,Nuclei|Texture_InfoMeas1_Phalloidin_3_00_256,Nuclei|Texture_InfoMeas1_Phalloidin_3_02_256,Nuclei|Texture_InfoMeas1_Phalloidin_3_01_256,Nuclei|Texture_InfoMeas1_Phalloidin_5_00_256,Nuclei|Texture_InfoMeas1_Phalloidin_5_01_256,Nuclei|Texture_InfoMeas1_Phalloidin_5_03_256,Nuclei|Texture_InfoMeas1_Phalloidin_5_02_256,Nuclei|Texture_InfoMeas1_Phalloidin_10_02_256,Nuclei|Texture_InfoMeas1_Phalloidin_10_00_256,Nuclei|Texture_InfoMeas1_Phalloidin_10_01_256,Nuclei|Texture_InfoMeas1_Phalloidin_10_03_256,Nuclei|Texture_Contrast_Vinculin_10_00_256,Nuclei|Texture_Contrast_Vinculin_10_03_256,Nuclei|Texture_Contrast_Vinculin_10_02_256,Nuclei|Texture_Contrast_Vinculin_10_01_256,Nuclei|Texture_Contrast_Vinculin_20_02_256,Nuclei|Texture_Contrast_Vinculin_20_01_256,Nuclei|Texture_Contrast_Vinculin_20_03_256,Nuclei|Texture_Contrast_Vinculin_20_00_256,Nuclei|Texture_Contrast_Vinculin_5_02_256,Nuclei|Texture_Contrast_Vinculin_5_01_256,Nuclei|Texture_Contrast_Vinculin_5_00_256,Nuclei|Texture_Contrast_Vinculin_5_03_256,Nuclei|Texture_Contrast_Vinculin_3_02_256,Nuclei|Texture_Contrast_Vinculin_3_01_256,Nuclei|Texture_Contrast_Vinculin_3_03_256,Nuclei|Texture_Contrast_Vinculin_3_00_256,Nuclei|Texture_Contrast_DNA_5_00_256,Nuclei|Texture_Contrast_DNA_5_01_256,Nuclei|Texture_Contrast_DNA_5_03_256,Nuclei|Texture_Contrast_DNA_5_02_256,Nuclei|Texture_Contrast_DNA_10_02_256,Nuclei|Texture_Contrast_DNA_10_01_256,Nuclei|Texture_Contrast_DNA_10_00_256,Nuclei|Texture_

Contrast_DNA_10_03_256,Nuclei|Texture_Contrast_DNA_3_01_256,Nuclei|Texture_Contrast_DNA_3_02_256,Nucle
i|Texture_Contrast_DNA_3_00_256,Nuclei|Texture_Contrast_DNA_3_03_256,Nuclei|Texture_Contrast_DNA_20_02_
256,Nuclei|Texture_Contrast_DNA_20_00_256,Nuclei|Texture_Contrast_DNA_20_01_256,Nuclei|Texture_Contrast_
DNA_20_03_256,Nuclei|Texture_Contrast_Phalloidin_5_03_256,Nuclei|Texture_Contrast_Phalloidin_5_01_256,Nucle
i|Texture_Contrast_Phalloidin_5_02_256,Nuclei|Texture_Contrast_Phalloidin_5_00_256,Nuclei|Texture_Contrast_Phal
loidin_3_01_256,Nuclei|Texture_Contrast_Phalloidin_3_02_256,Nuclei|Texture_Contrast_Phalloidin_3_03_256,Nuclei
|Texture_Contrast_Phalloidin_3_00_256,Nuclei|Texture_Contrast_Phalloidin_20_03_256,Nuclei|Texture_Contrast_Pha
lloidin_20_02_256,Nuclei|Texture_Contrast_Phalloidin_20_00_256,Nuclei|Texture_Contrast_Phalloidin_20_01_256,N
uclei|Texture_Contrast_Phalloidin_10_00_256,Nuclei|Texture_Contrast_Phalloidin_10_02_256,Nuclei|Texture_Contra
st_Phalloidin_10_01_256,Nuclei|Texture_Contrast_Phalloidin_10_03_256,Nuclei|Texture_Contrast_WGA_3_00_256,
Nuclei|Texture_Contrast_WGA_3_01_256,Nuclei|Texture_Contrast_WGA_3_03_256,Nuclei|Texture_Contrast_WGA_
3_02_256,Nuclei|Texture_Contrast_WGA_5_03_256,Nuclei|Texture_Contrast_WGA_5_01_256,Nuclei|Texture_Contr
ast_WGA_5_00_256,Nuclei|Texture_Contrast_WGA_5_02_256,Nuclei|Texture_Contrast_WGA_20_03_256,Nuclei|Te
xture_Contrast_WGA_20_02_256,Nuclei|Texture_Contrast_WGA_20_01_256,Nuclei|Texture_Contrast_WGA_20_00
_256,Nuclei|Texture_Contrast_WGA_10_00_256,Nuclei|Texture_Contrast_WGA_10_01_256,Nuclei|Texture_Contrast
_WGA_10_03_256,Nuclei|Texture_Contrast_WGA_10_02_256,Nuclei|Texture_InverseDifferenceMoment_DNA_5_0
3_256,Nuclei|Texture_InverseDifferenceMoment_DNA_5_02_256,Nuclei|Texture_InverseDifferenceMoment_DNA_5
_01_256,Nuclei|Texture_InverseDifferenceMoment_DNA_5_00_256,Nuclei|Texture_InverseDifferenceMoment_DNA
_3_00_256,Nuclei|Texture_InverseDifferenceMoment_DNA_3_01_256,Nuclei|Texture_InverseDifferenceMoment_DN
A_3_03_256,Nuclei|Texture_InverseDifferenceMoment_DNA_3_02_256,Nuclei|Texture_InverseDifferenceMoment_D
NA_20_01_256,Nuclei|Texture_InverseDifferenceMoment_DNA_20_00_256,Nuclei|Texture_InverseDifferenceMome
nt_DNA_20_02_256,Nuclei|Texture_InverseDifferenceMoment_DNA_20_03_256,Nuclei|Texture_InverseDifferenceM
oment_DNA_10_03_256,Nuclei|Texture_InverseDifferenceMoment_DNA_10_01_256,Nuclei|Texture_InverseDifferen
ceMoment_DNA_10_02_256,Nuclei|Texture_InverseDifferenceMoment_DNA_10_00_256,Nuclei|Texture_InverseDiff
erenceMoment_Vinculin_5_02_256,Nuclei|Texture_InverseDifferenceMoment_Vinculin_5_00_256,Nuclei|Texture_In
verseDifferenceMoment_Vinculin_5_03_256,Nuclei|Texture_InverseDifferenceMoment_Vinculin_5_01_256,Nuclei|Te
xture_InverseDifferenceMoment_Vinculin_10_03_256,Nuclei|Texture_InverseDifferenceMoment_Vinculin_10_00_25
6,Nuclei|Texture_InverseDifferenceMoment_Vinculin_10_02_256,Nuclei|Texture_InverseDifferenceMoment_Vinculin
_10_01_256,Nuclei|Texture_InverseDifferenceMoment_Vinculin_20_00_256,Nuclei|Texture_InverseDifferenceMome
nt_Vinculin_20_02_256,Nuclei|Texture_InverseDifferenceMoment_Vinculin_20_03_256,Nuclei|Texture_InverseDiffer
enceMoment_Vinculin_20_01_256,Nuclei|Texture_InverseDifferenceMoment_Vinculin_3_02_256,Nuclei|Texture_Inv
erseDifferenceMoment_Vinculin_3_01_256,Nuclei|Texture_InverseDifferenceMoment_Vinculin_3_00_256,Nuclei|Tex
ture_InverseDifferenceMoment_Vinculin_3_03_256,Nuclei|Texture_InverseDifferenceMoment_WGA_3_01_256,Nucl
ei|Texture_InverseDifferenceMoment_WGA_3_03_256,Nuclei|Texture_InverseDifferenceMoment_WGA_3_02_256,N
uclei|Texture_InverseDifferenceMoment_WGA_3_00_256,Nuclei|Texture_InverseDifferenceMoment_WGA_10_01_2
56,Nuclei|Texture_InverseDifferenceMoment_WGA_10_00_256,Nuclei|Texture_InverseDifferenceMoment_WGA_10
_03_256,Nuclei|Texture_InverseDifferenceMoment_WGA_10_02_256,Nuclei|Texture_InverseDifferenceMoment_WG
A_20_00_256,Nuclei|Texture_InverseDifferenceMoment_WGA_20_01_256,Nuclei|Texture_InverseDifferenceMoment
_WGA_20_02_256,Nuclei|Texture_InverseDifferenceMoment_WGA_20_03_256,Nuclei|Texture_InverseDifferenceM
oment_WGA_5_01_256,Nuclei|Texture_InverseDifferenceMoment_WGA_5_00_256,Nuclei|Texture_InverseDifferenc
eMoment_WGA_5_02_256,Nuclei|Texture_InverseDifferenceMoment_WGA_5_03_256,Nuclei|Texture_InverseDiffer
enceMoment_Phalloidin_3_03_256,Nuclei|Texture_InverseDifferenceMoment_Phalloidin_3_00_256,Nuclei|Texture_I
nverseDifferenceMoment_Phalloidin_3_01_256,Nuclei|Texture_InverseDifferenceMoment_Phalloidin_3_02_256,Nucl
ei|Texture_InverseDifferenceMoment_Phalloidin_10_03_256,Nuclei|Texture_InverseDifferenceMoment_Phalloidin_10
_01_256,Nuclei|Texture_InverseDifferenceMoment_Phalloidin_10_02_256,Nuclei|Texture_InverseDifferenceMoment_
Phalloidin_10_00_256,Nuclei|Texture_InverseDifferenceMoment_Phalloidin_5_03_256,Nuclei|Texture_InverseDiffere
nceMoment_Phalloidin_5_00_256,Nuclei|Texture_InverseDifferenceMoment_Phalloidin_5_01_256,Nuclei|Texture_In
verseDifferenceMoment_Phalloidin_5_02_256,Nuclei|Texture_InverseDifferenceMoment_Phalloidin_20_03_256,Nucl
ei|Texture_InverseDifferenceMoment_Phalloidin_20_02_256,Nuclei|Texture_InverseDifferenceMoment_Phalloidin_20
_00_256,Nuclei|Texture_InverseDifferenceMoment_Phalloidin_20_01_256,Nuclei|Texture_AngularSecondMoment_Vi
nculin_10_02_256,Nuclei|Texture_AngularSecondMoment_Vinculin_10_00_256,Nuclei|Texture_AngularSecondMom
ent_Vinculin_10_03_256,Nuclei|Texture_AngularSecondMoment_Vinculin_10_01_256,Nuclei|Texture_AngularSecon
dMoment_Vinculin_3_00_256,Nuclei|Texture_AngularSecondMoment_Vinculin_3_03_256,Nuclei|Texture_AngularSe
condMoment_Vinculin_3_02_256,Nuclei|Texture_AngularSecondMoment_Vinculin_3_01_256,Nuclei|Texture_Angul

arSecondMoment_Vinculin_20_02_256,Nuclei|Texture_AngularSecondMoment_Vinculin_20_01_256,Nuclei|Texture_
AngularSecondMoment_Vinculin_20_03_256,Nuclei|Texture_AngularSecondMoment_Vinculin_20_00_256,Nuclei|Te
xture_AngularSecondMoment_Vinculin_5_03_256,Nuclei|Texture_AngularSecondMoment_Vinculin_5_01_256,Nucle
i|Texture_AngularSecondMoment_Vinculin_5_00_256,Nuclei|Texture_AngularSecondMoment_Vinculin_5_02_256,N
uclei|Texture_AngularSecondMoment_Phalloidin_3_03_256,Nuclei|Texture_AngularSecondMoment_Phalloidin_3_01
_256,Nuclei|Texture_AngularSecondMoment_Phalloidin_3_02_256,Nuclei|Texture_AngularSecondMoment_Phalloidi
n_3_00_256,Nuclei|Texture_AngularSecondMoment_Phalloidin_5_02_256,Nuclei|Texture_AngularSecondMoment_P
halloidin_5_01_256,Nuclei|Texture_AngularSecondMoment_Phalloidin_5_03_256,Nuclei|Texture_AngularSecondMo
ment_Phalloidin_5_00_256,Nuclei|Texture_AngularSecondMoment_Phalloidin_20_01_256,Nuclei|Texture_AngularSe
condMoment_Phalloidin_20_02_256,Nuclei|Texture_AngularSecondMoment_Phalloidin_20_00_256,Nuclei|Texture_
AngularSecondMoment_Phalloidin_20_03_256,Nuclei|Texture_AngularSecondMoment_Phalloidin_10_03_256,Nuclei|
Texture_AngularSecondMoment_Phalloidin_10_01_256,Nuclei|Texture_AngularSecondMoment_Phalloidin_10_02_25
6,Nuclei|Texture_AngularSecondMoment_Phalloidin_10_00_256,Nuclei|Texture_AngularSecondMoment_WGA_20_0
3_256,Nuclei|Texture_AngularSecondMoment_WGA_20_02_256,Nuclei|Texture_AngularSecondMoment_WGA_20_
00_256,Nuclei|Texture_AngularSecondMoment_WGA_20_01_256,Nuclei|Texture_AngularSecondMoment_WGA_5_
01_256,Nuclei|Texture_AngularSecondMoment_WGA_5_02_256,Nuclei|Texture_AngularSecondMoment_WGA_5_0
0_256,Nuclei|Texture_AngularSecondMoment_WGA_5_03_256,Nuclei|Texture_AngularSecondMoment_WGA_3_01
_256,Nuclei|Texture_AngularSecondMoment_WGA_3_00_256,Nuclei|Texture_AngularSecondMoment_WGA_3_03_
256,Nuclei|Texture_AngularSecondMoment_WGA_3_02_256,Nuclei|Texture_AngularSecondMoment_WGA_10_03_
256,Nuclei|Texture_AngularSecondMoment_WGA_10_02_256,Nuclei|Texture_AngularSecondMoment_WGA_10_01
_256,Nuclei|Texture_AngularSecondMoment_WGA_10_00_256,Nuclei|Texture_AngularSecondMoment_DNA_5_03_
256,Nuclei|Texture_AngularSecondMoment_DNA_5_01_256,Nuclei|Texture_AngularSecondMoment_DNA_5_00_25
6,Nuclei|Texture_AngularSecondMoment_DNA_5_02_256,Nuclei|Texture_AngularSecondMoment_DNA_10_02_256,
Nuclei|Texture_AngularSecondMoment_DNA_10_00_256,Nuclei|Texture_AngularSecondMoment_DNA_10_03_256,
Nuclei|Texture_AngularSecondMoment_DNA_10_01_256,Nuclei|Texture_AngularSecondMoment_DNA_20_01_256,
Nuclei|Texture_AngularSecondMoment_DNA_20_02_256,Nuclei|Texture_AngularSecondMoment_DNA_20_00_256,
Nuclei|Texture_AngularSecondMoment_DNA_20_03_256,Nuclei|Texture_AngularSecondMoment_DNA_3_02_256,N
uclei|Texture_AngularSecondMoment_DNA_3_01_256,Nuclei|Texture_AngularSecondMoment_DNA_3_03_256,Nucl
ei|Texture_AngularSecondMoment_DNA_3_00_256,Nuclei|Texture_SumAverage_Vinculin_3_00_256,Nuclei|Texture
_SumAverage_Vinculin_3_03_256,Nuclei|Texture_SumAverage_Vinculin_3_01_256,Nuclei|Texture_SumAverage_Vi
nculin_3_02_256,Nuclei|Texture_SumAverage_Vinculin_5_02_256,Nuclei|Texture_SumAverage_Vinculin_5_00_256,
Nuclei|Texture_SumAverage_Vinculin_5_01_256,Nuclei|Texture_SumAverage_Vinculin_5_03_256,Nuclei|Texture_S
umAverage_Vinculin_10_00_256,Nuclei|Texture_SumAverage_Vinculin_10_02_256,Nuclei|Texture_SumAverage_Vi
nculin_10_01_256,Nuclei|Texture_SumAverage_Vinculin_10_03_256,Nuclei|Texture_SumAverage_Vinculin_20_02_
256,Nuclei|Texture_SumAverage_Vinculin_20_00_256,Nuclei|Texture_SumAverage_Vinculin_20_03_256,Nuclei|Tex
ture_SumAverage_Vinculin_20_01_256,Nuclei|Texture_SumAverage_Phalloidin_5_00_256,Nuclei|Texture_SumAver
age_Phalloidin_5_01_256,Nuclei|Texture_SumAverage_Phalloidin_5_03_256,Nuclei|Texture_SumAverage_Phalloidin
_5_02_256,Nuclei|Texture_SumAverage_Phalloidin_3_02_256,Nuclei|Texture_SumAverage_Phalloidin_3_00_256,Nu
clei|Texture_SumAverage_Phalloidin_3_01_256,Nuclei|Texture_SumAverage_Phalloidin_3_03_256,Nuclei|Texture_S
umAverage_Phalloidin_20_01_256,Nuclei|Texture_SumAverage_Phalloidin_20_00_256,Nuclei|Texture_SumAverage_
Phalloidin_20_03_256,Nuclei|Texture_SumAverage_Phalloidin_20_02_256,Nuclei|Texture_SumAverage_Phalloidin_1
0_00_256,Nuclei|Texture_SumAverage_Phalloidin_10_01_256,Nuclei|Texture_SumAverage_Phalloidin_10_02_256,N
uclei|Texture_SumAverage_Phalloidin_10_03_256,Nuclei|Texture_SumAverage_WGA_20_03_256,Nuclei|Texture_Su
mAverage_WGA_20_00_256,Nuclei|Texture_SumAverage_WGA_20_01_256,Nuclei|Texture_SumAverage_WGA_20
_02_256,Nuclei|Texture_SumAverage_WGA_10_03_256,Nuclei|Texture_SumAverage_WGA_10_02_256,Nuclei|Text
ure_SumAverage_WGA_10_00_256,Nuclei|Texture_SumAverage_WGA_10_01_256,Nuclei|Texture_SumAverage_W
GA_5_02_256,Nuclei|Texture_SumAverage_WGA_5_03_256,Nuclei|Texture_SumAverage_WGA_5_01_256,Nuclei|
Texture_SumAverage_WGA_5_00_256,Nuclei|Texture_SumAverage_WGA_3_01_256,Nuclei|Texture_SumAverage_
WGA_3_02_256,Nuclei|Texture_SumAverage_WGA_3_00_256,Nuclei|Texture_SumAverage_WGA_3_03_256,Nucle
i|Texture_SumAverage_DNA_10_01_256,Nuclei|Texture_SumAverage_DNA_10_00_256,Nuclei|Texture_SumAverag
e_DNA_10_03_256,Nuclei|Texture_SumAverage_DNA_10_02_256,Nuclei|Texture_SumAverage_DNA_20_01_256,N
uclei|Texture_SumAverage_DNA_20_03_256,Nuclei|Texture_SumAverage_DNA_20_00_256,Nuclei|Texture_SumAv
erage_DNA_20_02_256,Nuclei|Texture_SumAverage_DNA_5_01_256,Nuclei|Texture_SumAverage_DNA_5_02_256,
Nuclei|Texture_SumAverage_DNA_5_00_256,Nuclei|Texture_SumAverage_DNA_5_03_256,Nuclei|Texture_SumAve

rage_DNA_3_01_256,Nuclei|Texture_SumAverage_DNA_3_03_256,Nuclei|Texture_SumAverage_DNA_3_02_256,Nuclei|Texture_SumAverage_DNA_3_00_256,Nuclei|Granularity_8_Vinculin,Nuclei|Granularity_8_Phalloidin,Nuclei|Granularity_8_WGA,Nuclei|Granularity_8_DNA,Nuclei|Granularity_13_DNA,Nuclei|Granularity_13_Phalloidin,Nuclei|Granularity_13_WGA,Nuclei|Granularity_13_Vinculin,Nuclei|Granularity_2_DNA,Nuclei|Granularity_2_WGA,Nuclei|Granularity_2_Phalloidin,Nuclei|Granularity_2_Vinculin,Nuclei|Granularity_5_DNA,Nuclei|Granularity_5_WGA,Nuclei|Granularity_5_Phalloidin,Nuclei|Granularity_5_Vinculin,Nuclei|Granularity_6_Vinculin,Nuclei|Granularity_6_WGA,Nuclei|Granularity_6_Phalloidin,Nuclei|Granularity_6_DNA,Nuclei|Granularity_1_Phalloidin,Nuclei|Granularity_1_Vinculin,Nuclei|Granularity_1_WGA,Nuclei|Granularity_1_DNA,Nuclei|Granularity_16_DNA,Nuclei|Granularity_16_Vinculin,Nuclei|Granularity_16_Phalloidin,Nuclei|Granularity_16_WGA,Nuclei|Granularity_14_WGA,Nuclei|Granularity_14_DNA,Nuclei|Granularity_14_Phalloidin,Nuclei|Granularity_14_Vinculin,Nuclei|Granularity_12_DNA,Nuclei|Granularity_12_Vinculin,Nuclei|Granularity_12_WGA,Nuclei|Granularity_12_Phalloidin,Nuclei|Granularity_4_Vinculin,Nuclei|Granularity_4_DNA,Nuclei|Granularity_4_Phalloidin,Nuclei|Granularity_4_WGA,Nuclei|Granularity_9_Phalloidin,Nuclei|Granularity_9_WGA,Nuclei|Granularity_9_Vinculin,Nuclei|Granularity_9_DNA,Nuclei|Granularity_11_Vinculin,Nuclei|Granularity_11_WGA,Nuclei|Granularity_11_DNA,Nuclei|Granularity_11_Phalloidin,Nuclei|Granularity_15_Phalloidin,Nuclei|Granularity_15_DNA,Nuclei|Granularity_15_Vinculin,Nuclei|Granularity_15_WGA,Nuclei|Granularity_7_WGA,Nuclei|Granularity_7_Phalloidin,Nuclei|Granularity_7_Vinculin,Nuclei|Granularity_7_DNA,Nuclei|Granularity_10_Vinculin,Nuclei|Granularity_10_Phalloidin,Nuclei|Granularity_10_DNA,Nuclei|Granularity_10_WGA,Nuclei|Granularity_3_Phalloidin,Nuclei|Granularity_3_DNA,Nuclei|Granularity_3_WGA,Nuclei|Granularity_3_Vinculin,Nuclei|Correlation_K_Phalloidin_WGA,Nuclei|Correlation_K_Phalloidin_DNA,Nuclei|Correlation_K_Phalloidin_Vinculin,Nuclei|Correlation_K_Vinculin_WGA,Nuclei|Correlation_K_Vinculin_DNA,Nuclei|Correlation_K_Vinculin_Phalloidin,Nuclei|Correlation_K_WGA_Vinculin,Nuclei|Correlation_K_WGA_Phalloidin,Nuclei|Correlation_K_WGA_DNA,Nuclei|Correlation_K_DNA_Phalloidin,Nuclei|Correlation_K_DNA_WGA,Nuclei|Correlation_K_DNA_Vinculin,Nuclei|Correlation_Manders_DNA_Vinculin,Nuclei|Correlation_Manders_DNA_Phalloidin,Nuclei|Correlation_Manders_DNA_WGA,Nuclei|Correlation_Manders_WGA_DNA,Nuclei|Correlation_Manders_WGA_Phalloidin,Nuclei|Correlation_Manders_WGA_Vinculin,Nuclei|Correlation_Manders_Vinculin_DNA,Nuclei|Correlation_Manders_Vinculin_Phalloidin,Nuclei|Correlation_Manders_Vinculin_WGA,Nuclei|Correlation_Manders_Phalloidin_DNA,Nuclei|Correlation_Manders_Phalloidin_WGA,Nuclei|Correlation_Manders_Phalloidin_Vinculin,Nuclei|Correlation_RWC_DNA_WGA,Nuclei|Correlation_RWC_DNA_Vinculin,Nuclei|Correlation_RWC_DNA_Phalloidin,Nuclei|Correlation_RWC_Vinculin_DNA,Nuclei|Correlation_RWC_Vinculin_Phalloidin,Nuclei|Correlation_RWC_Vinculin_WGA,Nuclei|Correlation_RWC_Phalloidin_DNA,Nuclei|Correlation_RWC_Phalloidin_Vinculin,Nuclei|Correlation_RWC_Phalloidin_WGA,Nuclei|Correlation_RWC_WGA_Phalloidin,Nuclei|Correlation_RWC_WGA_Vinculin,Nuclei|Correlation_RWC_WGA_DNA,Nuclei|Correlation_Overlap_Phalloidin_WGA,Nuclei|Correlation_Overlap_Phalloidin_Vinculin,Nuclei|Correlation_Overlap_Vinculin_WGA,Nuclei|Correlation_Overlap_DNA_Vinculin,Nuclei|Correlation_Overlap_DNA_WGA,Nuclei|Correlation_Overlap_DNA_Phalloidin,Nuclei|Correlation_Correlation_DNA_WGA,Nuclei|Correlation_Correlation_DNA_Phalloidin,Nuclei|Correlation_Correlation_DNA_Vinculin,Nuclei|Correlation_Correlation_Phalloidin_Vinculin,Nuclei|Correlation_Correlation_Phalloidin_WGA,Nuclei|Correlation_Correlation_Vinculin_WGA,Nuclei|RadialDistribution_RadialCV_WGA_3of4,Nuclei|RadialDistribution_RadialCV_WGA_1of4,Nuclei|RadialDistribution_RadialCV_WGA_2of4,Nuclei|RadialDistribution_RadialCV_WGA_4of4,Nuclei|RadialDistribution_RadialCV_Vinculin_4of4,Nuclei|RadialDistribution_RadialCV_Vinculin_1of4,Nuclei|RadialDistribution_RadialCV_Vinculin_3of4,Nuclei|RadialDistribution_RadialCV_Vinculin_2of4,Nuclei|RadialDistribution_RadialCV_DNA_2of4,Nuclei|RadialDistribution_RadialCV_DNA_4of4,Nuclei|RadialDistribution_RadialCV_DNA_1of4,Nuclei|RadialDistribution_RadialCV_DNA_3of4,Nuclei|RadialDistribution_RadialCV_Phalloidin_2of4,Nuclei|RadialDistribution_RadialCV_Phalloidin_4of4,Nuclei|RadialDistribution_RadialCV_Phalloidin_1of4,Nuclei|RadialDistribution_RadialCV_Phalloidin_3of4,Nuclei|RadialDistribution_FracAtD_WGA_1of4,Nuclei|RadialDistribution_FracAtD_WGA_4of4,Nuclei|RadialDistribution_FracAtD_WGA_3of4,Nuclei|RadialDistribution_FracAtD_WGA_2of4,Nuclei|RadialDistribution_FracAtD_Phalloidin_2of4,Nuclei|RadialDistribution_FracAtD_Phalloidin_3of4,Nuclei|RadialDistribution_FracAtD_Phalloidin_1of4,Nuclei|RadialDistribution_FracAtD_Phalloidin_4of4,Nuclei|RadialDistribution_FracAtD_Vinculin_1of4,Nuclei|RadialDistribution_FracAtD_Vinculin_2of4,Nuclei|RadialDistribution_FracAtD_Vinculin_3of4,Nuclei|RadialDistribution_FracAtD_Vinculin_4of4,Nuclei|RadialDistribution_FracAtD_DNA_2of4,Nuclei|RadialDistribution_FracAtD_DNA_1of4,Nuclei|RadialDistribution_FracAtD_DNA_4of4,Nuclei|RadialDistribution_FracAtD_DNA_3of4,Nuclei|RadialDistribution_MeanFrac_Phalloidin_2of4,Nuclei|RadialDistribution_MeanFrac_Phalloidin_4of4,Nuclei|RadialDistribution_MeanFrac_Phalloidin_3of4,Nuclei|RadialDistribution_MeanFrac_Phalloidin_1of4,Nuclei|RadialDistribution_MeanFrac_DNA_1of4,Nuclei|RadialDistribution_MeanFrac_DNA_3of4,Nuclei|RadialDistribution_MeanFrac_DNA_4of4,Nuclei|RadialDistribution_MeanFrac_DNA_2of4,Nuclei|RadialDistribution_MeanFrac_Vinculin_3of4,Nuclei|RadialDistribution_MeanFrac_Vinculin_4of

4,Nuclei|RadialDistribution_MeanFrac_Vinculin_2of4,Nuclei|RadialDistribution_MeanFrac_Vinculin_1of4,Nuclei|RadialDistribution_MeanFrac_WGA_2of4,Nuclei|RadialDistribution_MeanFrac_WGA_4of4,Nuclei|RadialDistribution_MeanFrac_WGA_3of4,Nuclei|RadialDistribution_MeanFrac_WGA_1of4,Nuclei|AreaShape_MaximumRadius,Nuclei|AreaShape_Zernike_6_4,Nuclei|AreaShape_Zernike_6_2,Nuclei|AreaShape_Zernike_6_6,Nuclei|AreaShape_Zernike_6_0,Nuclei|AreaShape_Zernike_2_2,Nuclei|AreaShape_Zernike_2_0,Nuclei|AreaShape_Zernike_9_9,Nuclei|AreaShape_Zernike_9_1,Nuclei|AreaShape_Zernike_9_3,Nuclei|AreaShape_Zernike_9_7,Nuclei|AreaShape_Zernike_9_5,Nuclei|AreaShape_Zernike_5_3,Nuclei|AreaShape_Zernike_5_1,Nuclei|AreaShape_Zernike_5_5,Nuclei|AreaShape_Zernike_8_6,Nuclei|AreaShape_Zernike_8_8,Nuclei|AreaShape_Zernike_8_4,Nuclei|AreaShape_Zernike_8_0,Nuclei|AreaShape_Zernike_8_2,Nuclei|AreaShape_Zernike_1_1,Nuclei|AreaShape_Zernike_4_2,Nuclei|AreaShape_Zernike_4_4,Nuclei|AreaShape_Zernike_4_0,Nuclei|AreaShape_Zernike_7_1,Nuclei|AreaShape_Zernike_7_3,Nuclei|AreaShape_Zernike_7_7,Nuclei|AreaShape_Zernike_7_5,Nuclei|AreaShape_Zernike_0_0,Nuclei|AreaShape_Zernike_3_1,Nuclei|AreaShape_Zernike_3_3,Nuclei|AreaShape_BoundingBoxArea,Nuclei|AreaShape_BoundingBoxMaximum_X,Nuclei|AreaShape_BoundingBoxMaximum_Y,Nuclei|AreaShape_EquivalentDiameter,Nuclei|AreaShape_ConvexArea,Nuclei|AreaShape_MinorAxisLength,Nuclei|AreaShape_MajorAxisLength,Nuclei|AreaShape_Area,Nuclei|AreaShape_Eccentricity,Nuclei|AreaShape_FormFactor,Nuclei|AreaShape_Extent,Nuclei|AreaShape_MaxFeretDiameter,Nuclei|AreaShape_MeanRadius,Nuclei|AreaShape_MedianRadius,Nuclei|AreaShape_Orientation,Nuclei|AreaShape_MinFeretDiameter,Nuclei|AreaShape_Center_X,Nuclei|AreaShape_Center_Y,Nuclei|AreaShape_BoundingBoxMinimum_Y,Nuclei|AreaShape_BoundingBoxMinimum_X,Nuclei|AreaShape_EulerNumber,Nuclei|AreaShape_Compactness,Nuclei|AreaShape_Perimeter,Nuclei|AreaShape_Solidity,Nuclei|Location_MaxIntensity_Z_WGA,Nuclei|Location_MaxIntensity_Z_Phalloidin,Nuclei|Location_MaxIntensity_Z_DNA,Nuclei|Location_MaxIntensity_Z_Vinculin,Nuclei|Location_MaxIntensity_Y_WGA,Nuclei|Location_MaxIntensity_Y_Phalloidin,Nuclei|Location_MaxIntensity_Y_DNA,Nuclei|Location_MaxIntensity_Y_Vinculin,Nuclei|Location_MaxIntensity_X_Vinculin,Nuclei|Location_MaxIntensity_X_DNA,Nuclei|Location_MaxIntensity_X_WGA,Nuclei|Location_MaxIntensity_X_Phalloidin,Nuclei|Location_Center_Y,Nuclei|Location_Center_Z,Nuclei|Location_Center_X,Nuclei|Location_CenterMassIntensity_Y_WGA,Nuclei|Location_CenterMassIntensity_Y_DNA,Nuclei|Location_CenterMassIntensity_Y_Vinculin,Nuclei|Location_CenterMassIntensity_Y_Phalloidin,Nuclei|Location_CenterMassIntensity_X_DNA,Nuclei|Location_CenterMassIntensity_X_WGA,Nuclei|Location_CenterMassIntensity_X_Vinculin,Nuclei|Location_CenterMassIntensity_X_Phalloidin,Nuclei|Location_CenterMassIntensity_Z_WGA,Nuclei|Location_CenterMassIntensity_Z_DNA,Nuclei|Location_CenterMassIntensity_Z_Phalloidin,Nuclei|Location_CenterMassIntensity_Z_Vinculin,Nuclei|Neighbors_FirstClosestObjectNumber_2,Nuclei|Neighbors_PercentTouching_2,Nuclei|Neighbors_FirstClosestDistance_2,Nuclei|Neighbors_SecondClosestDistance_2,Nuclei|Neighbors_NumberOfNeighbors_2,Nuclei|Neighbors_AngleBetweenNeighbors_2,Nuclei|Neighbors_SecondClosestObjectNumber_2,Nuclei|Number_Object_Number,Nuclei|Children_Cytoplasm_Count,Nuclei|Parent_NucleiIncludingEdges,Cytoplasm|Texture_DifferenceVariance_WGA_5_01_256,Cytoplasm|Texture_DifferenceVariance_WGA_5_03_256,Cytoplasm|Texture_DifferenceVariance_WGA_5_02_256,Cytoplasm|Texture_DifferenceVariance_WGA_5_00_256,Cytoplasm|Texture_DifferenceVariance_WGA_3_03_256,Cytoplasm|Texture_DifferenceVariance_WGA_3_01_256,Cytoplasm|Texture_DifferenceVariance_WGA_3_02_256,Cytoplasm|Texture_DifferenceVariance_WGA_3_00_256,Cytoplasm|Texture_DifferenceVariance_WGA_20_02_256,Cytoplasm|Texture_DifferenceVariance_WGA_20_03_256,Cytoplasm|Texture_DifferenceVariance_WGA_20_01_256,Cytoplasm|Texture_DifferenceVariance_WGA_20_00_256,Cytoplasm|Texture_DifferenceVariance_WGA_10_02_256,Cytoplasm|Texture_DifferenceVariance_WGA_10_03_256,Cytoplasm|Texture_DifferenceVariance_WGA_10_00_256,Cytoplasm|Texture_DifferenceVariance_WGA_10_01_256,Cytoplasm|Texture_DifferenceVariance_Phalloidin_20_00_256,Cytoplasm|Texture_DifferenceVariance_Phalloidin_20_02_256,Cytoplasm|Texture_DifferenceVariance_Phalloidin_20_03_256,Cytoplasm|Texture_DifferenceVariance_Phalloidin_20_01_256,Cytoplasm|Texture_DifferenceVariance_Phalloidin_5_00_256,Cytoplasm|Texture_DifferenceVariance_Phalloidin_5_02_256,Cytoplasm|Texture_DifferenceVariance_Phalloidin_5_03_256,Cytoplasm|Texture_DifferenceVariance_Phalloidin_5_01_256,Cytoplasm|Texture_DifferenceVariance_Phalloidin_3_03_256,Cytoplasm|Texture_DifferenceVariance_Phalloidin_3_02_256,Cytoplasm|Texture_DifferenceVariance_Phalloidin_3_00_256,Cytoplasm|Texture_DifferenceVariance_Phalloidin_3_01_256,Cytoplasm|Texture_DifferenceVariance_Phalloidin_10_02_256,Cytoplasm|Texture_DifferenceVariance_Phalloidin_10_03_256,Cytoplasm|Texture_DifferenceVariance_Phalloidin_10_00_256,Cytoplasm|Texture_DifferenceVariance_Phalloidin_10_01_256,Cytoplasm|Texture_DifferenceVariance_DNA_5_00_256,Cytoplasm|Texture_DifferenceVariance_DNA_5_02_256,Cytoplasm|Texture_DifferenceVariance_DNA_5_03_256,Cytoplasm|Texture_DifferenceVariance_DNA_5_01_256,Cytoplasm|Texture_DifferenceVariance_DNA_3_03_256,Cytoplasm|Texture_DifferenceVariance_DNA_3_01_256,Cytoplasm|Texture_DifferenceVariance_DNA_3_02_256,Cytoplasm|Texture_DifferenceVariance_DNA_3_00_256,Cytoplasm|Texture_DifferenceVariance_DNA_20_01_256,Cytoplasm|Texture_DifferenceVariance_DNA_20_00_256,Cytoplasm|Texture_DifferenceVariance_DNA_20_02_256,Cytoplasm|Texture_DifferenceVariance_DN

A_20_03_256,Cytoplasm|Texture_DifferenceVariance_DNA_10_02_256,Cytoplasm|Texture_DifferenceVariance_DN
A_10_03_256,Cytoplasm|Texture_DifferenceVariance_DNA_10_00_256,Cytoplasm|Texture_DifferenceVariance_DN
A_10_01_256,Cytoplasm|Texture_DifferenceVariance_Vinculin_10_01_256,Cytoplasm|Texture_DifferenceVariance_
Vinculin_10_02_256,Cytoplasm|Texture_DifferenceVariance_Vinculin_10_00_256,Cytoplasm|Texture_DifferenceVari
ance_Vinculin_10_03_256,Cytoplasm|Texture_DifferenceVariance_Vinculin_5_01_256,Cytoplasm|Texture_Difference
Variance_Vinculin_5_02_256,Cytoplasm|Texture_DifferenceVariance_Vinculin_5_03_256,Cytoplasm|Texture_Differe
nceVariance_Vinculin_5_00_256,Cytoplasm|Texture_DifferenceVariance_Vinculin_3_00_256,Cytoplasm|Texture_Dif
ferenceVariance_Vinculin_3_03_256,Cytoplasm|Texture_DifferenceVariance_Vinculin_3_01_256,Cytoplasm|Texture_
DifferenceVariance_Vinculin_3_02_256,Cytoplasm|Texture_DifferenceVariance_Vinculin_20_03_256,Cytoplasm|Text
ure_DifferenceVariance_Vinculin_20_02_256,Cytoplasm|Texture_DifferenceVariance_Vinculin_20_00_256,Cytoplas
m|Texture_DifferenceVariance_Vinculin_20_01_256,Cytoplasm|Texture_SumAverage_WGA_5_01_256,Cytoplasm|Te
xture_SumAverage_WGA_5_02_256,Cytoplasm|Texture_SumAverage_WGA_5_03_256,Cytoplasm|Texture_SumAve
rage_WGA_5_00_256,Cytoplasm|Texture_SumAverage_WGA_10_00_256,Cytoplasm|Texture_SumAverage_WGA_1
0_01_256,Cytoplasm|Texture_SumAverage_WGA_10_03_256,Cytoplasm|Texture_SumAverage_WGA_10_02_256,C
ytoplasm|Texture_SumAverage_WGA_3_00_256,Cytoplasm|Texture_SumAverage_WGA_3_03_256,Cytoplasm|Textu
re_SumAverage_WGA_3_02_256,Cytoplasm|Texture_SumAverage_WGA_3_01_256,Cytoplasm|Texture_SumAverag
e_WGA_20_01_256,Cytoplasm|Texture_SumAverage_WGA_20_02_256,Cytoplasm|Texture_SumAverage_WGA_20
_03_256,Cytoplasm|Texture_SumAverage_WGA_20_00_256,Cytoplasm|Texture_SumAverage_Vinculin_10_01_256,
Cytoplasm|Texture_SumAverage_Vinculin_10_02_256,Cytoplasm|Texture_SumAverage_Vinculin_10_03_256,Cytopl
asm|Texture_SumAverage_Vinculin_10_00_256,Cytoplasm|Texture_SumAverage_Vinculin_20_00_256,Cytoplasm|Te
xture_SumAverage_Vinculin_20_02_256,Cytoplasm|Texture_SumAverage_Vinculin_20_03_256,Cytoplasm|Texture_
SumAverage_Vinculin_20_01_256,Cytoplasm|Texture_SumAverage_Vinculin_3_00_256,Cytoplasm|Texture_SumAve
rage_Vinculin_3_03_256,Cytoplasm|Texture_SumAverage_Vinculin_3_01_256,Cytoplasm|Texture_SumAverage_Vin
culin_3_02_256,Cytoplasm|Texture_SumAverage_Vinculin_5_02_256,Cytoplasm|Texture_SumAverage_Vinculin_5_0
0_256,Cytoplasm|Texture_SumAverage_Vinculin_5_03_256,Cytoplasm|Texture_SumAverage_Vinculin_5_01_256,Cy
toplasm|Texture_SumAverage_DNA_20_01_256,Cytoplasm|Texture_SumAverage_DNA_20_00_256,Cytoplasm|Textu
re_SumAverage_DNA_20_02_256,Cytoplasm|Texture_SumAverage_DNA_20_03_256,Cytoplasm|Texture_SumAvera
ge_DNA_10_01_256,Cytoplasm|Texture_SumAverage_DNA_10_02_256,Cytoplasm|Texture_SumAverage_DNA_10_
00_256,Cytoplasm|Texture_SumAverage_DNA_10_03_256,Cytoplasm|Texture_SumAverage_DNA_5_02_256,Cytopl
asm|Texture_SumAverage_DNA_5_01_256,Cytoplasm|Texture_SumAverage_DNA_5_00_256,Cytoplasm|Texture_Su
mAverage_DNA_5_03_256,Cytoplasm|Texture_SumAverage_DNA_3_01_256,Cytoplasm|Texture_SumAverage_DN
A_3_03_256,Cytoplasm|Texture_SumAverage_DNA_3_02_256,Cytoplasm|Texture_SumAverage_DNA_3_00_256,Cy
toplasm|Texture_SumAverage_Phalloidin_5_03_256,Cytoplasm|Texture_SumAverage_Phalloidin_5_00_256,Cytoplas
m|Texture_SumAverage_Phalloidin_5_02_256,Cytoplasm|Texture_SumAverage_Phalloidin_5_01_256,Cytoplasm|Text
ure_SumAverage_Phalloidin_3_00_256,Cytoplasm|Texture_SumAverage_Phalloidin_3_03_256,Cytoplasm|Texture_Su
mAverage_Phalloidin_3_02_256,Cytoplasm|Texture_SumAverage_Phalloidin_3_01_256,Cytoplasm|Texture_SumAver
age_Phalloidin_10_01_256,Cytoplasm|Texture_SumAverage_Phalloidin_10_03_256,Cytoplasm|Texture_SumAverage
_Phalloidin_10_02_256,Cytoplasm|Texture_SumAverage_Phalloidin_10_00_256,Cytoplasm|Texture_SumAverage_Ph
alloidin_20_02_256,Cytoplasm|Texture_SumAverage_Phalloidin_20_01_256,Cytoplasm|Texture_SumAverage_Phalloi
din_20_00_256,Cytoplasm|Texture_SumAverage_Phalloidin_20_03_256,Cytoplasm|Texture_DifferenceEntropy_DNA
_5_03_256,Cytoplasm|Texture_DifferenceEntropy_DNA_5_01_256,Cytoplasm|Texture_DifferenceEntropy_DNA_5_0
2_256,Cytoplasm|Texture_DifferenceEntropy_DNA_5_00_256,Cytoplasm|Texture_DifferenceEntropy_DNA_10_01_2
56,Cytoplasm|Texture_DifferenceEntropy_DNA_10_02_256,Cytoplasm|Texture_DifferenceEntropy_DNA_10_00_256
,Cytoplasm|Texture_DifferenceEntropy_DNA_10_03_256,Cytoplasm|Texture_DifferenceEntropy_DNA_20_01_256,C
ytoplasm|Texture_DifferenceEntropy_DNA_20_03_256,Cytoplasm|Texture_DifferenceEntropy_DNA_20_02_256,Cyt
oplasm|Texture_DifferenceEntropy_DNA_20_00_256,Cytoplasm|Texture_DifferenceEntropy_DNA_3_03_256,Cytopl
asm|Texture_DifferenceEntropy_DNA_3_02_256,Cytoplasm|Texture_DifferenceEntropy_DNA_3_01_256,Cytoplasm|
Texture_DifferenceEntropy_DNA_3_00_256,Cytoplasm|Texture_DifferenceEntropy_Vinculin_20_02_256,Cytoplasm|
Texture_DifferenceEntropy_Vinculin_20_00_256,Cytoplasm|Texture_DifferenceEntropy_Vinculin_20_01_256,Cytopl
asm|Texture_DifferenceEntropy_Vinculin_20_03_256,Cytoplasm|Texture_DifferenceEntropy_Vinculin_3_01_256,Cyt
oplasm|Texture_DifferenceEntropy_Vinculin_3_00_256,Cytoplasm|Texture_DifferenceEntropy_Vinculin_3_03_256,C
ytoplasm|Texture_DifferenceEntropy_Vinculin_3_02_256,Cytoplasm|Texture_DifferenceEntropy_Vinculin_5_03_256,
Cytoplasm|Texture_DifferenceEntropy_Vinculin_5_01_256,Cytoplasm|Texture_DifferenceEntropy_Vinculin_5_00_25
6,Cytoplasm|Texture_DifferenceEntropy_Vinculin_5_02_256,Cytoplasm|Texture_DifferenceEntropy_Vinculin_10_03_

256,Cytoplasm|Texture_DifferenceEntropy_Vinculin_10_00_256,Cytoplasm|Texture_DifferenceEntropy_Vinculin_10_
02_256,Cytoplasm|Texture_DifferenceEntropy_Vinculin_10_01_256,Cytoplasm|Texture_DifferenceEntropy_WGA_3_
02_256,Cytoplasm|Texture_DifferenceEntropy_WGA_3_01_256,Cytoplasm|Texture_DifferenceEntropy_WGA_3_03_
256,Cytoplasm|Texture_DifferenceEntropy_WGA_3_00_256,Cytoplasm|Texture_DifferenceEntropy_WGA_10_01_25
6,Cytoplasm|Texture_DifferenceEntropy_WGA_10_00_256,Cytoplasm|Texture_DifferenceEntropy_WGA_10_03_256,
Cytoplasm|Texture_DifferenceEntropy_WGA_10_02_256,Cytoplasm|Texture_DifferenceEntropy_WGA_20_01_256,C
ytoplasm|Texture_DifferenceEntropy_WGA_20_00_256,Cytoplasm|Texture_DifferenceEntropy_WGA_20_03_256,Cy
toplasm|Texture_DifferenceEntropy_WGA_20_02_256,Cytoplasm|Texture_DifferenceEntropy_WGA_5_00_256,Cyto
plasm|Texture_DifferenceEntropy_WGA_5_01_256,Cytoplasm|Texture_DifferenceEntropy_WGA_5_03_256,Cytoplas
m|Texture_DifferenceEntropy_WGA_5_02_256,Cytoplasm|Texture_DifferenceEntropy_Phalloidin_10_02_256,Cytopl
asm|Texture_DifferenceEntropy_Phalloidin_10_00_256,Cytoplasm|Texture_DifferenceEntropy_Phalloidin_10_03_256,
Cytoplasm|Texture_DifferenceEntropy_Phalloidin_10_01_256,Cytoplasm|Texture_DifferenceEntropy_Phalloidin_3_03
_256,Cytoplasm|Texture_DifferenceEntropy_Phalloidin_3_02_256,Cytoplasm|Texture_DifferenceEntropy_Phalloidin_
3_01_256,Cytoplasm|Texture_DifferenceEntropy_Phalloidin_3_00_256,Cytoplasm|Texture_DifferenceEntropy_Phalloi
din_5_01_256,Cytoplasm|Texture_DifferenceEntropy_Phalloidin_5_03_256,Cytoplasm|Texture_DifferenceEntropy_Ph
alloidin_5_02_256,Cytoplasm|Texture_DifferenceEntropy_Phalloidin_5_00_256,Cytoplasm|Texture_DifferenceEntrop
y_Phalloidin_20_01_256,Cytoplasm|Texture_DifferenceEntropy_Phalloidin_20_03_256,Cytoplasm|Texture_Difference
Entropy_Phalloidin_20_00_256,Cytoplasm|Texture_DifferenceEntropy_Phalloidin_20_02_256,Cytoplasm|Texture_Cor
relation_Vinculin_20_01_256,Cytoplasm|Texture_Correlation_Vinculin_20_03_256,Cytoplasm|Texture_Correlation_V
inculin_20_02_256,Cytoplasm|Texture_Correlation_Vinculin_20_00_256,Cytoplasm|Texture_Correlation_Vinculin_3_
00_256,Cytoplasm|Texture_Correlation_Vinculin_3_01_256,Cytoplasm|Texture_Correlation_Vinculin_3_02_256,Cyto
plasm|Texture_Correlation_Vinculin_3_03_256,Cytoplasm|Texture_Correlation_Vinculin_10_01_256,Cytoplasm|Text
ure_Correlation_Vinculin_10_02_256,Cytoplasm|Texture_Correlation_Vinculin_10_03_256,Cytoplasm|Texture_Correl
ation_Vinculin_10_00_256,Cytoplasm|Texture_Correlation_Vinculin_5_03_256,Cytoplasm|Texture_Correlation_Vinc
ulin_5_00_256,Cytoplasm|Texture_Correlation_Vinculin_5_01_256,Cytoplasm|Texture_Correlation_Vinculin_5_02_2
56,Cytoplasm|Texture_Correlation_WGA_10_03_256,Cytoplasm|Texture_Correlation_WGA_10_00_256,Cytoplasm|T
exture_Correlation_WGA_10_01_256,Cytoplasm|Texture_Correlation_WGA_10_02_256,Cytoplasm|Texture_Correlati
on_WGA_3_02_256,Cytoplasm|Texture_Correlation_WGA_3_01_256,Cytoplasm|Texture_Correlation_WGA_3_03_2
56,Cytoplasm|Texture_Correlation_WGA_3_00_256,Cytoplasm|Texture_Correlation_WGA_5_00_256,Cytoplasm|Text
ure_Correlation_WGA_5_01_256,Cytoplasm|Texture_Correlation_WGA_5_02_256,Cytoplasm|Texture_Correlation_
WGA_5_03_256,Cytoplasm|Texture_Correlation_WGA_20_02_256,Cytoplasm|Texture_Correlation_WGA_20_01_25
6,Cytoplasm|Texture_Correlation_WGA_20_00_256,Cytoplasm|Texture_Correlation_WGA_20_03_256,Cytoplasm|Te
xture_Correlation_DNA_20_02_256,Cytoplasm|Texture_Correlation_DNA_20_00_256,Cytoplasm|Texture_Correlatio
n_DNA_20_03_256,Cytoplasm|Texture_Correlation_DNA_20_01_256,Cytoplasm|Texture_Correlation_DNA_5_01_2
56,Cytoplasm|Texture_Correlation_DNA_5_03_256,Cytoplasm|Texture_Correlation_DNA_5_02_256,Cytoplasm|Text
ure_Correlation_DNA_5_00_256,Cytoplasm|Texture_Correlation_DNA_3_03_256,Cytoplasm|Texture_Correlation_D
NA_3_02_256,Cytoplasm|Texture_Correlation_DNA_3_00_256,Cytoplasm|Texture_Correlation_DNA_3_01_256,Cyt
oplasm|Texture_Correlation_DNA_10_00_256,Cytoplasm|Texture_Correlation_DNA_10_01_256,Cytoplasm|Texture_
Correlation_DNA_10_02_256,Cytoplasm|Texture_Correlation_DNA_10_03_256,Cytoplasm|Texture_Correlation_Phal
loidin_3_00_256,Cytoplasm|Texture_Correlation_Phalloidin_3_02_256,Cytoplasm|Texture_Correlation_Phalloidin_3_
03_256,Cytoplasm|Texture_Correlation_Phalloidin_3_01_256,Cytoplasm|Texture_Correlation_Phalloidin_10_03_256,
Cytoplasm|Texture_Correlation_Phalloidin_10_01_256,Cytoplasm|Texture_Correlation_Phalloidin_10_00_256,Cytopla
sm|Texture_Correlation_Phalloidin_10_02_256,Cytoplasm|Texture_Correlation_Phalloidin_20_02_256,Cytoplasm|Tex
ture_Correlation_Phalloidin_20_01_256,Cytoplasm|Texture_Correlation_Phalloidin_20_00_256,Cytoplasm|Texture_C
orrelation_Phalloidin_20_03_256,Cytoplasm|Texture_Correlation_Phalloidin_5_02_256,Cytoplasm|Texture_Correlatio
n_Phalloidin_5_00_256,Cytoplasm|Texture_Correlation_Phalloidin_5_03_256,Cytoplasm|Texture_Correlation_Phalloi
din_5_01_256,Cytoplasm|Texture_Entropy_Phalloidin_20_01_256,Cytoplasm|Texture_Entropy_Phalloidin_20_03_256
,Cytoplasm|Texture_Entropy_Phalloidin_20_00_256,Cytoplasm|Texture_Entropy_Phalloidin_20_02_256,Cytoplasm|Te
xture_Entropy_Phalloidin_10_02_256,Cytoplasm|Texture_Entropy_Phalloidin_10_01_256,Cytoplasm|Texture_Entropy
_Phalloidin_10_03_256,Cytoplasm|Texture_Entropy_Phalloidin_10_00_256,Cytoplasm|Texture_Entropy_Phalloidin_5
_00_256,Cytoplasm|Texture_Entropy_Phalloidin_5_02_256,Cytoplasm|Texture_Entropy_Phalloidin_5_03_256,Cytopl
asm|Texture_Entropy_Phalloidin_5_01_256,Cytoplasm|Texture_Entropy_Phalloidin_3_03_256,Cytoplasm|Texture_Ent
ropy_Phalloidin_3_00_256,Cytoplasm|Texture_Entropy_Phalloidin_3_02_256,Cytoplasm|Texture_Entropy_Phalloidin
_3_01_256,Cytoplasm|Texture_Entropy_DNA_5_00_256,Cytoplasm|Texture_Entropy_DNA_5_03_256,Cytoplasm|Te

xture_Entropy_DNA_5_02_256,Cytoplasm|Texture_Entropy_DNA_5_01_256,Cytoplasm|Texture_Entropy_DNA_20_01_256,Cytoplasm|Texture_Entropy_DNA_20_02_256,Cytoplasm|Texture_Entropy_DNA_20_00_256,Cytoplasm|Texture_Entropy_DNA_20_03_256,Cytoplasm|Texture_Entropy_DNA_3_03_256,Cytoplasm|Texture_Entropy_DNA_3_01_256,Cytoplasm|Texture_Entropy_DNA_3_02_256,Cytoplasm|Texture_Entropy_DNA_3_00_256,Cytoplasm|Texture_Entropy_DNA_10_03_256,Cytoplasm|Texture_Entropy_DNA_10_02_256,Cytoplasm|Texture_Entropy_DNA_10_01_256,Cytoplasm|Texture_Entropy_DNA_10_00_256,Cytoplasm|Texture_Entropy_Vinculin_10_02_256,Cytoplasm|Texture_Entropy_Vinculin_10_03_256,Cytoplasm|Texture_Entropy_Vinculin_10_01_256,Cytoplasm|Texture_Entropy_Vinculin_10_00_256,Cytoplasm|Texture_Entropy_Vinculin_20_01_256,Cytoplasm|Texture_Entropy_Vinculin_20_00_256,Cytoplasm|Texture_Entropy_Vinculin_20_02_256,Cytoplasm|Texture_Entropy_Vinculin_20_03_256,Cytoplasm|Texture_Entropy_Vinculin_3_01_256,Cytoplasm|Texture_Entropy_Vinculin_3_02_256,Cytoplasm|Texture_Entropy_Vinculin_3_00_256,Cytoplasm|Texture_Entropy_Vinculin_3_03_256,Cytoplasm|Texture_Entropy_Vinculin_5_03_256,Cytoplasm|Texture_Entropy_Vinculin_5_00_256,Cytoplasm|Texture_Entropy_Vinculin_5_01_256,Cytoplasm|Texture_Entropy_Vinculin_5_02_256,Cytoplasm|Texture_Entropy_WGA_20_00_256,Cytoplasm|Texture_Entropy_WGA_20_02_256,Cytoplasm|Texture_Entropy_WGA_20_03_256,Cytoplasm|Texture_Entropy_WGA_20_01_256,Cytoplasm|Texture_Entropy_WGA_5_00_256,Cytoplasm|Texture_Entropy_WGA_5_01_256,Cytoplasm|Texture_Entropy_WGA_5_03_256,Cytoplasm|Texture_Entropy_WGA_5_02_256,Cytoplasm|Texture_Entropy_WGA_3_01_256,Cytoplasm|Texture_Entropy_WGA_3_03_256,Cytoplasm|Texture_Entropy_WGA_3_02_256,Cytoplasm|Texture_Entropy_WGA_3_00_256,Cytoplasm|Texture_Entropy_WGA_10_03_256,Cytoplasm|Texture_Entropy_WGA_10_02_256,Cytoplasm|Texture_Entropy_WGA_10_01_256,Cytoplasm|Texture_Entropy_WGA_10_00_256,Cytoplasm|Texture_AngularSecondMoment_Phalloidin_20_00_256,Cytoplasm|Texture_AngularSecondMoment_Phalloidin_20_03_256,Cytoplasm|Texture_AngularSecondMoment_Phalloidin_20_01_256,Cytoplasm|Texture_AngularSecondMoment_Phalloidin_20_02_256,Cytoplasm|Texture_AngularSecondMoment_Phalloidin_3_01_256,Cytoplasm|Texture_AngularSecondMoment_Phalloidin_3_00_256,Cytoplasm|Texture_AngularSecondMoment_Phalloidin_3_03_256,Cytoplasm|Texture_AngularSecondMoment_Phalloidin_3_02_256,Cytoplasm|Texture_AngularSecondMoment_Phalloidin_10_01_256,Cytoplasm|Texture_AngularSecondMoment_Phalloidin_10_02_256,Cytoplasm|Texture_AngularSecondMoment_Phalloidin_10_00_256,Cytoplasm|Texture_AngularSecondMoment_Phalloidin_10_03_256,Cytoplasm|Texture_AngularSecondMoment_Phalloidin_5_02_256,Cytoplasm|Texture_AngularSecondMoment_Phalloidin_5_03_256,Cytoplasm|Texture_AngularSecondMoment_Phalloidin_5_00_256,Cytoplasm|Texture_AngularSecondMoment_Phalloidin_5_01_256,Cytoplasm|Texture_AngularSecondMoment_DNA_3_03_256,Cytoplasm|Texture_AngularSecondMoment_DNA_3_02_256,Cytoplasm|Texture_AngularSecondMoment_DNA_3_01_256,Cytoplasm|Texture_AngularSecondMoment_DNA_3_00_256,Cytoplasm|Texture_AngularSecondMoment_DNA_10_03_256,Cytoplasm|Texture_AngularSecondMoment_DNA_10_00_256,Cytoplasm|Texture_AngularSecondMoment_DNA_10_02_256,Cytoplasm|Texture_AngularSecondMoment_DNA_10_01_256,Cytoplasm|Texture_AngularSecondMoment_DNA_20_00_256,Cytoplasm|Texture_AngularSecondMoment_DNA_20_03_256,Cytoplasm|Texture_AngularSecondMoment_DNA_20_02_256,Cytoplasm|Texture_AngularSecondMoment_DNA_20_01_256,Cytoplasm|Texture_AngularSecondMoment_DNA_5_01_256,Cytoplasm|Texture_AngularSecondMoment_DNA_5_02_256,Cytoplasm|Texture_AngularSecondMoment_DNA_5_03_256,Cytoplasm|Texture_AngularSecondMoment_DNA_5_00_256,Cytoplasm|Texture_AngularSecondMoment_WGA_20_00_256,Cytoplasm|Texture_AngularSecondMoment_WGA_20_01_256,Cytoplasm|Texture_AngularSecondMoment_WGA_20_02_256,Cytoplasm|Texture_AngularSecondMoment_WGA_20_03_256,Cytoplasm|Texture_AngularSecondMoment_WGA_3_02_256,Cytoplasm|Texture_AngularSecondMoment_WGA_3_00_256,Cytoplasm|Texture_AngularSecondMoment_WGA_3_03_256,Cytoplasm|Texture_AngularSecondMoment_WGA_3_01_256,Cytoplasm|Texture_AngularSecondMoment_WGA_10_03_256,Cytoplasm|Texture_AngularSecondMoment_WGA_10_00_256,Cytoplasm|Texture_AngularSecondMoment_WGA_10_02_256,Cytoplasm|Texture_AngularSecondMoment_WGA_10_01_256,Cytoplasm|Texture_AngularSecondMoment_WGA_5_00_256,Cytoplasm|Texture_AngularSecondMoment_WGA_5_01_256,Cytoplasm|Texture_AngularSecondMoment_WGA_5_02_256,Cytoplasm|Texture_AngularSecondMoment_WGA_5_03_256,Cytoplasm|Texture_AngularSecondMoment_Vinculin_3_00_256,Cytoplasm|Texture_AngularSecondMoment_Vinculin_3_03_256,Cytoplasm|Texture_AngularSecondMoment_Vinculin_3_02_256,Cytoplasm|Texture_AngularSecondMoment_Vinculin_3_01_256,Cytoplasm|Texture_AngularSecondMoment_Vinculin_5_00_256,Cytoplasm|Texture_AngularSecondMoment_Vinculin_5_02_256,Cytoplasm|Texture_AngularSecondMoment_Vinculin_5_01_256,Cytoplasm|Texture_AngularSecondMoment_Vinculin_5_03_256,Cytoplasm|Texture_AngularSecondMoment_Vinculin_20_02_256,Cytoplasm|Texture_AngularSecondMoment_Vinculin_20_01_256,Cytoplasm|Texture_AngularSecondMoment_Vinculin_20_03_256,Cytoplasm|Texture_AngularSecondMoment_Vinculin_20_00_256,Cytoplasm|Texture_AngularSecondMoment_Vinculin_10_03_256,Cytoplasm|Texture_AngularSecondMoment_Vinculin_10_00_256,Cytoplasm|Texture_AngularSecondMoment_Vinculin_10_02_256,Cytoplasm|Texture_AngularSecondMoment_Vinculin_10_01_256,Cytoplasm|Texture_InfoMeas2_DNA_10_0

2_256,Cytoplasm|Texture_InfoMeas2_DNA_10_01_256,Cytoplasm|Texture_InfoMeas2_DNA_10_03_256,Cytoplasm|
Texture_InfoMeas2_DNA_10_00_256,Cytoplasm|Texture_InfoMeas2_DNA_5_01_256,Cytoplasm|Texture_InfoMeas2
_DNA_5_02_256,Cytoplasm|Texture_InfoMeas2_DNA_5_03_256,Cytoplasm|Texture_InfoMeas2_DNA_5_00_256,C
ytoplasm|Texture_InfoMeas2_DNA_20_01_256,Cytoplasm|Texture_InfoMeas2_DNA_20_03_256,Cytoplasm|Texture_
InfoMeas2_DNA_20_02_256,Cytoplasm|Texture_InfoMeas2_DNA_20_00_256,Cytoplasm|Texture_InfoMeas2_DNA_
3_00_256,Cytoplasm|Texture_InfoMeas2_DNA_3_03_256,Cytoplasm|Texture_InfoMeas2_DNA_3_01_256,Cytoplas
m|Texture_InfoMeas2_DNA_3_02_256,Cytoplasm|Texture_InfoMeas2_Vinculin_20_02_256,Cytoplasm|Texture_Info
Meas2_Vinculin_20_03_256,Cytoplasm|Texture_InfoMeas2_Vinculin_20_01_256,Cytoplasm|Texture_InfoMeas2_Vin
culin_20_00_256,Cytoplasm|Texture_InfoMeas2_Vinculin_10_00_256,Cytoplasm|Texture_InfoMeas2_Vinculin_10_0
3_256,Cytoplasm|Texture_InfoMeas2_Vinculin_10_02_256,Cytoplasm|Texture_InfoMeas2_Vinculin_10_01_256,Cyto
plasm|Texture_InfoMeas2_Vinculin_3_00_256,Cytoplasm|Texture_InfoMeas2_Vinculin_3_03_256,Cytoplasm|Texture
_InfoMeas2_Vinculin_3_01_256,Cytoplasm|Texture_InfoMeas2_Vinculin_3_02_256,Cytoplasm|Texture_InfoMeas2_
Vinculin_5_02_256,Cytoplasm|Texture_InfoMeas2_Vinculin_5_00_256,Cytoplasm|Texture_InfoMeas2_Vinculin_5_0
1_256,Cytoplasm|Texture_InfoMeas2_Vinculin_5_03_256,Cytoplasm|Texture_InfoMeas2_WGA_10_02_256,Cytoplas
m|Texture_InfoMeas2_WGA_10_00_256,Cytoplasm|Texture_InfoMeas2_WGA_10_01_256,Cytoplasm|Texture_Info
Meas2_WGA_10_03_256,Cytoplasm|Texture_InfoMeas2_WGA_20_03_256,Cytoplasm|Texture_InfoMeas2_WGA_20
_01_256,Cytoplasm|Texture_InfoMeas2_WGA_20_00_256,Cytoplasm|Texture_InfoMeas2_WGA_20_02_256,Cytopla
sm|Texture_InfoMeas2_WGA_3_02_256,Cytoplasm|Texture_InfoMeas2_WGA_3_01_256,Cytoplasm|Texture_InfoMe
as2_WGA_3_00_256,Cytoplasm|Texture_InfoMeas2_WGA_3_03_256,Cytoplasm|Texture_InfoMeas2_WGA_5_03_2
56,Cytoplasm|Texture_InfoMeas2_WGA_5_00_256,Cytoplasm|Texture_InfoMeas2_WGA_5_02_256,Cytoplasm|Text
ure_InfoMeas2_WGA_5_01_256,Cytoplasm|Texture_InfoMeas2_Phalloidin_10_02_256,Cytoplasm|Texture_InfoMeas
2_Phalloidin_10_00_256,Cytoplasm|Texture_InfoMeas2_Phalloidin_10_01_256,Cytoplasm|Texture_InfoMeas2_Phallo
idin_10_03_256,Cytoplasm|Texture_InfoMeas2_Phalloidin_20_00_256,Cytoplasm|Texture_InfoMeas2_Phalloidin_20_
02_256,Cytoplasm|Texture_InfoMeas2_Phalloidin_20_03_256,Cytoplasm|Texture_InfoMeas2_Phalloidin_20_01_256,
Cytoplasm|Texture_InfoMeas2_Phalloidin_5_01_256,Cytoplasm|Texture_InfoMeas2_Phalloidin_5_03_256,Cytoplasm|
Texture_InfoMeas2_Phalloidin_5_00_256,Cytoplasm|Texture_InfoMeas2_Phalloidin_5_02_256,Cytoplasm|Texture_In
foMeas2_Phalloidin_3_00_256,Cytoplasm|Texture_InfoMeas2_Phalloidin_3_01_256,Cytoplasm|Texture_InfoMeas2_P
halloidin_3_03_256,Cytoplasm|Texture_InfoMeas2_Phalloidin_3_02_256,Cytoplasm|Texture_InfoMeas1_Phalloidin_5
_00_256,Cytoplasm|Texture_InfoMeas1_Phalloidin_5_03_256,Cytoplasm|Texture_InfoMeas1_Phalloidin_5_02_256,C
ytoplasm|Texture_InfoMeas1_Phalloidin_5_01_256,Cytoplasm|Texture_InfoMeas1_Phalloidin_10_01_256,Cytoplasm|
Texture_InfoMeas1_Phalloidin_10_02_256,Cytoplasm|Texture_InfoMeas1_Phalloidin_10_00_256,Cytoplasm|Texture_
InfoMeas1_Phalloidin_10_03_256,Cytoplasm|Texture_InfoMeas1_Phalloidin_3_03_256,Cytoplasm|Texture_InfoMeas
1_Phalloidin_3_02_256,Cytoplasm|Texture_InfoMeas1_Phalloidin_3_00_256,Cytoplasm|Texture_InfoMeas1_Phalloidi
n_3_01_256,Cytoplasm|Texture_InfoMeas1_Phalloidin_20_03_256,Cytoplasm|Texture_InfoMeas1_Phalloidin_20_02_
256,Cytoplasm|Texture_InfoMeas1_Phalloidin_20_01_256,Cytoplasm|Texture_InfoMeas1_Phalloidin_20_00_256,Cyt
oplasm|Texture_InfoMeas1_Vinculin_5_03_256,Cytoplasm|Texture_InfoMeas1_Vinculin_5_02_256,Cytoplasm|Textur
e_InfoMeas1_Vinculin_5_01_256,Cytoplasm|Texture_InfoMeas1_Vinculin_5_00_256,Cytoplasm|Texture_InfoMeas1_
Vinculin_10_03_256,Cytoplasm|Texture_InfoMeas1_Vinculin_10_00_256,Cytoplasm|Texture_InfoMeas1_Vinculin_1
0_02_256,Cytoplasm|Texture_InfoMeas1_Vinculin_10_01_256,Cytoplasm|Texture_InfoMeas1_Vinculin_3_02_256,C
ytoplasm|Texture_InfoMeas1_Vinculin_3_00_256,Cytoplasm|Texture_InfoMeas1_Vinculin_3_03_256,Cytoplasm|Text
ure_InfoMeas1_Vinculin_3_01_256,Cytoplasm|Texture_InfoMeas1_Vinculin_20_00_256,Cytoplasm|Texture_InfoMea
s1_Vinculin_20_03_256,Cytoplasm|Texture_InfoMeas1_Vinculin_20_01_256,Cytoplasm|Texture_InfoMeas1_Vinculi
n_20_02_256,Cytoplasm|Texture_InfoMeas1_DNA_10_03_256,Cytoplasm|Texture_InfoMeas1_DNA_10_00_256,Cyt
oplasm|Texture_InfoMeas1_DNA_10_02_256,Cytoplasm|Texture_InfoMeas1_DNA_10_01_256,Cytoplasm|Texture_I
nfoMeas1_DNA_3_02_256,Cytoplasm|Texture_InfoMeas1_DNA_3_03_256,Cytoplasm|Texture_InfoMeas1_DNA_3_
00_256,Cytoplasm|Texture_InfoMeas1_DNA_3_01_256,Cytoplasm|Texture_InfoMeas1_DNA_5_00_256,Cytoplasm|T
exture_InfoMeas1_DNA_5_02_256,Cytoplasm|Texture_InfoMeas1_DNA_5_03_256,Cytoplasm|Texture_InfoMeas1_
DNA_5_01_256,Cytoplasm|Texture_InfoMeas1_DNA_20_02_256,Cytoplasm|Texture_InfoMeas1_DNA_20_01_256,
Cytoplasm|Texture_InfoMeas1_DNA_20_00_256,Cytoplasm|Texture_InfoMeas1_DNA_20_03_256,Cytoplasm|Textur
e_InfoMeas1_WGA_20_00_256,Cytoplasm|Texture_InfoMeas1_WGA_20_02_256,Cytoplasm|Texture_InfoMeas1_W
GA_20_01_256,Cytoplasm|Texture_InfoMeas1_WGA_20_03_256,Cytoplasm|Texture_InfoMeas1_WGA_10_00_256,
Cytoplasm|Texture_InfoMeas1_WGA_10_01_256,Cytoplasm|Texture_InfoMeas1_WGA_10_02_256,Cytoplasm|Textu
re_InfoMeas1_WGA_10_03_256,Cytoplasm|Texture_InfoMeas1_WGA_5_01_256,Cytoplasm|Texture_InfoMeas1_W
GA_5_00_256,Cytoplasm|Texture_InfoMeas1_WGA_5_03_256,Cytoplasm|Texture_InfoMeas1_WGA_5_02_256,Cyt

oplasm|Texture_InfoMeas1_WGA_3_02_256,Cytoplasm|Texture_InfoMeas1_WGA_3_00_256,Cytoplasm|Texture_Inf
oMeas1_WGA_3_03_256,Cytoplasm|Texture_InfoMeas1_WGA_3_01_256,Cytoplasm|Texture_SumVariance_Phalloi
din_3_02_256,Cytoplasm|Texture_SumVariance_Phalloidin_3_00_256,Cytoplasm|Texture_SumVariance_Phalloidin_3
_03_256,Cytoplasm|Texture_SumVariance_Phalloidin_3_01_256,Cytoplasm|Texture_SumVariance_Phalloidin_5_03_
256,Cytoplasm|Texture_SumVariance_Phalloidin_5_01_256,Cytoplasm|Texture_SumVariance_Phalloidin_5_00_256,C
ytoplasm|Texture_SumVariance_Phalloidin_5_02_256,Cytoplasm|Texture_SumVariance_Phalloidin_20_03_256,Cytop
lasm|Texture_SumVariance_Phalloidin_20_00_256,Cytoplasm|Texture_SumVariance_Phalloidin_20_01_256,Cytoplas
m|Texture_SumVariance_Phalloidin_20_02_256,Cytoplasm|Texture_SumVariance_Phalloidin_10_03_256,Cytoplasm|
Texture_SumVariance_Phalloidin_10_00_256,Cytoplasm|Texture_SumVariance_Phalloidin_10_02_256,Cytoplasm|Te
xture_SumVariance_Phalloidin_10_01_256,Cytoplasm|Texture_SumVariance_DNA_10_00_256,Cytoplasm|Texture_S
umVariance_DNA_10_02_256,Cytoplasm|Texture_SumVariance_DNA_10_01_256,Cytoplasm|Texture_SumVariance
_DNA_10_03_256,Cytoplasm|Texture_SumVariance_DNA_5_00_256,Cytoplasm|Texture_SumVariance_DNA_5_01_
256,Cytoplasm|Texture_SumVariance_DNA_5_03_256,Cytoplasm|Texture_SumVariance_DNA_5_02_256,Cytoplasm|
Texture_SumVariance_DNA_20_03_256,Cytoplasm|Texture_SumVariance_DNA_20_02_256,Cytoplasm|Texture_Su
mVariance_DNA_20_00_256,Cytoplasm|Texture_SumVariance_DNA_20_01_256,Cytoplasm|Texture_SumVariance_
DNA_3_03_256,Cytoplasm|Texture_SumVariance_DNA_3_02_256,Cytoplasm|Texture_SumVariance_DNA_3_00_25
6,Cytoplasm|Texture_SumVariance_DNA_3_01_256,Cytoplasm|Texture_SumVariance_WGA_10_03_256,Cytoplasm|
Texture_SumVariance_WGA_10_01_256,Cytoplasm|Texture_SumVariance_WGA_10_00_256,Cytoplasm|Texture_Su
mVariance_WGA_10_02_256,Cytoplasm|Texture_SumVariance_WGA_5_00_256,Cytoplasm|Texture_SumVariance_
WGA_5_03_256,Cytoplasm|Texture_SumVariance_WGA_5_02_256,Cytoplasm|Texture_SumVariance_WGA_5_01_
256,Cytoplasm|Texture_SumVariance_WGA_3_03_256,Cytoplasm|Texture_SumVariance_WGA_3_01_256,Cytoplas
m|Texture_SumVariance_WGA_3_02_256,Cytoplasm|Texture_SumVariance_WGA_3_00_256,Cytoplasm|Texture_Su
mVariance_WGA_20_02_256,Cytoplasm|Texture_SumVariance_WGA_20_00_256,Cytoplasm|Texture_SumVariance
_WGA_20_03_256,Cytoplasm|Texture_SumVariance_WGA_20_01_256,Cytoplasm|Texture_SumVariance_Vinculin_
20_01_256,Cytoplasm|Texture_SumVariance_Vinculin_20_03_256,Cytoplasm|Texture_SumVariance_Vinculin_20_00
_256,Cytoplasm|Texture_SumVariance_Vinculin_20_02_256,Cytoplasm|Texture_SumVariance_Vinculin_10_03_256,
Cytoplasm|Texture_SumVariance_Vinculin_10_01_256,Cytoplasm|Texture_SumVariance_Vinculin_10_02_256,Cytop
lasm|Texture_SumVariance_Vinculin_10_00_256,Cytoplasm|Texture_SumVariance_Vinculin_5_03_256,Cytoplasm|T
exture_SumVariance_Vinculin_5_02_256,Cytoplasm|Texture_SumVariance_Vinculin_5_01_256,Cytoplasm|Texture_S
umVariance_Vinculin_5_00_256,Cytoplasm|Texture_SumVariance_Vinculin_3_03_256,Cytoplasm|Texture_SumVaria
nce_Vinculin_3_00_256,Cytoplasm|Texture_SumVariance_Vinculin_3_01_256,Cytoplasm|Texture_SumVariance_Vin
culin_3_02_256,Cytoplasm|Texture_Variance_Phalloidin_20_02_256,Cytoplasm|Texture_Variance_Phalloidin_20_00_
256,Cytoplasm|Texture_Variance_Phalloidin_20_03_256,Cytoplasm|Texture_Variance_Phalloidin_20_01_256,Cytopla
sm|Texture_Variance_Phalloidin_10_01_256,Cytoplasm|Texture_Variance_Phalloidin_10_03_256,Cytoplasm|Texture_
Variance_Phalloidin_10_00_256,Cytoplasm|Texture_Variance_Phalloidin_10_02_256,Cytoplasm|Texture_Variance_P
halloidin_3_01_256,Cytoplasm|Texture_Variance_Phalloidin_3_00_256,Cytoplasm|Texture_Variance_Phalloidin_3_02
_256,Cytoplasm|Texture_Variance_Phalloidin_3_03_256,Cytoplasm|Texture_Variance_Phalloidin_5_03_256,Cytoplas
m|Texture_Variance_Phalloidin_5_00_256,Cytoplasm|Texture_Variance_Phalloidin_5_02_256,Cytoplasm|Texture_Var
iance_Phalloidin_5_01_256,Cytoplasm|Texture_Variance_WGA_10_02_256,Cytoplasm|Texture_Variance_WGA_10_
01_256,Cytoplasm|Texture_Variance_WGA_10_03_256,Cytoplasm|Texture_Variance_WGA_10_00_256,Cytoplasm|T
exture_Variance_WGA_5_02_256,Cytoplasm|Texture_Variance_WGA_5_00_256,Cytoplasm|Texture_Variance_WGA
_5_01_256,Cytoplasm|Texture_Variance_WGA_5_03_256,Cytoplasm|Texture_Variance_WGA_3_01_256,Cytoplasm|
Texture_Variance_WGA_3_02_256,Cytoplasm|Texture_Variance_WGA_3_03_256,Cytoplasm|Texture_Variance_WG
A_3_00_256,Cytoplasm|Texture_Variance_WGA_20_02_256,Cytoplasm|Texture_Variance_WGA_20_01_256,Cytopl
asm|Texture_Variance_WGA_20_00_256,Cytoplasm|Texture_Variance_WGA_20_03_256,Cytoplasm|Texture_Varian
ce_DNA_20_02_256,Cytoplasm|Texture_Variance_DNA_20_00_256,Cytoplasm|Texture_Variance_DNA_20_01_256,
Cytoplasm|Texture_Variance_DNA_20_03_256,Cytoplasm|Texture_Variance_DNA_5_03_256,Cytoplasm|Texture_Va
riance_DNA_5_01_256,Cytoplasm|Texture_Variance_DNA_5_00_256,Cytoplasm|Texture_Variance_DNA_5_02_256,
Cytoplasm|Texture_Variance_DNA_10_01_256,Cytoplasm|Texture_Variance_DNA_10_00_256,Cytoplasm|Texture_V
ariance_DNA_10_02_256,Cytoplasm|Texture_Variance_DNA_10_03_256,Cytoplasm|Texture_Variance_DNA_3_03_
256,Cytoplasm|Texture_Variance_DNA_3_02_256,Cytoplasm|Texture_Variance_DNA_3_00_256,Cytoplasm|Texture_
Variance_DNA_3_01_256,Cytoplasm|Texture_Variance_Vinculin_10_01_256,Cytoplasm|Texture_Variance_Vinculin_
10_03_256,Cytoplasm|Texture_Variance_Vinculin_10_00_256,Cytoplasm|Texture_Variance_Vinculin_10_02_256,Cyt
oplasm|Texture_Variance_Vinculin_3_02_256,Cytoplasm|Texture_Variance_Vinculin_3_00_256,Cytoplasm|Texture_

Variance_Vinculin_3_03_256,Cytoplasm|Texture_Variance_Vinculin_3_01_256,Cytoplasm|Texture_Variance_Vinculi
n_20_01_256,Cytoplasm|Texture_Variance_Vinculin_20_03_256,Cytoplasm|Texture_Variance_Vinculin_20_02_256,
Cytoplasm|Texture_Variance_Vinculin_20_00_256,Cytoplasm|Texture_Variance_Vinculin_5_03_256,Cytoplasm|Text
ure_Variance_Vinculin_5_02_256,Cytoplasm|Texture_Variance_Vinculin_5_00_256,Cytoplasm|Texture_Variance_Vi
nculin_5_01_256,Cytoplasm|Texture_Contrast_WGA_10_01_256,Cytoplasm|Texture_Contrast_WGA_10_03_256,Cyt
oplasm|Texture_Contrast_WGA_10_00_256,Cytoplasm|Texture_Contrast_WGA_10_02_256,Cytoplasm|Texture_Cont
rast_WGA_5_03_256,Cytoplasm|Texture_Contrast_WGA_5_01_256,Cytoplasm|Texture_Contrast_WGA_5_00_256,C
ytoplasm|Texture_Contrast_WGA_5_02_256,Cytoplasm|Texture_Contrast_WGA_20_03_256,Cytoplasm|Texture_Cont
rast_WGA_20_01_256,Cytoplasm|Texture_Contrast_WGA_20_02_256,Cytoplasm|Texture_Contrast_WGA_20_00_25
6,Cytoplasm|Texture_Contrast_WGA_3_01_256,Cytoplasm|Texture_Contrast_WGA_3_03_256,Cytoplasm|Texture_C
ontrast_WGA_3_00_256,Cytoplasm|Texture_Contrast_WGA_3_02_256,Cytoplasm|Texture_Contrast_DNA_3_03_256
,Cytoplasm|Texture_Contrast_DNA_3_00_256,Cytoplasm|Texture_Contrast_DNA_3_02_256,Cytoplasm|Texture_Cont
rast_DNA_3_01_256,Cytoplasm|Texture_Contrast_DNA_5_02_256,Cytoplasm|Texture_Contrast_DNA_5_00_256,Cyt
oplasm|Texture_Contrast_DNA_5_03_256,Cytoplasm|Texture_Contrast_DNA_5_01_256,Cytoplasm|Texture_Contrast
_DNA_10_01_256,Cytoplasm|Texture_Contrast_DNA_10_03_256,Cytoplasm|Texture_Contrast_DNA_10_00_256,Cyt
oplasm|Texture_Contrast_DNA_10_02_256,Cytoplasm|Texture_Contrast_DNA_20_03_256,Cytoplasm|Texture_Contr
ast_DNA_20_00_256,Cytoplasm|Texture_Contrast_DNA_20_02_256,Cytoplasm|Texture_Contrast_DNA_20_01_256,
Cytoplasm|Texture_Contrast_Vinculin_3_03_256,Cytoplasm|Texture_Contrast_Vinculin_3_02_256,Cytoplasm|Texture
_Contrast_Vinculin_3_00_256,Cytoplasm|Texture_Contrast_Vinculin_3_01_256,Cytoplasm|Texture_Contrast_Vinculi
n_20_00_256,Cytoplasm|Texture_Contrast_Vinculin_20_01_256,Cytoplasm|Texture_Contrast_Vinculin_20_03_256,C
ytoplasm|Texture_Contrast_Vinculin_20_02_256,Cytoplasm|Texture_Contrast_Vinculin_5_00_256,Cytoplasm|Texture
_Contrast_Vinculin_5_03_256,Cytoplasm|Texture_Contrast_Vinculin_5_02_256,Cytoplasm|Texture_Contrast_Vinculi
n_5_01_256,Cytoplasm|Texture_Contrast_Vinculin_10_00_256,Cytoplasm|Texture_Contrast_Vinculin_10_02_256,Cyt
oplasm|Texture_Contrast_Vinculin_10_01_256,Cytoplasm|Texture_Contrast_Vinculin_10_03_256,Cytoplasm|Texture_
Contrast_Phalloidin_20_03_256,Cytoplasm|Texture_Contrast_Phalloidin_20_02_256,Cytoplasm|Texture_Contrast_Pha
lloidin_20_01_256,Cytoplasm|Texture_Contrast_Phalloidin_20_00_256,Cytoplasm|Texture_Contrast_Phalloidin_5_01
_256,Cytoplasm|Texture_Contrast_Phalloidin_5_03_256,Cytoplasm|Texture_Contrast_Phalloidin_5_02_256,Cytoplas
m|Texture_Contrast_Phalloidin_5_00_256,Cytoplasm|Texture_Contrast_Phalloidin_10_03_256,Cytoplasm|Texture_Co
ntrast_Phalloidin_10_01_256,Cytoplasm|Texture_Contrast_Phalloidin_10_00_256,Cytoplasm|Texture_Contrast_Phallo
idin_10_02_256,Cytoplasm|Texture_Contrast_Phalloidin_3_02_256,Cytoplasm|Texture_Contrast_Phalloidin_3_01_25
6,Cytoplasm|Texture_Contrast_Phalloidin_3_00_256,Cytoplasm|Texture_Contrast_Phalloidin_3_03_256,Cytoplasm|Te
xture_InverseDifferenceMoment_WGA_20_01_256,Cytoplasm|Texture_InverseDifferenceMoment_WGA_20_02_256,
Cytoplasm|Texture_InverseDifferenceMoment_WGA_20_00_256,Cytoplasm|Texture_InverseDifferenceMoment_WG
A_20_03_256,Cytoplasm|Texture_InverseDifferenceMoment_WGA_5_01_256,Cytoplasm|Texture_InverseDifference
Moment_WGA_5_02_256,Cytoplasm|Texture_InverseDifferenceMoment_WGA_5_00_256,Cytoplasm|Texture_Invers
eDifferenceMoment_WGA_5_03_256,Cytoplasm|Texture_InverseDifferenceMoment_WGA_3_01_256,Cytoplasm|Tex
ture_InverseDifferenceMoment_WGA_3_00_256,Cytoplasm|Texture_InverseDifferenceMoment_WGA_3_03_256,Cyt
oplasm|Texture_InverseDifferenceMoment_WGA_3_02_256,Cytoplasm|Texture_InverseDifferenceMoment_WGA_10
_03_256,Cytoplasm|Texture_InverseDifferenceMoment_WGA_10_02_256,Cytoplasm|Texture_InverseDifferenceMom
ent_WGA_10_01_256,Cytoplasm|Texture_InverseDifferenceMoment_WGA_10_00_256,Cytoplasm|Texture_InverseD
ifferenceMoment_Phalloidin_10_01_256,Cytoplasm|Texture_InverseDifferenceMoment_Phalloidin_10_02_256,Cytopl
asm|Texture_InverseDifferenceMoment_Phalloidin_10_00_256,Cytoplasm|Texture_InverseDifferenceMoment_Phalloi
din_10_03_256,Cytoplasm|Texture_InverseDifferenceMoment_Phalloidin_5_00_256,Cytoplasm|Texture_InverseDiffer
enceMoment_Phalloidin_5_01_256,Cytoplasm|Texture_InverseDifferenceMoment_Phalloidin_5_02_256,Cytoplasm|Te
xture_InverseDifferenceMoment_Phalloidin_5_03_256,Cytoplasm|Texture_InverseDifferenceMoment_Phalloidin_3_0
1_256,Cytoplasm|Texture_InverseDifferenceMoment_Phalloidin_3_02_256,Cytoplasm|Texture_InverseDifferenceMo
ment_Phalloidin_3_03_256,Cytoplasm|Texture_InverseDifferenceMoment_Phalloidin_3_00_256,Cytoplasm|Texture_I
nverseDifferenceMoment_Phalloidin_20_02_256,Cytoplasm|Texture_InverseDifferenceMoment_Phalloidin_20_01_25
6,Cytoplasm|Texture_InverseDifferenceMoment_Phalloidin_20_03_256,Cytoplasm|Texture_InverseDifferenceMoment
_Phalloidin_20_00_256,Cytoplasm|Texture_InverseDifferenceMoment_Vinculin_10_03_256,Cytoplasm|Texture_Inver
seDifferenceMoment_Vinculin_10_02_256,Cytoplasm|Texture_InverseDifferenceMoment_Vinculin_10_00_256,Cytop
lasm|Texture_InverseDifferenceMoment_Vinculin_10_01_256,Cytoplasm|Texture_InverseDifferenceMoment_Vinculin
_3_03_256,Cytoplasm|Texture_InverseDifferenceMoment_Vinculin_3_00_256,Cytoplasm|Texture_InverseDifference
Moment_Vinculin_3_01_256,Cytoplasm|Texture_InverseDifferenceMoment_Vinculin_3_02_256,Cytoplasm|Texture_I

nverseDifferenceMoment_Vinculin_20_02_256,Cytoplasm|Texture_InverseDifferenceMoment_Vinculin_20_03_256,C
ytoplasm|Texture_InverseDifferenceMoment_Vinculin_20_01_256,Cytoplasm|Texture_InverseDifferenceMoment_Vin
culin_20_00_256,Cytoplasm|Texture_InverseDifferenceMoment_Vinculin_5_03_256,Cytoplasm|Texture_InverseDiffer
enceMoment_Vinculin_5_02_256,Cytoplasm|Texture_InverseDifferenceMoment_Vinculin_5_01_256,Cytoplasm|Text
ure_InverseDifferenceMoment_Vinculin_5_00_256,Cytoplasm|Texture_InverseDifferenceMoment_DNA_10_02_256,
Cytoplasm|Texture_InverseDifferenceMoment_DNA_10_01_256,Cytoplasm|Texture_InverseDifferenceMoment_DNA
_10_03_256,Cytoplasm|Texture_InverseDifferenceMoment_DNA_10_00_256,Cytoplasm|Texture_InverseDifferenceM
oment_DNA_20_00_256,Cytoplasm|Texture_InverseDifferenceMoment_DNA_20_02_256,Cytoplasm|Texture_Inverse
DifferenceMoment_DNA_20_03_256,Cytoplasm|Texture_InverseDifferenceMoment_DNA_20_01_256,Cytoplasm|Te
xture_InverseDifferenceMoment_DNA_3_01_256,Cytoplasm|Texture_InverseDifferenceMoment_DNA_3_02_256,Cyt
oplasm|Texture_InverseDifferenceMoment_DNA_3_03_256,Cytoplasm|Texture_InverseDifferenceMoment_DNA_3_0
0_256,Cytoplasm|Texture_InverseDifferenceMoment_DNA_5_01_256,Cytoplasm|Texture_InverseDifferenceMoment_
DNA_5_02_256,Cytoplasm|Texture_InverseDifferenceMoment_DNA_5_00_256,Cytoplasm|Texture_InverseDifferenc
eMoment_DNA_5_03_256,Cytoplasm|Texture_SumEntropy_Phalloidin_5_00_256,Cytoplasm|Texture_SumEntropy_P
halloidin_5_02_256,Cytoplasm|Texture_SumEntropy_Phalloidin_5_01_256,Cytoplasm|Texture_SumEntropy_Phalloidi
n_5_03_256,Cytoplasm|Texture_SumEntropy_Phalloidin_3_02_256,Cytoplasm|Texture_SumEntropy_Phalloidin_3_01
_256,Cytoplasm|Texture_SumEntropy_Phalloidin_3_03_256,Cytoplasm|Texture_SumEntropy_Phalloidin_3_00_256,C
ytoplasm|Texture_SumEntropy_Phalloidin_10_00_256,Cytoplasm|Texture_SumEntropy_Phalloidin_10_01_256,Cytopl
asm|Texture_SumEntropy_Phalloidin_10_02_256,Cytoplasm|Texture_SumEntropy_Phalloidin_10_03_256,Cytoplasm|
Texture_SumEntropy_Phalloidin_20_03_256,Cytoplasm|Texture_SumEntropy_Phalloidin_20_01_256,Cytoplasm|Text
ure_SumEntropy_Phalloidin_20_00_256,Cytoplasm|Texture_SumEntropy_Phalloidin_20_02_256,Cytoplasm|Texture_
SumEntropy_Vinculin_10_01_256,Cytoplasm|Texture_SumEntropy_Vinculin_10_00_256,Cytoplasm|Texture_SumEnt
ropy_Vinculin_10_02_256,Cytoplasm|Texture_SumEntropy_Vinculin_10_03_256,Cytoplasm|Texture_SumEntropy_Vi
nculin_3_02_256,Cytoplasm|Texture_SumEntropy_Vinculin_3_01_256,Cytoplasm|Texture_SumEntropy_Vinculin_3_
00_256,Cytoplasm|Texture_SumEntropy_Vinculin_3_03_256,Cytoplasm|Texture_SumEntropy_Vinculin_20_03_256,
Cytoplasm|Texture_SumEntropy_Vinculin_20_02_256,Cytoplasm|Texture_SumEntropy_Vinculin_20_00_256,Cytopla
sm|Texture_SumEntropy_Vinculin_20_01_256,Cytoplasm|Texture_SumEntropy_Vinculin_5_01_256,Cytoplasm|Textu
re_SumEntropy_Vinculin_5_00_256,Cytoplasm|Texture_SumEntropy_Vinculin_5_02_256,Cytoplasm|Texture_SumEn
tropy_Vinculin_5_03_256,Cytoplasm|Texture_SumEntropy_DNA_3_00_256,Cytoplasm|Texture_SumEntropy_DNA_
3_03_256,Cytoplasm|Texture_SumEntropy_DNA_3_02_256,Cytoplasm|Texture_SumEntropy_DNA_3_01_256,Cytopl
asm|Texture_SumEntropy_DNA_20_02_256,Cytoplasm|Texture_SumEntropy_DNA_20_03_256,Cytoplasm|Texture_S
umEntropy_DNA_20_00_256,Cytoplasm|Texture_SumEntropy_DNA_20_01_256,Cytoplasm|Texture_SumEntropy_D
NA_5_02_256,Cytoplasm|Texture_SumEntropy_DNA_5_01_256,Cytoplasm|Texture_SumEntropy_DNA_5_00_256,C
ytoplasm|Texture_SumEntropy_DNA_5_03_256,Cytoplasm|Texture_SumEntropy_DNA_10_03_256,Cytoplasm|Textur
e_SumEntropy_DNA_10_02_256,Cytoplasm|Texture_SumEntropy_DNA_10_01_256,Cytoplasm|Texture_SumEntropy
_DNA_10_00_256,Cytoplasm|Texture_SumEntropy_WGA_10_02_256,Cytoplasm|Texture_SumEntropy_WGA_10_0
0_256,Cytoplasm|Texture_SumEntropy_WGA_10_01_256,Cytoplasm|Texture_SumEntropy_WGA_10_03_256,Cytopl
asm|Texture_SumEntropy_WGA_5_00_256,Cytoplasm|Texture_SumEntropy_WGA_5_03_256,Cytoplasm|Texture_Su
mEntropy_WGA_5_01_256,Cytoplasm|Texture_SumEntropy_WGA_5_02_256,Cytoplasm|Texture_SumEntropy_WG
A_3_03_256,Cytoplasm|Texture_SumEntropy_WGA_3_00_256,Cytoplasm|Texture_SumEntropy_WGA_3_02_256,C
ytoplasm|Texture_SumEntropy_WGA_3_01_256,Cytoplasm|Texture_SumEntropy_WGA_20_01_256,Cytoplasm|Text
ure_SumEntropy_WGA_20_02_256,Cytoplasm|Texture_SumEntropy_WGA_20_00_256,Cytoplasm|Texture_SumEntr
opy_WGA_20_03_256,Cytoplasm|Intensity_IntegratedIntensityEdge_Phalloidin,Cytoplasm|Intensity_IntegratedIntensit
yEdge_DNA,Cytoplasm|Intensity_IntegratedIntensityEdge_Vinculin,Cytoplasm|Intensity_IntegratedIntensityEdge_WG
A,Cytoplasm|Intensity_MaxIntensityEdge_Vinculin,Cytoplasm|Intensity_MaxIntensityEdge_DNA,Cytoplasm|Intensity
_MaxIntensityEdge_WGA,Cytoplasm|Intensity_MaxIntensityEdge_Phalloidin,Cytoplasm|Intensity_MedianIntensity_P
halloidin,Cytoplasm|Intensity_MedianIntensity_Vinculin,Cytoplasm|Intensity_MedianIntensity_DNA,Cytoplasm|Intens
ity_MedianIntensity_WGA,Cytoplasm|Intensity_StdIntensity_Phalloidin,Cytoplasm|Intensity_StdIntensity_WGA,Cyto
plasm|Intensity_StdIntensity_Vinculin,Cytoplasm|Intensity_StdIntensity_DNA,Cytoplasm|Intensity_UpperQuartileInte
nsity_Phalloidin,Cytoplasm|Intensity_UpperQuartileIntensity_DNA,Cytoplasm|Intensity_UpperQuartileIntensity_WGA
,Cytoplasm|Intensity_UpperQuartileIntensity_Vinculin,Cytoplasm|Intensity_MADIntensity_WGA,Cytoplasm|Intensity
_MADIntensity_Vinculin,Cytoplasm|Intensity_MADIntensity_Phalloidin,Cytoplasm|Intensity_MADIntensity_DNA,Cy
toplasm|Intensity_MaxIntensity_Phalloidin,Cytoplasm|Intensity_MaxIntensity_Vinculin,Cytoplasm|Intensity_MaxInten
sity_WGA,Cytoplasm|Intensity_MaxIntensity_DNA,Cytoplasm|Intensity_IntegratedIntensity_Phalloidin,Cytoplasm|Int

ensity_IntegratedIntensity_DNA,Cytoplasm|Intensity_IntegratedIntensity_WGA,Cytoplasm|Intensity_IntegratedIntensit
y_Vinculin,Cytoplasm|Intensity_MassDisplacement_WGA,Cytoplasm|Intensity_MassDisplacement_Phalloidin,Cytopla
sm|Intensity_MassDisplacement_Vinculin,Cytoplasm|Intensity_MassDisplacement_DNA,Cytoplasm|Intensity_StdInten
sityEdge_DNA,Cytoplasm|Intensity_StdIntensityEdge_Vinculin,Cytoplasm|Intensity_StdIntensityEdge_Phalloidin,Cyt
oplasm|Intensity_StdIntensityEdge_WGA,Cytoplasm|Intensity_MinIntensity_WGA,Cytoplasm|Intensity_MinIntensity_
Phalloidin,Cytoplasm|Intensity_MinIntensity_DNA,Cytoplasm|Intensity_MinIntensity_Vinculin,Cytoplasm|Intensity_
MinIntensityEdge_Vinculin,Cytoplasm|Intensity_MinIntensityEdge_Phalloidin,Cytoplasm|Intensity_MinIntensityEdge
_WGA,Cytoplasm|Intensity_MinIntensityEdge_DNA,Cytoplasm|Intensity_LowerQuartileIntensity_Phalloidin,Cytoplas
m|Intensity_LowerQuartileIntensity_DNA,Cytoplasm|Intensity_LowerQuartileIntensity_Vinculin,Cytoplasm|Intensity_
LowerQuartileIntensity_WGA,Cytoplasm|Intensity_MeanIntensity_DNA,Cytoplasm|Intensity_MeanIntensity_Phalloidi
n,Cytoplasm|Intensity_MeanIntensity_WGA,Cytoplasm|Intensity_MeanIntensity_Vinculin,Cytoplasm|Intensity_MeanI
ntensityEdge_WGA,Cytoplasm|Intensity_MeanIntensityEdge_DNA,Cytoplasm|Intensity_MeanIntensityEdge_Phalloidi
n,Cytoplasm|Intensity_MeanIntensityEdge_Vinculin,Cytoplasm|Granularity_12_WGA,Cytoplasm|Granularity_12_Phal
loidin,Cytoplasm|Granularity_12_DNA,Cytoplasm|Granularity_12_Vinculin,Cytoplasm|Granularity_15_Phalloidin,Cyt
oplasm|Granularity_15_WGA,Cytoplasm|Granularity_15_Vinculin,Cytoplasm|Granularity_15_DNA,Cytoplasm|Granul
arity_16_Vinculin,Cytoplasm|Granularity_16_Phalloidin,Cytoplasm|Granularity_16_DNA,Cytoplasm|Granularity_16_
WGA,Cytoplasm|Granularity_4_Phalloidin,Cytoplasm|Granularity_4_WGA,Cytoplasm|Granularity_4_Vinculin,Cytopl
asm|Granularity_4_DNA,Cytoplasm|Granularity_14_DNA,Cytoplasm|Granularity_14_Phalloidin,Cytoplasm|Granularit
y_14_WGA,Cytoplasm|Granularity_14_Vinculin,Cytoplasm|Granularity_2_DNA,Cytoplasm|Granularity_2_WGA,Cyt
oplasm|Granularity_2_Vinculin,Cytoplasm|Granularity_2_Phalloidin,Cytoplasm|Granularity_10_DNA,Cytoplasm|Gran
ularity_10_Vinculin,Cytoplasm|Granularity_10_Phalloidin,Cytoplasm|Granularity_10_WGA,Cytoplasm|Granularity_7_
DNA,Cytoplasm|Granularity_7_WGA,Cytoplasm|Granularity_7_Vinculin,Cytoplasm|Granularity_7_Phalloidin,Cytopla
sm|Granularity_9_WGA,Cytoplasm|Granularity_9_Phalloidin,Cytoplasm|Granularity_9_Vinculin,Cytoplasm|Granularit
y_9_DNA,Cytoplasm|Granularity_11_Phalloidin,Cytoplasm|Granularity_11_Vinculin,Cytoplasm|Granularity_11_DNA
,Cytoplasm|Granularity_11_WGA,Cytoplasm|Granularity_1_Phalloidin,Cytoplasm|Granularity_1_DNA,Cytoplasm|Gra
nularity_1_Vinculin,Cytoplasm|Granularity_1_WGA,Cytoplasm|Granularity_3_WGA,Cytoplasm|Granularity_3_DNA,
Cytoplasm|Granularity_3_Vinculin,Cytoplasm|Granularity_3_Phalloidin,Cytoplasm|Granularity_6_Phalloidin,Cytoplas
m|Granularity_6_DNA,Cytoplasm|Granularity_6_Vinculin,Cytoplasm|Granularity_6_WGA,Cytoplasm|Granularity_5_
Phalloidin,Cytoplasm|Granularity_5_WGA,Cytoplasm|Granularity_5_Vinculin,Cytoplasm|Granularity_5_DNA,Cytopla
sm|Granularity_8_Phalloidin,Cytoplasm|Granularity_8_WGA,Cytoplasm|Granularity_8_DNA,Cytoplasm|Granularity_
8_Vinculin,Cytoplasm|Granularity_13_DNA,Cytoplasm|Granularity_13_WGA,Cytoplasm|Granularity_13_Phalloidin,C
ytoplasm|Granularity_13_Vinculin,Cytoplasm|AreaShape_BoundingBoxMaximum_X,Cytoplasm|AreaShape_Bounding
BoxMaximum_Y,Cytoplasm|AreaShape_Zernike_8_0,Cytoplasm|AreaShape_Zernike_8_6,Cytoplasm|AreaShape_Zern
ike_8_2,Cytoplasm|AreaShape_Zernike_8_4,Cytoplasm|AreaShape_Zernike_8_8,Cytoplasm|AreaShape_Zernike_7_1,
Cytoplasm|AreaShape_Zernike_7_3,Cytoplasm|AreaShape_Zernike_7_7,Cytoplasm|AreaShape_Zernike_7_5,Cytoplas
m|AreaShape_Zernike_3_1,Cytoplasm|AreaShape_Zernike_3_3,Cytoplasm|AreaShape_Zernike_4_2,Cytoplasm|AreaS
hape_Zernike_4_4,Cytoplasm|AreaShape_Zernike_4_0,Cytoplasm|AreaShape_Zernike_6_4,Cytoplasm|AreaShape_Zer
nike_6_6,Cytoplasm|AreaShape_Zernike_6_2,Cytoplasm|AreaShape_Zernike_6_0,Cytoplasm|AreaShape_Zernike_9_9,
Cytoplasm|AreaShape_Zernike_9_1,Cytoplasm|AreaShape_Zernike_9_7,Cytoplasm|AreaShape_Zernike_9_5,Cytoplas
m|AreaShape_Zernike_9_3,Cytoplasm|AreaShape_Zernike_2_0,Cytoplasm|AreaShape_Zernike_2_2,Cytoplasm|AreaS
hape_Zernike_5_3,Cytoplasm|AreaShape_Zernike_5_1,Cytoplasm|AreaShape_Zernike_5_5,Cytoplasm|AreaShape_Zer
nike_1_1,Cytoplasm|AreaShape_Zernike_0_0,Cytoplasm|AreaShape_Center_X,Cytoplasm|AreaShape_Center_Y,Cytop
lasm|AreaShape_MaximumRadius,Cytoplasm|AreaShape_MeanRadius,Cytoplasm|AreaShape_Solidity,Cytoplasm|Area
Shape_Orientation,Cytoplasm|AreaShape_BoundingBoxArea,Cytoplasm|AreaShape_Eccentricity,Cytoplasm|AreaShap
e_ConvexArea,Cytoplasm|AreaShape_Area,Cytoplasm|AreaShape_Extent,Cytoplasm|AreaShape_EquivalentDiameter,
Cytoplasm|AreaShape_MinFeretDiameter,Cytoplasm|AreaShape_Compactness,Cytoplasm|AreaShape_MajorAxisLengt
h,Cytoplasm|AreaShape_FormFactor,Cytoplasm|AreaShape_Perimeter,Cytoplasm|AreaShape_MaxFeretDiameter,Cyto
plasm|AreaShape_MedianRadius,Cytoplasm|AreaShape_BoundingBoxMinimum_Y,Cytoplasm|AreaShape_BoundingB
oxMinimum_X,Cytoplasm|AreaShape_MinorAxisLength,Cytoplasm|AreaShape_EulerNumber,Cytoplasm|RadialDistri
bution_RadialCV_Phalloidin_2of4,Cytoplasm|RadialDistribution_RadialCV_Phalloidin_4of4,Cytoplasm|RadialDistrib
ution_RadialCV_Phalloidin_3of4,Cytoplasm|RadialDistribution_RadialCV_Phalloidin_1of4,Cytoplasm|RadialDistribut
ion_RadialCV_DNA_1of4,Cytoplasm|RadialDistribution_RadialCV_DNA_3of4,Cytoplasm|RadialDistribution_Radial
CV_DNA_4of4,Cytoplasm|RadialDistribution_RadialCV_DNA_2of4,Cytoplasm|RadialDistribution_RadialCV_Vincul
in_2of4,Cytoplasm|RadialDistribution_RadialCV_Vinculin_3of4,Cytoplasm|RadialDistribution_RadialCV_Vinculin_4

of4,Cytoplasm|RadialDistribution_RadialCV_Vinculin_1of4,Cytoplasm|RadialDistribution_RadialCV_WGA_1of4,Cyt
oplasm|RadialDistribution_RadialCV_WGA_2of4,Cytoplasm|RadialDistribution_RadialCV_WGA_3of4,Cytoplasm|Ra
dialDistribution_RadialCV_WGA_4of4,Cytoplasm|RadialDistribution_MeanFrac_Vinculin_2of4,Cytoplasm|RadialDist
ribution_MeanFrac_Vinculin_1of4,Cytoplasm|RadialDistribution_MeanFrac_Vinculin_3of4,Cytoplasm|RadialDistribut
ion_MeanFrac_Vinculin_4of4,Cytoplasm|RadialDistribution_MeanFrac_DNA_3of4,Cytoplasm|RadialDistribution_Me
anFrac_DNA_1of4,Cytoplasm|RadialDistribution_MeanFrac_DNA_2of4,Cytoplasm|RadialDistribution_MeanFrac_DN
A_4of4,Cytoplasm|RadialDistribution_MeanFrac_WGA_3of4,Cytoplasm|RadialDistribution_MeanFrac_WGA_1of4,C
ytoplasm|RadialDistribution_MeanFrac_WGA_4of4,Cytoplasm|RadialDistribution_MeanFrac_WGA_2of4,Cytoplasm|
RadialDistribution_MeanFrac_Phalloidin_4of4,Cytoplasm|RadialDistribution_MeanFrac_Phalloidin_2of4,Cytoplasm|R
adialDistribution_MeanFrac_Phalloidin_3of4,Cytoplasm|RadialDistribution_MeanFrac_Phalloidin_1of4,Cytoplasm|Ra
dialDistribution_FracAtD_DNA_4of4,Cytoplasm|RadialDistribution_FracAtD_DNA_2of4,Cytoplasm|RadialDistributio
n_FracAtD_DNA_3of4,Cytoplasm|RadialDistribution_FracAtD_DNA_1of4,Cytoplasm|RadialDistribution_FracAtD_V
inculin_3of4,Cytoplasm|RadialDistribution_FracAtD_Vinculin_4of4,Cytoplasm|RadialDistribution_FracAtD_Vinculin
_2of4,Cytoplasm|RadialDistribution_FracAtD_Vinculin_1of4,Cytoplasm|RadialDistribution_FracAtD_WGA_1of4,Cyt
oplasm|RadialDistribution_FracAtD_WGA_3of4,Cytoplasm|RadialDistribution_FracAtD_WGA_2of4,Cytoplasm|Radi
alDistribution_FracAtD_WGA_4of4,Cytoplasm|RadialDistribution_FracAtD_Phalloidin_3of4,Cytoplasm|RadialDistrib
ution_FracAtD_Phalloidin_4of4,Cytoplasm|RadialDistribution_FracAtD_Phalloidin_2of4,Cytoplasm|RadialDistributio
n_FracAtD_Phalloidin_1of4,Cytoplasm|Correlation_Correlation_DNA_Phalloidin,Cytoplasm|Correlation_Correlation_
DNA_WGA,Cytoplasm|Correlation_Correlation_DNA_Vinculin,Cytoplasm|Correlation_Correlation_Vinculin_WGA,C
ytoplasm|Correlation_Correlation_Phalloidin_Vinculin,Cytoplasm|Correlation_Correlation_Phalloidin_WGA,Cytoplas
m|Correlation_K_DNA_Phalloidin,Cytoplasm|Correlation_K_DNA_WGA,Cytoplasm|Correlation_K_DNA_Vinculin,C
ytoplasm|Correlation_K_WGA_Vinculin,Cytoplasm|Correlation_K_WGA_Phalloidin,Cytoplasm|Correlation_K_WGA
_DNA,Cytoplasm|Correlation_K_Vinculin_WGA,Cytoplasm|Correlation_K_Vinculin_DNA,Cytoplasm|Correlation_K
_Vinculin_Phalloidin,Cytoplasm|Correlation_K_Phalloidin_Vinculin,Cytoplasm|Correlation_K_Phalloidin_WGA,Cyto
plasm|Correlation_K_Phalloidin_DNA,Cytoplasm|Correlation_RWC_Phalloidin_Vinculin,Cytoplasm|Correlation_RW
C_Phalloidin_DNA,Cytoplasm|Correlation_RWC_Phalloidin_WGA,Cytoplasm|Correlation_RWC_WGA_Phalloidin,C
ytoplasm|Correlation_RWC_WGA_DNA,Cytoplasm|Correlation_RWC_WGA_Vinculin,Cytoplasm|Correlation_RWC
_Vinculin_Phalloidin,Cytoplasm|Correlation_RWC_Vinculin_DNA,Cytoplasm|Correlation_RWC_Vinculin_WGA,Cyt
oplasm|Correlation_RWC_DNA_Phalloidin,Cytoplasm|Correlation_RWC_DNA_WGA,Cytoplasm|Correlation_RWC_
DNA_Vinculin,Cytoplasm|Correlation_Manders_Phalloidin_DNA,Cytoplasm|Correlation_Manders_Phalloidin_Vinculi
n,Cytoplasm|Correlation_Manders_Phalloidin_WGA,Cytoplasm|Correlation_Manders_WGA_DNA,Cytoplasm|Correlat
ion_Manders_WGA_Vinculin,Cytoplasm|Correlation_Manders_WGA_Phalloidin,Cytoplasm|Correlation_Manders_Vi
nculin_WGA,Cytoplasm|Correlation_Manders_Vinculin_Phalloidin,Cytoplasm|Correlation_Manders_Vinculin_DNA,C
ytoplasm|Correlation_Manders_DNA_Vinculin,Cytoplasm|Correlation_Manders_DNA_WGA,Cytoplasm|Correlation_
Manders_DNA_Phalloidin,Cytoplasm|Correlation_Overlap_DNA_WGA,Cytoplasm|Correlation_Overlap_DNA_Phallo
idin,Cytoplasm|Correlation_Overlap_DNA_Vinculin,Cytoplasm|Correlation_Overlap_Vinculin_WGA,Cytoplasm|Corre
lation_Overlap_Phalloidin_Vinculin,Cytoplasm|Correlation_Overlap_Phalloidin_WGA,Cytoplasm|Location_CenterMa
ssIntensity_Y_DNA,Cytoplasm|Location_CenterMassIntensity_Y_WGA,Cytoplasm|Location_CenterMassIntensity_Y_
Vinculin,Cytoplasm|Location_CenterMassIntensity_Y_Phalloidin,Cytoplasm|Location_CenterMassIntensity_X_WGA,
Cytoplasm|Location_CenterMassIntensity_X_DNA,Cytoplasm|Location_CenterMassIntensity_X_Phalloidin,Cytoplasm
|Location_CenterMassIntensity_X_Vinculin,Cytoplasm|Location_CenterMassIntensity_Z_Phalloidin,Cytoplasm|Locati
on_CenterMassIntensity_Z_Vinculin,Cytoplasm|Location_CenterMassIntensity_Z_DNA,Cytoplasm|Location_CenterM
assIntensity_Z_WGA,Cytoplasm|Location_MaxIntensity_X_Phalloidin,Cytoplasm|Location_MaxIntensity_X_Vinculin
,Cytoplasm|Location_MaxIntensity_X_DNA,Cytoplasm|Location_MaxIntensity_X_WGA,Cytoplasm|Location_MaxInt
ensity_Z_DNA,Cytoplasm|Location_MaxIntensity_Z_WGA,Cytoplasm|Location_MaxIntensity_Z_Vinculin,Cytoplasm
|Location_MaxIntensity_Z_Phalloidin,Cytoplasm|Location_MaxIntensity_Y_Vinculin,Cytoplasm|Location_MaxIntensi
ty_Y_WGA,Cytoplasm|Location_MaxIntensity_Y_Phalloidin,Cytoplasm|Location_MaxIntensity_Y_DNA,Cytoplasm|
Location_Center_X,Cytoplasm|Location_Center_Y,Cytoplasm|Parent_Cells,Cytoplasm|Parent_Nuclei,Cytoplasm|Num
ber_Object_Number,Experiment|Run_Timestamp,Experiment|ImageQuality_ThresholdStdOtsu_OrigDNA_2W,Experi
ment|ImageQuality_ThresholdStdOtsu_OrigVinculin_3FW,Experiment|ImageQuality_ThresholdMedianOtsu_OrigDN
A_2W,Experiment|ImageQuality_ThresholdMedianOtsu_OrigVinculin_3FW,Experiment|ImageQuality_ThresholdMea
nOtsu_OrigVinculin_3FW,Experiment|ImageQuality_ThresholdMeanOtsu_OrigDNA_2W,Experiment|Modification_Ti
mestamp,Experiment|CellProfiler_Version,Experiment|Pipeline_Pipeline
Representation of Nan/Inf:NaN

Add a prefix to file names?:No
Filename prefix:MyExpt_
Overwrite existing files without warning?:No
Data to export:Image
Combine these object measurements with those of the previous object?:No
File name:DATA.csv
Use the object name for the file name?:Yes
Data to export:Experiment
Combine these object measurements with those of the previous object?:No
File name:DATA.csv
Use the object name for the file name?:Yes
Data to export:TransfectedCells
Combine these object measurements with those of the previous object?:No
File name:Cells.csv
Use the object name for the file name?:No
Data to export:TransfectedCytoplasm
Combine these object measurements with those of the previous object?:No
File name:Cytoplasm.csv
Use the object name for the file name?:No
Data to export:TransfectedNuclei
Combine these object measurements with those of the previous object?:No
File name:Nuclei.csv
Use the object name for the file name?:No
Data to export:UntransfectedCells
Combine these object measurements with those of the previous object?:No
File name:DATA.csv
Use the object name for the file name?:Yes
Data to export:UntransfectedCytoplasm
Combine these object measurements with those of the previous object?:No
File name:DATA.csv
Use the object name for the file name?:Yes
Data to export:UntransfectedNuclei
Combine these object measurements with those of the previous object?:No
File name:DATA.csv
Use the object name for the file name?:Yes
